# MAP2 is Differentially Phosphorylated in Schizophrenia, Altering its Function

**DOI:** 10.1101/683912

**Authors:** MJ Grubisha, X Sun, ML MacDonald, M Garver, Z Sun, KA Paris, DS Patel, RA DeGiosio, DA Lewis, NA Yates, C Camacho, GE Homanics, Y Ding, RA Sweet

## Abstract

Schizophrenia (Sz) is a highly polygenic disorder, with common, rare, and structural variants each contributing only a small fraction of overall disease risk. Thus, there is a need to identify downstream points of convergence that can be targeted with therapeutics. Reduction of Microtubule-associated Protein 2 (MAP2) immunoreactivity (MAP2-IR) is present in individuals with Sz, despite no change in MAP2 protein levels. MAP2 is phosphorylated downstream of multiple receptors and kinases identified as Sz risk genes, altering its immunoreactivity and function. Using an unbiased phosphoproteomics approach we quantified 18 MAP2 phosphopeptides, 9 of which were significantly altered in Sz subjects. Network analysis grouped MAP2 phosphopeptides into 3 modules, each with a distinct relationship to dendritic spine loss, synaptic protein levels, and clinical function in Sz subjects. We then investigated the most hyperphosphorylated site in Sz, phosphoserine1782 (pS1782). Computational modeling predicted phosphorylation of S1782 reduces binding of MAP2 to microtubules, which was confirmed experimentally. We generated a transgenic mouse containing a phosphomimetic mutation at S1782 (S1782E) and found reductions in basilar dendritic length and complexity along with reduced spine density. Because only a limited number of MAP2 interacting proteins have been previously identified, we combined co-immunoprecipitation with mass spectrometry to characterize the MAP2 interactome in mouse brain. The MAP2 interactome was enriched for proteins involved in protein translation. These associations were shown to be functional as overexpression of wildtype and phosphomimetic MAP2 reduced protein synthesis *in vitro*. Finally, we found that Sz subjects with low MAP2-IR had reductions in the levels of synaptic proteins relative to nonpsychiatric control (NPC) subjects and to Sz subjects with normal and MAP2-IR, and this same pattern was recapitulated in S1782E mice. These findings suggest a new conceptual framework for Sz - that a large proportion of individuals have a “MAP2opathy” - in which MAP function is altered by phosphorylation, leading to impairments of neuronal structure, synaptic protein synthesis, and function.

## Introduction

Schizophrenia (Sz) is highly polygenic, with common, rare, and structural variants each contributing only a small fraction of overall disease risk^1^. Thus, there is a need to identify points of convergence, i.e. pathologies shared by large proportions of individuals with Sz, that can be therapeutically targeted to halt or reverse the synaptic^2^ and functional deterioration^3,4^ that occurs in affected individuals. Reduced Microtubule-Associated Protein 2 (MAP2) immunoreactivity (IR) has been reported in multiple cortical regions in Sz^5–10^, leading one recent author to characterize it as a hallmark pathology in Sz^11^. Reduced MAP2-IR was not a consequence of common technical and clinical confounds of postmortem studies, suggesting it reflects an underlying pathogenic process ^10,12^.

In ~60% of individuals with Sz, the reduction in MAP2-IR is profound ^10,12,13^ Even in these cases in which MAP2-IR is markedly reduced, it is not due to loss of MAP2 protein itself^10^, and thus must reflect reduced detectability of MAP2 by antibody. Phosphorylation of MAP2 is known to alter its structure, function, and immunoreactivity (IR)^14^, and to be regulated downstream of multiple genes now established by unbiased methods as Sz risk genes^15–23^. As the predominant regulator of the dendritic microtubule cytoskeleton, as well as a noted coordinator between microtubule and actin dynamics^24^, alterations to MAP2 function could explain a variety of abnormalities in dendritic structure which have been observed in Sz. Many groups have reported reduced dendritic arborization^25–29^, dendritic spine density ^10,27,28,30–33^, and dendritic spine number^33^ to be present in multiple cortical regions in Sz. These include our studies of primary auditory cortex^10,32,34^.

Thus, we hypothesized that MAP2 phosphorylation would be altered in Sz and alter MAP2 functions, providing a potential hub for pathogenesis. To evaluate this, we undertook a phosphoproteomic analysis of MAP2 in primary auditory cortex of Sz and nonpsychiatric control (NPC) subjects. We identified 18 unique phosphopeptides. Levels of 9 of these were significantly altered in Sz. The 18 phosphopeptides grouped into 3 modules using network analysis, each with a distinct relationship to clinicopathologic correlates of disease. We designed a series of experiments to further investigate the most altered phosphorylation site, serine 1782 (pS1782), finding that phosphomimetic mutation at this site reduces microtuble binding. In a phosphomimetic knock-in mouse model of pS1782 (S1782E), we found reduced dendritic length, complexity, and spine density. Using MAP2 co-immunoprecipitation (co-IP) coupled with mass spectrometry, we found that the MAP2 interactome was enriched for proteins regulating translation. This association was shown to be functional as overexpression of wildtype and phosphomimetic MAP2 reduced protein synthesis *in vitro*. Finally, we found that Sz with profound reductions in MAP2-IR (MAP2-IR_LOW_) was characterized by reductions in the levels of synaptic proteins relative to both NPC and SZ with normal MAP2-IR (MAP2-IR_NL_) subjects, and this same pattern was recapitulated in S1782E mice, although to a lesser degree.

## Results

### MAP2 is differentially phosphorylated in auditory cortex in Sz

We conducted an unbiased proteomic survey of MAP2 phosphorylation in tissue obtained from right Hechl’s gyrus (primary auditory cortex (A1)) obtained from Sz subjects with either profoundly reduced MAP2-IR in left Heschl’s gyrus (MAP2-IR_LOW_, N=11)^10^ or normal MAP2-IR (MAP2-IR_NL_, N=5), and from non-psychiatric control (NPC, N=16) subjects (see Fig 1a and Table S1 for subject characteristics). We identified 18 unique MAP2 phosphopeptides in our samples (Table S2), confirmed by manual sequencing of MS2 spectra, 8 of which were significantly altered (q<0.05) in MAP2-IR_LOW_ Sz subjects relative to NPC subjects (Table S2). In MAP2-IR_NL_ Sz subjects, changes in phosphopeptide levels were mostly attenuated, although in the combined Sz group a total of 9 phosphopeptides now reached significance (Figure 1b upper panel, Table S2). The most altered phosphopeptide, VDHGAEIITQS[+80]PGR, containing a single phosphorylation at serine 1782 (pS1782, numbering per the canonical MAP2b isoform, Uniprot identifier: P11137-1), was increased up to nearly 7 fold relative to NPC subjects. Elevated levels of MAP2 phosphopeptides were largely unassociated with potential tissue and clinical confounds (Table S3 and Figure S4) with two notable exceptions. First, levels of VDHGAEIITQS[+80]PGR were significantly lower in Sz subjects taking antipsychotics at time of death than in those subjects off antipsychotics, raising the possibility that antipsychotic treatment could act, in part, through effects on phosphorylation of S1782. However, this effect was not recapitulated by a monkey model of longterm antipsychotic exposure^35^, nor did this model induce elevations in any of the phosphopeptides found to be elevated in Sz in our current study (Fig 1a, lower panel). Second, phosphopeptide abundance may be affected by postmortem interval (PMI). Thus, we further evaluated this potential confound in 2 ways. We first tested the effect of PMI on phosphopeptide abundance in our subjects, finding no significant associations (all q>0.1, Table S3). Second, we surveyed MAP2 phosphopeptide abundance in a mouse model of increasing PMI. We detected 9 MAP2 phosphopeptides in this model. Fig 1c shows the detected MAP2 phosphopeptides, 8 of which demonstrate stable or linear changes over time (and thus readily accounted for by PMI matching in our subjects). The exception was peptide (RL*S[+80]*NVS*S[+80]*SG*S[+80]*INLLESPQLATLAEDVTAALAK) which had a nonlinear decay.

**Fig 1.**
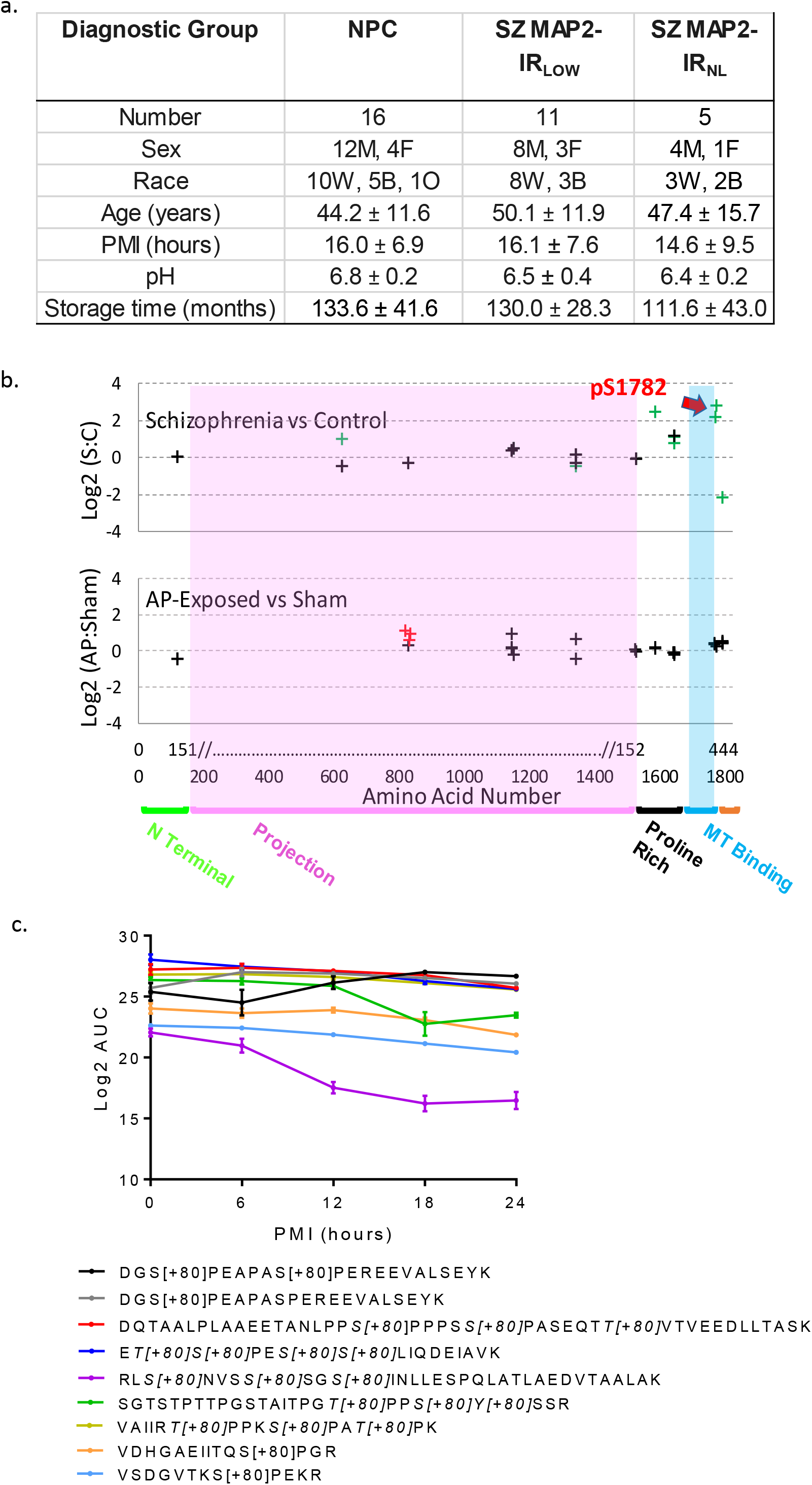
MAP2 is differentially phosphorylated in Sz. (a) Data for continuous variables are presented as group average ± standard deviation for each group (NPC = nonpsychiatric control, SZ MAP2-IR_LOW_, SZ MAP2-IR_NL_). There was no significant difference amongst groups on any of the variables (all p>0.1). (b) Top panel shows Sz vs NPC, phosphopeptides are denoted by crosses at the position of their starting amino acid. Nine phosphopeptides had FDR significant alterations (q<0.05, green crosses). Note significantly altered phosphopeptides were present in the Projection domain (unique to high MW MAP2A/B) and surrounding the MT binding domain. Red arrow denotes pS1782. No phosphopeptides were detected in the MT binding domain. Bottom panel shows antipsychotic-exposed (AP) vs sham-exposed monkey. Three peptides were nominally significant (p<0.05, red crosses). Most peptides were identified in both species (indicated by alignment on x-axis), but peptides significantly altered in Sz did not correspond to those increased by antipsychotic exposure. X-axis shows amino acid location in MAP2C and in canonical MAP2B. MAP2 functional domains are denoted. (c) Log2 area under the curve (AUC) values for significant MAP2 phosphopeptides across increasing postmortem intervals in mouse tissue demonstrates relatively linear stability that is easily accounted for in our PMI matching of subjects. The one exception to these linear trends is RLS[+80]NVSS[+80]SGS[+80]INLLESPQLATLAEDVTAALAK. Individual peptides indicating phosphorylated residue(s) are listed in the legend below the graph. Data shown are ±SEM from N=4 samples per timepoint.

We next applied Weighted Gene Co-Expression Network Analysis (WGCNA)^36^ to the MAP2 phosphopeptide data from the Sz and NPC subjects. MAP2 phosphopeptides clustered into three modules (Fig S1). We then used these three modules to compute eigenpeptide values for all subjects and tested these values for their associations with known pathological features of disease, namely decreased dendritic spine density and decreased synaptic protein levels, both of which had been previously measured in these subjects^37,38^. Furthermore, socioeconomic status (SES) was included as a proxy measure for functional impairment in Sz^39^, thus we also tested the eigenpeptide values for their association with subject SES (Figure 2a). Levels of phosphorylation of blue module sites, which include pS1782, are associated with lower spine density, reduced synaptic protein levels, and poorer outcomes (lower SES). In contrast, increasing phosphorylation of brown module sites, which are only present in the MAP2a and MAP2b isoforms, are correlated with better phenotypic outcomes. Turquoise module phosphopeptides show little relationship to the measured phenotypes.

**Fig 2.**
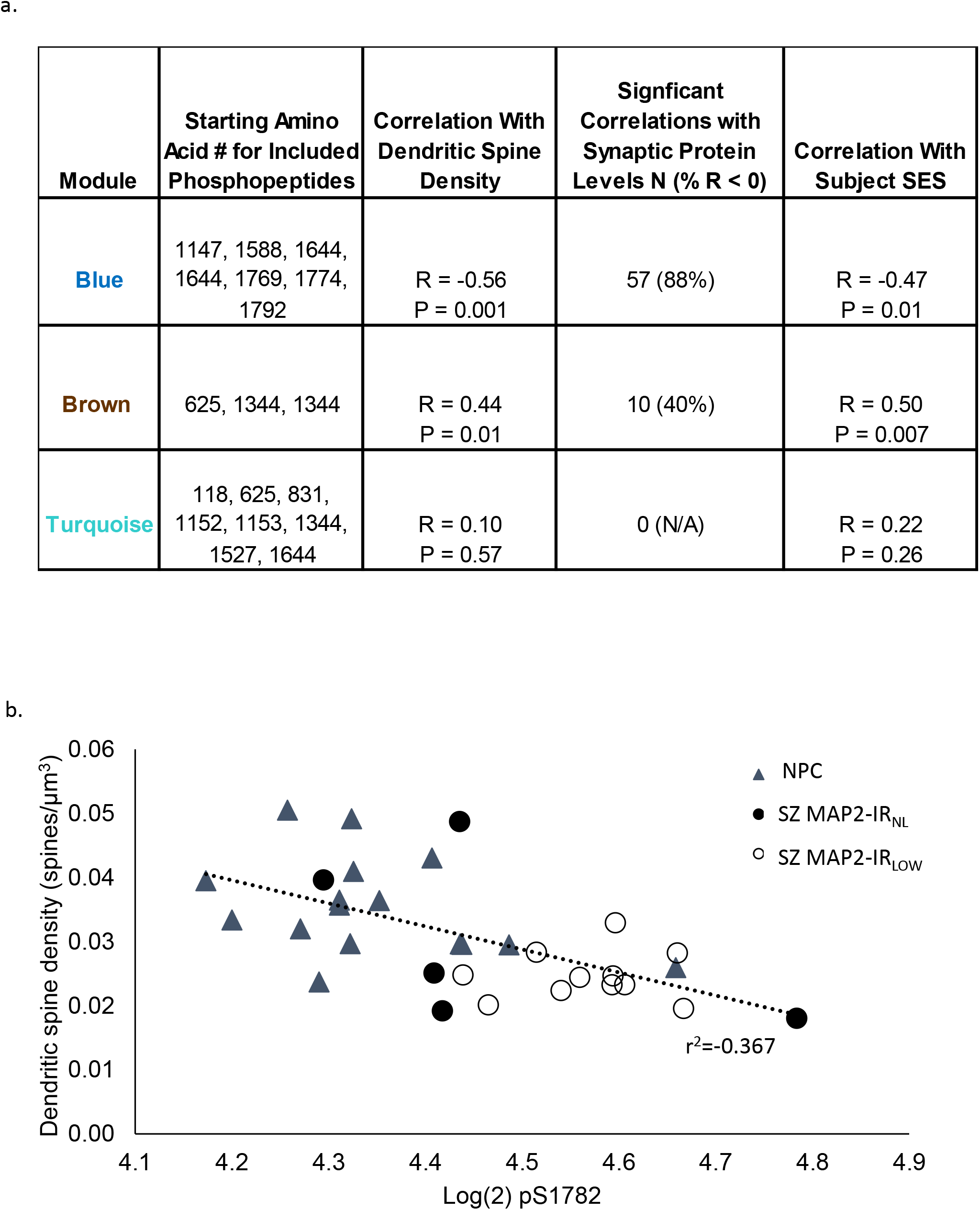
Association of phosphopeptide modules with clinical and pathologic phenotypes. (a) For each module the association with small spine density, synaptic protein levels and SES are shown. Significant correlations with synaptic protein levels were defined as q<0.05, and the percent of those significantly correlated synaptic proteins that were negatively correlated with each module is shown (% R<0). (b) pS1782 negatively correlates (r = −0.61, p = 0.0002) with dendritic spine density. NPC = nonpsychiatric control, Sz subjects grouped by MAP2 immunoreactivity as either normal (MAP2-IR_NL_) or low (MAP2-IR_LOW_)

### Phosphorylation of MAP2 at S1782 alters its predicted structure and reduces microtubule binding

The phosphopeptide with the most elevated levels in Sz was VDHGAEIITQS^[P]^PGR, phosphorylated at S1782. This peptide belongs to the blue module, which includes negative correlations with dendritic spine density and synaptic protein levels. Figure 2b shows the negative correlation (r = −0.61, p<0.001) between dendritic spine density and pS1782 levels. Furthermore, S1782 is conserved in the MAP2 homolog microtubule-associated protein tau (MAPT), as S396 (numbering by common convention as per Isoform Tau-F, Uniprot identifier: P10636-8), which when phosphorylated confers synaptic pathology^40–43^. We used all atom Molecular Dynamics simulations (MDS) to predict potential effects of pS1782 on MAP2 structure. Like its homolog Tau, structure of the intrinsically disordered MAP2 is difficult to characterize. Because MAP2 and Tau share high sequence homology, to validate our model we first checked for consistency with available information in both MAP2^44,45^ and Tau^46,47^. For this validation we relied on the known effect of phosphorylation at T231 in Tau (homologous to T1649 in MAP2). For example, our model of pT1649 MAP2 revealed a salt bridge interaction between pT1649 and R1648 and helical propensity for the region ^299^ATPKQLRLR^307^, both of which predictions recapitulated prior observations for the corresponding residues in Tau (data not shown)^48–51^. We then examined phosphorylation at S1782 by simulating a region of MAP2 containing residues 1718 to 1827 (covering the C-terminal domain). Our models indicated two structural changes upon phosphorylation of S1782: (a) an antiparallel β hairpin structure observed in wild-type (WT) MAP2 opened up to create a solvent-exposed extended β structural conformation between A1776 and S1782 in pS1782 phosphorylated MAP2; and (b) a region with helical propensity in WT MAP2 was instead found to prefer a solvent-exposed extended β structural conformation between residues V1755 and A1770 in the pS1782 ensemble (Fig 3). The homologous residues K369 to A384 in Tau are part of the C-terminal (residues K369 to S400) that flank the MT binding region and have been proposed for positioning Tau on the MT surface allowing MT binding^52–54^. More recently, NMR showed that pS396 Tau had weaker tubulin affinity than the wild type Tau peptide with signal attenuation centered around residues H374, F378, H388 and Y394^55^. Based on these data, we predicted that MAP2 pS1782 decreases MT binding.

**Fig 3.**
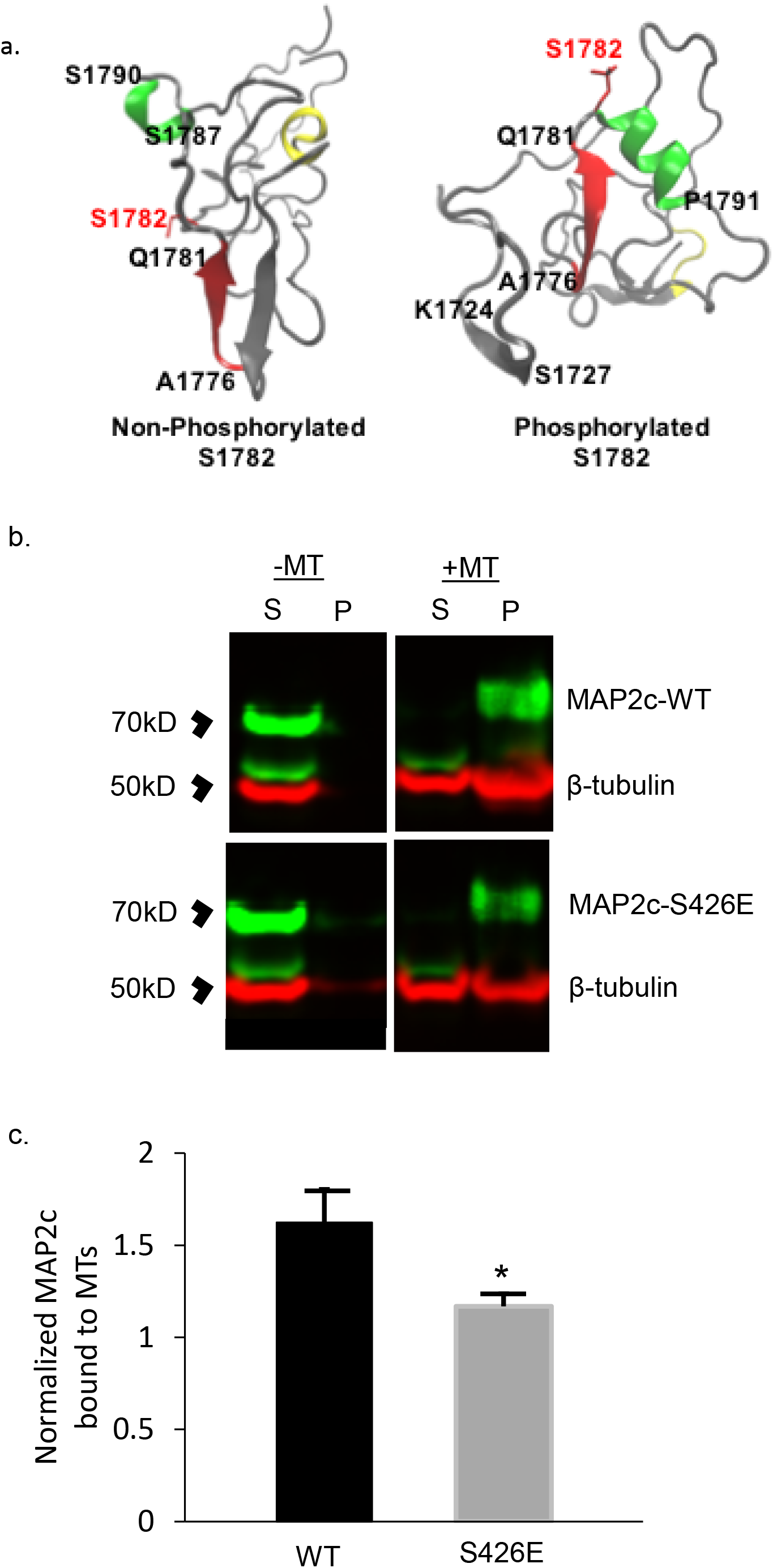
Computational modeling of MAP2 pS1782 alters predicted structure and reduces microtubule binding. (a) Snapshot of the C-terminal domain of MAP2. Amino acids A1776 to Q1781 form an extended β strand (red β strand) that is occluded by an opposing β strand. In much of the nonphosphorylated ensemble, there is found to be helical propensity in or near regions S1759 to R1765 (yellow) which is not found in the phosphorylated ensembles. In pMAP2 (pS1782) the β strand at A1776 to Q1781 is no longer occluded by another β strand; this is predicted to reduce microtubule binding and enhance interactions with other proteins. The effect of phosphorylation is recapitulated by phosphomimetic mutation S1782E (data not shown). (b) Reduced MT-binding in MAP2c with the phosphomimetic mutation S426E, homologous to S1782 in full length MAP2b relative to MAP2c-WT. 30μL of cell lysate from transiently transfected HEK293 cells was subjected to *in vitro* MT-binding assay and supernatant (S) and pellet (P) fractions were subjected to SDS-PAGE. (c) Densitometric analysis of gels shows decreased MT binding in the S426E mutant compared to WT. Data shown are from 3 independent experiments, ± SEM. *p < 0.05

To empirically evaluate these predictions, HEK293 cells, which lack endogenous MAP2, were transfected with plasmid containing either wildtype MAP2c (Uniprot identifier: P11137-2) or MAP2c with the phosphomimetic mutation S426E, homologous to S1782 in full length MAP2b (see Methods for plasmid construct information), and a microtubule (MT) binding assay was performed. The results of the MT binding assay are shown in Figure 3b&c and confirm the computationally-predicted reduction of MT binding by S426E MAP2c.

### S1782E reduces dendritic length, complexity, and spine density in vivo

Our studies in the non-neuronal HEK293 cells, while able to inform on MT binding properties of phosphomimetic MAP2, could not inform whether such effects would have an impact in the specific intracellular environment of neurons. Because MAP2 is a critical determinant of dendritic structure via its canonical function of regulating microtubule polymerization dynamics^14,56–58^, we hypothesized that the disrupted MT binding caused by pS1782 would cause reductions in dendritic architecture similar to those seen in Sz (reviewed in ^59^). To test this, we used CRISPR/Cas9 gene editing to introduce the phosphomimetic mutation (serine to glutamate, S1782E) in a mouse model (Fig S2) to allow for *in vivo* studies of dendritic structure. Reduced dendritic spine density has been consistently reported in auditory cortex in Sz^59^, thus we initially focused on layer 3 PCs in auditory cortex for dendritic reconstructions. Using Golgi staining, we compared dendritic arbor size in 12-week heterozygous S1782E+/- mice (N=10, 5 male, 5 female) with WT littermates (N=10, 5 male, 5 female). A total of 5 neurons per animal were collected. Dendritic reconstructions were manually traced from image stacks offline using NeuroLucida software (Microbrightfield, Inc) and analyzed via Sholl analysis using NeuroExplorer (Microbrightfield, Inc). There were no significant effects of sex or phase of estrus cycle, thus sexes were combined to generate N=10 animals/genotype for final analyses. We found that S1782E+/- mice had significant reductions in basal dendritic length, complexity, and spine density compared to WT (Fig 4). Despite the effects on dendritic structure, phosphomimetic mutation at S1782 did not reduce MAP2 immunoreactivity in our mouse model. We quantified MAP2-IR in a small cohort of wildtype and MAP2 S/E +/+ mice and did not find a significant difference between genotypes (mean change in MAP2 intensity 1.96 + 16%, data not shown).

**Fig 4.**
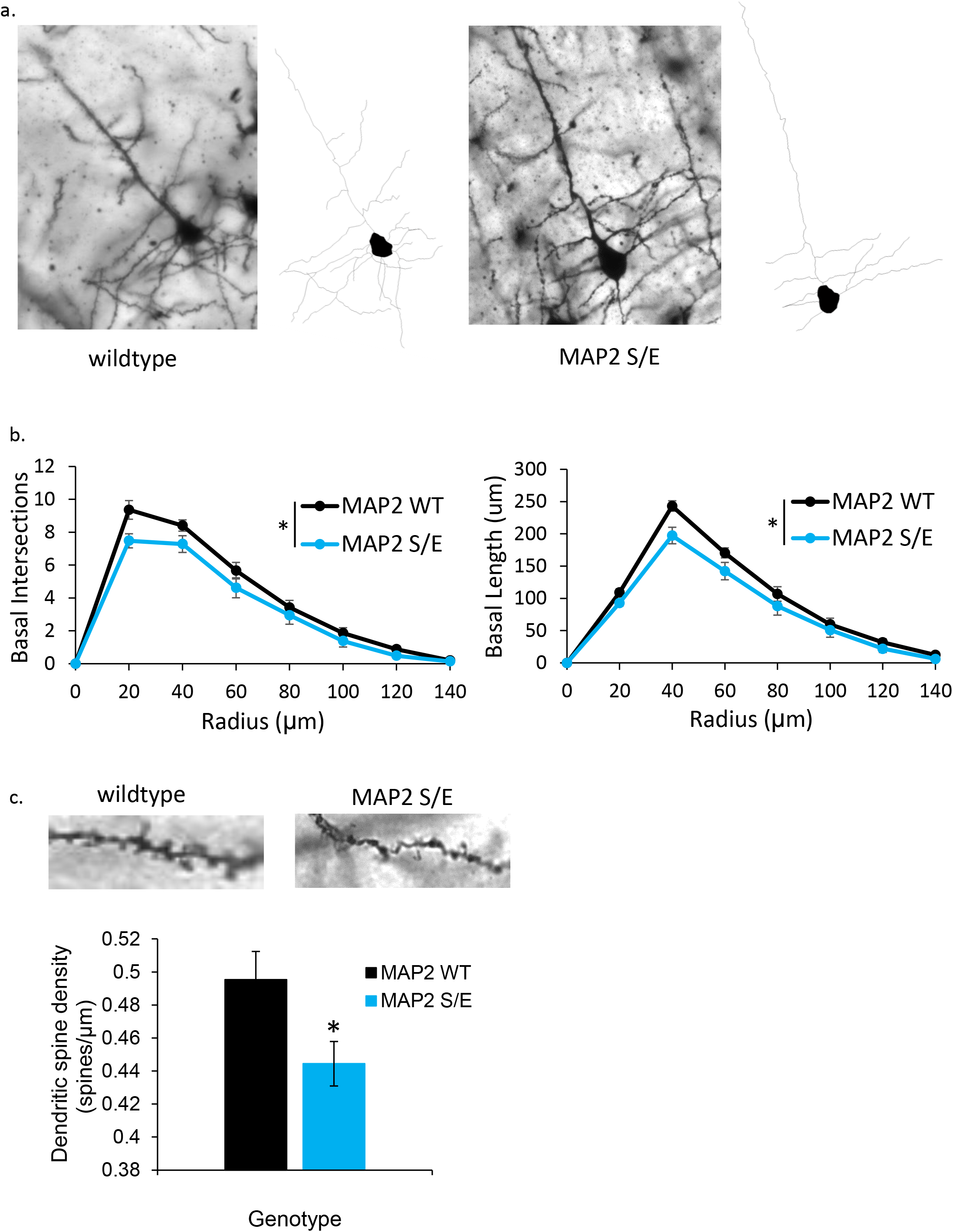
S1782E +/- mice demonstrate reduced basilar arbor branching and length. (a) Representative single plane Golgi images and corresponding reconstructions of L3 PCs generated from wildtype (left) and S1782E +/- (right) mice. (b) S1782E +/- mice show reduced branching and length in basilar arbors compared to wildtype. (c) Representative single plane Golgi images of a segment of basilar dendrite from a wildtype (left) and S1782E +/- mouse (right). Quantitiatively, S1782 +/- mice show a reduction in spine density along basilar branches compared to wildtype. Sholl data shown are averaged across N=10 animals/genotype (animal values generated by averaging 5 neurons/animal) ± SEM. Spine data shown are generated from the average value per animal across N=10 animals/genotype (animal values generated from 3-5 neurons/animal) ± SEM. *p<.05

### MAP2 interacts with ribosomal complexes and inhibits protein synthesis

MAP2 binding to MTs can sequester MAP2 and its binding partners^60^. Conversely, reduced MAP2 binding to MTs increases the availability of MAP2 to associate with alternate interaction partners^61^, creating what in essence is a gain of function of these latter effects. We therefore sought to use an unbiased proteomic screen of the MAP2 interactome to identify additional potential interaction partners. Co-IP of MAP2 from RNAse-treated whole brain homogenate from C57Bl6J mice (N=3 mice, 6 hemispheres total as biological samples) resulted in identification of 590 proteins that were potential interactors of MAP2 (q<.05) for MAP2 IP compared to no antibody control (Fig 5, see also Fig S3 and Table S4). Our unbiased approach highlighted an interaction previously observed in neurons, although underappreciated, between MAP2 and ribosomal complexes^62^. Functional annotation analysis^63^ of these proteins revealed enrichment for terms related to translation (e.g. GO:0006412~translation, corrected p=1.4E^-30^, N=71 proteins) and ribosome (e.g. GO:0005840~ribosome, corrected p=3.2E^-27^, N=46 proteins). Table S5 shows the 3 highest scored GO categories within the top 8 functional annotation clusters.

**Fig 5.**
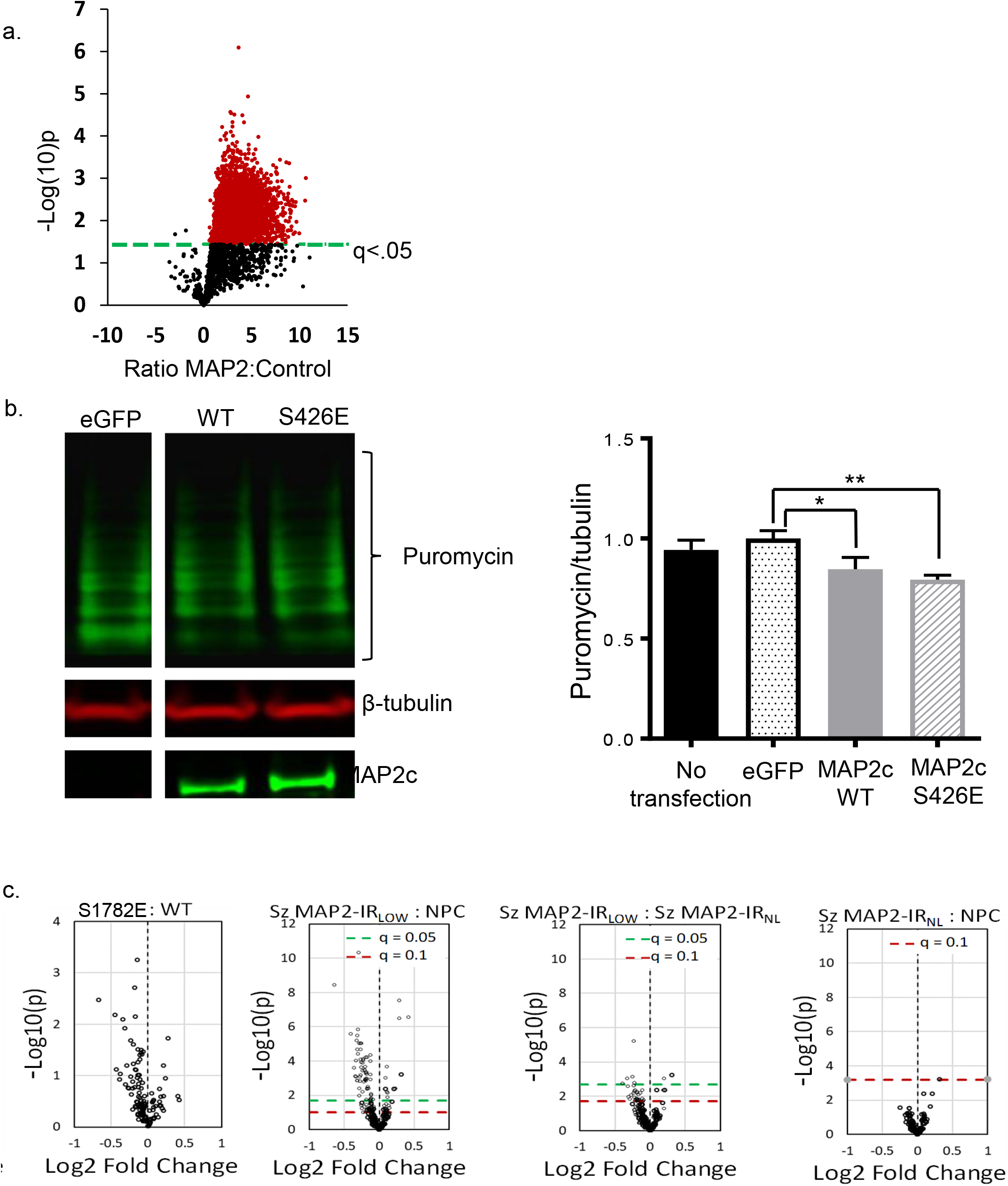
Co-IP of MAP2 from mouse brain and functional effects of S1782E on protein synthesis. (a) Using conservative cutoff criteria of q<.05, 6095 peptides (red) from 590 proteins were identified as potential MAP2 interactors. (b) Puromycin incorporation was detected via western blotting following a 1h treatment of transiently transfected HEK293 cells expressing MAP2c-WT or MAP2c-S426E in comparison to cells transfected with eGFP control for 48h. Densitometric quantification of relative puromycin incorporation normalized to MAP2 expression levels. N=4, data shown are average across experiments ± SEM. *p < 0.1, **p<.05. (c) Reductions in the synaptic proteome are seen in both MAP-IR_LOW_ Sz and S1782E mice. Volcano plots showing distribution of 159 synaptically enriched proteins in NPC, MAP2-IR_LOW_ and MAP2-IR_NL_ Sz (N=45 pairs; N=19 MAP2-IR_NL_, N=26 MAP2-IR_LOW_). Mice with phosphomimetic mutation S1782E have synaptic proteome alterations similar to those seen in MAP2-IR_LOW_ Sz (N=4 mice/genotype). Green dashed line indicates FDR-corrected p-value (q-value) of 0.05, red dashed line indicates q-value of 0.1.

Based on the above association of MAP2 with ribosomal complexes, we next hypothesized that MAP2 may play a role in translational regulation. To test this, we measured incorporation of puromycin, an indicator of new protein synthesis^64^. Expression of both WT or S426E MAP2c in HEK293 cells suppressed new protein synthesis compared to expression of eGFP alone, though the magnitude of this effect was slightly greater in S426E (Fig 5b). Because this finding was derived from study of HEK293 cells, we once again undertook to assess whether similar effects might be observed in neurons. Previous work from our lab using a targeted proteomics assay in synaptosomes prepared from A1 of Sz and NPC subjects found widespread reductions in the levels of multiple synaptic proteins^65^. We then generated synaptosome preparations from cerebral cortex of WT and S1782E+/+ mice and assayed protein levels with the same targeted proteomics assay. Fig 5c shows that mice with the phosphomimetic mutation S1782E have reduced levels of multiple synaptic proteins. We then undertook to re-analyze the data in our Sz subjects, dividing them into those in whom MAP2-IR was reduced (MAP2-IR_LOW_) or intact (MAP2-IR_NL_). Reduced synaptosome protein levels were restricted to the MAP2-IR_LOW_ Sz group (Fig 5c), similar to that seen in S1782E mice, although with only N=4 mice/genotype the p values are higher in the mice than in the postmortem study.

## Discussion

MAP2 is the predominant dendritic MAP in the brain, where its modification by phosphorylation downstream of glutamate signaling^17^ has been postulated to be a requisite step linking experience to structural plasticity of dendrites^66^. We implemented a phosphoproteomics approach to evaluate this post-translational modification of MAP2 in Sz subjects in comparison to NPC subjects. We identified multiple abnormally phosphorylated MAP2 peptides, located predominantly in the Proline Rich and C-terminal domains which flank the MT Binding domain and are present in all isoforms. The most altered site was pS1782, which was found to be increased nearly 8-fold in Sz subjects compared to NPC subjects. We carefully considered potential confounds (e.g. PMI, pH, tissue storage time, age, race, sex, antipsychotic treatment, substance use, tobacco use) in our human subjects, none of which accounted for the elevation in phosphopeptides in Sz. Instead, a module of cooccurring phosphopeptides including pS1782, correlated with increasing severity of structural, proteomic, and functional measures in our subjects, suggesting the possibility that some or all of these co-occurring phosphorylations may causally contribute to the pathogenesis of Sz. To evaluate potential causality, we chose the most elevated phosphorylation, pS1782, as a candidate for further study.

Prior studies have shown that phosphorylation of MAP2 within its MT binding domain and proline rich domains reduce MT binding by MAP2,^14^ and thus alter their dynamics.^67^ However, the effects of phosphorylation are anticipated to be site specific, and prior studies have not evaluated phosphorylation at S1782. Given the proximity of S1782 to the microtubule binding domain of MAP2, we utilized computational modeling to predict the effects of this modification on structure. We now show that phosphorylation of S1782 in the c-terminal domain of MAP2 can act similarly to phosphorylation at the homologous S396 in Tau, which (in combination with pS404) is primarily responsible for Tau dissociating from MTs.^68^ Subsequent *in vitro* studies confirmed this finding, demonstrating reduced MT-binding in an S426E (MAP2c) phosphomimetic construct.

Dendritic length and branching determine a PCs receptive field^69,70^, help to segment computational compartments^71^, and contribute substantially to how the received signals are integrated and transmitted to the cell body^72,73^. Similarly, dendritic spines serve to segregate synaptic afferents^74^. Reduced dendritic arborization and dendritic spine density (DSD) have consistently been found in postmortem studies of Sz^59^, and impaired frequency discrimination is a hallmark of the disease^75^. Since dendrites are a principal component of cortical gray matter, it is possible that their reduction in Sz is a contributing factor to the observed reductions in gray matter volumes in disease^76–81^. Our data suggest that pathologic hyperphosphorylation at S1782 in MAP2 leads to reductions in basilar dendritic length and complexity similar to those seen in Sz and likely directly contributes to disease pathology. Similarly, reduced DSD has more recently been shown to directly correlate with MAP2-IR^34^. Our analysis of S1782E mice demonstrates similar reductions in DSD to those described in Sz, suggesting that hyperphosphorylation at this site directly contributes to impairments in DSD.

While our current findings demonstrate the functional consequences of S1782E specifically, they should not be interpreted as a model of schizophrenia per se, in which other MAP2 phosphorylations are also present. For example, we did not replicate in S1782E mice the loss of MAP2-IR present in SZ, suggesting that other phosphorylation events on MAP2, either in isolation or in combination with S1782, are likely responsible for the loss of MAP2-IR in SZ. In keeping with this observation, we found that MAP2-IR in human brain was most highly correlated with levels of a different phosphopeptide RLS[+80]NVSS[+80]SGS[+80]INLLESPQLATLAEDVTAALAK (r=0.62; p<001). Thus, each pathologically altered MAP2 phosphorylation event may affect MAP2 structure and/or function differently, ultimately converging onto a commonality of disruption of normal MAP2 function and subsequent disease pathology. Moreover, at least one prior study has reported changes in immunoreactivity of an alternate MAP, MAP1B, in schizophrenia^13^. MAP1B can partially compensate for loss of MAP2, and correspondingly, concurrent knockdown of both generates a more severe impairment in neuron structure^82^.

Our computational models similarly identified a set of β-strands induced by S1782 phosphorylation between K368-V372 and A420-Q425 (K1724-V1728 and A1776-Q1781 in canonical MAP2b numbering), a more open configuration that provides additional conformational combinations required for protein-protein interaction and thus the potential for an additional gain of pathologic functions. Specifically, we hypothesized that the reduced MT-binding affinity caused by (pseudo)phosphorylation of S1782 would lead to increased MT-unbound MAP2 levels. Previous work has shown binding by some proteins to only MT-unbound MAP2^83^, suggesting that a careful balance between MT-bound and MT-unbound MAP2 may be necessary for proper cellular functioning. Hence this increase, in combination with new protein interactions enabled by the conformational changes, provides a means by which phosphorylation of MAP2 may yield additional pathological gains of function in schizophrenia. To determine the nature of these additional functions, we combined immunoenrichment with LC-MS/MS to characterize the MAP2 interactome. These studies revealed that the MAP2 interactome was enriched for interactions with ribosome and RNA-binding proteins forming the translational machinery (Tables S3 and S4). This finding is unlikely to be artifactual, as we eliminated the possibility of residual RNA binding through RNAse treatment prior to the co-IP. This finding suggests a novel role for MAP2 in regulation of protein translation. To determine the nature of this alternative function of MAP2 we used puromycin incorporation to measure protein synthesis. We found that overexpression of MAP2c suppresses protein translation *in vitro*, an effect which is slightly, though non-signficantly, enhanced by pseudophosphorylation at S426 MAP2c (S1782 in MA2P2b). However, because this observation was obtained in HEK293 cells, which are not a neuronal cell line, we looked for convergent evidence of this effect in neurons, without the potential bias of overexpression of WT and phosphomimetic MAP2. We therefore evaluated a complementary measure, synaptic protein levels (rather than measuring protein synthesis per se), in mouse brain tissue. We found reduced levels of multiple synaptic proteins in synaptosomes of S1782E in comparison to WT mice. The possibility that repression of synaptic protein synthesis by pMAP2 has a role in the pathogenesis of Sz is further supported by our findings of reduced levels of multiple synaptic proteins in synaptosomes of MAP2-IR_Low_, but not MAP2-IR_NL_, subjects with Sz. Taken together, these findings are the first to suggest that MAP2 may have an additional function in regulating protein translation and that phosphorylation at this site, alone or in combination with other phosphorylation sites, may alter this novel function of MAP2.

Long-standing findings have established that MAP2 is phosphorylated in response to synaptic activity that induces plasticity^84,85^. Recent studies have illuminated these prior observations, including direct demonstration that MAP2 is required for long term potentiation (LTP)^86^ a process dependent on new protein synthesis. Our replication of findings of an association of MAP2 with the protein translation machinery ^62,87,88^, and more importantly our novel demonstration that overexpression of WT or S426E MAP2 inhibits protein synthesis, provide a potential mechanism by which phosphorylation of MAP2 may impact LTP. A number of mRNAs are translated explicitly in dendrites after synaptic stimulation to provide rapid regulation of local protein levels in support of structural and functional plasticity.^89^ Our findings that WT MAP2 and S426E/S1782E MAP2 inhibits protein synthesis and reduces synaptic protein levels suggests MAP2 may impact this process. However, the presence of multiple ribosomal proteins in our co-ip of MAP2 likely reflects many indirect interactions of proteins within the ribosomal complexes. MAP2 has been previously shown to interact directly with KH domains that are present in some RNA binding proteins ^88^. Our MAP2 co-ip from mouse brain included multiple such KH domain containing proteins (HNRNPK_Mouse, FMR1_Mouse, PCBP1_Mouse, PCBP2_Mouse, PCBP3_Mouse), identifying them as candidates for mediating the interaction of MAP2 with ribosomal complexes and the inhibition of protein translation.

In summary, we identified a novel pathology of Sz, differential phosphorylation of multiple peptides within MAP2. We further demonstrated that at least one of these phosphorylations, pS1782, impairs MAP2 MT binding, reduces dendritic length, complexity, and DSD, and inhibits new protein synthesis-a function critical for dendritic spine plasticity. MAP2-IR has been shown to be reduced in multiple different regions of the cerebral cortex in schizophrenia^59,35^ and we recently showed that the magnitude of this reduction is conserved across multiple cortical regions within individuals with schizophrenia^90^. Similarly, reduced dendritic length, complexity, and DSD have been described across regions of the cerebral cortex in schizophrenia^59^. Thus, although the current study findings were obtained in auditory cortex, they are likely to generalize across the cerebral cortex and provide a new conceptual framework for schizophrenia as a “MAP2opathy”. That is, differential phosphorylation of MAP2, like its homolog Tau in dementia pathogenesis, may mediate impairments of neuronal structure and function downstream of many different genetic (and possibly environmental) risk factors. The potential impact of this conceptualization is that it identifies a bottleneck of molecular pathology providing a narrow set of targets for prevention and treatment for a large proportion of individuals with schizophrenia, regardless of the polygenic origins of their illness. Identification of the specific binding partner(s) of MAP2 within the protein translational machinery may provide leads to small molecules or peptides that disrupt this interaction and could then be tested as novel therapies for these individuals.

## Acknowledgements

MH071533 (RAS), NARSAD Distinguished Investigator Grant from the Brain & Behavior Research Foundation (RAS), MH118513-01 (MJG). We thank Duncan R. Groebe, PhD, MBA for assistance in the editing of this manuscript.

## Author Contributions

MJG performed co-IP experiments, Golgi reconstructions, data analysis and interpretation, and writing of the manuscript. XS performed MTB assay, puromycin incorporation, and contributed to the conceptualization of the project and first drafts of parts of the manuscript. MLM, MG, and NAY performed the phosphoproteomics and LC/MS-MS on co-IP samples. ZS and YD performed all advanced statistical and network analyses. RAD performed confirmatory experiments and contributed to the data interpretation and analysis. DAL contributed to data interpretation and analysis. GEH generated the transgenic mouse model and contributed to data interpretation and analysis. CC, KP, and DP performed computational modeling. RAS oversaw the project conceptualization, all data analysis and interpretation, and manuscript writing.

## Competing Interests

None to report (MJG, MLM, MG, ZS, RAD, DAL, NAY, KP, DP, CC, GH, YD). The Tsinghua Educational Foundation North America and the University of Pittsburgh have an ongoing collaboration to allow visiting scholars from Tshingua University to gain laboratory experience within a Univeristy of Pittsburgh affiliated laboratory. XS, a University of Pittsburgh-affiliated visiting research scholar working in Dr. Sweet’s lab from August 1, 2016 through July 31, 2018, received no financial support from Dr. Sweet and did not contribute to any NIH-funded aspects of this work.

Corresponding Author: Robert A. Sweet, University of Pittsburgh

## Methods

### Human Subjects and Animals

Tissue was obtained from postmortem brains recovered during autopsies conducted at the Allegheny County Medical Examiner’s Office, Pittsburgh, PA, following informed consent from the next of kin. All procedures were approved by the University of Pittsburgh Committee for the Oversight of Research and Clinical Trials Involving the Dead and the Institutional Review Board for Biomedical Research as previously described^37,92^. An independent committee of experienced clinicians made consensus Diagnostic and Statistical Manual of Mental Disorders, Fourth Edition diagnoses, or absence thereof, for each subject using medical records, structured interviews with surviving relatives, and, when available, toxicology reports. For the phosphoproteomics, 16 individuals with a consensus diagnosis of schizophrenia (Sz) and 16 matched non-psychiatric control (NPC) subjects, all of whom had been previously characterized for MAP2 immunoreactivity (MAP2-IR) and dendritic spine density^34^, were studied (Table S1). The Sz subjects in this cohort were further divided by MAP2-IR as either low (MAP2-IRLOW) or normal (MAP2-IRNL)^33,34^.

In order to account for the effects of chronic antipsychotic drug administration, a cohort of 18 male rhesus monkeys, n = 6 per group, were administered therapeutically relevant doses of haloperidol, olanzapine, or vehicle, and monkeys were sacrificed in matched triads and tissue was immediately harvested and processed^35^. The tissue utilized here has been extensively described elsewhere^93,94^. Briefly, tissue slabs containing the superior temporal gyrus were removed in a single block and 20 mg of gray matter was harvested as described above. A mouse model of postmortem interval (PMI) was generated using wildtype C57/Bl6J mice purchased from Jackson Laboratories (Bar Harbor, ME, USA). Adult mice, n= 2 per timepoint (0, 6, 12, 18, 24 hrs), were sacrificed by CO_2_ asphyxiation followed by cervical dislocation. The carcasses were incubated at room temperature for 2/3 the PMI duration and then placed at 4°C for the final 1/3. The brain was then removed from the skull, the cerebellum discarded, and the hemispheres separated (giving n = 4 hemispheres/time point), flash frozen in isopentane on dry ice, and stored at −80°C until use.

### Phosphoproteomics

#### Sample Preparation – Phosphopeptide enrichment

Gray matter from frozen right hemisphere tissue blocks containing Heshl’s gyrus was harvested as previously described^95^. Protein quantification was determined by using a micro bicinchoninic acid assay. The samples were processed across multiple runs, using a balanced block design to evenly distribute schizophrenia and control subjects. Each run was processed identically. To account for biological and experimental variation, pooled controls were included in the designed and interspersed throughout. 500μg of protein from each gray matter homogenate sample were in-solution digested overnight at 37C with trypsin at a ratio of 1:50 (ug trypsin: ug total protein). Samples were then desalted using Oasis cartridges. Phosphorylated peptides were enriched using Titanium Dioxide Phosphoenrichment. To perform the titanium dioxide phosphoenrichment, ToptTips TiO2 + ZrO2 (Glygen Corporation, Columbia, MD, USA)) were washed three times with loading buffer (19% lactic acid, 80% ACN, 1% formic acid) and centrifuged 30 sec x 0.5g. Beads were suspended to 50% slurry with the loading buffer and 20μL of bead slurry was added to each sample. The sample + bead mixtures were rotated for 1 hour. Samples were spun 30sec x 0.5g and supernatant was removed. 1 mL of the loading buffer was added to each sample, and rotated for 5 minutes. Samples were centrifuged and supernatant removed. Sample was suspended in 200μL washing buffer (80% ACN, 5%TFA) and centrifuged through a C18 Stage tip. C18 stage tip was subsequently washed x4 with washing buffer. Phosphopeptides were eluted with 50μL elution buffer (450μL Ammonium Hydroxide (28%) in 4.5 mL DI H_2_0 pH 11) 2 times. Elution buffer was evaporated and samples re-suspended in 20μL 98% buffer A (0.1% formic acid/DI H_2_0) 2% buffer B (0.1% formic acid/ACN). Samples were then analyzed by LC-MS/MS.

#### Liquid Chromatography Tandem Mass Spectrometry: Phosphopeptides

Phosphopeptide enrichments were analyzed by reversed phase nanoflow liquid chromatography tandem mass spectrometry (nLC-MS/MS) using a nanoACQUITY (Waters Corporation, Milford, MA, USA) coupled to an Orbitrap XL hybrid ion trap mass spectrometer (ThermoFisher Scientific, Waltham, MA, USA). ~ 1 ug of phosphopeptides were eluted over a 60-minute gradient from 95% buffer A (2% Acetonitrile, 0.1% formic acid) to 35% buffer B (Acetonitrile, 0.1% formic acid) on a 3 μm 120A; 205mm REPROSIL-Pur C18 Picochip (New Objective). Sample eluate was electrosprayed (2000V) into a Thermo Scientific Orbitrap XL mass spectrometer for analysis. The instrument was operated in MS2, top 4. MS1 spectra were acquired at a resolving power of 60,000. MS2 spectra acquired in the ion trap with CID, normalized collision energy = 35. Dynamic exclusion was enabled to minimize the redundant selection of peptides previously selected for MS/MS. Ion chromatograms of 10 selected peptides were extracted using Skyline software (version 3.5)^96^ and used to monitor during the instrumental analysis to ensure robust instrument performance.

#### MS/MS data processing

MS/MS spectra from the phosphopeptide experiment were searched using MASCOT search engine (Version 2.4.0, Matrix Science Ltd, Boston, MA, USA) against the UniProt human proteome database. The following modifications were used: static modification of cysteine (carboxyamidomethylation, +57.05 Da), variable modification of methionine (oxidation, +15.99Da), variable modification of serine, threonine, and tyrosine (phosphorylation, +79.67Da). The mass tolerance was set at 20ppm for the precursor ions and 0.8 Da for the fragment ions. Peptide identifications were filtered using PeptideProphet^™^ and ProteinProphet^®^ algorithms with a protein threshold cutoff of 99% and peptide threshold cutoff of 90% implemented in Scaffold^™^ (v4.7.5; Proteome Software, Portland, Oregon, USA). Skyline MS1 filtering^97^ was used for batched integration of ion chromatograms of all identified peptides. MS2 spectra for MAP2 were manually sequenced to confirm the phosphopeptide-spectra match and confidence of phospho-site assignment.

### Preparation of expression constructs

IRES-MAP2c-EGFP plasmids containing cDNAs encoding human MAP2c (NM_031845) were purchased from GeneCopoeiaTM (Cat#: CS-E2438-M61-01). Although MAP2c is a shorter isoform, we use the amino acid numbering designated by full-length MAP2 throughout unless specifically referring to an experiment using the MAP2c plasmid (i.e. S426 in MAP2c corresponds to S1782 using numbering based on the canonical MAP2b isoform). MAP2c phosphomimetic mutant S426E was generated using the QuikChange Lightning site-directed mutagenesis kit (Agilent Technologies) and mutagenized primers (Table S5). Plasmids were purified using the EndoFree Plasmid Maxi Kit (Qiagen). Presence of the mutation at the desired site without additional off-target effects was evaluated by Sanger sequencing (Table S5). Transfection with the IRES-EGFP plasmid (eGFP) lacking the MAP2c construct was used as a control.

### Cell culture and transfection

HEK293 cells were grown in RPMI-1640 containing 5% fetal bovine serum (Cellgro, Manassas, VA) without antibiotics. They were grown in a humidified chamber at 37 degrees Celsius with 5% CO_2_. They were passaged appropriately during linear growth phase to maintain sub-confluent culture. For transfections they were seeded at 1.5×10^5^ cells/well into 12-well culture-treated plates. They were serum starved for 16h overnight, then replenished with complete media at the time of transfection the following morning.

#### Transfection

All transfections were performed using Lipofectamine LTX with Plus reagent (ThermoFisher Scientific, Grand Island, NY) in the absence of antibiotics. Media was changed 16h after transfection.

### Western Blotting

Cell-free protein lysates were obtained from cells using radioimmune precipitation assay buffer (10 mM Tris-HCl, pH 8.0, 1 mM EDTA, pH 8.0, 0.5 mM EGTA, 140 mM NaCl, 1% Triton X-100, 0.1% sodium deoxycholate, 0.1% SDS), and total protein levels were quantified using the Lowry protein assay (Bio-Rad). 15 μg of total protein was loaded per well. Samples were electrophoresed on a NuPage Bis-Tris 4-12% gradient polyacrylamide gel (ThermoFisher Scientific) and transferred to a PVDF membrane (Millipore, Danvers, MA) at 100V for 1h. Membranes were blocked in Licor Blocking buffer (Odyssey, Lincoln, NE) for 1h at room temperature and primary antibody was added and incubated overnight at 4 degrees Celsius. The following primary antibodies were used in this study: mouse anti-MAP2 (HM-2, Abcam, 1:1000), rabbit anti-β tubulin (Abcam, 1:10,000), and mouse anti-puromycin (12D10, Millipore Sigma, 1: 25,000). Membranes were washed 5×5min in TBS containing 0.1% Tween and incubated with fluorescent-conjugated secondary antibodies (1:10,000) for 1h at room temperature. Five more washes were performed and membranes were imaged using the Odyssey Licor System. Densitometric analyses were performed using MCID Core Analysis software (UK).

### CRISPR/Cas9 Mouse Generation

#### Reagent Production

An sgRNA targeting MAP2 was selected using CRISPOR ^98^. An sgRNA specific forward primer and a common overlapping reverse PCR primer (see Table S6) were used to generate a T7 promoter containing sgRNA template as described ^99^. This DNA template was transcribed *in vitro* using a MEGAshortscript Kit (Ambion). The Cas9 coding sequence was amplified from pX330 ^100^ using a T7 promoter containing forward primer and reverse primer (see Table S6) and subcloned into pCR2.1-TOPO. This plasmid was digested with EcoRI and the ~4.3 kb fragment was *in vitro* transcribed and polyA tailed using the mMessage mMachine T7 Ultra Kit (Ambion). Following synthesis, the sgRNAs and Cas9 mRNA were purified using the MEGAclear Kit (Ambion), ethanol precipitated, and resuspended in DEPC treated water. A single stranded 130 nucleotide DNA repair template (see Table S6) harboring the knockin sequences (TCC changed to GAG) and sequences homologous to MAP2 were purchased as Ultramer DNA (Integrated DNA Technologies, Coralville, IA). The oligo included 3 phosphorothioate linkages on each end ^101^, was asymmetrically oriented with respect to the PAM, and was complementary to the nontarget strand ^102^.

#### Knockin Mouse Production

sgRNA (200ng/μL), Cas9 mRNA (200ng/μL, NEB cat#M0646), and the repair oligo (200ng/μL) were mixed for electroporation using a Biorad Gene Pulser XCell and custom electroporation chamber (Protech International Inc., cat# LF501-P1X10). sgRNAs and protein were diluted with Opti-MEM (ThermoFisher Scientific) prior to electroporation; OMEM constituted at least 50% of the electroporation solution. Electroporation conditions consisted of five repeats of 3 msec square wave pulses of 25V with 100 msec intervals. Following electroporation, embryos were washed in M2 and transferred to KSOM for culture prior to embryo transfer. Embryos were transferred as 2-cells to the oviducts of day 0.5 postcoitum pseudopregnant CD-1 females.

#### Mouse Genotype Analysis

Mice were genotyped by PCR amplification (see Table S6 for primer sequences, MAP2 F1 and MAP2 R1) followed by restriction digestion with BspLI (FastDigest) to produce PCR amplicons that were either 487bp (wildtype) or 337bp + 150bp (knockin). Undigested PCR amplicons of founder and all F1 offspring were also analyzed by Sanger sequencing.

#### Off target analysis

The top 10 predicted off-target sites (see Table S7) using CRISPOR software were analyzed by PCR amplification followed by Sanger sequencing.

#### Mouse Production

A single male that was confirmed to be heterozygous for the mutation and free of off-target effects was chosen as the founder for line generation. He was mated with wildtype C57BL/6J females purchased from Jackson Laboratories (Bar Harbor, ME) and all F1 mice were genotyped by both PCR and restriction digest as well as confirmatory Sanger sequencing to establish the MAP2 S/E line. Heterozygotes were backcrossed into purchased wildtype C57BL/6J strain to produce animals for all experimental studies mitigating against genetic drift of the line. Throughtout the text animals are referred to as S1782E mice.

Animals were identified with metal ear tags and tail snip DNA samples were obtained for genotyping by PCR prior to weaning at approximately postnatal day 21. Genotypes of all mice included in studies were confirmed following sacrifice. Both male and female animals were included in all studies. Animals were under specific pathogen free conditions and housed in standard microisolator caging (Allentown Caging Equipment, Allentown, NJ, USA) in groups of up to 4 males or 5 females, maintained on a 12-h light/dark cycle (lights on at 7 AM), and were provided with food and water ad libitum. All experimental procedures were approved by the Institutional Animal Care and Use Committee at the University of Pittsburgh.

### Golgi

#### Tissue Generation

Cohorts of mice were sacrificed at age 12 weeks ± 3 days. Age-matched cohorts that included males and females of the 2 genotypes of interest (wildtype and S1782E +/-) were processed together throughout. Vaginal cytology was collected on all female mice at the time of sacrifice. Mice were sacrificed by lethal CO_2_ inhalation followed by decapitation and whole brain extraction. Golgi staining was performed using the FD Rapid GolgiStain Kit (FD Neurotechnologies, Inc) according to manufacturer’s instructions. Brains were sectioned (150 μm) on a cryostat. Tissue sections were placed onto gelatin-coated slides and allowed to dry overnight at room temperature. The following day sections were then rehydrated with distilled and deionized water, reacted in a developing solution (FD Rapid GolgiStain Kit, FD Neurotechnologies), and dehydrated through graded ethanols. Finally, sections were cleared in xylene and coverslipped using Permount (Fisher Scientific). Auditory cortex was defined using gross anatomical landmarks and alignment with the corresponding Nissl plate containing A1 in the mouse brain atlas^103^. Within A1, deep layer 3 was defined as 30-50% of the distance between the pial surface and white matter border and sampling of pyramidal cells was performed as described below.

#### Imaging

For the analysis of dendritic structure *in vivo*, image acquisition was performed on a custom-built, wide-field epifluorescence Olympus IX73 microscope equipped with XYZ-encoded prior stage, real-time autofocus, and a Hamamatsu ORCA-Flash4.0 sCMOS camera controlled with SlideBook 6.0 using an Olympus PlanAPOS 40x 0.90 NA air objective. Brightfield exposure time was optimized for each image. Image stacks, 1 μm step-size, through the entire tissue depth were captured of pyramidal shaped cells within the region of interest. Pyramidal cells were included if they did not have overlapping soma from neighboring cells and their soma was located roughly within 40-60% of the z-depth. Collected image planes were 2048×2048 pixels with a resolution of 0.161μm/pixel. A total of 5 neurons per animal distributed across 2-3 sections were collected from the right hemisphere. For dendritic reconstruction, image stacks were manually traced offline using NeuroLucida software (Microbrightfield, Inc) and total dendritic content was analyzed via Sholl analysis using NeuroExplorer (Microbrightfield, Inc). Five animals per sex per genotype were included. Experimenter was blinded to the genotype throughout image acquisition and processing.

For the analysis of dendritic spine density, the same imaging system as described above was employed. PCs meeting the above criteria were identified at 20x magnification, and image stacks were subsequently collected using an Olympus PlanAPOS 60x 1.42 NA oil objective at a step size of 0.7 μM. For spine counting, image stacks were manually evaluated offline using StereoInvestigator software (MBF Bioscience). For the analysis of spine density, we chose to focus on basilar dendrites. To be included in analysis a basilar dendrite was required to have >75μm total length, branch at least once, and be free of overlapping dendrites that obscured accurate visualization. As dendritic spine density is not uniform across the entirety of the arbor^104^, we ensured there was no difference across genotypes in average length (MAP2-WT 144.9 ± 54.1 μm, MAP2-S/E 145.8 ± 72.2 μm, p=0.95) or number of branches (MAP2-WT 2.4 ± 0.64, MAP2-S/E 2.6 ± 0.79, p=0.35) of included dendrites. Three to five neurons per animal were included in the final analyses.

### MAP2 Co-immunoprecipitation

#### Tissue preparation and Co-IP

For co-immunoprecipitation (Co-IP) from 4-week old mouse brain tissue, mice were sacrificed by lethal CO2 followed by rapid decapitation. Whole mouse brains were then extracted, snap frozen and stored at −80 °C until use. Frozen brain tissue was manually homogenized in ice-cold 0.5% NP-40 IP lysis buffer with RNAse (100uL per 1mL of lysis buffer) by micro-pestle and passing through 23G needle multiple times. The lysate was then incubated on ice for 10 min followed by centrifugation at 14,000g for 20 min at 4 °C to remove the insoluble fraction. The supernatant was centrifuged again at 12,000g for 10 min at 4 °C. The supernatant was transferred to a new tube, 50μl of which was saved as input, and the rest of the supernatant was precleared by incubation with 1mg 1xPBS pre-washed Dynabeads (Thermo Fisher Scientific) on a rotator at 4°C for 1h. To prepare the antibody-coupled beads, HM-2 antibody (Abcam, Cambridge, MA, USA) was incubated together with washed Dynabeads (8μg antibody per mg beads used) in 1x PBS buffer with 0.1M citrate (pH 3.1) on a rotator at 4°C overnight. The precleared supernatant was incubated with antibody-coupled beads on a rotator at 4°C overnight. Controls were generated using washed beads not coupled to antibody. Following overnight incubation, the supernatant was collected and saved in a new tube. The Co-IP beads were washed three times in ice-cold 1xPBS buffer by gentle pipetting and directly boiled in 60 μl 1x SDS sample buffer (30 mM Tris-HCl, pH 7.4, 1% SDS, 12.5% glycerol, 2.5% β-mercaptoethanol, 0.05% Orange G) at 95 °C for 10 min to elute protein complexes from the beads.

#### In-gel trypsin digestion

Immunoprecipitated samples were shortly separated (1cm) by SDS-PAGE and stained with SimplyBlue^™^ SafeStain (Thermo Fisher Scientific). The stained gel regions were excised and in-gel trypsin digested as previously described ^105^. In brief, the excised gel bands were washed with HPLC water and destained with 50% acetonitrile (ACN)/25mM ammonium bicarbonate until no visible staining. Gel pieces were dehydrated with 100% ACN, reduced with 10mM dithiothreitol (DTT) at 56°C for 1 hour, followed by alkylation with 55mM iodoacetamide (IAA) at room temperature for 45min in the dark. Gel pieces were then again dehydrated with 100% ACN to remove excess DTT and IAA, and rehydrated with 20ng/μl trypsin/25mM ammonium bicarbonate and digested overnight at 37°C. The resulting tryptic peptides were desalted with Pierce C18 Spin Columns, speed-vac dried and resuspended in 18μL 0.1% formic acid. A pooled instrument control (PIC) was made by taking 3μL from each of the samples.

#### LC--MS/MS

Extracted tryptic peptides were analyzed by nano reversed-phase liquid chromatography tandem mass spectrometry (nLC-MS/MS) using a nanoACQUITY (Waters Corporation, Milford, MA, USA) online coupled with an Orbitrap Velos Pro hybrid ion trap mass spectrometer (Thermo Fisher Scientific). For each LC-MS/MS analysis, 1 μL of peptides was injected onto a C18 column (PicoChip^™^ 25 cm column packed with Reprosil C18 3 μm 120Å chromatography media with a 75 μm ID column and 15 μm tip; New Objective, Inc., Woburn, MA, USA) and then eluted off into the mass spectrometer with a 66 min linear gradient of 2-35% ACN/0.1% FA at a 300 nL/min. A data-dependent top 13 method was used with full scan resolution set at 60,000. Ions were isolated for MS/MS analysis with a 2.0 Da window. Dynamic exclusion and chromatograms from 20 selected peptides were utilized as described above.

#### MS/MS data processing

MS/MS spectra from the MAP2 co-IP experiment were searched and processed as described above for phosphoproteomic study using the mouse proteome database.

#### Functional Annotation Analysis

Peptides positively identified as potential MAP2 interactors (Table S4) were imported into DAVID (https://david.ncifcrf.gov/) and the conversion tool was used to identify 567 unique DAVID protein IDs within species mus musculus. Mus musculus was selected as the background and functional annotation analysis was performed using the default settings (Table S5). ^63^

### Microtubule-binding (MTB) assay

Transfected HEK293 cells were harvested 48 h post transfection with ice-cold microtubule-binding buffer (80 mM Pipes, 5 mM MgCl2, 0.5 mM EGTA) supplemented with 1% Triton X-100 and protease/phosphatase inhibitors (Millipore, Burlington, MA, USA; and Sigma, St. Louis, MO, USA, respectively) and lysed by vortex and passing through a 26G needle multiple times. Protein lysates were first centrifuged at 17,000g for 10 min at 4 °C to remove insoluble cell debris, after which the supernatant was collected and further pre-cleared by ultracentrifugation at 100,000 g for 40 min at 4°C. *In vitro* microtubule-binding assay was performed using the microtubule-binding protein spin down assay kit (Cytoskeleton, Denver, CO, USA) according to the manufacturer’s instructions. Briefly, microtubules were freshly polymerized and kept at room temperature prior to use. For each microtubule binding reaction, 20μl freshly prepared microtubules were incubated with a series of volumes (10 μL /20 μL /30μL) of pre-cleared cell lysate for 30 min at room temperature. Pre-cleared cell lysate alone in incubation reaction was included as negative control. Moreover, MAP fraction (contains 60% MAP2 at 280 kDa and 40% tau proteins at 40-70 kDa) and BSA, each incubated with the polymerized microtubules, served as a technical positive and negative control, respectively. Reaction solution was then placed on top of 100 μl taxol supplemented Cushion buffer (80mM PIPES pH 7.0, 1 mM MgCl2, 1 mM EGTA, 60% Glycerol) and centrifuged at 100,000 g for 40 min at room temperature. The pellets containing microtubules and their binding proteins were resuspended in 1x SDS sample buffer as described above. Supernatant and pellet samples were analyzed by SDS-PAGE and Western blot.

### MD simulation

The molecular dynamic (MD) systems were built using cubic TIP3P water box with 12 Å buffer around the solute following the general procedure of protein system building in CHARMM-GUI^106^. Cl- and Na+ ions were added to the solvent to neutralize the charge of the systems. All MD simulations were performed using NAMD^107^ and CHARMM C36m FF (optimized for disordered proteins)^108^. Secondary structure analysis of MD simulation trajectories were carried out using Timeline Plugin available within VMDv1.9.2 (that utilizes STRIDE^109^ for secondary structure analysis and an in-house script for identifying extended β conformations that are not included as part of a β sheet or hairpin using serial successive phi/psi angles. Initially all systems were minimized twice and then equilibrated them before beginning production runs. In the first minimization solute, atoms were held fixed through 500 steps of steepest descent and 500 steps of conjugate gradient minimization. In the second minimization, only the solute backbone atoms were restrained through 2000 steps of steepest descent and 3000 steps of conjugate gradient. After minimization, the system temperature was raised to 300 K over the course of a 100 ps constant volume simulation during which the solute is fixed with weak (10.0 kcal/mol) restraints. For the production run, the 500 ns simulations was held at constant temperature 300 K and under constant pressure of 1 bar without any restraints used during equilibration step. For this, Langevin dynamics was used to maintain constant temperature with a Langevin coupling coefficient of 1 ps–1, and a Nosé-Hoover Langevin piston was used to maintain constant pressure with a piston period of 50 fs and a piston decay time of 25 fs while keeping barostating isotropic. A 2-fs timestep was used for integration together with the SHAKE algorithm where all covalent bonds between hydrogen and the heavy atom are constrained. The van der Waals interactions were smoothly switched off at 10 – 12 Å by a force-switching function while the long-range electrostatic interactions were calculated using the particle-mesh Ewald method.

### Surface sensing of translation (SUnSET) Assay

Surface sensing of translation (SUnSET) assay was performed as previously described^64^ with minor modifications: HEK293 cells were incubated with 10 μg/ml of puromycin in cell culture media for 1 h before harvest. As a negative control, cells were pretreated with 10 μL of cycloheximide (CHX) to inhibit protein synthesis for 1h before puromycin treatment. No puromycin incorporation was detected under these conditions (data not shown). Levels of newly synthesized proteins incorporating puromycin were measured using western blots as described above.

### Synaptosome Preparations

Full methodological information for the generation of the synaptosome data on the human subject cohort has been previously published in ^65^. To generate the synaptosome data from the S1782E transgenic mouse line, gray matter from reproductively capable mice (N=4 WT, 7.5 ± 0.6 weeks; N=4 S1782E +/+, 8.3 ± 3.3 weeks) was homogenized and subjected to a sucrose density gradient centrifugation method ^110^. Detailed methods for sample preparation, MS/MS, and data processing are fully described in ^65^.

### Statistical analysis

ANCOVA was performed to detect the difference across Sz with MAP2-IR_LOW_, Sz with MAP2-IR_NL_, and NPC samples in 18 MAP2 phosphopetides. PMI and assay run were adjusted as covariates. Fold changes, p-values, and q-values between each Sz group and control samples are presented (Figure 2a).

We also analyzed the association between MAP2 phosphopetides and each demographical or clinical variable using a similar ANCOVA model to detect potential confounders, where diagnosis, PMI and assay run were adjusted. P-values and q-values for each variable are presented (Table S3).

Weighted Gene Co-expression Network Analysis (WGCNA)^36^ was used to investigate the pattern of coregulated phosphopeptide expression in all samples. Topological overlap matrix will be used to identify modules with clustered phosphopetides. The three identified networks were visualized using Cytoscape. To quantitate the representation of expression levels in each module, we calculated the first principal component of modules given by WGCNA, referred to as the module-specific eigenpeptide. Linear regression models were fitted to model the correlation between each module-specific eigenpeptide and dendritic spine density, synaptic protein levels, and socioeconomic status (SES), with diagnosis (Sz or control) adjusted.

For Sholl analyses, we averaged the neurons within each animal. Then we plotted the average length or intersection count across all radii up to 140 μm, separated by genotype. We also plotted the trajectory of length or intersection count over radius for each genotype group. A linear-mixed effects model ^111^ was fitted for each length or intersection count, where radius (treated categorical), sex, genotype, and radius-by-genotype interaction were treated as the fixed effects, and mouse was treated as the random effect. To control for estrus phase, instead of the 2-category sex, a 4-category “sex” including male and three vaginal cytology for the female cohort was included as a covariate in the final model. The analysis was performed with the PROC MIXED procedure in SAS 9.4. Spine density analysis was performed using a student’s t-test.

Statistical analyses for the MTB and puromycin incorporation assays were performed in SPSS software with One-way ANOVA and Bonferroni post-hoc test to determine statistically significant differences for all the comparisons. All statistical tests were two-sided with α= 0.05. Data are displayed ± standard error of mean (SEM).

Student’s t-test on log2 transformed peptide peak area was used for the statistical inference to select Map2 interacting partners. A peptide was considered differentially ID’d if the q-value was <.1 and a positive Map2 CoIP:Control ratio value.

ANCOVA was performed to detect the difference in synaptosome proteins among three phenotype groups (Sz MAP2-IR_low_, Sz MAP2-IR_NL_, NPC). PMI and assay run were adjusted as covariates. P-values and fold change values were generated and plotted for each group comparison.

