## Supplemental Figures for "MAP2 is Differentially Phosphorylated in Schizophrenia, Altering its Function"

Figure S1

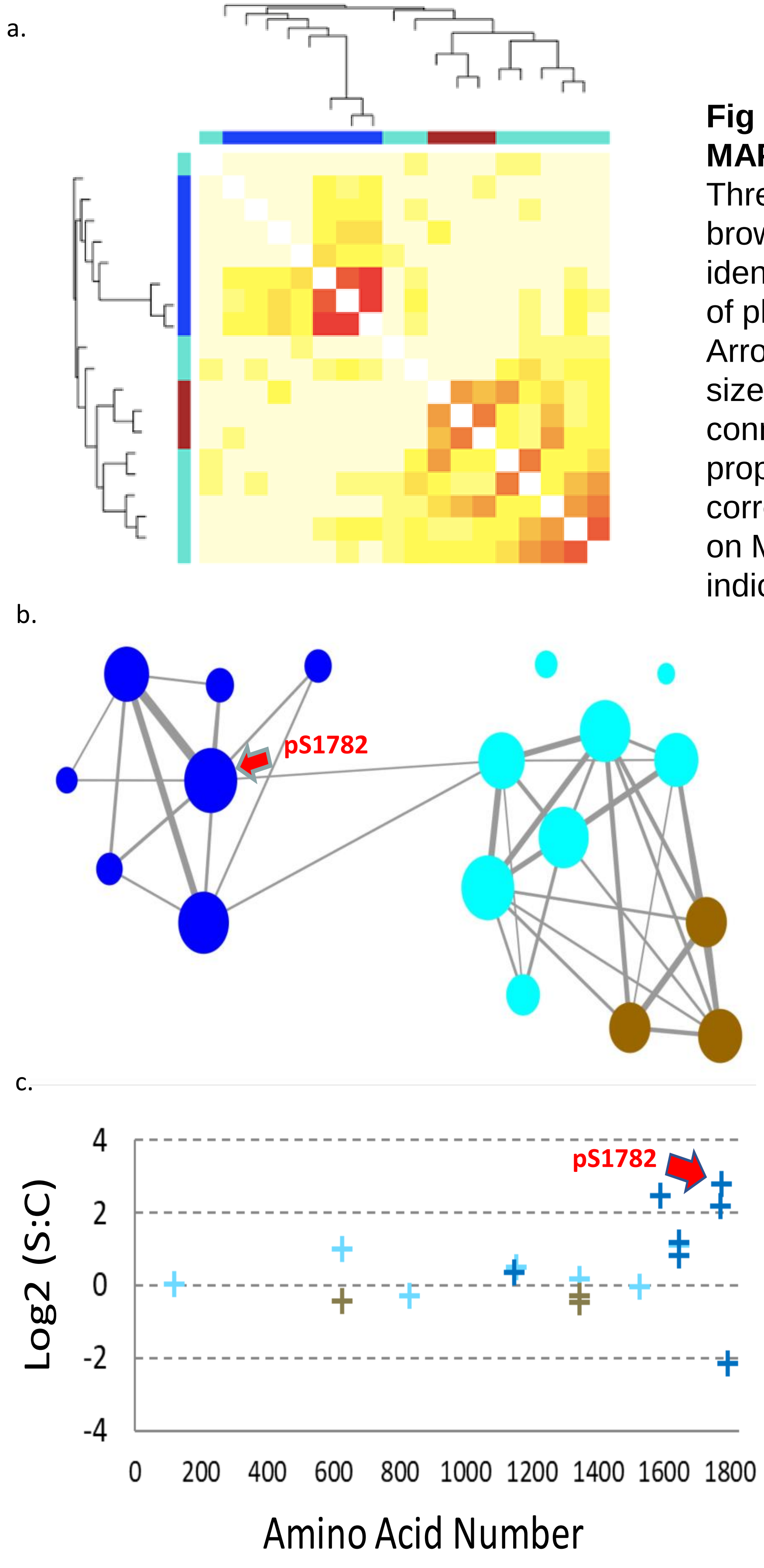


**Fig S1. WGCNA analysis of MAP2 phosphopeptides.** (a) Three distinct modules (blue, brown, and turquoise) were identified. (b) Network diagram of phosphopeptide modules. Arrow indicates S1782. Node size is proportional to node connectivity, line thickness is proportional to pairwise correlation. (c) Modules shown on MAP2B sequence. Arrow indicates S1782.

Figure S2

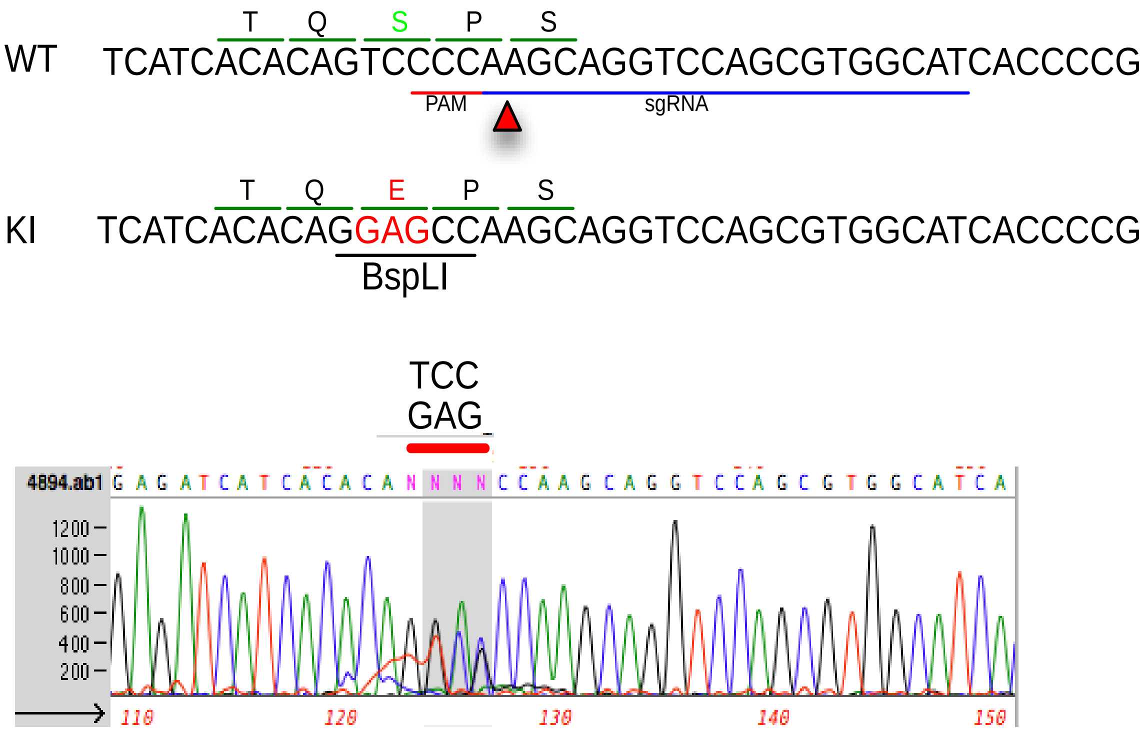


**Figure S2. S1782E knockin mouse production.** Sequence of wildtype and knockin genes with targeted codon indicated. Target site of PAM and sgRNA indicated below genetic sequence on wildtype, BspLI restriction site indicated on KI sequence.

Figure S3

a.

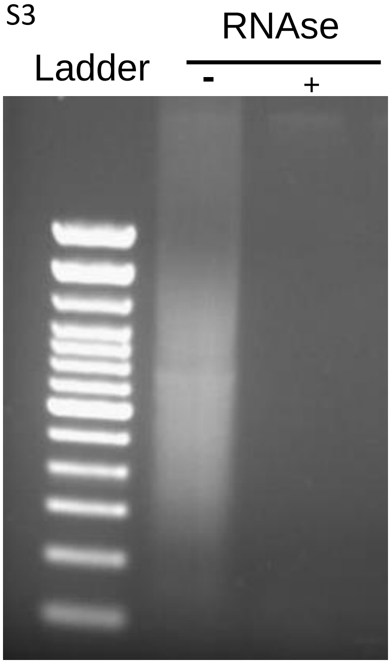


b.

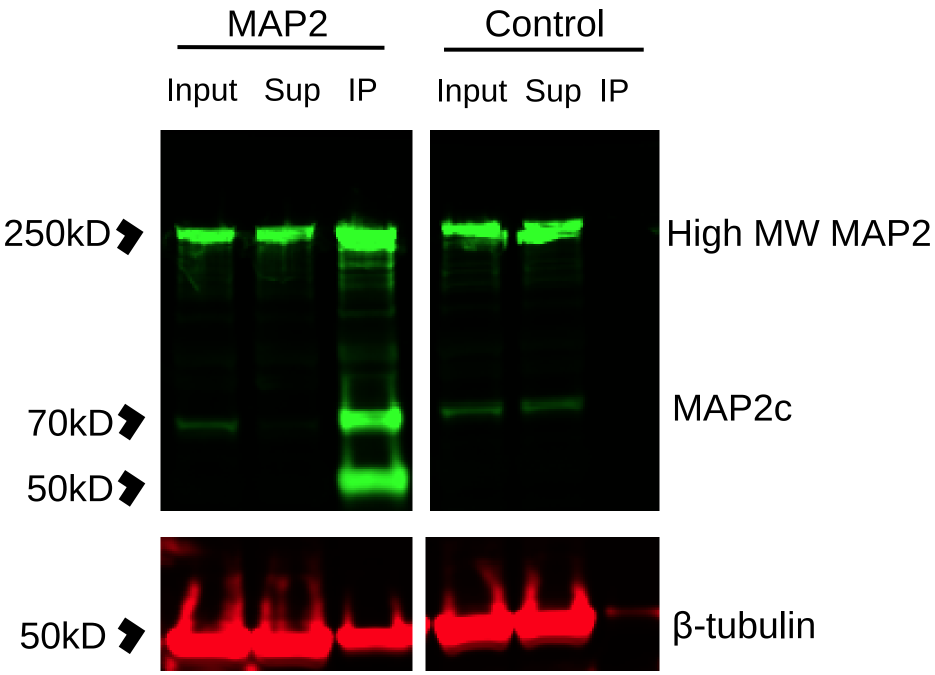


**Fig S3. Co-IP of MAP2 from mouse brain.** (a) Agarose gel image confirming degradation of RNA in tissue lysate prior to co-IP. (b) MAP2c was IP'd from RNAse treated mouse brain cortical homogenate (sup = supernatant, IP = immunoprecipitant). Beads without antibody coupling were used as a control. Individual channels are shown vertically (MAP2 in 800 channel, β-tubulin in 700 channel).

Figure S4

a.

**VDHGAEIITQS[+80]PGR**

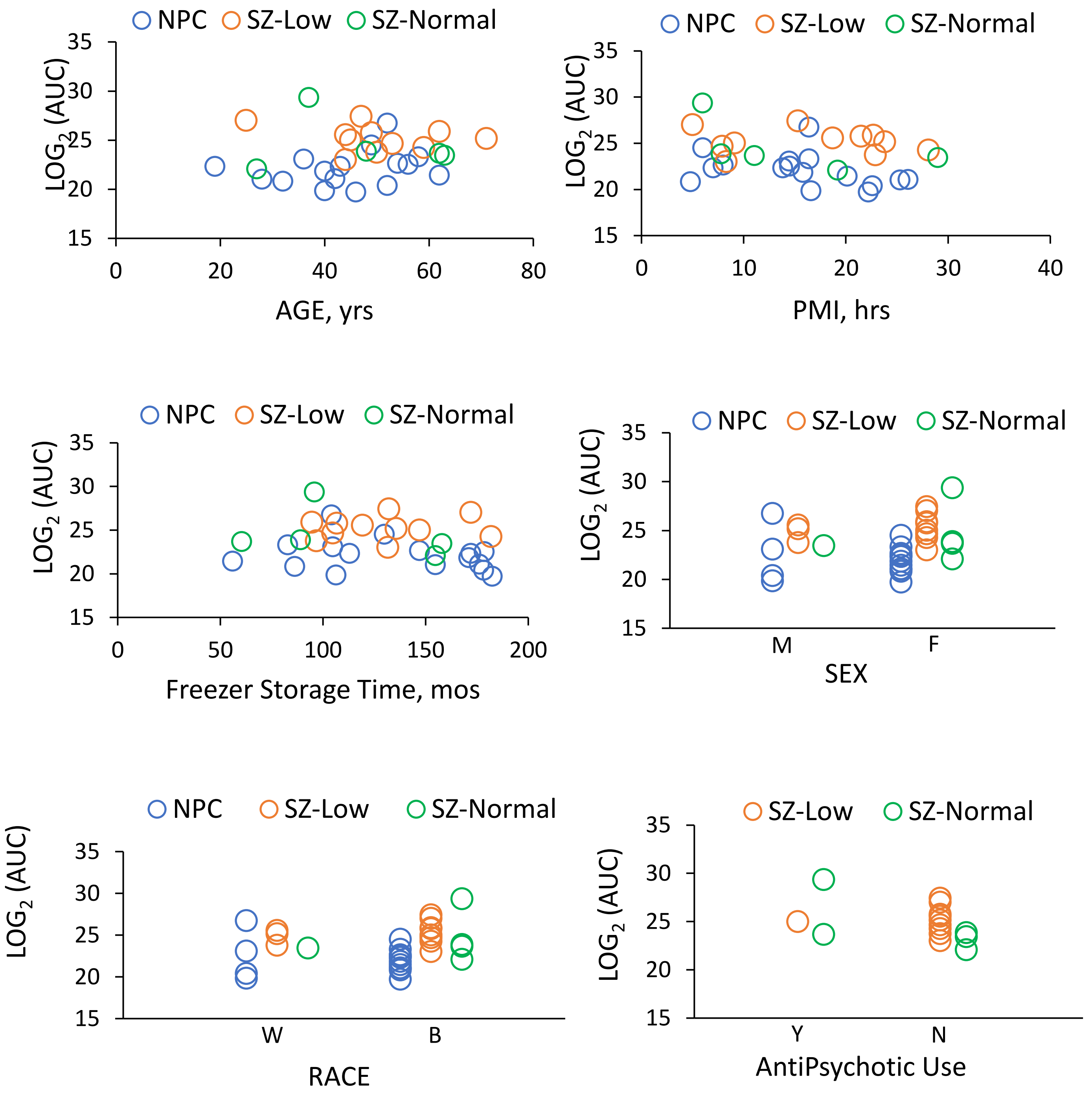

b.

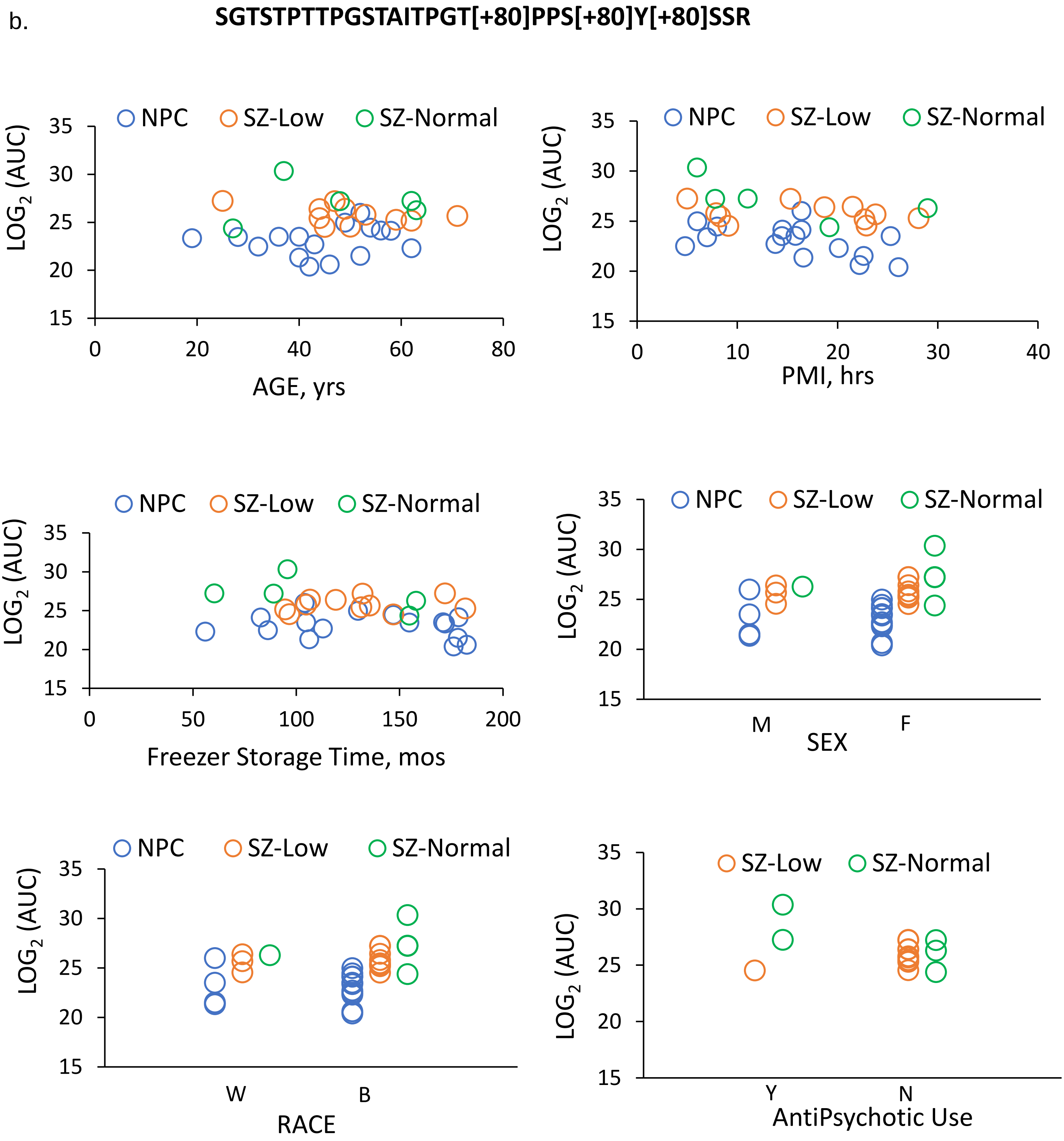


**Fig S4. Potential tissue and clinical confounds for the 2 most elevated MAP2 phosphopeptides.** (a) Association of age, PMI, freezer storage time, sex, race, and antipsychotic use at time of death for peptide VDHGAEIITQS[+80]PGR (b) Association of age, PMI, freezer storage time, sex, race, and antipsychotic use at time of death for peptide SGTSTPTTPGSTAITPGT[+80]PPS[+80]Y[+80]SSR
