## Supplementary material for "MAP2 is Differentially Phosphorylated in Schizophrenia, Altering its Function": Table S4

**Table S4.** Using cutoff criteria to define significance of q<.05 and fold change >0, 3,264 peptides representing 590 proteins were identified in a co-IP-MS assay performed on whole brain homogenate from wildtype C57/Bl6J mice. Data shown include accession number (column A), protein description (column B), modified peptide sequence (Column C), fold change (column D), the associated p-value (column E), corrected p-value (q value, Column F), and the number of significant peptides from that protein that were identified (Column G). Data are sorted in descending order of number of peptides identified for a protein. Note MAP2 is the protein with the most coverage, consistent with expected results from a MAP2 co-IP.

| Protein Accession | Protein Description | Modified Peptide Sequence | co-IP:Ctrl | p value | q value | Sig Peptide Count |
|---|---|---|---|---|---|---|
| P20357 | MTAP2_MOUSE | VSDFGQM[+16]ASGM[+16]NVDAGK | 5.57 | 4.27E-04 | 0.0143 | 128 |
| P20357 | MTAP2_MOUSE | M[+16]PC[+57]FPIESKEEEDKAEQAK | 8.89 | 4.40E-04 | 0.0143 | 128 |
| P20357 | MTAP2_MOUSE | APHWTSASLTEAAAHPHSPEM[+16]K | 7.63 | 7.50E-04 | 0.0143 | 128 |
| P20357 | MTAP2_MOUSE | ARVDHGAEIITQSPSR | 6.54 | 7.71E-04 | 0.0143 | 128 |
| P20357 | MTAP2_MOUSE | QDSFPISLEQAVTDAAM[+16]TSK | 7.62 | 7.92E-04 | 0.0143 | 128 |
| P20357 | MTAP2_MOUSE | M[+16]PSKPGEDFEHAALVPDTSK | 5.64 | 8.74E-04 | 0.0143 | 128 |
| P20357 | MTAP2_MOUSE | GEVQM[+16]EFIQLPK | 2.02 | 9.46E-04 | 0.0143 | 128 |
| P20357 | MTAP2_MOUSE | VDHGAEIITQSPSR | 6.34 | 9.49E-04 | 0.0143 | 128 |
| P20357 | MTAP2_MOUSE | M[+16]PC[+57]FPIESK | 5.34 | 1.04E-03 | 0.0143 | 128 |
| P20357 | MTAP2_MOUSE | FPSSFAEPLDK | 4.85 | 1.04E-03 | 0.0143 | 128 |
| P20357 | MTAP2_MOUSE | NGTVM[+16]APDLPEMLDLAGTR | 8.14 | 1.16E-03 | 0.0143 | 128 |
| P20357 | MTAP2_MOUSE | VSDFGQMASGM[+16]NVDAGK | 7.68 | 1.18E-03 | 0.0143 | 128 |
| P20357 | MTAP2_MOUSE | KETSAPSVQEPTLTETEPQTK | 6.88 | 1.28E-03 | 0.0143 | 128 |
| P20357 | MTAP2_MOUSE | DLATDLSLIEVK | 5.96 | 1.31E-03 | 0.0143 | 128 |
| P20357 | MTAP2_MOUSE | DWFIEM[+16]PTESK | 6.00 | 1.34E-03 | 0.0143 | 128 |
| P20357 | MTAP2_MOUSE | VSDFGQM[+16]ASGMNVDAGK | 6.29 | 1.36E-03 | 0.0143 | 128 |
| P20357 | MTAP2_MOUSE | ETSAPSVQEPTLTETEPQTK | 6.38 | 1.50E-03 | 0.0143 | 128 |
| P20357 | MTAP2_MOUSE | TPGTPGTPSYPR | 6.10 | 1.50E-03 | 0.0143 | 128 |
| P20357 | MTAP2_MOUSE | QFDSPM[+16]PSPFHGGSFTLPLDTM[+16]K | 5.11 | 1.51E-03 | 0.0143 | 128 |
| P20357 | MTAP2_MOUSE | SM[+16]SINLPM[+16]SC[+57]LDSIALGFNFGR | 8.94 | 1.57E-03 | 0.0143 | 128 |
| P20357 | MTAP2_MOUSE | QSTEPSIVM[+16]PSIGLSAEPPAPK | 5.97 | 1.68E-03 | 0.0143 | 128 |
| P20357 | MTAP2_MOUSE | LEGAGSATIAEVEM[+16]PFYEDK | 5.31 | 1.73E-03 | 0.0143 | 128 |
| P20357 | MTAP2_MOUSE | NKLEGAGSATIAEVEM[+16]PFYEDK | 6.83 | 1.75E-03 | 0.0143 | 128 |
| P20357 | MTAP2_MOUSE | SM[+16]SINLPMSC[+57]LDSIALGFNFGR | 7.44 | 1.77E-03 | 0.0143 | 128 |
| P20357 | MTAP2_MOUSE | FAAPAQPEEER | 6.08 | 1.79E-03 | 0.0143 | 128 |
| P20357 | MTAP2_MOUSE | STELGSDYYELSDSR | 6.54 | 1.84E-03 | 0.0143 | 128 |
| P20357 | MTAP2_MOUSE | DDKTGVIQTSTEQSFSK | 7.16 | 1.90E-03 | 0.0143 | 128 |
| P20357 | MTAP2_MOUSE | SGTSTPTTPGSTAITPGTPPSYSSR | 6.52 | 1.95E-03 | 0.0143 | 128 |
| P20357 | MTAP2_MOUSE | KIDLSHVTSK | 6.46 | 1.97E-03 | 0.0143 | 128 |
| P20357 | MTAP2_MOUSE | ANDKLDTVLEK | 5.43 | 2.02E-03 | 0.0143 | 128 |
| P20357 | MTAP2_MOUSE | FPSSFAEPLDKGEM[+16]EFK | 6.13 | 2.04E-03 | 0.0143 | 128 |
| P20357 | MTAP2_MOUSE | LILKPAIK | 6.50 | 2.06E-03 | 0.0143 | 128 |
| P20357 | MTAP2_MOUSE | YFETSALKEDM[+16]TR | 8.00 | 2.09E-03 | 0.0143 | 128 |
| P20357 | MTAP2_MOUSE | DGSPDAPATPEKEEVAFSEYK | 5.43 | 2.19E-03 | 0.0143 | 128 |
| P20357 | MTAP2_MOUSE | EIQGLFEEK | 6.04 | 2.26E-03 | 0.0143 | 128 |
| P20357 | MTAP2_MOUSE | KSEVQAHSPSR | 5.58 | 2.32E-03 | 0.0143 | 128 |
| P20357 | MTAP2_MOUSE | EKDVLEDIPR | 6.47 | 2.47E-03 | 0.0143 | 128 |
| P20357 | MTAP2_MOUSE | IGSTDNIK | 6.34 | 2.48E-03 | 0.0143 | 128 |
| P20357 | MTAP2_MOUSE | EEFVETC[+57]PGELK | 6.21 | 2.50E-03 | 0.0143 | 128 |
| P20357 | MTAP2_MOUSE | DEWGLAAPISPGPLTPMR | 4.44 | 2.50E-03 | 0.0143 | 128 |
| P20357 | MTAP2_MOUSE | DVLEDIPR | 5.59 | 2.53E-03 | 0.0143 | 128 |
| P20357 | MTAP2_MOUSE | QFDSPMPSPFHGGSFTLPLDTM[+16]K | 5.26 | 2.57E-03 | 0.0143 | 128 |
| P20357 | MTAP2_MOUSE | GQEHTIDELKQDSFPISLEQAVTDAAMTSK | 7.74 | 2.58E-03 | 0.0143 | 128 |
| P20357 | MTAP2_MOUSE | SQGTYSDTK | 6.09 | 2.66E-03 | 0.0143 | 128 |
| P20357 | MTAP2_MOUSE | TTAASGDLAQAPGAFK | 6.38 | 2.72E-03 | 0.0143 | 128 |
| P20357 | MTAP2_MOUSE | TGVIQTSTEQSFSK | 6.25 | 2.81E-03 | 0.0143 | 128 |
| P20357 | MTAP2_MOUSE | GLSSVPEVAEVEPTTK | 6.43 | 2.82E-03 | 0.0143 | 128 |
| P20357 | MTAP2_MOUSE | KDEWGLAAPISPGPLTPM[+16]R | 6.64 | 2.83E-03 | 0.0143 | 128 |
| P20357 | MTAP2_MOUSE | ESEEM[+16]GGKVELFGLGITYDQASTK | 6.83 | 2.85E-03 | 0.0143 | 128 |
| P20357 | MTAP2_MOUSE | NANGFPYREEEEGAFGEHR | 8.38 | 2.88E-03 | 0.0143 | 128 |
| P20357 | MTAP2_MOUSE | LINQPLPDLK | 6.14 | 2.90E-03 | 0.0143 | 128 |
| P20357 | MTAP2_MOUSE | VSLQDPSALATSK | 6.40 | 2.94E-03 | 0.0143 | 128 |
| P20357 | MTAP2_MOUSE | GGQVQIVTK | 6.29 | 2.95E-03 | 0.0143 | 128 |
| P20357 | MTAP2_MOUSE | ENGINGELTSADRETAEEVSAR | 7.09 | 3.18E-03 | 0.0143 | 128 |
| P20357 | MTAP2_MOUSE | DQGGAGEGLSR | 6.64 | 3.22E-03 | 0.0143 | 128 |
| P20357 | MTAP2_MOUSE | VNETEVK | 5.90 | 3.22E-03 | 0.0143 | 128 |
| P20357 | MTAP2_MOUSE | SGILVPSEK | 5.88 | 3.25E-03 | 0.0143 | 128 |
| P20357 | MTAP2_MOUSE | QFDSPM[+16]PSPFHGGSFTLPLDTMK | 4.86 | 3.32E-03 | 0.0143 | 128 |
| P20357 | MTAP2_MOUSE | KANDKLDTVLEK | 6.37 | 3.32E-03 | 0.0143 | 128 |
| P20357 | MTAP2_MOUSE | STVSIEEAVAK | 5.98 | 3.34E-03 | 0.0143 | 128 |
| P20357 | MTAP2_MOUSE | VTSEPEAVSER | 6.12 | 3.35E-03 | 0.0143 | 128 |
| P20357 | MTAP2_MOUSE | MPC[+57]FPIESK | 6.39 | 3.63E-03 | 0.0146 | 128 |

| Protein Accession | Protein Description | Modified Peptide Sequence | co-IP:Ctrl | p value | q value | Sig Peptide Count |
|---|---|---|---|---|---|---|
| P20357 | MTAP2_MOUSE | MPSKPGEDFEHAALVPDTSK | 7.22 | 3.77E-03 | 0.0147 | 128 |
| P20357 | MTAP2_MOUSE | ETSPETSLIQDEVALK | 7.23 | 3.78E-03 | 0.0147 | 128 |
| P20357 | MTAP2_MOUSE | QFDSPMPSPFHGGSFTLPLDTMK | 8.06 | 3.80E-03 | 0.0147 | 128 |
| P20357 | MTAP2_MOUSE | EENSFSLNSSISSAR | 6.42 | 3.83E-03 | 0.0147 | 128 |
| P20357 | MTAP2_MOUSE | ENGINGELTSADR | 4.88 | 3.84E-03 | 0.0147 | 128 |
| P20357 | MTAP2_MOUSE | MPC[+57]FPIESKEEEDKAEQAK | 7.79 | 3.89E-03 | 0.0147 | 128 |
| P20357 | MTAP2_MOUSE | FEVAQELTLSSEAPQEADSFM[+16]GVESGHIK | 6.20 | 4.01E-03 | 0.0148 | 128 |
| P20357 | MTAP2_MOUSE | KQSTEPSIVM[+16]PSIGLSAEPPAPK | 6.10 | 4.09E-03 | 0.0148 | 128 |
| P20357 | MTAP2_MOUSE | FPSSFAEPLDKGEMEFK | 5.91 | 4.24E-03 | 0.0149 | 128 |
| P20357 | MTAP2_MOUSE | NGTVMAPDLPEMLDLAGTR | 8.95 | 4.30E-03 | 0.0150 | 128 |
| P20357 | MTAP2_MOUSE | VSDFGQMASGMNVDAGK | 7.57 | 4.32E-03 | 0.0150 | 128 |
| P20357 | MTAP2_MOUSE | DLQGMEGEKLPPVPFAQTFGTNLEDR | 4.49 | 4.54E-03 | 0.0152 | 128 |
| P20357 | MTAP2_MOUSE | VSEGPRPFAPVFFQSDDK | 4.63 | 4.55E-03 | 0.0152 | 128 |
| P20357 | MTAP2_MOUSE | YFETSALKEDMTR | 6.73 | 4.82E-03 | 0.0154 | 128 |
| P20357 | MTAP2_MOUSE | SILTEQLETIPK | 6.48 | 4.83E-03 | 0.0154 | 128 |
| P20357 | MTAP2_MOUSE | YTVPLPSPVQDSENLSGESGSFYEGTDDK | 6.96 | 4.98E-03 | 0.0155 | 128 |
| P20357 | MTAP2_MOUSE | ETAEEVSAR | 5.83 | 5.04E-03 | 0.0155 | 128 |
| P20357 | MTAP2_MOUSE | VTGGQTIQVETSSESPFPAK | 6.19 | 5.09E-03 | 0.0155 | 128 |
| P20357 | MTAP2_MOUSE | KLILKPAIK | 7.52 | 5.14E-03 | 0.0155 | 128 |
| P20357 | MTAP2_MOUSE | RPRPHDEELEIEM[+16]AAEAQAEPK | 2.89 | 5.69E-03 | 0.0161 | 128 |
| P20357 | MTAP2_MOUSE | KTTAASGDLAQAPGAFK | 3.96 | 5.70E-03 | 0.0161 | 128 |
| P20357 | MTAP2_MOUSE | SMSINLPM[+16]SC[+57]LDSIALGFNFGR | 9.09 | 5.93E-03 | 0.0163 | 128 |
| P20357 | MTAP2_MOUSE | SILTEQLETIPKEER | 6.44 | 6.01E-03 | 0.0164 | 128 |
| P20357 | MTAP2_MOUSE | GEVQMEFIQLPK | 5.95 | 6.52E-03 | 0.0171 | 128 |
| P20357 | MTAP2_MOUSE | SDTLQISDLLVSESREEFVETC[+57]PGELK | 3.26 | 6.61E-03 | 0.0171 | 128 |
| P20357 | MTAP2_MOUSE | LPPVPFAQTFGTNLEDR | 7.34 | 6.66E-03 | 0.0172 | 128 |
| P20357 | MTAP2_MOUSE | RDLATDLSLIEVK | 8.49 | 6.67E-03 | 0.0172 | 128 |
| P20357 | MTAP2_MOUSE | DLQGM[+16]EGEKLPPVPFAQTFGTNLEDRK | 4.64 | 6.94E-03 | 0.0174 | 128 |
| P20357 | MTAP2_MOUSE | DWFIEMPTESK | 6.69 | 6.97E-03 | 0.0175 | 128 |
| P20357 | MTAP2_MOUSE | VTTDQEKK | 5.52 | 7.03E-03 | 0.0175 | 128 |
| P20357 | MTAP2_MOUSE | SDTLQISDLLVSESR | 6.10 | 7.26E-03 | 0.0177 | 128 |
| P20357 | MTAP2_MOUSE | LEGAGSATIAEVEMPFYEDK | 7.70 | 8.06E-03 | 0.0184 | 128 |
| P20357 | MTAP2_MOUSE | IVQVVTAEAVAVLK | 6.09 | 8.48E-03 | 0.0189 | 128 |
| P20357 | MTAP2_MOUSE | LRDDKTGVIQTSTEQSFSK | 2.43 | 8.48E-03 | 0.0189 | 128 |
| P20357 | MTAP2_MOUSE | AEPSQLDIK | 7.29 | 8.64E-03 | 0.0191 | 128 |
| P20357 | MTAP2_MOUSE | KDEWGLAAPISPGPLTPMR | 7.45 | 9.01E-03 | 0.0196 | 128 |
| P20357 | MTAP2_MOUSE | VELFGLGITYDQASTK | 6.36 | 9.08E-03 | 0.0197 | 128 |
| P20357 | MTAP2_MOUSE | VNETEVKEK | 5.41 | 9.12E-03 | 0.0197 | 128 |
| P20357 | MTAP2_MOUSE | NKLEGAGSATIAEVEMPFYEDK | 3.70 | 9.54E-03 | 0.0202 | 128 |
| P20357 | MTAP2_MOUSE | RGVSGDREENSFSLNSSISSAR | 8.79 | 1.04E-02 | 0.0211 | 128 |
| P20357 | MTAP2_MOUSE | YTRPTHLSC[+57]VK | 9.65 | 1.06E-02 | 0.0213 | 128 |
| P20357 | MTAP2_MOUSE | VGSLDNAHHVPGGGNVK | 5.82 | 1.17E-02 | 0.0225 | 128 |
| P20357 | MTAP2_MOUSE | ESEEMGGKVELFGLGITYDQASTK | 5.64 | 1.23E-02 | 0.0231 | 128 |
| P20357 | MTAP2_MOUSE | REIQGLFEEK | 1.64 | 1.28E-02 | 0.0235 | 128 |
| P20357 | MTAP2_MOUSE | NQHDEKELQAK | 8.00 | 1.30E-02 | 0.0238 | 128 |
| P20357 | MTAP2_MOUSE | SSLPRPSSILPPR | 7.89 | 1.40E-02 | 0.0249 | 128 |
| P20357 | MTAP2_MOUSE | GQEHTIDELK | 2.74 | 1.40E-02 | 0.0249 | 128 |
| P20357 | MTAP2_MOUSE | RPRPHDEELEIEMAAEAQAEPK | 5.12 | 1.44E-02 | 0.0254 | 128 |
| P20357 | MTAP2_MOUSE | VADVSISEVTTLLGNVHSPVVEGYVGENISGEVK | 3.37 | 1.47E-02 | 0.0257 | 128 |
| P20357 | MTAP2_MOUSE | GVSGDREENSFSLNSSISSAR | 6.00 | 1.49E-02 | 0.0259 | 128 |
| P20357 | MTAP2_MOUSE | DEWGLAAPISPGPLTPM[+16]R | 5.93 | 1.52E-02 | 0.0263 | 128 |
| P20357 | MTAP2_MOUSE | LPPVPFAQTFGTNLEDRK | 3.37 | 1.53E-02 | 0.0264 | 128 |
| P20357 | MTAP2_MOUSE | LSVEIPC[+57]PPPVSEADLSTDEK | 2.86 | 1.56E-02 | 0.0267 | 128 |
| P20357 | MTAP2_MOUSE | KQSTEPSIVMPSIGLSAEPPAPK | 6.08 | 1.60E-02 | 0.0271 | 128 |
| P20357 | MTAP2_MOUSE | FEVAQELTLSSEAPQEADSFMGVESGHIK | 6.27 | 1.61E-02 | 0.0272 | 128 |
| P20357 | MTAP2_MOUSE | DDLTLSR | 2.43 | 1.65E-02 | 0.0276 | 128 |
| P20357 | MTAP2_MOUSE | QDSFPISLEQAVTDAAMTSK | 8.85 | 1.69E-02 | 0.0281 | 128 |
| P47911 | RL6_MOUSE 60 | SSITPGTVLIILTGR | 4.34 | 1.78E-02 | 0.0290 | 128 |
| P52480 | KPYM_MOUSE | FGVEQDVDMVFASFIR | 9.35 | 2.06E-02 | 0.0323 | 128 |
| O08638 | MYH11_MOUSE | SLEADLM[+16]QLQEDLAAAER | 5.62 | 2.09E-02 | 0.0326 | 128 |
| P16546 | SPTN1_MOUSE | KVEDLFLTFAK | 3.36 | 2.12E-02 | 0.0330 | 128 |
| Q8VDM4 | PSMD2_MOUSE | MNLASSFVNGFVNAAFGQDK | 7.90 | 2.32E-02 | 0.0352 | 128 |

| Protein Accession | Protein Description | Modified Peptide Sequence | co-IP:Ctrl | p value | q value | Sig Peptide Count |
|---|---|---|---|---|---|---|
| E9Q912 | E9Q912_MOUSE | SVAQQASLTEQR | 4.19 | 2.38E-02 | 0.0359 | 128 |
| Q9D0F9 | PGM1_MOUSE | TIEEYAIC[+57]PDLK | 5.62 | 2.51E-02 | 0.0372 | 128 |
| P80314 | TCPB_MOUSE | LAVEAVLR | 5.61 | 3.58E-02 | 0.0494 | 128 |
| Q8VDD5 | MYH9_MOUSE | LQLQEQLQAETELC[+57]AEAEELR | 5.64 | 2.09E-02 | 0.0326 | 128 |
| Q61879 | MYH10_MOUSE | EEELQGALAR | 3.03 | 9.82E-05 | 0.0143 | 125 |
| Q61879 | MYH10_MOUSE | NKQEVM[+16]ISDLEER | 4.68 | 1.61E-04 | 0.0143 | 125 |
| Q61879 | MYH10_MOUSE | NTDQASM[+16]PENTVAQK | 2.66 | 3.10E-04 | 0.0143 | 125 |
| Q61879 | MYH10_MOUSE | ADM[+16]EDLM[+16]SSKDDVGK | 3.78 | 5.13E-04 | 0.0143 | 125 |
| Q61879 | MYH10_MOUSE | NILAEQLQAETELFAEAEEM[+16]R | 3.44 | 5.34E-04 | 0.0143 | 125 |
| Q61879 | MYH10_MOUSE | VEGELEEM[+16]ER | 3.94 | 5.35E-04 | 0.0143 | 125 |
| Q61879 | MYH10_MOUSE | ALEDETKNHEAQIQDM[+16]R | 5.25 | 5.54E-04 | 0.0143 | 125 |
| Q61879 | MYH10_MOUSE | VEEEEERNQILQNEK | 4.12 | 8.28E-04 | 0.0143 | 125 |
| Q61879 | MYH10_MOUSE | QGLETDNKELAC[+57]EVK | 4.15 | 8.45E-04 | 0.0143 | 125 |
| Q61879 | MYH10_MOUSE | ADM[+16]EDLM[+16]SSK | 5.24 | 8.53E-04 | 0.0143 | 125 |
| Q61879 | MYH10_MOUSE | C[+57]M[+16]LQDREDQSILC[+57]TGESGAGK | 5.09 | 9.29E-04 | 0.0143 | 125 |
| Q61879 | MYH10_MOUSE | ELDDATEANEGLSR | 4.33 | 9.35E-04 | 0.0143 | 125 |
| Q61879 | MYH10_MOUSE | LDAQVQELHAK | 4.24 | 9.59E-04 | 0.0143 | 125 |
| Q61879 | MYH10_MOUSE | KEEELQGALAR | 4.07 | 9.78E-04 | 0.0143 | 125 |
| Q61879 | MYH10_MOUSE | M[+16]EEEVLLLEDQNSK | 4.53 | 1.03E-03 | 0.0143 | 125 |
| Q61879 | MYH10_MOUSE | MEIDLKDLEAQIEAANK | 5.40 | 1.04E-03 | 0.0143 | 125 |
| Q61879 | MYH10_MOUSE | ELQAQIAELQEDFESEK | 4.63 | 1.07E-03 | 0.0143 | 125 |
| Q61879 | MYH10_MOUSE | TSDVNDTQPPQSE | 4.65 | 1.08E-03 | 0.0143 | 125 |
| Q61879 | MYH10_MOUSE | QLEEEKNSLQEQQEEEEEARK | 5.93 | 1.09E-03 | 0.0143 | 125 |
| Q61879 | MYH10_MOUSE | DAAGLESQLQDTQELLQEETR | 4.62 | 1.10E-03 | 0.0143 | 125 |
| Q61879 | MYH10_MOUSE | IAEC[+57]SSQLAEEEEK | 4.96 | 1.13E-03 | 0.0143 | 125 |
| Q61879 | MYH10_MOUSE | QLEEAEEEATR | 4.21 | 1.14E-03 | 0.0143 | 125 |
| Q61879 | MYH10_MOUSE | IAQLEEELEEEQSNM[+16]ELLNDR | 3.79 | 1.14E-03 | 0.0143 | 125 |
| Q61879 | MYH10_MOUSE | KLDAQVQELHAK | 4.12 | 1.14E-03 | 0.0143 | 125 |
| Q61879 | MYH10_MOUSE | LQNELDNVSTLLEEAEK | 4.48 | 1.16E-03 | 0.0143 | 125 |
| Q61879 | MYH10_MOUSE | SLEAEILQLQEELASSER | 6.55 | 1.19E-03 | 0.0143 | 125 |
| Q61879 | MYH10_MOUSE | NLPIYSENIIEM[+16]YR | 4.89 | 1.22E-03 | 0.0143 | 125 |
| Q61879 | MYH10_MOUSE | ASRDEIFAQSK | 3.91 | 1.24E-03 | 0.0143 | 125 |
| Q61879 | MYH10_MOUSE | HGFEAASIKEER | 4.69 | 1.35E-03 | 0.0143 | 125 |
| Q61879 | MYH10_MOUSE | FLSNGYIPIPGQQDK | 4.74 | 1.40E-03 | 0.0143 | 125 |
| Q61879 | MYH10_MOUSE | QLEEEKNSLQEQQEEEEEAR | 5.12 | 1.47E-03 | 0.0143 | 125 |
| Q61879 | MYH10_MOUSE | GGPISFSSSR | 4.16 | 1.51E-03 | 0.0143 | 125 |
| Q61879 | MYH10_MOUSE | M[+16]EIDLKDLEAQIEAANK | 4.51 | 1.51E-03 | 0.0143 | 125 |
| Q61879 | MYH10_MOUSE | DLSEELEALK | 4.63 | 1.54E-03 | 0.0143 | 125 |
| Q61879 | MYH10_MOUSE | QIVSNLEK | 4.58 | 1.57E-03 | 0.0143 | 125 |
| Q61879 | MYH10_MOUSE | VEEEEERNQILQNEKK | 5.68 | 1.58E-03 | 0.0143 | 125 |
| Q61879 | MYH10_MOUSE | ATISALEAK | 4.02 | 1.58E-03 | 0.0143 | 125 |
| Q61879 | MYH10_MOUSE | KM[+16]EEEVLLLEDQNSK | 3.54 | 1.76E-03 | 0.0143 | 125 |
| Q61879 | MYH10_MOUSE | NC[+57]AAYLK | 4.77 | 1.84E-03 | 0.0143 | 125 |
| Q61879 | MYH10_MOUSE | HEM[+16]PPHIYAISESAYR | 5.05 | 1.84E-03 | 0.0143 | 125 |
| Q61879 | MYH10_MOUSE | IGQLEEQLEQEAK | 4.77 | 1.86E-03 | 0.0143 | 125 |
| Q61879 | MYH10_MOUSE | TGLEDPER | 4.26 | 1.88E-03 | 0.0143 | 125 |
| Q61879 | MYH10_MOUSE | LQELEGAVK | 3.78 | 1.93E-03 | 0.0143 | 125 |
| Q61879 | MYH10_MOUSE | ELEAELEDER | 4.29 | 1.94E-03 | 0.0143 | 125 |
| Q61879 | MYH10_MOUSE | KLDGETTDLQDQIAELQAQVDELK | 6.09 | 1.94E-03 | 0.0143 | 125 |
| Q61879 | MYH10_MOUSE | NSLQEQQEEEEEAR | 5.00 | 1.95E-03 | 0.0143 | 125 |
| Q61879 | MYH10_MOUSE | HAEQERDELADEIANSASGK | 4.70 | 1.95E-03 | 0.0143 | 125 |
| Q61879 | MYH10_MOUSE | ALEQQVEEMR | 4.43 | 2.01E-03 | 0.0143 | 125 |
| Q61879 | MYH10_MOUSE | DVEALSQR | 3.62 | 2.02E-03 | 0.0143 | 125 |
| Q61879 | MYH10_MOUSE | DEIFAQSK | 4.16 | 2.05E-03 | 0.0143 | 125 |
| Q61879 | MYH10_MOUSE | VLAYDKLEK | 4.31 | 2.18E-03 | 0.0143 | 125 |
| Q61879 | MYH10_MOUSE | ALEEALEAKEEFER | 4.70 | 2.19E-03 | 0.0143 | 125 |
| Q61879 | MYH10_MOUSE | ALELDPNLYR | 4.76 | 2.22E-03 | 0.0143 | 125 |
| Q61879 | MYH10_MOUSE | NM[+16]DPLNDNVATLLHQSSDR | 4.44 | 2.28E-03 | 0.0143 | 125 |
| Q61879 | MYH10_MOUSE | YEILTPNAIPK | 4.57 | 2.32E-03 | 0.0143 | 125 |
| Q61879 | MYH10_MOUSE | LC[+57]HLLGMNVM[+16]EFTR | 4.61 | 2.46E-03 | 0.0143 | 125 |
| Q61879 | MYH10_MOUSE | NTDQASMPENTVAQK | 5.31 | 2.46E-03 | 0.0143 | 125 |
| Q61879 | MYH10_MOUSE | VDDDLGTIESLEEAK | 3.57 | 2.50E-03 | 0.0143 | 125 |

| Protein Accession | Protein Description | Modified Peptide Sequence | co-IP:Ctrl | p value | q value | Sig Peptide Count |
|---|---|---|---|---|---|---|
| Q61879 | MYH10_MOUSE | M[+16]QAHIQDLEEQLDEEEGAR | 4.23 | 2.51E-03 | 0.0143 | 125 |
| Q61879 | MYH10_MOUSE | QELEEILHDLESR | 6.86 | 2.57E-03 | 0.0143 | 125 |
| Q61879 | MYH10_MOUSE | KMQAHIQDLEEQLDEEEGAR | 5.05 | 2.58E-03 | 0.0143 | 125 |
| Q61879 | MYH10_MOUSE | QVLALQSQLADTK | 4.44 | 2.72E-03 | 0.0143 | 125 |
| Q61879 | MYH10_MOUSE | TELEDTLDTTAAQQELR | 5.79 | 2.72E-03 | 0.0143 | 125 |
| Q61879 | MYH10_MOUSE | ALEEALEAK | 3.97 | 2.76E-03 | 0.0143 | 125 |
| Q61879 | MYH10_MOUSE | NLPIYSENIIEMYR | 5.75 | 2.79E-03 | 0.0143 | 125 |
| Q61879 | MYH10_MOUSE | KLVWIPSER | 4.17 | 2.80E-03 | 0.0143 | 125 |
| Q61879 | MYH10_MOUSE | KKVDDDLGTIESLEEAK | 4.65 | 2.90E-03 | 0.0143 | 125 |
| Q61879 | MYH10_MOUSE | QQQLSALK | 4.56 | 2.99E-03 | 0.0143 | 125 |
| Q61879 | MYH10_MOUSE | ELEAELEDERK | 4.60 | 3.02E-03 | 0.0143 | 125 |
| Q61879 | MYH10_MOUSE | SALLDEKR | 4.31 | 3.02E-03 | 0.0143 | 125 |
| Q61879 | MYH10_MOUSE | KVDDDLGTIESLEEAKK | 5.04 | 3.11E-03 | 0.0143 | 125 |
| Q61879 | MYH10_MOUSE | IVFQEFR | 4.58 | 3.16E-03 | 0.0143 | 125 |
| Q61879 | MYH10_MOUSE | IAQLEEELEEEQSNMELLNDR | 5.06 | 3.28E-03 | 0.0143 | 125 |
| Q61879 | MYH10_MOUSE | IVGLDQVTGMTETAFGSAYK | 5.47 | 3.32E-03 | 0.0143 | 125 |
| Q61879 | MYH10_MOUSE | TTLQVDTLNTELAAER | 4.73 | 3.38E-03 | 0.0143 | 125 |
| Q61879 | MYH10_MOUSE | HATALEELSEQLEQAK | 4.97 | 3.54E-03 | 0.0144 | 125 |
| Q61879 | MYH10_MOUSE | EQADFAVEALAK | 4.53 | 3.56E-03 | 0.0145 | 125 |
| Q61879 | MYH10_MOUSE | QEEELQAKDEELLK | 5.29 | 3.61E-03 | 0.0145 | 125 |
| Q61879 | MYH10_MOUSE | KVDDDLGTIESLEEAK | 4.70 | 3.65E-03 | 0.0146 | 125 |
| Q61879 | MYH10_MOUSE | NILAEQLQAETELFAEAEEMR | 5.16 | 3.69E-03 | 0.0147 | 125 |
| Q61879 | MYH10_MOUSE | SDLLLEGFNNYR | 3.60 | 3.72E-03 | 0.0147 | 125 |
| Q61879 | MYH10_MOUSE | AVIYNPATQADWTAK | 4.35 | 3.72E-03 | 0.0147 | 125 |
| Q61879 | MYH10_MOUSE | LC[+57]HLLGM[+16]NVM[+16]EFTR | 3.70 | 3.76E-03 | 0.0147 | 125 |
| Q61879 | MYH10_MOUSE | KQQQLSALK | 4.86 | 4.02E-03 | 0.0148 | 125 |
| Q61879 | MYH10_MOUSE | DELADEIANSASGK | 4.29 | 4.14E-03 | 0.0148 | 125 |
| Q61879 | MYH10_MOUSE | AMVNKDDIQK | 4.26 | 4.23E-03 | 0.0149 | 125 |
| Q61879 | MYH10_MOUSE | QLHIEGASLELSDDDTESK | 4.37 | 4.27E-03 | 0.0150 | 125 |
| Q61879 | MYH10_MOUSE | HEMPPHIYAISESAYR | 4.22 | 4.37E-03 | 0.0150 | 125 |
| Q61879 | MYH10_MOUSE | VEDMAELTC[+57]LNEASVLHNLKDR | 4.91 | 5.12E-03 | 0.0155 | 125 |
| Q61879 | MYH10_MOUSE | IVGLDQVTGM[+16]TETAFGSAYK | 3.80 | 5.17E-03 | 0.0155 | 125 |
| Q61879 | MYH10_MOUSE | GDEVMVELAENGKK | 6.76 | 5.29E-03 | 0.0158 | 125 |
| Q61879 | MYH10_MOUSE | KMEEEVLLLEDQNSK | 4.59 | 5.30E-03 | 0.0158 | 125 |
| Q61879 | MYH10_MOUSE | DLEAQIEAANK | 4.32 | 5.48E-03 | 0.0159 | 125 |
| Q61879 | MYH10_MOUSE | AM[+16]VNKDDIQK | 3.69 | 5.49E-03 | 0.0159 | 125 |
| Q61879 | MYH10_MOUSE | DHNIPGELER | 4.46 | 5.84E-03 | 0.0163 | 125 |
| Q61879 | MYH10_MOUSE | RQLEEAEEEATR | 5.16 | 5.89E-03 | 0.0163 | 125 |
| Q61879 | MYH10_MOUSE | VVSSVLQFGNISFK | 5.07 | 6.15E-03 | 0.0166 | 125 |
| Q61879 | MYH10_MOUSE | VEGELEEMER | 3.90 | 6.19E-03 | 0.0167 | 125 |
| Q61879 | MYH10_MOUSE | HATALEELSEQLEQAKR | 5.49 | 6.58E-03 | 0.0171 | 125 |
| Q61879 | MYH10_MOUSE | LDGETTDLQDQIAELQAQVDELK | 5.69 | 6.70E-03 | 0.0172 | 125 |
| Q61879 | MYH10_MOUSE | QLLQANPILESFGNAK | 5.09 | 7.02E-03 | 0.0175 | 125 |
| Q61879 | MYH10_MOUSE | LEVNMQAM[+16]K | 3.09 | 7.25E-03 | 0.0177 | 125 |
| Q61879 | MYH10_MOUSE | KMEIDLKDLEAQIEAANK | 6.78 | 7.29E-03 | 0.0177 | 125 |
| Q61879 | MYH10_MOUSE | LEVNMQAMK | 4.03 | 7.55E-03 | 0.0180 | 125 |
| Q61879 | MYH10_MOUSE | NKQEVMISDLEER | 4.43 | 7.71E-03 | 0.0181 | 125 |
| Q61879 | MYH10_MOUSE | MEEEVLLLEDQNSK | 4.22 | 7.80E-03 | 0.0182 | 125 |
| Q61879 | MYH10_MOUSE | LC[+57]HLLGM[+16]NVMEFTR | 5.48 | 7.89E-03 | 0.0183 | 125 |
| Q61879 | MYH10_MOUSE | LQNELDNVSTLLEEAEKK | 7.24 | 7.91E-03 | 0.0183 | 125 |
| Q61879 | MYH10_MOUSE | HGFEAASIK | 4.47 | 8.11E-03 | 0.0184 | 125 |
| Q61879 | MYH10_MOUSE | ADMEDLMSSKDDVGK | 5.66 | 8.22E-03 | 0.0186 | 125 |
| Q61879 | MYH10_MOUSE | ADMEDLMSSK | 4.18 | 8.27E-03 | 0.0187 | 125 |
| Q61879 | MYH10_MOUSE | GDEVMVELAENGK | 3.83 | 1.03E-02 | 0.0209 | 125 |
| Q61879 | MYH10_MOUSE | LEVNM[+16]QAM[+16]K | 3.08 | 1.03E-02 | 0.0209 | 125 |
| Q61879 | MYH10_MOUSE | NSLQEQQEEEEARK | 7.88 | 1.23E-02 | 0.0231 | 125 |
| Q61879 | MYH10_MOUSE | LC[+57]HLLGMNVMEFTR | 6.01 | 1.27E-02 | 0.0234 | 125 |
| Q61879 | MYH10_MOUSE | TFHIFYQLLSGAGEHLK | 6.48 | 1.32E-02 | 0.0240 | 125 |
| Q61879 | MYH10_MOUSE | QEVM[+16]ISDLEER | 2.39 | 1.37E-02 | 0.0246 | 125 |
| Q61879 | MYH10_MOUSE | ITDIIIFFQAVC[+57]R | 5.71 | 1.39E-02 | 0.0248 | 125 |
| Q61879 | MYH10_MOUSE | KTTLQVDTLNTELAAER | 5.03 | 1.40E-02 | 0.0249 | 125 |
| P14873 | MAP1B_MOUSE | SPPLLGSESPYEDFLSADSK | 4.69 | 2.12E-02 | 0.0330 | 125 |

| Protein Accession | Protein Description | Modified Peptide Sequence | co-IP:Ctrl | p value | q value | Sig Peptide Count |
|---|---|---|---|---|---|---|
| Q62261 | SPTB2_MOUSE | HLLGVEDLLQK | 5.30 | 2.24E-02 | 0.0344 | 125 |
| Q99104 | MYO5A_MOUSE | NTMTDSTILLEDVQK | 4.92 | 2.43E-02 | 0.0364 | 125 |
| P16546 | SPTN1_MOUSE | GVIDM[+16]GNSLIER | 2.92 | 2.50E-02 | 0.0371 | 125 |
| A2AJI0 | MA7D1_MOUSE | SLQLSAWESSIVDR | 5.05 | 2.67E-02 | 0.0390 | 125 |
| Q8K1M6 | DNM1L_MOUSE | ALQGASQIIAEIR | 4.82 | 3.10E-02 | 0.0437 | 125 |
| Q8VDD5 | MYH9_MOUSE | QIATLHAQVTDM[+16]KK | 4.31 | 4.81E-04 | 0.0143 | 117 |
| Q8VDD5 | MYH9_MOUSE | DLGEELEALK | 4.29 | 5.23E-04 | 0.0143 | 117 |
| Q8VDD5 | MYH9_MOUSE | ALEQQVEEM[+16]KTQLEELEDELQATEDAK | 7.52 | 7.06E-04 | 0.0143 | 117 |
| Q8VDD5 | MYH9_MOUSE | KLEEDQIIM[+16]EDQNC[+57]K | 4.23 | 7.18E-04 | 0.0143 | 117 |
| Q8VDD5 | MYH9_MOUSE | TEM[+16]EDLM[+16]SSKDDVGK | 3.39 | 9.94E-04 | 0.0143 | 117 |
| Q8VDD5 | MYH9_MOUSE | QAQQERDELADEIANSSGK | 4.11 | 1.09E-03 | 0.0143 | 117 |
| Q8VDD5 | MYH9_MOUSE | KM[+16]EDGVGC[+57]LETAEEAK | 4.04 | 1.12E-03 | 0.0143 | 117 |
| Q8VDD5 | MYH9_MOUSE | IIGLDQVAGM[+16]SETALPGAFK | 3.79 | 1.14E-03 | 0.0143 | 117 |
| Q8VDD5 | MYH9_MOUSE | LQEM[+16]ESAVK | 3.76 | 1.22E-03 | 0.0143 | 117 |
| Q8VDD5 | MYH9_MOUSE | ELEDATETADAM[+16]NR | 3.88 | 1.23E-03 | 0.0143 | 117 |
| Q8VDD5 | MYH9_MOUSE | KM[+16]QQNIQELEEQLEEEESAR | 4.54 | 1.25E-03 | 0.0143 | 117 |
| Q8VDD5 | MYH9_MOUSE | SM[+16]M[+16]QDREDQSILC[+57]TGESGAGK | 4.31 | 1.28E-03 | 0.0143 | 117 |
| Q8VDD5 | MYH9_MOUSE | QLLQANPILEAFGNAK | 4.48 | 1.31E-03 | 0.0143 | 117 |
| Q8VDD5 | MYH9_MOUSE | LQQELDDLLVDLDHQR | 5.26 | 1.31E-03 | 0.0143 | 117 |
| Q8VDD5 | MYH9_MOUSE | IM[+16]GIPEDEQM[+16]GLLR | 4.28 | 1.32E-03 | 0.0143 | 117 |
| Q8VDD5 | MYH9_MOUSE | DFSALESQLQDTQELLQEENR | 4.71 | 1.45E-03 | 0.0143 | 117 |
| Q8VDD5 | MYH9_MOUSE | SM[+16]EAEMIQLQEELAAAER | 4.31 | 1.48E-03 | 0.0143 | 117 |
| Q8VDD5 | MYH9_MOUSE | VAEFTTNLM[+16]EEEEK | 3.61 | 1.50E-03 | 0.0143 | 117 |
| Q8VDD5 | MYH9_MOUSE | KVEAQLQELQVK | 3.29 | 1.57E-03 | 0.0143 | 117 |
| Q8VDD5 | MYH9_MOUSE | EDQSILC[+57]TGESGAGK | 4.49 | 1.57E-03 | 0.0143 | 117 |
| Q8VDD5 | MYH9_MOUSE | ALELDSNLYR | 4.11 | 1.64E-03 | 0.0143 | 117 |
| Q8VDD5 | MYH9_MOUSE | VISGVLQLGNIAFK | 4.56 | 1.64E-03 | 0.0143 | 117 |
| Q8VDD5 | MYH9_MOUSE | LEVNLQAM[+16]K | 3.98 | 1.70E-03 | 0.0143 | 117 |
| Q8VDD5 | MYH9_MOUSE | NC[+57]AAYLR | 4.65 | 1.70E-03 | 0.0143 | 117 |
| Q8VDD5 | MYH9_MOUSE | FDQLLAEEK | 3.65 | 1.72E-03 | 0.0143 | 117 |
| Q8VDD5 | MYH9_MOUSE | VEDM[+16]AELTC[+57]LNEASVLHNLK | 2.87 | 1.74E-03 | 0.0143 | 117 |
| Q8VDD5 | MYH9_MOUSE | NTNPNFVR | 4.41 | 1.76E-03 | 0.0143 | 117 |
| Q8VDD5 | MYH9_MOUSE | MQQNIQELEEQLEEEESAR | 4.93 | 1.77E-03 | 0.0143 | 117 |
| Q8VDD5 | MYH9_MOUSE | ALEQQVEEM[+16]K | 3.39 | 1.78E-03 | 0.0143 | 117 |
| Q8VDD5 | MYH9_MOUSE | HSQAVEELADQLEQTKR | 4.67 | 1.79E-03 | 0.0143 | 117 |
| Q8VDD5 | MYH9_MOUSE | QTLENERGELANEVK | 5.95 | 1.79E-03 | 0.0143 | 117 |
| Q8VDD5 | MYH9_MOUSE | DVLLQVEDERR | 4.84 | 1.83E-03 | 0.0143 | 117 |
| Q8VDD5 | MYH9_MOUSE | C[+57]NGVLEGIR | 4.00 | 1.86E-03 | 0.0143 | 117 |
| Q8VDD5 | MYH9_MOUSE | VIQYLAHVASSHK | 4.57 | 1.86E-03 | 0.0143 | 117 |
| Q8VDD5 | MYH9_MOUSE | KANLQIDQINTDLNLER | 4.52 | 1.87E-03 | 0.0143 | 117 |
| Q8VDD5 | MYH9_MOUSE | IAQLEEELEEEQGNTELINDR | 4.47 | 1.89E-03 | 0.0143 | 117 |
| Q8VDD5 | MYH9_MOUSE | GDLPFVVTR | 3.46 | 1.90E-03 | 0.0143 | 117 |
| Q8VDD5 | MYH9_MOUSE | QLEEAEEEAQR | 3.97 | 1.93E-03 | 0.0143 | 117 |
| Q8VDD5 | MYH9_MOUSE | YEILTPNSIPK | 4.47 | 1.96E-03 | 0.0143 | 117 |
| Q8VDD5 | MYH9_MOUSE | ITDVIIGFQAC[+57]C[+57]R | 4.79 | 1.99E-03 | 0.0143 | 117 |
| Q8VDD5 | MYH9_MOUSE | KQELEEIC[+57]HDLEAR | 4.59 | 2.04E-03 | 0.0143 | 117 |
| Q8VDD5 | MYH9_MOUSE | ELETQISELQEDLESER | 4.45 | 2.06E-03 | 0.0143 | 117 |
| Q8VDD5 | MYH9_MOUSE | NLPIYSEEIVEM[+16]YK | 4.36 | 2.06E-03 | 0.0143 | 117 |
| Q8VDD5 | MYH9_MOUSE | TELEDTLDSTAAQQELR | 5.84 | 2.12E-03 | 0.0143 | 117 |
| Q8VDD5 | MYH9_MOUSE | NFINNPLAQADWAAK | 4.76 | 2.13E-03 | 0.0143 | 117 |
| Q8VDD5 | MYH9_MOUSE | HEAMITDLEER | 4.78 | 2.15E-03 | 0.0143 | 117 |
| Q8VDD5 | MYH9_MOUSE | TDLLLEPYNK | 4.05 | 2.19E-03 | 0.0143 | 117 |
| Q8VDD5 | MYH9_MOUSE | IAQLEEQLDNETK | 4.09 | 2.22E-03 | 0.0143 | 117 |
| Q8VDD5 | MYH9_MOUSE | EQADFAIEALAK | 4.15 | 2.23E-03 | 0.0143 | 117 |
| Q8VDD5 | MYH9_MOUSE | IRELETQISELQEDLESER | 5.59 | 2.24E-03 | 0.0143 | 117 |
| Q8VDD5 | MYH9_MOUSE | TQLEELEDELQATEDAK | 4.57 | 2.32E-03 | 0.0143 | 117 |
| Q8VDD5 | MYH9_MOUSE | IIGLDQVAGMSETALPGAFK | 5.00 | 2.32E-03 | 0.0143 | 117 |
| Q8VDD5 | MYH9_MOUSE | C[+57]QYLQAEK | 3.97 | 2.34E-03 | 0.0143 | 117 |
| Q8VDD5 | MYH9_MOUSE | RKLEGDSTDLSDQIAELQAQIAELK | 5.17 | 2.41E-03 | 0.0143 | 117 |
| Q8VDD5 | MYH9_MOUSE | KEEELQAALAR | 3.81 | 2.45E-03 | 0.0143 | 117 |
| Q8VDD5 | MYH9_MOUSE | ANLQIDQINTDLNLER | 4.40 | 2.45E-03 | 0.0143 | 117 |
| Q8VDD5 | MYH9_MOUSE | VAAYDKLEK | 4.25 | 2.47E-03 | 0.0143 | 117 |

| Protein Accession | Protein Description | Modified Peptide Sequence | co-IP:Ctrl | p value | q value | Sig Peptide Count |
|---|---|---|---|---|---|---|
| Q8VDD5 | MYH9_MOUSE | LQVELDSVTGLLSQSDSK | 4.40 | 2.49E-03 | 0.0143 | 117 |
| Q8VDD5 | MYH9_MOUSE | NKHEAMITDLEER | 4.26 | 2.59E-03 | 0.0143 | 117 |
| Q8VDD5 | MYH9_MOUSE | GALALEEKR | 4.65 | 2.64E-03 | 0.0143 | 117 |
| Q8VDD5 | MYH9_MOUSE | HEDELLAK | 4.09 | 2.67E-03 | 0.0143 | 117 |
| Q8VDD5 | MYH9_MOUSE | HSQAVEELADQLEQTK | 4.64 | 2.72E-03 | 0.0143 | 117 |
| Q8VDD5 | MYH9_MOUSE | AGVLAHLEEER | 5.22 | 2.76E-03 | 0.0143 | 117 |
| Q8VDD5 | MYH9_MOUSE | LDPHLVLDQLR | 5.04 | 2.93E-03 | 0.0143 | 117 |
| Q8VDD5 | MYH9_MOUSE | VEAQLQELQVK | 3.79 | 3.01E-03 | 0.0143 | 117 |
| Q8VDD5 | MYH9_MOUSE | MEDGVGC[+57]LETAEEAK | 4.89 | 3.09E-03 | 0.0143 | 117 |
| Q8VDD5 | MYH9_MOUSE | LEGDSTDLSDQIAELQAQIAELK | 6.21 | 3.09E-03 | 0.0143 | 117 |
| Q8VDD5 | MYH9_MOUSE | ADEWLM[+16]K | 5.02 | 3.11E-03 | 0.0143 | 117 |
| Q8VDD5 | MYH9_MOUSE | SMEAEMIQLQEELAAAER | 5.53 | 3.16E-03 | 0.0143 | 117 |
| Q8VDD5 | MYH9_MOUSE | LTEMETM[+16]QSQLMAEK | 4.78 | 3.17E-03 | 0.0143 | 117 |
| Q8VDD5 | MYH9_MOUSE | KFDQLLAEEK | 3.98 | 3.17E-03 | 0.0143 | 117 |
| Q8VDD5 | MYH9_MOUSE | VAEFTTNLMEEEEK | 4.98 | 3.24E-03 | 0.0143 | 117 |
| Q8VDD5 | MYH9_MOUSE | NAEQFKDQADK | 4.74 | 3.44E-03 | 0.0144 | 117 |
| Q8VDD5 | MYH9_MOUSE | NLPIYSEEIVEMYK | 5.25 | 3.51E-03 | 0.0144 | 117 |
| Q8VDD5 | MYH9_MOUSE | NTDQASM[+16]PDNTAAQK | 4.38 | 3.56E-03 | 0.0145 | 117 |
| Q8VDD5 | MYH9_MOUSE | VSHLLGINVTDFTR | 5.00 | 3.60E-03 | 0.0145 | 117 |
| Q8VDD5 | MYH9_MOUSE | VVFQEFR | 3.80 | 3.66E-03 | 0.0146 | 117 |
| Q8VDD5 | MYH9_MOUSE | ADFC[+57]IIHYAGK | 4.99 | 3.82E-03 | 0.0147 | 117 |
| Q8VDD5 | MYH9_MOUSE | KLVWVPSSK | 3.52 | 3.85E-03 | 0.0147 | 117 |
| Q8VDD5 | MYH9_MOUSE | LQEMESAVK | 4.43 | 3.93E-03 | 0.0147 | 117 |
| Q8VDD5 | MYH9_MOUSE | KLEEDQIIMEDQNC[+57]K | 4.35 | 3.97E-03 | 0.0148 | 117 |
| Q8VDD5 | MYH9_MOUSE | DLQGRDEQSEEK | 6.37 | 4.04E-03 | 0.0148 | 117 |
| Q8VDD5 | MYH9_MOUSE | DVLLQVEDER | 2.71 | 4.05E-03 | 0.0148 | 117 |
| Q8VDD5 | MYH9_MOUSE | EMEAELEDER | 4.78 | 4.06E-03 | 0.0148 | 117 |
| Q8VDD5 | MYH9_MOUSE | ALEQQVEEMKTQLEELEDELQATEDAK | 8.79 | 4.08E-03 | 0.0148 | 117 |
| Q8VDD5 | MYH9_MOUSE | LQQLFNHTM[+16]FILEQEEYQR | 5.08 | 4.14E-03 | 0.0148 | 117 |
| Q8VDD5 | MYH9_MOUSE | KM[+16]EDGVGC[+57]LETAEEAKR | 4.81 | 4.16E-03 | 0.0149 | 117 |
| Q8VDD5 | MYH9_MOUSE | ELEDATETADAMNR | 3.85 | 4.20E-03 | 0.0149 | 117 |
| Q8VDD5 | MYH9_MOUSE | KMQQNIQELEEQLEEEESAR | 4.40 | 4.34E-03 | 0.0150 | 117 |
| Q8VDD5 | MYH9_MOUSE | RGDLPFVVTR | 4.10 | 4.90E-03 | 0.0155 | 117 |
| Q8VDD5 | MYH9_MOUSE | THEAQIQEMR | 5.12 | 4.93E-03 | 0.0155 | 117 |
| Q8VDD5 | MYH9_MOUSE | ALEEAMEQK | 3.63 | 5.32E-03 | 0.0158 | 117 |
| Q8VDD5 | MYH9_MOUSE | EQLEEEEEAKR | 4.35 | 5.44E-03 | 0.0158 | 117 |
| Q8VDD5 | MYH9_MOUSE | NGFEPASLKEEVGEEAIVELVENGK | 5.35 | 5.97E-03 | 0.0164 | 117 |
| Q8VDD5 | MYH9_MOUSE | VEEEEER | 4.66 | 6.73E-03 | 0.0172 | 117 |
| Q8VDD5 | MYH9_MOUSE | QAC[+57]VLMIK | 4.39 | 6.76E-03 | 0.0172 | 117 |
| Q8VDD5 | MYH9_MOUSE | LEEDQIIMEDQNC[+57]K | 4.46 | 6.80E-03 | 0.0172 | 117 |
| Q8VDD5 | MYH9_MOUSE | GTGDC[+57]SDEEVDGKADGADAK | 4.86 | 6.89E-03 | 0.0174 | 117 |
| Q8VDD5 | MYH9_MOUSE | NTDQASMPDNTAAQK | 2.66 | 7.05E-03 | 0.0175 | 117 |
| Q8VDD5 | MYH9_MOUSE | LTEMETMQSQLM[+16]AEK | 4.51 | 7.08E-03 | 0.0175 | 117 |
| Q8VDD5 | MYH9_MOUSE | SMEAEM[+16]IQLQEELAAAER | 4.34 | 7.11E-03 | 0.0176 | 117 |
| Q8VDD5 | MYH9_MOUSE | YLYVDKNFINNPLAQADWAAK | 5.46 | 7.14E-03 | 0.0176 | 117 |
| Q8VDD5 | MYH9_MOUSE | TFHIFYYLLSGAGEHLK | 6.14 | 7.42E-03 | 0.0179 | 117 |
| Q8VDD5 | MYH9_MOUSE | SMM[+16]QDREDQSILC[+57]TGESGAGK | 6.30 | 7.63E-03 | 0.0181 | 117 |
| Q8VDD5 | MYH9_MOUSE | RQLEEAEEEAQR | 7.02 | 7.97E-03 | 0.0183 | 117 |
| Q8VDD5 | MYH9_MOUSE | SMMQDREDQSILC[+57]TGESGAGK | 4.96 | 9.25E-03 | 0.0198 | 117 |
| Q8VDD5 | MYH9_MOUSE | LQQLFNHTMFILEQEEYQR | 7.25 | 9.95E-03 | 0.0206 | 117 |
| Q8VDD5 | MYH9_MOUSE | SM[+16]EAEM[+16]IQLQEELAAAER | 1.46 | 1.00E-02 | 0.0206 | 117 |
| Q8VDD5 | MYH9_MOUSE | KLEGDSTDLSDQIAELQAQIAELK | 6.04 | 1.24E-02 | 0.0231 | 117 |
| Q8VDD5 | MYH9_MOUSE | ADEWLMK | 4.67 | 1.62E-02 | 0.0273 | 117 |
| Q8VDD5 | MYH9_MOUSE | ALEQQVEEMK | 1.82 | 1.68E-02 | 0.0279 | 117 |
| Q7TQI3 | OTUB1_MOUSE | AFGFSHLEALLDDSKELQR | 2.46 | 1.92E-02 | 0.0306 | 117 |
| P67984 | RL22_MOUSE 6 | AGNLGGGVVTIER | 3.01 | 2.02E-02 | 0.0318 | 117 |
| P62754 | RS6_MOUSE 4( | GC[+57]IVDANLSVLNLVIVK | 2.30 | 2.02E-02 | 0.0318 | 117 |
| Q9Z1B3 | PLCB1_MOUSE | LNEILYPPLKQEQVQVLIEK | 5.47 | 2.27E-02 | 0.0347 | 117 |
| Q91ZU6 | DYST_MOUSE I | VGGGWMALDEFLVK | 4.14 | 2.37E-02 | 0.0358 | 117 |
| O08709 | PRDX6_MOUSE | DINAYNGETPTEK | 5.54 | 2.95E-02 | 0.0420 | 117 |
| Q99104 | MYO5A_MOUSE | M[+16]LPELFQDDEK | 4.11 | 1.34E-03 | 0.0143 | 98 |
| Q99104 | MYO5A_MOUSE | GEIQSLKEENNR | 4.22 | 1.54E-03 | 0.0143 | 98 |

| Protein Accession | Protein Description | Modified Peptide Sequence | co-IP:Ctrl | p value | q value | Sig Peptide Count |
|---|---|---|---|---|---|---|
| Q99104 | MYO5A_MOUSE | TDDDAEAIC[+57]SM[+16]C[+57]NALTTAQIVK | 5.62 | 1.76E-03 | 0.0143 | 98 |
| Q99104 | MYO5A_MOUSE | QYSGEEGFM[+16]K | 4.96 | 1.83E-03 | 0.0143 | 98 |
| Q99104 | MYO5A_MOUSE | LGILDLLDEEC[+57]KM[+16]PK | 4.79 | 1.98E-03 | 0.0143 | 98 |
| Q99104 | MYO5A_MOUSE | TSSIADEGTYTLDSILR | 4.73 | 2.22E-03 | 0.0143 | 98 |
| Q99104 | MYO5A_MOUSE | AAITVQR | 4.69 | 2.30E-03 | 0.0143 | 98 |
| Q99104 | MYO5A_MOUSE | VPLDMSLFLK | 5.52 | 2.30E-03 | 0.0143 | 98 |
| Q99104 | MYO5A_MOUSE | ETLEPLIQAAQLLQVK | 4.77 | 2.46E-03 | 0.0143 | 98 |
| Q99104 | MYO5A_MOUSE | QGGSPM[+16]IEGVDDAK | 3.94 | 2.61E-03 | 0.0143 | 98 |
| Q99104 | MYO5A_MOUSE | NTM[+16]TDSTILLEDVQK | 2.56 | 2.64E-03 | 0.0143 | 98 |
| Q99104 | MYO5A_MOUSE | LLESQLQSQK | 4.27 | 2.76E-03 | 0.0143 | 98 |
| Q99104 | MYO5A_MOUSE | QAC[+57]TLLGISESYQM[+16]GIFR | 5.87 | 2.78E-03 | 0.0143 | 98 |
| Q99104 | MYO5A_MOUSE | EQIPWTLIDFYDNQPC[+57]INLIESK | 8.73 | 2.82E-03 | 0.0143 | 98 |
| Q99104 | MYO5A_MOUSE | AISPTSATSSGR | 4.63 | 2.86E-03 | 0.0143 | 98 |
| Q99104 | MYO5A_MOUSE | QGGSPMIEGVDDAK | 2.99 | 2.88E-03 | 0.0143 | 98 |
| Q99104 | MYO5A_MOUSE | DDKNTM[+16]TDSTILLEDVQK | 4.01 | 2.97E-03 | 0.0143 | 98 |
| Q99104 | MYO5A_MOUSE | KLATATETYIKPISK | 4.47 | 3.06E-03 | 0.0143 | 98 |
| Q99104 | MYO5A_MOUSE | SHENEAEALR | 3.90 | 3.08E-03 | 0.0143 | 98 |
| Q99104 | MYO5A_MOUSE | FPFTFDEK | 5.01 | 3.51E-03 | 0.0144 | 98 |
| Q99104 | MYO5A_MOUSE | YNVSQLEEWLR | 5.55 | 3.52E-03 | 0.0144 | 98 |
| Q99104 | MYO5A_MOUSE | NQSIIVSGESGAGK | 4.46 | 3.78E-03 | 0.0147 | 98 |
| Q99104 | MYO5A_MOUSE | NKDTVFEEQIK | 4.46 | 3.79E-03 | 0.0147 | 98 |
| Q99104 | MYO5A_MOUSE | SAPEVTAPGAPAYR | 4.89 | 3.84E-03 | 0.0147 | 98 |
| Q99104 | MYO5A_MOUSE | IGELEVGQM[+16]ENISPGQIIDEPIRPVNIPR | 6.21 | 3.90E-03 | 0.0147 | 98 |
| Q99104 | MYO5A_MOUSE | DFQGMLEYK | 1.05 | 3.94E-03 | 0.0147 | 98 |
| Q99104 | MYO5A_MOUSE | TDSTHSSNESEYTFSSEFAETEDIAPR | 5.71 | 4.07E-03 | 0.0148 | 98 |
| Q99104 | MYO5A_MOUSE | VWIPDPEEVWK | 4.48 | 4.08E-03 | 0.0148 | 98 |
| Q99104 | MYO5A_MOUSE | ISAAGFPSR | 4.33 | 4.13E-03 | 0.0148 | 98 |
| Q99104 | MYO5A_MOUSE | IEASLQHEITR | 4.80 | 4.26E-03 | 0.0150 | 98 |
| Q99104 | MYO5A_MOUSE | GTDDTWAQK | 3.88 | 4.55E-03 | 0.0152 | 98 |
| Q99104 | MYO5A_MOUSE | AGQVAYLEK | 4.57 | 4.72E-03 | 0.0153 | 98 |
| Q99104 | MYO5A_MOUSE | DSPQLLMDAK | 3.54 | 4.79E-03 | 0.0154 | 98 |
| Q99104 | MYO5A_MOUSE | IGELEVGQMENISPGQIIDEPIRPVNIPR | 6.72 | 4.96E-03 | 0.0155 | 98 |
| Q99104 | MYO5A_MOUSE | KEEVLILR | 4.48 | 5.05E-03 | 0.0155 | 98 |
| Q99104 | MYO5A_MOUSE | HADYLNDDQK | 5.04 | 5.12E-03 | 0.0155 | 98 |
| Q99104 | MYO5A_MOUSE | SLLTSTINSIKK | 5.86 | 5.33E-03 | 0.0158 | 98 |
| Q99104 | MYO5A_MOUSE | YQNLLNEFSR | 4.81 | 5.35E-03 | 0.0158 | 98 |
| Q99104 | MYO5A_MOUSE | LGILDLLDEEC[+57]KMPK | 5.93 | 5.40E-03 | 0.0158 | 98 |
| Q99104 | MYO5A_MOUSE | LILDKDKYQFGK | 3.06 | 5.43E-03 | 0.0158 | 98 |
| Q99104 | MYO5A_MOUSE | GVAVNLIPGLPAYILFM[+16]C[+57]VR | 8.96 | 5.46E-03 | 0.0158 | 98 |
| Q99104 | MYO5A_MOUSE | QAC[+57]TLLGISESYQMGIFR | 4.07 | 5.48E-03 | 0.0159 | 98 |
| Q99104 | MYO5A_MOUSE | LATATETYIKPISK | 6.88 | 5.62E-03 | 0.0160 | 98 |
| Q99104 | MYO5A_MOUSE | GAELEYESLK | 2.76 | 5.69E-03 | 0.0161 | 98 |
| Q99104 | MYO5A_MOUSE | TDDDAEAIC[+57]SMC[+57]NALTTAQIVK | 5.71 | 5.86E-03 | 0.0163 | 98 |
| Q99104 | MYO5A_MOUSE | VVFQAEEER | 3.50 | 5.88E-03 | 0.0163 | 98 |
| Q99104 | MYO5A_MOUSE | EM[+16]TETM[+16]ER | 4.27 | 5.94E-03 | 0.0163 | 98 |
| Q99104 | MYO5A_MOUSE | YFATVSGSASEANVEEK | 5.23 | 5.98E-03 | 0.0164 | 98 |
| Q99104 | MYO5A_MOUSE | GEIAQAYIGLK | 4.56 | 6.40E-03 | 0.0169 | 98 |
| Q99104 | MYO5A_MOUSE | AC[+57]GVLETIR | 3.43 | 6.41E-03 | 0.0169 | 98 |
| Q99104 | MYO5A_MOUSE | SLLTSTINSIK | 4.94 | 6.46E-03 | 0.0170 | 98 |
| Q99104 | MYO5A_MOUSE | VLSLQEEIAK | 4.23 | 6.73E-03 | 0.0172 | 98 |
| Q99104 | MYO5A_MOUSE | QETDQLVSNLKEENTLLK | 4.84 | 7.06E-03 | 0.0175 | 98 |
| Q99104 | MYO5A_MOUSE | VEYQC[+57]EGFLEK | 5.00 | 7.11E-03 | 0.0176 | 98 |
| Q99104 | MYO5A_MOUSE | QLELDLNDER | 2.72 | 7.26E-03 | 0.0177 | 98 |
| Q99104 | MYO5A_MOUSE | LTNLEGVYNSETEK | 4.33 | 7.61E-03 | 0.0181 | 98 |
| Q99104 | MYO5A_MOUSE | KVPLDM[+16]SLFLK | 2.90 | 8.04E-03 | 0.0184 | 98 |
| Q99104 | MYO5A_MOUSE | RGDDFETVSFWLSNTC[+57]R | 7.16 | 8.21E-03 | 0.0186 | 98 |
| Q99104 | MYO5A_MOUSE | AIVYLQC[+57]C[+57]FR | 4.59 | 8.32E-03 | 0.0187 | 98 |
| Q99104 | MYO5A_MOUSE | MLPELFQDDEK | 4.68 | 8.34E-03 | 0.0188 | 98 |
| Q99104 | MYO5A_MOUSE | VLNLYTPVNEFEER | 5.45 | 8.38E-03 | 0.0188 | 98 |
| Q99104 | MYO5A_MOUSE | DKGEIAQAYIGLK | 3.87 | 8.39E-03 | 0.0188 | 98 |
| Q99104 | MYO5A_MOUSE | YIEIGFDK | 5.55 | 8.48E-03 | 0.0189 | 98 |
| Q99104 | MYO5A_MOUSE | LLESQLQSQKR | 3.05 | 8.56E-03 | 0.0190 | 98 |

| Protein Accession | Protein Description | Modified Peptide Sequence | co-IP:Ctrl | p value | q value | Sig Peptide Count |
|---|---|---|---|---|---|---|
| Q99104 | MYO5A_MOUSE | VLM[+16]EQLTSVSEELDVR | 4.14 | 8.86E-03 | 0.0193 | 98 |
| Q99104 | MYO5A_MOUSE | VLMEQLTSVSEELDVRK | 4.87 | 9.20E-03 | 0.0198 | 98 |
| Q99104 | MYO5A_MOUSE | QLMQDELDR | 4.02 | 9.21E-03 | 0.0198 | 98 |
| Q99104 | MYO5A_MOUSE | VSVSFIR | 4.59 | 9.27E-03 | 0.0199 | 98 |
| Q99104 | MYO5A_MOUSE | ILAGILHLGNVGFASR | 6.04 | 9.71E-03 | 0.0204 | 98 |
| Q99104 | MYO5A_MOUSE | VLASNPIMESIGNAK | 4.24 | 9.81E-03 | 0.0204 | 98 |
| Q99104 | MYO5A_MOUSE | VLASNPIM[+16]ESIGNAK | 2.45 | 1.00E-02 | 0.0206 | 98 |
| Q99104 | MYO5A_MOUSE | QVLSDLAIQIYQQLVR | 8.86 | 1.04E-02 | 0.0211 | 98 |
| Q99104 | MYO5A_MOUSE | EMTETMER | 5.92 | 1.05E-02 | 0.0212 | 98 |
| Q99104 | MYO5A_MOUSE | HFADKVEYQC[+57]EGFLEK | 5.73 | 1.06E-02 | 0.0213 | 98 |
| Q99104 | MYO5A_MOUSE | DDKNTMTDSTILLEDVQK | 4.77 | 1.08E-02 | 0.0215 | 98 |
| Q99104 | MYO5A_MOUSE | LGNADSFHYTK | 5.16 | 1.11E-02 | 0.0219 | 98 |
| Q99104 | MYO5A_MOUSE | LQQQFNMHVFK | 5.35 | 1.12E-02 | 0.0219 | 98 |
| Q99104 | MYO5A_MOUSE | LEQEEYMK | 1.87 | 1.15E-02 | 0.0223 | 98 |
| Q99104 | MYO5A_MOUSE | GAELEYESLKR | 4.21 | 1.18E-02 | 0.0226 | 98 |
| Q99104 | MYO5A_MOUSE | LGILDLLDEEC[+57]K | 4.90 | 1.23E-02 | 0.0231 | 98 |
| Q99104 | MYO5A_MOUSE | VLLLHLEEGKDLEYR | 4.58 | 1.24E-02 | 0.0231 | 98 |
| Q99104 | MYO5A_MOUSE | YKQETDQLVSNLKEENTLLK | 6.05 | 1.25E-02 | 0.0232 | 98 |
| Q99104 | MYO5A_MOUSE | AATIVIQSYLR | 5.42 | 1.30E-02 | 0.0238 | 98 |
| Q99104 | MYO5A_MOUSE | KVPLDMSLFLK | 5.37 | 1.43E-02 | 0.0253 | 98 |
| Q99104 | MYO5A_MOUSE | QNEHC[+57]LTNFDLAEYR | 4.87 | 1.47E-02 | 0.0257 | 98 |
| Q99104 | MYO5A_MOUSE | NYHIFYQLC[+57]ASAK | 5.83 | 1.57E-02 | 0.0268 | 98 |
| P43277 | H13_MOUSE Hi | SGVSLAALK | 6.09 | 1.72E-02 | 0.0284 | 98 |
| Q04447 | KCRB_MOUSE | LGFSEVELVQM[+16]VVDGVK | 4.01 | 1.72E-02 | 0.0284 | 98 |
| P61294 | RAB6B_MOUSE | QITIEEGEQR | 3.66 | 1.78E-02 | 0.0290 | 98 |
| P14873 | MAP1B_MOUSE | SIEEAC[+57]FTLQYLNK | 8.18 | 1.82E-02 | 0.0295 | 98 |
| P01869 | IGH1M_MOUSE | VNSAAFPAPIEK | 7.75 | 1.82E-02 | 0.0295 | 98 |
| Q61937 | NPM_MOUSE N | MTDQEAIQDLWQWR | 4.19 | 1.89E-02 | 0.0302 | 98 |
| O08539 | BIN1_MOUSE M | VGFYVNTFQSIAGLEENFHK | 3.80 | 2.19E-02 | 0.0338 | 98 |
| Q7TPR4 | ACTN1_MOUSE | GISQEQMNEFR | 4.32 | 2.43E-02 | 0.0364 | 98 |
| A0A0N5DKY8 | A0A0N5DKY8_TRI | AFMTADLPNELIEILEK | 4.01 | 2.50E-02 | 0.0371 | 98 |
| Q9Z0E0 | NCDN_MOUSE | NDSEQFAALLLVTK | 3.35 | 2.78E-02 | 0.0402 | 98 |
| P52480 | KPYM_MOUSE | AGKPVIC[+57]ATQMLESMIK | 6.47 | 2.92E-02 | 0.0417 | 98 |
| Q99M74 | KRT82_MOUSE | LAGLEEALQK | 3.52 | 3.02E-02 | 0.0428 | 98 |
| Q9JMH9 | MY18A_MOUSE | LQQELEDKM[+16]EVEQQSR | 3.85 | 5.45E-04 | 0.0143 | 63 |
| Q9JMH9 | MY18A_MOUSE | WQALSTLLEAFGNSPTIM[+16]NGSATR | 6.59 | 1.18E-03 | 0.0143 | 63 |
| Q9JMH9 | MY18A_MOUSE | DLALGLVPGDR | 2.87 | 1.87E-03 | 0.0143 | 63 |
| Q9JMH9 | MY18A_MOUSE | RFDSELSQAHEETQR | 5.54 | 2.05E-03 | 0.0143 | 63 |
| Q9JMH9 | MY18A_MOUSE | LQALQSQVEFLEQSMVDK | 5.30 | 2.48E-03 | 0.0143 | 63 |
| Q9JMH9 | MY18A_MOUSE | LQALQSQVEFLEQSM[+16]VDK | 4.26 | 2.51E-03 | 0.0143 | 63 |
| Q9JMH9 | MY18A_MOUSE | AVEELLESLDLEK | 4.51 | 2.79E-03 | 0.0143 | 63 |
| Q9JMH9 | MY18A_MOUSE | NQLEESEFTC[+57]AAAVK | 4.76 | 3.29E-03 | 0.0143 | 63 |
| Q9JMH9 | MY18A_MOUSE | VVSLEAELQDISSQESKDEASLAK | 5.36 | 3.45E-03 | 0.0144 | 63 |
| Q9JMH9 | MY18A_MOUSE | LQQELEDKMEVEQQSR | 5.54 | 3.60E-03 | 0.0145 | 63 |
| Q9JMH9 | MY18A_MOUSE | NTGESASQLLDAETAER | 5.59 | 3.83E-03 | 0.0147 | 63 |
| Q9JMH9 | MY18A_MOUSE | SEELSLPEGK | 3.97 | 3.89E-03 | 0.0147 | 63 |
| Q9JMH9 | MY18A_MOUSE | DEEVEEAR | 5.45 | 3.93E-03 | 0.0147 | 63 |
| Q9JMH9 | MY18A_MOUSE | LVEINGQNVENK | 3.93 | 4.27E-03 | 0.0150 | 63 |
| Q9JMH9 | MY18A_MOUSE | LHLEGQQVR | 6.08 | 4.29E-03 | 0.0150 | 63 |
| Q9JMH9 | MY18A_MOUSE | MSAAELR | 4.37 | 4.39E-03 | 0.0150 | 63 |
| Q9JMH9 | MY18A_MOUSE | TFLQELER | 4.46 | 4.77E-03 | 0.0154 | 63 |
| Q9JMH9 | MY18A_MOUSE | GFFNLNR | 4.90 | 4.86E-03 | 0.0155 | 63 |
| Q9JMH9 | MY18A_MOUSE | ALLADAQIMLDHLK | 5.80 | 4.89E-03 | 0.0155 | 63 |
| Q9JMH9 | MY18A_MOUSE | LTAELQDTK | 4.11 | 5.37E-03 | 0.0158 | 63 |
| Q9JMH9 | MY18A_MOUSE | ALLADAQIM[+16]LDHLK | 2.74 | 5.41E-03 | 0.0158 | 63 |
| Q9JMH9 | MY18A_MOUSE | LEISNPIPIK | 4.96 | 5.42E-03 | 0.0158 | 63 |
| Q9JMH9 | MY18A_MOUSE | LGDLQADSDESQR | 5.43 | 5.63E-03 | 0.0160 | 63 |
| Q9JMH9 | MY18A_MOUSE | QDQSIVLLGSSGSGK | 5.18 | 5.76E-03 | 0.0161 | 63 |
| Q9JMH9 | MY18A_MOUSE | QNPATQNAPR | 5.44 | 5.77E-03 | 0.0162 | 63 |
| Q9JMH9 | MY18A_MOUSE | AAEINGEVDDDDAGGEWR | 4.90 | 5.84E-03 | 0.0163 | 63 |
| Q9JMH9 | MY18A_MOUSE | RFDVLAPHLTK | 4.50 | 5.96E-03 | 0.0163 | 63 |
| Q9JMH9 | MY18A_MOUSE | NKDEEIQQLR | 5.61 | 5.98E-03 | 0.0164 | 63 |

| Protein Accession | Protein Description | Modified Peptide Sequence | co-IP:Ctrl | p value | q value | Sig Peptide Count |
|---|---|---|---|---|---|---|
| Q9JMH9 | MY18A_MOUSE | RPTGDFGFSLR | 5.79 | 5.99E-03 | 0.0164 | 63 |
| Q9JMH9 | MY18A_MOUSE | DGFSLASQLK | 4.74 | 5.99E-03 | 0.0164 | 63 |
| Q9JMH9 | MY18A_MOUSE | VVSLEAELQDISSQESK | 3.23 | 6.12E-03 | 0.0166 | 63 |
| Q9JMH9 | MY18A_MOUSE | IISNLFLGR | 5.81 | 6.32E-03 | 0.0169 | 63 |
| Q9JMH9 | MY18A_MOUSE | LEDLASLVYLNESSVLHTLR | 6.83 | 6.60E-03 | 0.0171 | 63 |
| Q9JMH9 | MY18A_MOUSE | QGPEESGLGEGTK | 4.76 | 6.66E-03 | 0.0172 | 63 |
| Q9JMH9 | MY18A_MOUSE | SSC[+57]C[+57]LGLSR | 3.68 | 6.79E-03 | 0.0172 | 63 |
| Q9JMH9 | MY18A_MOUSE | LPALVPPPPPALR | 5.79 | 6.89E-03 | 0.0174 | 63 |
| Q9JMH9 | MY18A_MOUSE | DTKEEMSELAR | 4.90 | 7.17E-03 | 0.0177 | 63 |
| Q9JMH9 | MY18A_MOUSE | EKDM[+16]LLAEAFSLK | 5.03 | 7.79E-03 | 0.0182 | 63 |
| Q9JMH9 | MY18A_MOUSE | YGASLLHTYAGPSLLVLSTR | 5.99 | 9.11E-03 | 0.0197 | 63 |
| Q9JMH9 | MY18A_MOUSE | VLAISPEEQK | 4.42 | 9.24E-03 | 0.0198 | 63 |
| Q9JMH9 | MY18A_MOUSE | LQVDALIDTIK | 6.27 | 1.15E-02 | 0.0223 | 63 |
| Q9JMH9 | MY18A_MOUSE | VSSSSELDLPPGDPC[+57]EAGLLQLDVSLLR | 7.54 | 1.24E-02 | 0.0231 | 63 |
| Q9JMH9 | MY18A_MOUSE | GSIILDSGHLSTASSSDDLKGEEGSFR | 7.15 | 1.27E-02 | 0.0234 | 63 |
| Q9JMH9 | MY18A_MOUSE | VWLVHR | 2.48 | 1.28E-02 | 0.0235 | 63 |
| Q9JMH9 | MY18A_MOUSE | VKDQEEELDEQAGSIQMLEQAK | 4.25 | 1.30E-02 | 0.0238 | 63 |
| Q9JMH9 | MY18A_MOUSE | QMEVQLEEEYEDKQK | 4.68 | 1.31E-02 | 0.0239 | 63 |
| Q9JMH9 | MY18A_MOUSE | EKDMLLAEAFSLK | 6.19 | 1.31E-02 | 0.0239 | 63 |
| Q9JMH9 | MY18A_MOUSE | LQVDALIDTIKR | 6.06 | 1.32E-02 | 0.0240 | 63 |
| Q9JMH9 | MY18A_MOUSE | FSHSYLSDSDTEAK | 4.85 | 1.35E-02 | 0.0243 | 63 |
| Q9JMH9 | MY18A_MOUSE | VKLDHDGAILDVDEDDIEK | 5.72 | 1.38E-02 | 0.0247 | 63 |
| Q9JMH9 | MY18A_MOUSE | DLDIAGFTQK | 2.35 | 1.49E-02 | 0.0259 | 63 |
| Q9JMH9 | MY18A_MOUSE | LFTTVRPLIQVQLSEEQIR | 7.13 | 1.52E-02 | 0.0263 | 63 |
| Q9JMH9 | MY18A_MOUSE | AAYLLGC[+57]SLEELSSAIFK | 8.02 | 1.54E-02 | 0.0265 | 63 |
| Q9JMH9 | MY18A_MOUSE | AGSATVLSGSIAGLEGGSQLALR | 7.09 | 1.63E-02 | 0.0274 | 63 |
| Q9JMH9 | MY18A_MOUSE | RAVEELLESLDLEK | 2.19 | 1.68E-02 | 0.0279 | 63 |
| P17426 | AP2A1_MOUSE | VAAQVDGGAQVQQVLNIEC[+57]LR | 8.26 | 1.76E-02 | 0.0288 | 63 |
| P16546 | SPTN1_MOUSE | DLTSWVTEMK | 5.88 | 2.05E-02 | 0.0322 | 63 |
| Q7TMM9 | TBB2A_MOUSE | RISEQFTAMFR | 4.29 | 2.22E-02 | 0.0342 | 63 |
| P50518 | VATE1_MOUSE | VSNTLESR | 5.83 | 2.71E-02 | 0.0394 | 63 |
| P05202 | AATM_MOUSE | NLDKEYLPIGGLAEFC[+57]K | 6.77 | 2.72E-02 | 0.0395 | 63 |
| Q6IME9 | K2C72_MOUSE | TAAENEFVVLK | 3.25 | 2.85E-02 | 0.0409 | 63 |
| P51863 | VA0D1_MOUSE | NIVWIAEC[+57]IAQR | 4.23 | 3.22E-02 | 0.0451 | 63 |
| P63038 | CH60_MOUSE 6 | ISSVQSIVPALEIANAHR | 2.46 | 3.22E-02 | 0.0451 | 63 |
| Q6URW6 | MYH14_MOUSE | RQEEEAAVLEAGEEAR | 4.03 | 3.24E-05 | 0.0143 | 61 |
| Q6URW6 | MYH14_MOUSE | AAEQAASDLR | 3.80 | 3.45E-04 | 0.0143 | 61 |
| Q6URW6 | MYH14_MOUSE | VLGLLPEEITAM[+16]LR | 4.57 | 3.46E-04 | 0.0143 | 61 |
| Q6URW6 | MYH14_MOUSE | RLESQLQEVQGR | 4.68 | 6.10E-04 | 0.0143 | 61 |
| Q6URW6 | MYH14_MOUSE | AEELLAQLGR | 3.40 | 9.44E-04 | 0.0143 | 61 |
| Q6URW6 | MYH14_MOUSE | AEAELC[+57]SEAEETR | 4.83 | 1.12E-03 | 0.0143 | 61 |
| Q6URW6 | MYH14_MOUSE | LAQAEEQLEQESR | 2.63 | 1.13E-03 | 0.0143 | 61 |
| Q6URW6 | MYH14_MOUSE | ELQSTQAQLSEWR | 2.00 | 1.32E-03 | 0.0143 | 61 |
| Q6URW6 | MYH14_MOUSE | AELSSLQTSR | 3.47 | 1.37E-03 | 0.0143 | 61 |
| Q6URW6 | MYH14_MOUSE | FLTNGPSSSPGQER | 5.15 | 1.40E-03 | 0.0143 | 61 |
| Q6URW6 | MYH14_MOUSE | AELEALLSSK | 4.61 | 1.49E-03 | 0.0143 | 61 |
| Q6URW6 | MYH14_MOUSE | QLEEAEEEASR | 5.99 | 1.62E-03 | 0.0143 | 61 |
| Q6URW6 | MYH14_MOUSE | VLGLLPEEITAMLR | 7.32 | 1.68E-03 | 0.0143 | 61 |
| Q6URW6 | MYH14_MOUSE | LSQLEEELEEEQNNSELLK | 5.99 | 1.70E-03 | 0.0143 | 61 |
| Q6URW6 | MYH14_MOUSE | LQQELDDATVDLGQQK | 6.85 | 1.72E-03 | 0.0143 | 61 |
| Q6URW6 | MYH14_MOUSE | TPNVGGPGGPQVEWTAR | 5.68 | 1.75E-03 | 0.0143 | 61 |
| Q6URW6 | MYH14_MOUSE | VGEEEEC[+57]SR | 6.14 | 1.79E-03 | 0.0143 | 61 |
| Q6URW6 | MYH14_MOUSE | LSLEAEVSELKAELSSLQTSR | 5.73 | 1.86E-03 | 0.0143 | 61 |
| Q6URW6 | MYH14_MOUSE | GLEAEVLR | 4.35 | 1.89E-03 | 0.0143 | 61 |
| Q6URW6 | MYH14_MOUSE | ELEDVTESAESMNR | 5.55 | 1.98E-03 | 0.0143 | 61 |
| Q6URW6 | MYH14_MOUSE | TVSAVLQFGNIVLK | 5.45 | 2.05E-03 | 0.0143 | 61 |
| Q6URW6 | MYH14_MOUSE | ELSSAESQLHDTQELLQEETR | 4.83 | 2.11E-03 | 0.0143 | 61 |
| Q6URW6 | MYH14_MOUSE | TQVTELEDELTAAEDAK | 4.75 | 2.21E-03 | 0.0143 | 61 |
| Q6URW6 | MYH14_MOUSE | EQADFALEALAK | 4.15 | 2.23E-03 | 0.0143 | 61 |
| Q6URW6 | MYH14_MOUSE | ILFQEFR | 5.50 | 2.29E-03 | 0.0143 | 61 |
| Q6URW6 | MYH14_MOUSE | GELEDTLDSTNAQQELR | 2.19 | 2.49E-03 | 0.0143 | 61 |
| Q6URW6 | MYH14_MOUSE | MIQALELDPNLYR | 4.61 | 2.54E-03 | 0.0143 | 61 |

| Protein Accession | Protein Description | Modified Peptide Sequence | co-IP:Ctrl | p value | q value | Sig Peptide Count |
|---|---|---|---|---|---|---|
| Q6URW6 | MYH14_MOUSE | LGEEDAGAR | 4.05 | 2.54E-03 | 0.0143 | 61 |
| Q6URW6 | MYH14_MOUSE | LLGLGVTDFSR | 4.64 | 2.80E-03 | 0.0143 | 61 |
| Q6URW6 | MYH14_MOUSE | LVLQVESLTTELSAER | 4.87 | 2.81E-03 | 0.0143 | 61 |
| Q6URW6 | MYH14_MOUSE | RLQQELDDATVDLGQQK | 4.22 | 2.83E-03 | 0.0143 | 61 |
| Q6URW6 | MYH14_MOUSE | LM[+16]ATLSNTNPSFVR | 2.26 | 2.92E-03 | 0.0143 | 61 |
| Q6URW6 | MYH14_MOUSE | DEC[+57]SFHIFYQLLGGAGEQLK | 5.83 | 2.98E-03 | 0.0143 | 61 |
| Q6URW6 | MYH14_MOUSE | LSLEAEVSELK | 4.49 | 3.09E-03 | 0.0143 | 61 |
| Q6URW6 | MYH14_MOUSE | LQQHIQELESHLEAEEGAR | 5.23 | 3.15E-03 | 0.0143 | 61 |
| Q6URW6 | MYH14_MOUSE | KFEEDLLLLEDQNSK | 5.12 | 3.16E-03 | 0.0143 | 61 |
| Q6URW6 | MYH14_MOUSE | ALEAEAAGLR | 4.55 | 3.18E-03 | 0.0143 | 61 |
| Q6URW6 | MYH14_MOUSE | KEDELQAALLR | 5.05 | 3.32E-03 | 0.0143 | 61 |
| Q6URW6 | MYH14_MOUSE | VIQYLAHVASSPK | 5.90 | 3.49E-03 | 0.0144 | 61 |
| Q6URW6 | MYH14_MOUSE | VASRPGPVPEAAQSFLYAPR | 6.46 | 3.76E-03 | 0.0147 | 61 |
| Q6URW6 | MYH14_MOUSE | KQELELVVTELEAR | 4.97 | 4.18E-03 | 0.0149 | 61 |
| Q6URW6 | MYH14_MOUSE | QAQQDRDEMAEEVASGNLSK | 5.01 | 4.19E-03 | 0.0149 | 61 |
| Q6URW6 | MYH14_MOUSE | AQAELESVSTALSEAESK | 2.67 | 4.51E-03 | 0.0152 | 61 |
| Q6URW6 | MYH14_MOUSE | ALEAEAAGLREQMEEEVVAR | 3.63 | 4.83E-03 | 0.0154 | 61 |
| Q6URW6 | MYH14_MOUSE | VAQLEEER | 3.54 | 5.56E-03 | 0.0159 | 61 |
| Q6URW6 | MYH14_MOUSE | DVEGIVGLEQVSSLGDGPPGGRPR | 5.76 | 5.70E-03 | 0.0161 | 61 |
| Q6URW6 | MYH14_MOUSE | QRAEELLAQLGR | 3.80 | 5.80E-03 | 0.0162 | 61 |
| Q6URW6 | MYH14_MOUSE | LMATLSNTNPSFVR | 5.63 | 5.84E-03 | 0.0163 | 61 |
| Q6URW6 | MYH14_MOUSE | EQMEEEVVAR | 2.71 | 5.95E-03 | 0.0163 | 61 |
| Q6URW6 | MYH14_MOUSE | VTDIIVSFQAAAR | 5.08 | 6.25E-03 | 0.0168 | 61 |
| Q6URW6 | MYH14_MOUSE | LQQLFNHTMFVLEQEEYQR | 5.06 | 7.05E-03 | 0.0175 | 61 |
| Q6URW6 | MYH14_MOUSE | EAQAGLAEAQEDLEAER | 3.49 | 7.13E-03 | 0.0176 | 61 |
| Q6URW6 | MYH14_MOUSE | LRLEVTVQALK | 7.28 | 7.95E-03 | 0.0183 | 61 |
| Q6URW6 | MYH14_MOUSE | LKYEATISDMEDR | 4.86 | 8.46E-03 | 0.0189 | 61 |
| Q6URW6 | MYH14_MOUSE | RRQEEEAAVLEAGEEAR | 3.30 | 1.27E-02 | 0.0234 | 61 |
| Q6URW6 | MYH14_MOUSE | VAQEQGSHPK | 3.34 | 1.46E-02 | 0.0256 | 61 |
| P12382 | PFKAL_MOUSE | LNIIIIAEGAIDR | 1.45 | 1.87E-02 | 0.0300 | 61 |
| P62259 | 1433E_MOUSE | HLIPAANTGESK | 2.74 | 2.46E-02 | 0.0367 | 61 |
| Q9CQE8 | RTRAF_MOUSE | INEAIVAVQAIIADPK | 8.15 | 2.49E-02 | 0.0370 | 61 |
| Q99104 | MYO5A_MOUSE | IMQLQR | 6.92 | 2.92E-02 | 0.0417 | 61 |
| Q8R1B4 | EIF3C_MOUSE | VWDLFPEADKVR | 4.68 | 3.47E-02 | 0.0481 | 61 |
| P16546 | SPTN1_MOUSE | VNDVC[+57]TNGQDLIK | 2.81 | 4.53E-04 | 0.0143 | 61 |
| P16546 | SPTN1_MOUSE | DVTGAEALLER | 2.96 | 1.24E-03 | 0.0143 | 61 |
| P16546 | SPTN1_MOUSE | MTLVASEDYGDTLAAIQGLLK | 4.28 | 1.34E-03 | 0.0143 | 61 |
| P16546 | SPTN1_MOUSE | DLASVNNLLK | 2.70 | 1.35E-03 | 0.0143 | 61 |
| P16546 | SPTN1_MOUSE | NQALNTDNYGHDLASVQALQR | 3.08 | 1.62E-03 | 0.0143 | 61 |
| P16546 | SPTN1_MOUSE | KIEDLGAAMEEALILDNK | 3.65 | 1.71E-03 | 0.0143 | 61 |
| P16546 | SPTN1_MOUSE | IAALQAFADQLIAVDHYAK | 4.68 | 1.75E-03 | 0.0143 | 61 |
| P16546 | SPTN1_MOUSE | DVEDEETWIR | 3.43 | 1.81E-03 | 0.0143 | 61 |
| P16546 | SPTN1_MOUSE | EAIVTSEELGQDLEHVEVLQK | 2.58 | 2.22E-03 | 0.0143 | 61 |
| P16546 | SPTN1_MOUSE | TKQDEVNAAWQR | 2.80 | 2.92E-03 | 0.0143 | 61 |
| P16546 | SPTN1_MOUSE | QVEELYQSLLELGEK | 4.43 | 3.14E-03 | 0.0143 | 61 |
| P16546 | SPTN1_MOUSE | LLVSSEDYGR | 2.06 | 3.45E-03 | 0.0144 | 61 |
| P16546 | SPTN1_MOUSE | DLAALGDKVNSLGETAQR | 2.02 | 3.56E-03 | 0.0145 | 61 |
| P16546 | SPTN1_MOUSE | LSDDNTIGQEEIQQR | 2.89 | 3.83E-03 | 0.0147 | 61 |
| P16546 | SPTN1_MOUSE | DLTNVQNLQK | 2.00 | 4.06E-03 | 0.0148 | 61 |
| P16546 | SPTN1_MOUSE | DLIGVQNLLK | 2.66 | 4.53E-03 | 0.0152 | 61 |
| P16546 | SPTN1_MOUSE | LIQSHPESAEDLKEK | 2.35 | 4.91E-03 | 0.0155 | 61 |
| P16546 | SPTN1_MOUSE | RQEENDKLR | 3.01 | 4.93E-03 | 0.0155 | 61 |
| P16546 | SPTN1_MOUSE | HQAFEAELSANQSR | 3.09 | 5.03E-03 | 0.0155 | 61 |
| P16546 | SPTN1_MOUSE | VNEVSQFAAK | 2.37 | 5.40E-03 | 0.0158 | 61 |
| P16546 | SPTN1_MOUSE | DLASVQALLR | 2.51 | 5.50E-03 | 0.0159 | 61 |
| P16546 | SPTN1_MOUSE | EAFLNTEDKGDSLDSVEALIK | 3.47 | 5.51E-03 | 0.0159 | 61 |
| P16546 | SPTN1_MOUSE | LGESQTLQQFSR | 2.73 | 5.54E-03 | 0.0159 | 61 |
| P16546 | SPTN1_MOUSE | SQLLGSAHEVQR | 2.36 | 5.55E-03 | 0.0159 | 61 |
| P16546 | SPTN1_MOUSE | HQAFEAELHANADR | 3.82 | 5.61E-03 | 0.0160 | 61 |
| P16546 | SPTN1_MOUSE | VLETAEDIQER | 2.44 | 5.74E-03 | 0.0161 | 61 |
| P16546 | SPTN1_MOUSE | ASAFNSWFENAEEDLTDPVR | 6.47 | 5.94E-03 | 0.0163 | 61 |
| P16546 | SPTN1_MOUSE | SLGYDLPM[+16]VEEGEPDPEFEAILDTVDPNR | 6.92 | 6.48E-03 | 0.0170 | 61 |

| Protein Accession | Protein Description | Modified Peptide Sequence | co-IP:Ctrl | p value | q value | Sig Peptide Count |
|---|---|---|---|---|---|---|
| P16546 | SPTN1_MOUSE | DLSSVQTLLTK | 3.15 | 6.85E-03 | 0.0173 | 61 |
| P16546 | SPTN1_MOUSE | LAALADQWQFLVQK | 3.87 | 6.90E-03 | 0.0174 | 61 |
| P16546 | SPTN1_MOUSE | QETFDAGLQAFQQEGIANITALKDQLLAAK | 8.04 | 7.12E-03 | 0.0176 | 61 |
| P16546 | SPTN1_MOUSE | GLVSSDELAK | 2.33 | 7.21E-03 | 0.0177 | 61 |
| P16546 | SPTN1_MOUSE | EANQQQQFNR | 3.36 | 7.53E-03 | 0.0180 | 61 |
| P16546 | SPTN1_MOUSE | LDENSAFLQFNWK | 4.05 | 7.55E-03 | 0.0180 | 61 |
| P16546 | SPTN1_MOUSE | VNSLGETAQR | 2.66 | 7.67E-03 | 0.0181 | 61 |
| P16546 | SPTN1_MOUSE | HQLLEADISAHEDR | 3.39 | 7.78E-03 | 0.0182 | 61 |
| P16546 | SPTN1_MOUSE | HQAFEAEVQANSGAIVK | 2.60 | 7.96E-03 | 0.0183 | 61 |
| P16546 | SPTN1_MOUSE | YTEHSTVGLAQQWDQLDQLGMR | 4.32 | 8.66E-03 | 0.0191 | 61 |
| P16546 | SPTN1_MOUSE | TALLELWELR | 4.72 | 8.92E-03 | 0.0194 | 61 |
| P16546 | SPTN1_MOUSE | LFGAAEVQR | 2.45 | 9.01E-03 | 0.0196 | 61 |
| P16546 | SPTN1_MOUSE | ADVVESWIGEK | 2.13 | 9.48E-03 | 0.0202 | 61 |
| P16546 | SPTN1_MOUSE | QEAFLLNEDLGDSLDSVEALLK | 8.52 | 1.01E-02 | 0.0207 | 61 |
| P16546 | SPTN1_MOUSE | SADESGQALLAASHYASDEVR | 3.12 | 1.03E-02 | 0.0209 | 61 |
| P16546 | SPTN1_MOUSE | DLMSWINGIR | 5.99 | 1.22E-02 | 0.0230 | 61 |
| P16546 | SPTN1_MOUSE | QETFDAGLQAFQQEGIANITALK | 5.06 | 1.25E-02 | 0.0232 | 61 |
| P16546 | SPTN1_MOUSE | LEESLEYQQFVANVEEEEAWINEK | 5.50 | 1.26E-02 | 0.0233 | 61 |
| P16546 | SPTN1_MOUSE | IDGITIQAR | 2.18 | 1.31E-02 | 0.0239 | 61 |
| P16546 | SPTN1_MOUSE | WTQLLANSATR | 2.73 | 1.35E-02 | 0.0243 | 61 |
| P16546 | SPTN1_MOUSE | TYLLDGSC[+57]MVEESGTLESQLEATK | 3.15 | 1.48E-02 | 0.0258 | 61 |
| P16546 | SPTN1_MOUSE | LM[+16]VHTVATFNSIK | 3.09 | 1.60E-02 | 0.0271 | 61 |
| P16546 | SPTN1_MOUSE | EKEPIVGSTDYGKDEDSAEALLK | 3.92 | 1.61E-02 | 0.0272 | 61 |
| P16546 | SPTN1_MOUSE | GKDLIGVQNLLK | 3.43 | 1.62E-02 | 0.0273 | 61 |
| P62751 | RL23A_MOUSE | LAPDYDALDVANK | 3.19 | 1.79E-02 | 0.0292 | 61 |
| P51410 | RL9_MOUSE 60 | DELILEGNDIELVSNSAALIQQATTVK | 3.70 | 1.80E-02 | 0.0293 | 61 |
| Q8BH44 | COR2B_MOUSE | SVVVNGIDLLENVPPR | 3.20 | 1.85E-02 | 0.0298 | 61 |
| Q99KI0 | ACON_MOUSE | QGLLPLTFADPSDYNK | 3.07 | 1.88E-02 | 0.0301 | 61 |
| P61205 | ARF3_MOUSE / | MLAEDELR | 7.44 | 1.94E-02 | 0.0308 | 61 |
| P63318 | KPCG_MOUSE | NDFMGAMSFGVSELLK | 3.15 | 2.04E-02 | 0.0321 | 61 |
| Q60875 | ARHG2_MOUSE | M[+16]QDIPEETESR | 3.24 | 2.12E-02 | 0.0330 | 61 |
| Q99104 | MYO5A_MOUSE | VLMEQLTSVSEELDVR | 1.79 | 2.50E-02 | 0.0371 | 61 |
| Q9WUM4 | COR1C_MOUSE | VGIVAWHPTAR | 4.06 | 2.64E-02 | 0.0387 | 61 |
| Q68FD5 | CLH1_MOUSE ( | ADDPSSYM[+16]EVVQAANASGNWEELVK | 2.05 | 5.21E-04 | 0.0143 | 49 |
| Q68FD5 | CLH1_MOUSE ( | AFM[+16]TADLPNELIELLEK | 4.22 | 5.52E-04 | 0.0143 | 49 |
| Q68FD5 | CLH1_MOUSE ( | VDKLDASESLR | 1.50 | 5.96E-04 | 0.0143 | 49 |
| Q68FD5 | CLH1_MOUSE ( | NLQNLLILTAIK | 2.18 | 6.01E-04 | 0.0143 | 49 |
| Q68FD5 | CLH1_MOUSE ( | LLLPWLEAR | 2.99 | 1.69E-03 | 0.0143 | 49 |
| Q68FD5 | CLH1_MOUSE ( | TLQIFNIEM[+16]K | 1.66 | 1.74E-03 | 0.0143 | 49 |
| Q68FD5 | CLH1_MOUSE ( | LPVVIGGLLDVDC[+57]SEDVIK | 2.58 | 1.77E-03 | 0.0143 | 49 |
| Q68FD5 | CLH1_MOUSE ( | SVNESLNNLFITEEDYQALR | 2.69 | 1.85E-03 | 0.0143 | 49 |
| Q68FD5 | CLH1_MOUSE ( | ESYVETELIFALAK | 3.79 | 1.87E-03 | 0.0143 | 49 |
| Q68FD5 | CLH1_MOUSE ( | SVDPTLALSVYLR | 2.90 | 1.88E-03 | 0.0143 | 49 |
| Q68FD5 | CLH1_MOUSE ( | AVDVFFPPEAQNDFPVAMQISEK | 3.82 | 2.26E-03 | 0.0143 | 49 |
| Q68FD5 | CLH1_MOUSE ( | HNIMDFAMPYFIQVMK | 7.73 | 2.46E-03 | 0.0143 | 49 |
| Q68FD5 | CLH1_MOUSE ( | NNRPSEGPLQTR | 1.19 | 2.65E-03 | 0.0143 | 49 |
| Q68FD5 | CLH1_MOUSE ( | VMEYINR | 2.56 | 3.07E-03 | 0.0143 | 49 |
| Q68FD5 | CLH1_MOUSE ( | ADDPSSYMEVVQAANASGNWEELVK | 2.65 | 3.39E-03 | 0.0143 | 49 |
| Q68FD5 | CLH1_MOUSE ( | VGYTPDWIFLLR | 3.79 | 3.43E-03 | 0.0144 | 49 |
| Q68FD5 | CLH1_MOUSE ( | AHIAQLC[+57]EK | 1.44 | 3.65E-03 | 0.0146 | 49 |
| Q68FD5 | CLH1_MOUSE ( | GQC[+57]DLELINVC[+57]NENSLFK | 2.21 | 3.72E-03 | 0.0147 | 49 |
| Q68FD5 | CLH1_MOUSE ( | IHEGC[+57]EEPATHNALAK | 1.27 | 3.78E-03 | 0.0147 | 49 |
| Q68FD5 | CLH1_MOUSE ( | NNLAGAEELFAR | 1.62 | 3.92E-03 | 0.0147 | 49 |
| Q68FD5 | CLH1_MOUSE ( | GQFSTDELVAEVEK | 3.85 | 4.15E-03 | 0.0149 | 49 |
| Q68FD5 | CLH1_MOUSE ( | FQSVPAQPGQTSPLLQYFGILLDQGQLNK | 9.07 | 4.26E-03 | 0.0150 | 49 |
| Q68FD5 | CLH1_MOUSE ( | AHMGMFTELAILYSK | 5.44 | 4.79E-03 | 0.0154 | 49 |
| Q68FD5 | CLH1_MOUSE ( | EVC[+57]FAC[+57]VDGK | 1.46 | 4.88E-03 | 0.0155 | 49 |
| Q68FD5 | CLH1_MOUSE ( | YHEQLSTQSLIELFESFK | 8.25 | 4.97E-03 | 0.0155 | 49 |
| Q68FD5 | CLH1_MOUSE ( | VIQC[+57]FAETGQVQK | 2.35 | 6.31E-03 | 0.0169 | 49 |
| Q68FD5 | CLH1_MOUSE ( | ISGETIFVTAPHEATAGIIGVNR | 2.47 | 6.73E-03 | 0.0172 | 49 |
| Q68FD5 | CLH1_MOUSE ( | LASTLVHLGEYQAAVDGAR | 2.20 | 7.50E-03 | 0.0180 | 49 |
| Q68FD5 | CLH1_MOUSE ( | KFDVNTSAVQVLIEHIGNLDR | 5.48 | 7.60E-03 | 0.0181 | 49 |

| Protein Accession | Protein Description | Modified Peptide Sequence | co-IP:Ctrl | p value | q value | Sig Peptide Count |
|---|---|---|---|---|---|---|
| Q68FD5 | CLH1_MOUSE ( | KDPELWGSVLLESNPYR | 2.47 | 7.76E-03 | 0.0182 | 49 |
| Q68FD5 | CLH1_MOUSE ( | KFNALFAQGNYSEAAK | 1.69 | 7.93E-03 | 0.0183 | 49 |
| Q68FD5 | CLH1_MOUSE ( | ALEHFTDLYDIK | 2.24 | 7.96E-03 | 0.0183 | 49 |
| Q68FD5 | CLH1_MOUSE ( | LHIIEVGTPPTGNQPFPK | 2.35 | 8.23E-03 | 0.0186 | 49 |
| Q68FD5 | CLH1_MOUSE ( | VVGAMQLYSVDR | 1.97 | 8.41E-03 | 0.0188 | 49 |
| Q68FD5 | CLH1_MOUSE ( | LLYNNVSNFGR | 1.66 | 8.96E-03 | 0.0195 | 49 |
| Q68FD5 | CLH1_MOUSE ( | ALEHFTDLYDIKR | 2.79 | 1.04E-02 | 0.0211 | 49 |
| Q68FD5 | CLH1_MOUSE ( | VVGAM[+16]QLYSVDR | 1.44 | 1.19E-02 | 0.0227 | 49 |
| Q68FD5 | CLH1_MOUSE ( | LAELEEFINGPNNAHIQQVGDR | 1.70 | 1.24E-02 | 0.0231 | 49 |
| Q68FD5 | CLH1_MOUSE ( | IAAYLFK | 1.72 | 1.25E-02 | 0.0232 | 49 |
| Q68FD5 | CLH1_MOUSE ( | VANVELYYK | 1.74 | 1.26E-02 | 0.0233 | 49 |
| Q68FD5 | CLH1_MOUSE ( | YIEIYVQK | 2.20 | 1.34E-02 | 0.0242 | 49 |
| Q68FD5 | CLH1_MOUSE ( | HDVVFLITK | 2.06 | 1.41E-02 | 0.0250 | 49 |
| Q68FD5 | CLH1_MOUSE ( | WLLLTGISAQQNR | 3.06 | 1.48E-02 | 0.0258 | 49 |
| Q68FD5 | CLH1_MOUSE ( | TLQIFNIEMK | 2.64 | 1.51E-02 | 0.0262 | 49 |
| Q68FD5 | CLH1_MOUSE ( | LTDQLPLIIVC[+57]DR | 2.79 | 1.66E-02 | 0.0277 | 49 |
| Q6URW6 | MYH14_MOUSE | DLGEELEALRGELEDTLDSTNAQQELR | 7.49 | 2.49E-02 | 0.0370 | 49 |
| P05213 | TBA1B_MOUSE | DVNAAIATIK | 3.37 | 2.56E-02 | 0.0377 | 49 |
| P28660 | NCKP1_MOUSE | LKEFLALASSSLLK | 1.71 | 2.56E-02 | 0.0377 | 49 |
| Q62261 | SPTB2_MOUSE | MWEVLESTTQTK | 1.70 | 2.74E-02 | 0.0397 | 49 |
| P14873 | MAP1B_MOUSE | SSYYVVSGNDPAAEEPSR | 5.33 | 3.44E-04 | 0.0143 | 44 |
| P14873 | MAP1B_MOUSE | NLISPDLGVVFLNVPENLK | 6.01 | 7.32E-04 | 0.0143 | 44 |
| P14873 | MAP1B_MOUSE | QQDLNIMVLASSSTVVMQDESFPAC[+57]K | 5.31 | 1.17E-03 | 0.0143 | 44 |
| P14873 | MAP1B_MOUSE | NVDVEFFK | 2.95 | 1.33E-03 | 0.0143 | 44 |
| P14873 | MAP1B_MOUSE | AAEAGVTEEQYGYLGTSAK | 3.40 | 1.69E-03 | 0.0143 | 44 |
| P14873 | MAP1B_MOUSE | TTEAAATAVGTAATTAAVVAAAGIAASGPV | 3.38 | 2.22E-03 | 0.0143 | 44 |
| P14873 | MAP1B_MOUSE | SDISPLTPR | 4.54 | 3.09E-03 | 0.0143 | 44 |
| P14873 | MAP1B_MOUSE | TSDVETMSSQSALALDER | 4.11 | 3.42E-03 | 0.0144 | 44 |
| P14873 | MAP1B_MOUSE | ESSPLYSPGFSDSTSAAK | 4.88 | 3.88E-03 | 0.0147 | 44 |
| P14873 | MAP1B_MOUSE | SPSLSPSPPSPIEK | 3.86 | 3.99E-03 | 0.0148 | 44 |
| P14873 | MAP1B_MOUSE | DLTTSSVEK | 3.34 | 4.65E-03 | 0.0153 | 44 |
| P14873 | MAP1B_MOUSE | VLFPGNSTQYNILEGLEK | 4.14 | 4.70E-03 | 0.0153 | 44 |
| P14873 | MAP1B_MOUSE | VDSILLTHIGDDNLPGINSM[+16]LQR | 3.69 | 5.00E-03 | 0.0155 | 44 |
| P14873 | MAP1B_MOUSE | TPEEGGYSYEISEK | 3.01 | 5.01E-03 | 0.0155 | 44 |
| P14873 | MAP1B_MOUSE | SWDTNLIEC[+57]NLDQELK | 4.86 | 5.11E-03 | 0.0155 | 44 |
| P14873 | MAP1B_MOUSE | TPEVSGYTYEK | 3.63 | 6.12E-03 | 0.0166 | 44 |
| P14873 | MAP1B_MOUSE | TPQASTYSYETSDR | 4.06 | 6.59E-03 | 0.0171 | 44 |
| P14873 | MAP1B_MOUSE | NLISPDLGVVFLNVPENLKDPEPNIK | 5.20 | 6.64E-03 | 0.0172 | 44 |
| P14873 | MAP1B_MOUSE | TPEDGGYTC[+57]EITEK | 4.56 | 6.92E-03 | 0.0174 | 44 |
| P14873 | MAP1B_MOUSE | ESPVSDLTSTGLYQDKQEEK | 4.15 | 7.28E-03 | 0.0177 | 44 |
| P14873 | MAP1B_MOUSE | SPC[+57]DSGYSYETIEK | 4.75 | 7.30E-03 | 0.0178 | 44 |
| P14873 | MAP1B_MOUSE | SDVLETVVLINPSDEAVSTEVR | 4.61 | 7.56E-03 | 0.0180 | 44 |
| P14873 | MAP1B_MOUSE | DFEELKAEEIDVAK | 3.86 | 7.57E-03 | 0.0180 | 44 |
| P14873 | MAP1B_MOUSE | VDSILLTHIGDDNLPGINSMLQR | 4.40 | 7.63E-03 | 0.0181 | 44 |
| P14873 | MAP1B_MOUSE | EEGYEPDKTEAEDYVM[+16]AVADK | 3.93 | 8.14E-03 | 0.0185 | 44 |
| P14873 | MAP1B_MOUSE | DLTGQVPTPPVK | 3.43 | 8.81E-03 | 0.0193 | 44 |
| P14873 | MAP1B_MOUSE | YESSLYSQEYSKPAVASFNGLSEGSK | 5.10 | 9.25E-03 | 0.0198 | 44 |
| P14873 | MAP1B_MOUSE | DIKPQLELIEDEEK | 3.45 | 9.37E-03 | 0.0200 | 44 |
| P14873 | MAP1B_MOUSE | GDSALFAVNGFNMLINGGSER | 6.38 | 1.09E-02 | 0.0216 | 44 |
| P14873 | MAP1B_MOUSE | ASLTLFC[+57]PEEGDWK | 4.06 | 1.15E-02 | 0.0223 | 44 |
| P14873 | MAP1B_MOUSE | SVNFSLTPNEIK | 3.81 | 1.17E-02 | 0.0225 | 44 |
| P14873 | MAP1B_MOUSE | EMSLYASLASEK | 5.44 | 1.19E-02 | 0.0227 | 44 |
| P14873 | MAP1B_MOUSE | SVGNTIEPVILFQK | 4.64 | 1.21E-02 | 0.0229 | 44 |
| P14873 | MAP1B_MOUSE | AIGNIELGIR | 4.04 | 1.21E-02 | 0.0229 | 44 |
| P14873 | MAP1B_MOUSE | HNLQDFINIK | 3.51 | 1.28E-02 | 0.0235 | 44 |
| P14873 | MAP1B_MOUSE | LEMYVLNPVK | 4.44 | 1.34E-02 | 0.0242 | 44 |
| P14873 | MAP1B_MOUSE | THDVGGYYYEK | 3.82 | 1.39E-02 | 0.0248 | 44 |
| P14873 | MAP1B_MOUSE | M[+16]SISEGTVSDK | 3.62 | 1.51E-02 | 0.0262 | 44 |
| P14873 | MAP1B_MOUSE | QDVDLC[+57]LVSSC[+57]EFK | 4.49 | 1.55E-02 | 0.0266 | 44 |
| Q8BP67 | RL24_MOUSE 6 | VELC[+57]SFSGYK | 4.34 | 1.81E-02 | 0.0294 | 44 |
| Q61171 | PRDX2_MOUSE | GLFIIDAK | 4.21 | 2.12E-02 | 0.0330 | 44 |
| Q61316 | HSP74_MOUSE | FLEMC[+57]DDLLAR | 3.73 | 2.58E-02 | 0.0380 | 44 |

| Protein Accession | Protein Description | Modified Peptide Sequence | co-IP:Ctrl | p value | q value | Sig Peptide Count |
|---|---|---|---|---|---|---|
| Q68FD5 | CLH1_MOUSE ( | NLILVVR | 4.64 | 2.75E-02 | 0.0398 | 44 |
| P62889 | RL30_MOUSE 6 | KSEIEYYAMLAK | 1.83 | 3.59E-02 | 0.0495 | 44 |
| Q62261 | SPTB2_MOUSE | TQTAIASEDM[+16]PNTLTEAEK | 2.58 | 1.77E-04 | 0.0143 | 44 |
| Q62261 | SPTB2_MOUSE | DASVAEAWLLGQEPYLSSR | 5.00 | 2.36E-03 | 0.0143 | 44 |
| Q62261 | SPTB2_MOUSE | FANSLVGVQQQLQAFNTYR | 3.40 | 2.82E-03 | 0.0143 | 44 |
| Q62261 | SPTB2_MOUSE | LVSQDNFGFDLPAVEAATK | 3.71 | 3.20E-03 | 0.0143 | 44 |
| Q62261 | SPTB2_MOUSE | EAEKLESEHPDQAQAILSR | 3.67 | 3.21E-03 | 0.0143 | 44 |
| Q62261 | SPTB2_MOUSE | EAASELLM[+16]R | 2.80 | 3.36E-03 | 0.0143 | 44 |
| Q62261 | SPTB2_MOUSE | VIESTQDLGNDLAGVMALQR | 3.25 | 3.79E-03 | 0.0147 | 44 |
| Q62261 | SPTB2_MOUSE | LLEVLSGER | 3.07 | 3.98E-03 | 0.0148 | 44 |
| Q62261 | SPTB2_MOUSE | EVDDLEQWIAER | 4.19 | 4.75E-03 | 0.0154 | 44 |
| Q62261 | SPTB2_MOUSE | VLDNAIETEK | 2.33 | 4.88E-03 | 0.0155 | 44 |
| Q62261 | SPTB2_MOUSE | DLDDFQSWLSR | 3.61 | 4.93E-03 | 0.0155 | 44 |
| Q62261 | SPTB2_MOUSE | EGMQLISEKPETEAVVK | 3.97 | 5.02E-03 | 0.0155 | 44 |
| Q62261 | SPTB2_MOUSE | IVSSNDVGHDEYSTQSLVK | 3.25 | 5.15E-03 | 0.0155 | 44 |
| Q62261 | SPTB2_MOUSE | VAVVNQIAR | 2.78 | 5.17E-03 | 0.0155 | 44 |
| Q62261 | SPTB2_MOUSE | KHEAIETDIAAYEER | 4.21 | 5.39E-03 | 0.0158 | 44 |
| Q62261 | SPTB2_MOUSE | SALPAQSAATLPAR | 3.27 | 5.49E-03 | 0.0159 | 44 |
| Q62261 | SPTB2_MOUSE | LSDGNEYLFQAK | 2.96 | 5.67E-03 | 0.0160 | 44 |
| Q62261 | SPTB2_MOUSE | QALQDTLALYK | 3.72 | 6.36E-03 | 0.0169 | 44 |
| Q62261 | SPTB2_MOUSE | LVSDGNINSDR | 2.69 | 6.82E-03 | 0.0172 | 44 |
| Q62261 | SPTB2_MOUSE | GNLEVLLFTIQSK | 4.36 | 6.92E-03 | 0.0174 | 44 |
| Q62261 | SPTB2_MOUSE | DALLSALSIQNYHLEC[+57]NETK | 3.44 | 6.94E-03 | 0.0174 | 44 |
| Q62261 | SPTB2_MOUSE | HQILEQAVEDYAETVHQLSK | 4.19 | 7.85E-03 | 0.0182 | 44 |
| Q62261 | SPTB2_MOUSE | LQALDTGWNELHK | 3.63 | 7.91E-03 | 0.0183 | 44 |
| Q62261 | SPTB2_MOUSE | VQAVVAVAR | 3.54 | 8.25E-03 | 0.0186 | 44 |
| Q62261 | SPTB2_MOUSE | DVEDEILWVGER | 5.08 | 9.12E-03 | 0.0197 | 44 |
| Q62261 | SPTB2_MOUSE | DLTSVNILLK | 3.37 | 9.24E-03 | 0.0198 | 44 |
| Q62261 | SPTB2_MOUSE | SQNIITDSSSLNAEAIR | 4.22 | 9.50E-03 | 0.0202 | 44 |
| Q62261 | SPTB2_MOUSE | ALVADSHPESER | 4.93 | 1.13E-02 | 0.0221 | 44 |
| Q62261 | SPTB2_MOUSE | VAHMEFC[+57]YQELC[+57]QLAAER | 3.35 | 1.15E-02 | 0.0223 | 44 |
| Q62261 | SPTB2_MOUSE | DQNTVETLQR | 2.32 | 1.28E-02 | 0.0235 | 44 |
| Q62261 | SPTB2_MOUSE | AELFTQSC[+57]ADLDK | 4.06 | 1.34E-02 | 0.0242 | 44 |
| Q62261 | SPTB2_MOUSE | LTTLELLEVR | 3.46 | 1.36E-02 | 0.0244 | 44 |
| Q62261 | SPTB2_MOUSE | LFQLNR | 2.96 | 1.49E-02 | 0.0259 | 44 |
| Q62261 | SPTB2_MOUSE | LQQFLR | 1.98 | 1.54E-02 | 0.0265 | 44 |
| Q9D8E6 | RL4_MOUSE 60 | APIRPDIVNFVHTNLR | 5.13 | 1.72E-02 | 0.0284 | 44 |
| O08788 | DCTN1_MOUSE | VDELTTDLEILK | 3.49 | 2.07E-02 | 0.0324 | 44 |
| Q9CWS5 | Q9CWS5_MOUSE | VADALANAAGHLDDLPGALSALSDLHAHK | 4.32 | 2.15E-02 | 0.0334 | 44 |
| P62717 | RL18A_MOUSE | FWYFVSQLK | 4.43 | 2.23E-02 | 0.0343 | 44 |
| P63005 | LIS1_MOUSE P | LLASC[+57]SADMTIK | 2.33 | 2.50E-02 | 0.0371 | 44 |
| P20357 | MTAP2_MOUSE | LASVSADAEVAR | 6.12 | 2.53E-02 | 0.0374 | 44 |
| Q68FD5 | CLH1_MOUSE ( | IYIDSNNNPER | 4.02 | 2.58E-02 | 0.0380 | 44 |
| Q9DBG3 | AP2B1_MOUSE | LLSTDPVTAK | 2.32 | 2.74E-02 | 0.0397 | 44 |
| Q6PIE5 | AT1A2_MOUSE | DTAGDASESALLK | 2.64 | 2.79E-02 | 0.0403 | 44 |
| P80317 | TCPZ_MOUSE ¯ | ALQFLEQVK | 0.93 | 2.93E-02 | 0.0418 | 44 |
| P14873 | MAP1B_MOUSE | LGGDVSPTQIDVSQFGSFK | #REF! | #REF! | 0.0495 | 43 |
| Q9QYR6 | MAP1A_MOUSE | SEPQDFQEDSWGDTK | 3.17 | 3.11E-05 | 0.0143 | 41 |
| Q9QYR6 | MAP1A_MOUSE | DAEQTEPEQREPTPYPDER | 2.61 | 2.05E-04 | 0.0143 | 41 |
| Q9QYR6 | MAP1A_MOUSE | VPSAPGQESPVPDTK | 2.13 | 2.89E-04 | 0.0143 | 41 |
| Q9QYR6 | MAP1A_MOUSE | TRHDEYLEVTK | 1.94 | 5.71E-04 | 0.0143 | 41 |
| Q9QYR6 | MAP1A_MOUSE | VPEVTESHTTR | 1.56 | 7.07E-04 | 0.0143 | 41 |
| Q9QYR6 | MAP1A_MOUSE | TADQDFFR | 2.12 | 7.64E-04 | 0.0143 | 41 |
| Q9QYR6 | MAP1A_MOUSE | AQWGENLQVTLIPTHDTEVTR | 4.02 | 9.70E-04 | 0.0143 | 41 |
| Q9QYR6 | MAP1A_MOUSE | AVLDALLEGK | 3.42 | 9.90E-04 | 0.0143 | 41 |
| Q9QYR6 | MAP1A_MOUSE | REEVLEEGAK | 3.57 | 1.14E-03 | 0.0143 | 41 |
| Q9QYR6 | MAP1A_MOUSE | SAPC[+57]GSLAFSGDR | 2.59 | 1.19E-03 | 0.0143 | 41 |
| Q9QYR6 | MAP1A_MOUSE | EQKDEASEEKEQVLEQK | 2.70 | 1.21E-03 | 0.0143 | 41 |
| Q9QYR6 | MAP1A_MOUSE | NLISPELGVVFFNVPDK | 3.38 | 1.48E-03 | 0.0143 | 41 |
| Q9QYR6 | MAP1A_MOUSE | GDSALFAVNGFNILVDGGSDRK | 2.67 | 1.52E-03 | 0.0143 | 41 |
| Q9QYR6 | MAP1A_MOUSE | IDSVLLTHIGADNLPGINGLLQR | 4.49 | 1.93E-03 | 0.0143 | 41 |
| Q9QYR6 | MAP1A_MOUSE | EAELTPENIAAAR | 2.09 | 2.05E-03 | 0.0143 | 41 |

| Protein Accession | Protein Description | Modified Peptide Sequence | co-IP:Ctrl | p value | q value | Sig Peptide Count |
|---|---|---|---|---|---|---|
| Q9QYR6 | MAP1A_MOUSE | SPWASDFKDFQEPLPQK | 2.87 | 2.12E-03 | 0.0143 | 41 |
| Q9QYR6 | MAP1A_MOUSE | VVSNTIEPLTLFHK | 2.98 | 2.66E-03 | 0.0143 | 41 |
| Q9QYR6 | MAP1A_MOUSE | GAALQQTQAPEPR | 2.03 | 3.19E-03 | 0.0143 | 41 |
| Q9QYR6 | MAP1A_MOUSE | SPQAQDTLGSLAGGQTGC[+57]TIQLLPEQDK | 2.90 | 3.70E-03 | 0.0147 | 41 |
| Q9QYR6 | MAP1A_MOUSE | AVVFETGEAGAASGAGSLPGEVR | 2.31 | 3.88E-03 | 0.0147 | 41 |
| Q9QYR6 | MAP1A_MOUSE | EGEGGAGAPDSSSFSSK | 1.97 | 4.16E-03 | 0.0149 | 41 |
| Q9QYR6 | MAP1A_MOUSE | ELALSSPEDLTQDFEELKR | 4.25 | 4.87E-03 | 0.0155 | 41 |
| Q9QYR6 | MAP1A_MOUSE | DLAAGAVPANLKPSK | 2.25 | 4.92E-03 | 0.0155 | 41 |
| Q9QYR6 | MAP1A_MOUSE | NEPTTPSWLAEIPPWVPK | 4.79 | 5.38E-03 | 0.0158 | 41 |
| Q9QYR6 | MAP1A_MOUSE | LSSFATSVAEDQSVASLTAPQTEETGK | 2.35 | 5.63E-03 | 0.0160 | 41 |
| Q9QYR6 | MAP1A_MOUSE | ALGLEESPEEEGK | 3.93 | 5.78E-03 | 0.0162 | 41 |
| Q9QYR6 | MAP1A_MOUSE | LGIQAEPLYR | 1.83 | 6.77E-03 | 0.0172 | 41 |
| Q9QYR6 | MAP1A_MOUSE | LSKPC[+57]C[+57]YIFPGGR | 4.49 | 7.61E-03 | 0.0181 | 41 |
| Q9QYR6 | MAP1A_MOUSE | VLFPGNAPQNK | 2.10 | 8.00E-03 | 0.0183 | 41 |
| Q9QYR6 | MAP1A_MOUSE | APDSGAEVER | 1.92 | 8.63E-03 | 0.0191 | 41 |
| Q9QYR6 | MAP1A_MOUSE | EQDVVQGWR | 2.15 | 9.53E-03 | 0.0202 | 41 |
| Q9QYR6 | MAP1A_MOUSE | ESTFLDEGPNEQEITPLQHTPR | 2.92 | 9.61E-03 | 0.0203 | 41 |
| Q9QYR6 | MAP1A_MOUSE | ALALVPGTPTR | 1.57 | 9.65E-03 | 0.0203 | 41 |
| Q9QYR6 | MAP1A_MOUSE | SSLLLDTVTSIPSSR | 2.40 | 1.13E-02 | 0.0221 | 41 |
| Q9QYR6 | MAP1A_MOUSE | FPTSTYDLSGPEGPGPFEASQSAESAVPASSSK | 4.00 | 1.16E-02 | 0.0224 | 41 |
| Q9QYR6 | MAP1A_MOUSE | SPPC[+57]EDFSVTGESEK | 2.82 | 1.18E-02 | 0.0226 | 41 |
| Q9QYR6 | MAP1A_MOUSE | GDSALFAVNGFNILVDGGSDR | 5.82 | 1.26E-02 | 0.0233 | 41 |
| Q9QYR6 | MAP1A_MOUSE | QEPEPGPNVEPSFTPPAVPPR | 2.55 | 1.27E-02 | 0.0234 | 41 |
| Q9QYR6 | MAP1A_MOUSE | LDMYVLNPVK | 3.52 | 1.39E-02 | 0.0248 | 41 |
| Q9QYR6 | MAP1A_MOUSE | SFQYADIYEQMMLTGLGPAC[+57]PTR | 3.54 | 1.52E-02 | 0.0263 | 41 |
| Q8R191 | SNG3_MOUSE | AGAAFDPVSFAR | 2.28 | 3.30E-02 | 0.0461 | 41 |
| Q7TMM9 | TBB2A_MOUSE | FWEVISDEHGIDPTGSYHGDSDLQLER | 5.95 | 6.15E-04 | 0.0143 | 38 |
| Q7TMM9 | TBB2A_MOUSE | M[+16]SATFIGNSTAIQELFK | 2.08 | 8.51E-04 | 0.0143 | 38 |
| Q7TMM9 | TBB2A_MOUSE | GHYTEGAELVDSVLDVVRK | 3.70 | 8.82E-04 | 0.0143 | 38 |
| Q7TMM9 | TBB2A_MOUSE | M[+16]REIVHIQAGQC[+57]GNQIGAK | 2.53 | 1.29E-03 | 0.0143 | 38 |
| Q7TMM9 | TBB2A_MOUSE | KLAVNM[+16]VPFPR | 2.26 | 1.46E-03 | 0.0143 | 38 |
| Q7TMM9 | TBB2A_MOUSE | LAVNM[+16]VPFPR | 1.92 | 1.57E-03 | 0.0143 | 38 |
| Q7TMM9 | TBB2A_MOUSE | NM[+16]MAAC[+57]DPR | 2.01 | 1.91E-03 | 0.0143 | 38 |
| Q7TMM9 | TBB2A_MOUSE | NMM[+16]AAC[+57]DPR | 1.95 | 2.11E-03 | 0.0143 | 38 |
| Q7TMM9 | TBB2A_MOUSE | MSATFIGNSTAIQELFKR | 3.32 | 2.79E-03 | 0.0143 | 38 |
| Q7TMM9 | TBB2A_MOUSE | MSATFIGNSTAIQELFK | 3.20 | 2.92E-03 | 0.0143 | 38 |
| Q7TMM9 | TBB2A_MOUSE | LTTPTYGDLNHLVSATM[+16]SGVTTC[+57]LR | 2.76 | 3.17E-03 | 0.0143 | 38 |
| Q7TMM9 | TBB2A_MOUSE | FPGQLNADLRK | 3.04 | 3.85E-03 | 0.0147 | 38 |
| Q7TMM9 | TBB2A_MOUSE | EIVHIQAGQC[+57]GNQIGAK | 2.28 | 4.11E-03 | 0.0148 | 38 |
| Q7TMM9 | TBB2A_MOUSE | ESESC[+57]DC[+57]LQGFQLTHSLGGGTGSGM[+16]GTLLISK | 3.11 | 4.17E-03 | 0.0149 | 38 |
| Q7TMM9 | TBB2A_MOUSE | LTTPTYGDLNHLVSATMSGVTTC[+57]LR | 4.28 | 4.55E-03 | 0.0152 | 38 |
| Q7TMM9 | TBB2A_MOUSE | SGPFGQIFRPDNFVFGQSGAGNNWAK | 5.44 | 4.85E-03 | 0.0154 | 38 |
| Q7TMM9 | TBB2A_MOUSE | ALTVPELTQQMFDSK | 2.66 | 4.93E-03 | 0.0155 | 38 |
| Q7TMM9 | TBB2A_MOUSE | ALTVPELTQQM[+16]FDSK | 2.41 | 5.09E-03 | 0.0155 | 38 |
| Q7TMM9 | TBB2A_MOUSE | ISEQFTAM[+16]FR | 2.19 | 5.64E-03 | 0.0160 | 38 |
| Q7TMM9 | TBB2A_MOUSE | INVYYNEAAGNK | 2.31 | 7.49E-03 | 0.0180 | 38 |
| Q7TMM9 | TBB2A_MOUSE | LHFFM[+16]PGFAPLTSR | 3.20 | 7.50E-03 | 0.0180 | 38 |
| Q7TMM9 | TBB2A_MOUSE | KLAVNMVPFPR | 3.13 | 7.78E-03 | 0.0182 | 38 |
| Q7TMM9 | TBB2A_MOUSE | MSM[+16]KEVDEQM[+16]LNVQNK | 1.88 | 7.90E-03 | 0.0183 | 38 |
| Q7TMM9 | TBB2A_MOUSE | LHFFMPGFAPLTSR | 4.22 | 8.06E-03 | 0.0184 | 38 |
| Q7TMM9 | TBB2A_MOUSE | IM[+16]NTFSVMPSPK | 1.73 | 8.20E-03 | 0.0186 | 38 |
| Q7TMM9 | TBB2A_MOUSE | FPGQLNADLR | 2.21 | 8.23E-03 | 0.0186 | 38 |
| Q7TMM9 | TBB2A_MOUSE | AILVDLEPGTM[+16]DSVR | 1.99 | 8.65E-03 | 0.0191 | 38 |
| Q7TMM9 | TBB2A_MOUSE | LAVNMVPFPR | 2.81 | 8.72E-03 | 0.0191 | 38 |
| Q7TMM9 | TBB2A_MOUSE | MREIVHIQAGQC[+57]GNQIGAK | 3.06 | 9.60E-03 | 0.0203 | 38 |
| Q7TMM9 | TBB2A_MOUSE | GHYTEGAELVDSVLDVVR | 4.40 | 9.70E-03 | 0.0204 | 38 |
| Q7TMM9 | TBB2A_MOUSE | AILVDLEPGTMDSVR | 2.65 | 1.00E-02 | 0.0206 | 38 |
| Q7TMM9 | TBB2A_MOUSE | ESESC[+57]DC[+57]LQGFQLTHSLGGGTGSGMGTLLISK | 3.72 | 1.29E-02 | 0.0237 | 38 |
| Q7TMM9 | TBB2A_MOUSE | ISEQFTAMFR | 3.55 | 1.38E-02 | 0.0247 | 38 |
| Q7TMM9 | TBB2A_MOUSE | IMNTFSVM[+16]PSPK | 1.80 | 1.40E-02 | 0.0249 | 38 |
| Q62167 | DDX3X_MOUSE | TAAFLLPILSQIYADGPGEALR | 2.56 | 1.73E-02 | 0.0284 | 38 |
| P16546 | SPTN1_MOUSE | ENLLEEQGSIALR | 3.03 | 1.88E-02 | 0.0301 | 38 |

| Protein Accession | Protein Description | Modified Peptide Sequence | co-IP:Ctrl | p value | q value | Sig Peptide Count |
|---|---|---|---|---|---|---|
| P26039 | TLN1_MOUSE | IGITNHDEYSLVR | 4.36 | 2.21E-02 | 0.0341 | 38 |
| P39053 | DYN1_MOUSE | GM[+16]EDLIPLVNR | 2.67 | 2.73E-02 | 0.0396 | 38 |
| P46460 | NSF_MOUSE | LFADAEEEQR | 3.05 | 2.19E-04 | 0.0143 | 30 |
| P46460 | NSF_MOUSE | GSM[+16]AGSTGVHDTVVNQLLSK | 2.59 | 1.68E-03 | 0.0143 | 30 |
| P46460 | NSF_MOUSE | IEVGLVVGNSQVAFEK | 3.85 | 2.20E-03 | 0.0143 | 30 |
| P46460 | NSF_MOUSE | M[+16]EIGLPDEK | 2.58 | 2.42E-03 | 0.0143 | 30 |
| P46460 | NSF_MOUSE | QKIEVGLVVGNSQVAFEK | 3.65 | 3.04E-03 | 0.0143 | 30 |
| P46460 | NSF_MOUSE | M[+16]GIGGLDKEFSDIFR | 3.04 | 3.34E-03 | 0.0143 | 30 |
| P46460 | NSF_MOUSE | QC[+57]IGTMTIEIDFLQK | 3.90 | 3.38E-03 | 0.0143 | 30 |
| P46460 | NSF_MOUSE | NFSGAELEGLVR | 2.89 | 4.04E-03 | 0.0148 | 30 |
| P46460 | NSF_MOUSE | TPLVSVLLEGPPHSGK | 3.18 | 4.10E-03 | 0.0148 | 30 |
| P46460 | NSF_MOUSE | AESLQVTR | 2.66 | 4.47E-03 | 0.0151 | 30 |
| P46460 | NSF_MOUSE | IAEESNFPFIK | 3.07 | 4.62E-03 | 0.0153 | 30 |
| P46460 | NSF_MOUSE | LLDYVPIGPR | 3.03 | 5.16E-03 | 0.0155 | 30 |
| P46460 | NSF_MOUSE | MGIGGLDKEFSDIFR | 5.19 | 5.24E-03 | 0.0157 | 30 |
| P46460 | NSF_MOUSE | GILLYGPPGC[+57]GK | 2.75 | 5.58E-03 | 0.0160 | 30 |
| P46460 | NSF_MOUSE | AENSSLNLIGK | 2.46 | 6.39E-03 | 0.0169 | 30 |
| P46460 | NSF_MOUSE | LGANSGLHIIIFDEIDAIC[+57]K | 4.25 | 6.47E-03 | 0.0170 | 30 |
| P46460 | NSF_MOUSE | SQLSC[+57]VVVDDIER | 2.70 | 6.57E-03 | 0.0171 | 30 |
| P46460 | NSF_MOUSE | KLLIIGTTSR | 3.73 | 6.64E-03 | 0.0172 | 30 |
| P46460 | NSF_MOUSE | VLDDGELLVQQTK | 2.26 | 6.68E-03 | 0.0172 | 30 |
| P46460 | NSF_MOUSE | C[+57]PTDELSLSNC[+57]AVVNEK | 3.34 | 6.69E-03 | 0.0172 | 30 |
| P46460 | NSF_MOUSE | YVGESEANIR | 2.68 | 6.73E-03 | 0.0172 | 30 |
| P46460 | NSF_MOUSE | NIDSNPYDTDK | 2.66 | 7.61E-03 | 0.0181 | 30 |
| P46460 | NSF_MOUSE | LFGLLVK | 3.76 | 7.77E-03 | 0.0182 | 30 |
| P46460 | NSF_MOUSE | VVNGPEILNK | 2.38 | 9.09E-03 | 0.0197 | 30 |
| P46460 | NSF_MOUSE | LLIIGTTSR | 2.49 | 1.16E-02 | 0.0224 | 30 |
| P46460 | NSF_MOUSE | GHQLLSADVDIK | 3.20 | 1.36E-02 | 0.0244 | 30 |
| P46460 | NSF_MOUSE | WGDPVTR | 2.73 | 1.58E-02 | 0.0269 | 30 |
| Q9Z1G4 | VPP1_MOUSE | QAEIENPLEDPVTGDYVHK | 1.88 | 1.92E-02 | 0.0306 | 30 |
| Q93092 | TALDO_MOUSE | LFVLFGAEILK | 5.55 | 2.39E-02 | 0.0360 | 30 |
| P12970 | RL7A_MOUSE | NFGIGQDIQPK | 4.46 | 2.43E-02 | 0.0364 | 30 |
| P63017 | HSP7C_MOUSE | RFDDAVVQSDMK | 3.51 | 2.36E-04 | 0.0143 | 30 |
| P63017 | HSP7C_MOUSE | C[+57]NEIISWLDK | 1.46 | 9.53E-04 | 0.0143 | 30 |
| P63017 | HSP7C_MOUSE | STAGDTHLGGEDFDNR | 1.96 | 1.27E-03 | 0.0143 | 30 |
| P63017 | HSP7C_MOUSE | NQVAM[+16]NPTNTVFDAK | 1.86 | 1.37E-03 | 0.0143 | 30 |
| P63017 | HSP7C_MOUSE | SINPDEAVAYGAAVQAAILSGDK | 3.09 | 2.01E-03 | 0.0143 | 30 |
| P63017 | HSP7C_MOUSE | NSLESYAFNM[+16]K | 2.18 | 2.24E-03 | 0.0143 | 30 |
| P63017 | HSP7C_MOUSE | MVNHFIAEFKR | 2.75 | 2.75E-03 | 0.0143 | 30 |
| P63017 | HSP7C_MOUSE | DAGTIAGLNVLR | 2.22 | 2.99E-03 | 0.0143 | 30 |
| P63017 | HSP7C_MOUSE | LLQDFFNGK | 2.31 | 3.28E-03 | 0.0143 | 30 |
| P63017 | HSP7C_MOUSE | VQVEYK | 2.31 | 3.34E-03 | 0.0143 | 30 |
| P63017 | HSP7C_MOUSE | FDDAVVQSDM[+16]K | 1.63 | 3.35E-03 | 0.0143 | 30 |
| P63017 | HSP7C_MOUSE | FELTGIPPAPR | 2.94 | 3.84E-03 | 0.0147 | 30 |
| P63017 | HSP7C_MOUSE | FEELNADLFR | 2.39 | 3.95E-03 | 0.0148 | 30 |
| P63017 | HSP7C_MOUSE | GPAVGIDLGTTYSC[+57]VGVFQHGK | 3.42 | 4.05E-03 | 0.0148 | 30 |
| P63017 | HSP7C_MOUSE | TTPSYVAFTDTER | 2.21 | 4.22E-03 | 0.0149 | 30 |
| P63017 | HSP7C_MOUSE | EIAEAYLGK | 1.74 | 4.46E-03 | 0.0151 | 30 |
| P63017 | HSP7C_MOUSE | FDDAVVQSDMK | 2.39 | 5.21E-03 | 0.0156 | 30 |
| P63017 | HSP7C_MOUSE | LDKSQIHDIVLVGGSTR | 3.22 | 5.65E-03 | 0.0160 | 30 |
| P63017 | HSP7C_MOUSE | SFYPEEVSSMVLTK | 2.58 | 6.65E-03 | 0.0172 | 30 |
| P63017 | HSP7C_MOUSE | NSLESYAFNMK | 2.60 | 6.79E-03 | 0.0172 | 30 |
| P63017 | HSP7C_MOUSE | SQIHDIVLVGGSTR | 2.06 | 7.22E-03 | 0.0177 | 30 |
| P63017 | HSP7C_MOUSE | QTQTFTTYSDNQPGVLIQVYEGER | 3.49 | 7.23E-03 | 0.0177 | 30 |
| P63017 | HSP7C_MOUSE | MKEIAEAYLGK | 2.54 | 7.65E-03 | 0.0181 | 30 |
| P63017 | HSP7C_MOUSE | ARFEELNADLFR | 3.32 | 8.71E-03 | 0.0191 | 30 |
| P63017 | HSP7C_MOUSE | HWPFMVVNDAGRPK | 4.01 | 9.34E-03 | 0.0199 | 30 |
| P63017 | HSP7C_MOUSE | HWPFM[+16]VVNDAGRPK | 2.84 | 9.68E-03 | 0.0204 | 30 |
| P63017 | HSP7C_MOUSE | TVTNAVVTVPAYFNDSQR | 2.68 | 1.11E-02 | 0.0219 | 30 |
| P63017 | HSP7C_MOUSE | M[+16]VNHFIAEFK | 2.42 | 1.29E-02 | 0.0237 | 30 |
| P63017 | HSP7C_MOUSE | MVNHFIAEFK | 2.39 | 1.57E-02 | 0.0268 | 30 |
| P63017 | HSP7C_MOUSE | NQVAMNPTNTVFDAK | 2.19 | 1.69E-02 | 0.0281 | 30 |

| Protein Accession | Protein Description | Modified Peptide Sequence | co-IP:Ctrl | p value | q value | Sig Peptide Count |
|---|---|---|---|---|---|---|
| Q9QXS6 | DREB_MOUSE | EQSIFGDQRDEEEESQM[+16]KK | 4.46 | 1.38E-03 | 0.0143 | 26 |
| Q9QXS6 | DREB_MOUSE | SPSDSSTASTPIAEQIER | 3.64 | 1.44E-03 | 0.0143 | 26 |
| Q9QXS6 | DREB_MOUSE | M[+16]APTPIPTR | 3.38 | 1.57E-03 | 0.0143 | 26 |
| Q9QXS6 | DREB_MOUSE | LREDENAEPVGTTYQK | 4.08 | 1.71E-03 | 0.0143 | 26 |
| Q9QXS6 | DREB_MOUSE | LKEQSIFGDQRDEEEESQM[+16]K | 5.59 | 1.96E-03 | 0.0143 | 26 |
| Q9QXS6 | DREB_MOUSE | ASDSGPSSSSSSSSSPPR | 6.45 | 2.90E-03 | 0.0143 | 26 |
| Q9QXS6 | DREB_MOUSE | TPNLSSSLPC[+57]SHLDSHR | 4.02 | 3.31E-03 | 0.0143 | 26 |
| Q9QXS6 | DREB_MOUSE | EQFWEQAK | 3.90 | 3.95E-03 | 0.0148 | 26 |
| Q9QXS6 | DREB_MOUSE | TPFPYITC[+57]HR | 4.13 | 4.16E-03 | 0.0149 | 26 |
| Q9QXS6 | DREB_MOUSE | SESEVEEAAAIIAQRPDNPR | 4.41 | 4.50E-03 | 0.0152 | 26 |
| Q9QXS6 | DREB_MOUSE | LAASGEGGLQELSGHFENQK | 4.66 | 4.54E-03 | 0.0152 | 26 |
| Q9QXS6 | DREB_MOUSE | LELLAAYEEVIR | 5.17 | 4.56E-03 | 0.0152 | 26 |
| Q9QXS6 | DREB_MOUSE | KSESEVEEAAAIIAQRPDNPR | 4.84 | 4.58E-03 | 0.0152 | 26 |
| Q9QXS6 | DREB_MOUSE | TDAAVEMK | 3.67 | 4.82E-03 | 0.0154 | 26 |
| Q9QXS6 | DREB_MOUSE | MAPTPIPTR | 4.02 | 4.84E-03 | 0.0154 | 26 |
| Q9QXS6 | DREB_MOUSE | KQQSLEAEEAK | 4.00 | 5.12E-03 | 0.0155 | 26 |
| Q9QXS6 | DREB_MOUSE | EGTQASEGYFSQSQEEEFAQSEEPC[+57]AK | 4.95 | 5.15E-03 | 0.0155 | 26 |
| Q9QXS6 | DREB_MOUSE | EESAADWALYTYEDGSDDLK | 4.61 | 5.95E-03 | 0.0163 | 26 |
| Q9QXS6 | DREB_MOUSE | EREQQIEEHR | 4.87 | 8.09E-03 | 0.0184 | 26 |
| Q9QXS6 | DREB_MOUSE | LKEQSIFGDQRDEEEESQMK | 5.60 | 8.18E-03 | 0.0186 | 26 |
| Q9QXS6 | DREB_MOUSE | EQSIFGDQRDEEEESQMK | 4.75 | 8.84E-03 | 0.0193 | 26 |
| Q9QXS6 | DREB_MOUSE | DSQAALPK | 3.70 | 9.10E-03 | 0.0197 | 26 |
| Q9QXS6 | DREB_MOUSE | YVLINWVGEDVPDARK | 4.42 | 1.16E-02 | 0.0224 | 26 |
| Q9QXS6 | DREB_MOUSE | YVLINWVGEDVPDAR | 5.14 | 1.25E-02 | 0.0232 | 26 |
| Q9QXS6 | DREB_MOUSE | VMYGFC[+57]SVK | 4.13 | 1.49E-02 | 0.0259 | 26 |
| Q9QXS6 | DREB_MOUSE | C[+57]AC[+57]ASHVAK | 4.58 | 1.57E-02 | 0.0268 | 26 |
| Q64331 | MYO6_MOUSE | QARPTYATAMLQNLLK | 3.49 | 7.79E-04 | 0.0143 | 25 |
| Q64331 | MYO6_MOUSE | ALGLNEVDYK | 2.98 | 1.21E-03 | 0.0143 | 25 |
| Q64331 | MYO6_MOUSE | QREEESQQQAVLAQEC[+57]R | 5.58 | 1.26E-03 | 0.0143 | 25 |
| Q64331 | MYO6_MOUSE | SSEDLLSALQK | 5.06 | 1.53E-03 | 0.0143 | 25 |
| Q64331 | MYO6_MOUSE | VNLWLVC[+57]SR | 2.92 | 1.67E-03 | 0.0143 | 25 |
| Q64331 | MYO6_MOUSE | SAPSLEYC[+57]AELLGLDQDDLR | 2.77 | 1.68E-03 | 0.0143 | 25 |
| Q64331 | MYO6_MOUSE | IVEANPLLEAFGNAK | 4.84 | 1.70E-03 | 0.0143 | 25 |
| Q64331 | MYO6_MOUSE | FNEVVSALK | 3.52 | 2.05E-03 | 0.0143 | 25 |
| Q64331 | MYO6_MOUSE | STGASFIR | 5.35 | 2.07E-03 | 0.0143 | 25 |
| Q64331 | MYO6_MOUSE | GAEILPR | 4.92 | 2.16E-03 | 0.0143 | 25 |
| Q64331 | MYO6_MOUSE | LC[+57]AGASEDIR | 3.51 | 2.49E-03 | 0.0143 | 25 |
| Q64331 | MYO6_MOUSE | LSFISVGNK | 4.65 | 2.72E-03 | 0.0143 | 25 |
| Q64331 | MYO6_MOUSE | NLEISIDALMAK | 5.85 | 2.78E-03 | 0.0143 | 25 |
| Q64331 | MYO6_MOUSE | ILKEEQELYQK | 6.75 | 2.78E-03 | 0.0143 | 25 |
| Q64331 | MYO6_MOUSE | TQLNLLLDK | 5.21 | 2.79E-03 | 0.0143 | 25 |
| Q64331 | MYO6_MOUSE | LQQFFNER | 4.21 | 2.89E-03 | 0.0143 | 25 |
| Q64331 | MYO6_MOUSE | IAQNESELISDEAQGDM[+16]ALR | 4.85 | 2.90E-03 | 0.0143 | 25 |
| Q64331 | MYO6_MOUSE | TVYSHLFDHVVNR | 5.35 | 3.39E-03 | 0.0143 | 25 |
| Q64331 | MYO6_MOUSE | LVGILDILDEENR | 5.67 | 3.94E-03 | 0.0147 | 25 |
| Q64331 | MYO6_MOUSE | NLRDDEGFIIR | 2.02 | 4.42E-03 | 0.0151 | 25 |
| Q64331 | MYO6_MOUSE | VQWC[+57]SLSVIK | 4.07 | 4.53E-03 | 0.0152 | 25 |
| Q64331 | MYO6_MOUSE | NNDALHMSLESLIC[+57]ESR | 6.21 | 6.15E-03 | 0.0166 | 25 |
| Q64331 | MYO6_MOUSE | C[+57]GGIQYLQSAIESR | 2.15 | 1.51E-02 | 0.0262 | 25 |
| Q64331 | MYO6_MOUSE | DTINTSC[+57]DIELLAAC[+57]R | 6.14 | 1.58E-02 | 0.0269 | 25 |
| Q9ESJ4 | SPN90_MOUSE | LLLLLNR | 3.49 | 1.92E-02 | 0.0306 | 25 |
| Q7TSJ2 | MAP6_MOUSE | AVAIETQPAQGESDAVAR | 3.42 | 3.32E-04 | 0.0143 | 23 |
| Q7TSJ2 | MAP6_MOUSE | TEGHEETPLPPAQSQTQEGGPAAGK | 4.21 | 5.17E-04 | 0.0143 | 23 |
| Q7TSJ2 | MAP6_MOUSE | ATGPAPGPSVDRETVAAPGR | 2.80 | 7.87E-04 | 0.0143 | 23 |
| Q7TSJ2 | MAP6_MOUSE | SEYQPSDAPFER | 3.24 | 1.42E-03 | 0.0143 | 23 |
| Q7TSJ2 | MAP6_MOUSE | NQDPIIPVPLK | 2.81 | 1.56E-03 | 0.0143 | 23 |
| Q7TSJ2 | MAP6_MOUSE | DQSFPTPAPR | 2.39 | 2.27E-03 | 0.0143 | 23 |
| Q7TSJ2 | MAP6_MOUSE | SQDPIIPALAK | 2.76 | 2.33E-03 | 0.0143 | 23 |
| Q7TSJ2 | MAP6_MOUSE | GQDPLVPAPTK | 1.84 | 2.44E-03 | 0.0143 | 23 |
| Q7TSJ2 | MAP6_MOUSE | ADIAVPLVFTK | 3.28 | 2.51E-03 | 0.0143 | 23 |
| Q7TSJ2 | MAP6_MOUSE | DQGAVLLGPVK | 2.39 | 2.56E-03 | 0.0143 | 23 |
| Q7TSJ2 | MAP6_MOUSE | VQDHIASELLK | 2.33 | 2.62E-03 | 0.0143 | 23 |

| Protein Accession | Protein Description | Modified Peptide Sequence | co-IP:Ctrl | p value | q value | Sig Peptide Count |
|---|---|---|---|---|---|---|
| Q7TSJ2 | MAP6_MOUSE | EEVASTVSSSYR | 3.00 | 2.88E-03 | 0.0143 | 23 |
| Q7TSJ2 | MAP6_MOUSE | NQGLAGPELVK | 2.23 | 3.13E-03 | 0.0143 | 23 |
| Q7TSJ2 | MAP6_MOUSE | DPEGAGGAGVLAAGK | 3.79 | 3.26E-03 | 0.0143 | 23 |
| Q7TSJ2 | MAP6_MOUSE | ATGPAPGPSVDR | 2.76 | 3.27E-03 | 0.0143 | 23 |
| Q7TSJ2 | MAP6_MOUSE | YSEATEHPGAPPQPPAPLQPALAPPSR | 4.18 | 3.30E-03 | 0.0143 | 23 |
| Q7TSJ2 | MAP6_MOUSE | GHDSVFVAPVK | 2.76 | 3.39E-03 | 0.0143 | 23 |
| Q7TSJ2 | MAP6_MOUSE | AGPAWM[+16]VTR | 2.05 | 3.88E-03 | 0.0147 | 23 |
| Q7TSJ2 | MAP6_MOUSE | SGLGLGAASASTSGSGPADSVM[+16]R | 3.62 | 4.65E-03 | 0.0153 | 23 |
| Q7TSJ2 | MAP6_MOUSE | AGPAWMVR | 2.63 | 6.08E-03 | 0.0165 | 23 |
| Q7TSJ2 | MAP6_MOUSE | AQSPLLPEPLKNQSPVVPASTK | 3.34 | 1.07E-02 | 0.0214 | 23 |
| Q7TSJ2 | MAP6_MOUSE | AGPAWMVTR | 2.74 | 1.46E-02 | 0.0256 | 23 |
| Q8BPN8 | DMXL2_MOUSE | VGC[+57]PVLALEVLSK | 2.99 | 3.10E-02 | 0.0437 | 23 |
| P17426 | AP2A1_MOUSE | LLGFGSALLDNVDPNPENFVGAGIIQTK | 4.14 | 7.44E-04 | 0.0143 | 22 |
| P17426 | AP2A1_MOUSE | NSGVLFENQLLQIGVK | 2.92 | 1.39E-03 | 0.0143 | 22 |
| P17426 | AP2A1_MOUSE | LPVTINK | 2.15 | 1.42E-03 | 0.0143 | 22 |
| P17426 | AP2A1_MOUSE | FINLFPETK | 2.17 | 1.84E-03 | 0.0143 | 22 |
| P17426 | AP2A1_MOUSE | IVSSASTDLQDYTYYFVPAPWLSVK | 4.10 | 2.07E-03 | 0.0143 | 22 |
| P17426 | AP2A1_MOUSE | VGGYILGEFGNLIAGDPR | 3.74 | 2.66E-03 | 0.0143 | 22 |
| P17426 | AP2A1_MOUSE | AC[+57]NQLGQFLQHR | 2.14 | 3.04E-03 | 0.0143 | 22 |
| P17426 | AP2A1_MOUSE | TSVQFQNFLPTVVHPGDLQTQLAVQTK | 4.38 | 3.41E-03 | 0.0143 | 22 |
| P17426 | AP2A1_MOUSE | NADVELQQR | 1.67 | 9.15E-03 | 0.0197 | 22 |
| P17426 | AP2A1_MOUSE | ALLLSTYIK | 2.10 | 9.97E-03 | 0.0206 | 22 |
| P17426 | AP2A1_MOUSE | HLC[+57]ELLAQQF | 2.65 | 1.03E-02 | 0.0209 | 22 |
| P17426 | AP2A1_MOUSE | VLQIVTNRDDVQGYAAK | 1.84 | 1.18E-02 | 0.0226 | 22 |
| P17426 | AP2A1_MOUSE | LVEC[+57]LETVLNK | 1.73 | 1.31E-02 | 0.0239 | 22 |
| P17426 | AP2A1_MOUSE | THIDTVINALK | 2.88 | 1.45E-02 | 0.0255 | 22 |
| P17426 | AP2A1_MOUSE | IAGDYVSEEVWYR | 2.69 | 1.46E-02 | 0.0256 | 22 |
| P17426 | AP2A1_MOUSE | GLAVFISDIR | 1.75 | 1.66E-02 | 0.0277 | 22 |
| Q8JZQ9 | EIF3B_MOUSE | FSHQGVQLIDFSPC[+57]ER | 2.95 | 1.71E-02 | 0.0283 | 22 |
| P17183 | ENOG_MOUSE | AAVPSGASTGIYEALELR | 1.21 | 1.76E-02 | 0.0288 | 22 |
| Q6ZWN5 | RS9_MOUSE 4( | LDYILGLK | 1.79 | 2.00E-02 | 0.0316 | 22 |
| Q9Z1B3 | PLCB1_MOUSE | RLPLEILEFVQEAMK | 2.04 | 2.06E-02 | 0.0323 | 22 |
| P63330 | PP2AA_MOUSE | ELDQWIEQLNEC[+57]K | 3.15 | 3.13E-02 | 0.0440 | 22 |
| P39053 | DYN1_MOUSE I | DMLMQFVTK | 1.26 | 3.40E-02 | 0.0473 | 22 |
| Q9JHU4 | DYHC1_MOUSE | KLVPLLLEDGGDAPAALEAALEEK | 3.63 | 8.11E-07 | 0.0037 | 21 |
| Q9JHU4 | DYHC1_MOUSE | FQSISTEFLALMK | 2.85 | 2.87E-05 | 0.0143 | 21 |
| Q9JHU4 | DYHC1_MOUSE | LNTQEIFDDWAR | 1.90 | 6.09E-05 | 0.0143 | 21 |
| Q9JHU4 | DYHC1_MOUSE | ENFIPTIVNFSAEEISDAIR | 6.26 | 9.84E-04 | 0.0143 | 21 |
| Q9JHU4 | DYHC1_MOUSE | VNFLPEIITLSK | 1.96 | 1.09E-03 | 0.0143 | 21 |
| Q9JHU4 | DYHC1_MOUSE | ILDDDTIITTLENLK | 3.05 | 1.72E-03 | 0.0143 | 21 |
| Q9JHU4 | DYHC1_MOUSE | LSLSNAISTVLPLTQLR | 2.60 | 2.06E-03 | 0.0143 | 21 |
| Q9JHU4 | DYHC1_MOUSE | LVPLLLEDGGDAPAALEAALEEK | 2.27 | 2.89E-03 | 0.0143 | 21 |
| Q9JHU4 | DYHC1_MOUSE | TYAEPLTAAMVEFYTMSQER | 3.46 | 3.08E-03 | 0.0143 | 21 |
| Q9JHU4 | DYHC1_MOUSE | HVPVVYVDYPGPASLTQIYGTFNR | 2.83 | 3.19E-03 | 0.0143 | 21 |
| Q9JHU4 | DYHC1_MOUSE | RAPVIDADKPVSSQLR | 1.65 | 3.21E-03 | 0.0143 | 21 |
| Q9JHU4 | DYHC1_MOUSE | IFVFEPPPGVK | 1.54 | 3.93E-03 | 0.0147 | 21 |
| Q9JHU4 | DYHC1_MOUSE | VTDFGDKVEDPTFLNQLQSGVNR | 1.93 | 4.97E-03 | 0.0155 | 21 |
| Q9JHU4 | DYHC1_MOUSE | VTFVNFTVTR | 2.01 | 5.01E-03 | 0.0155 | 21 |
| Q9JHU4 | DYHC1_MOUSE | QLQNISQAAASGGAK | 1.55 | 5.20E-03 | 0.0156 | 21 |
| Q9JHU4 | DYHC1_MOUSE | VLLTTQGVDM[+16]ISK | 2.85 | 5.87E-03 | 0.0163 | 21 |
| Q9JHU4 | DYHC1_MOUSE | LAETVFNFQEK | 1.55 | 1.11E-02 | 0.0219 | 21 |
| Q9JHU4 | DYHC1_MOUSE | FYFVGDEDLLEIIGNSK | 4.86 | 1.13E-02 | 0.0221 | 21 |
| P17427 | AP2A2_MOUSE | THIETVINALK | 1.09 | 2.30E-02 | 0.0350 | 21 |
| Q61879 | MYH10_MOUSE | C[+57]MLQDREDQSILC[+57]TGESGAGK | 4.31 | 2.68E-02 | 0.0391 | 21 |
| Q9Z0E0 | NCDN_MOUSE | IPILSTFLTAR | 3.88 | 2.85E-02 | 0.0409 | 21 |
| Q8BH44 | COR2B_MOUSE | NM[+16]TEALLELHGHSR | 5.19 | 1.50E-03 | 0.0143 | 19 |
| Q8BH44 | COR2B_MOUSE | VC[+57]GHQGNVLDIK | 4.37 | 1.56E-03 | 0.0143 | 19 |
| Q8BH44 | COR2B_MOUSE | NVHDNHFC[+57]AVNAR | 4.51 | 1.64E-03 | 0.0143 | 19 |
| Q8BH44 | COR2B_MOUSE | WNPFIDNIIASC[+57]SEDTSVR | 5.79 | 1.97E-03 | 0.0143 | 19 |
| Q8BH44 | COR2B_MOUSE | VLQEANC[+57]K | 4.64 | 2.37E-03 | 0.0143 | 19 |
| Q8BH44 | COR2B_MOUSE | YYEISTEKPYLSYLM[+16]EFR | 5.31 | 2.70E-03 | 0.0143 | 19 |
| Q8BH44 | COR2B_MOUSE | TENELLR | 4.28 | 2.93E-03 | 0.0143 | 19 |

| Protein Accession | Protein Description | Modified Peptide Sequence | co-IP:Ctrl | p value | q value | Sig Peptide Count |
|---|---|---|---|---|---|---|
| Q8BH44 | COR2B_MOUSE | DPVLM[+16]SLK | 3.44 | 3.04E-03 | 0.0143 | 19 |
| Q8BH44 | COR2B_MOUSE | GLIEPISMIVPR | 5.10 | 3.83E-03 | 0.0147 | 19 |
| Q8BH44 | COR2B_MOUSE | VLIWNLDIGEPVK | 5.81 | 4.49E-03 | 0.0151 | 19 |
| Q8BH44 | COR2B_MOUSE | IWEIPDGGLK | 4.19 | 6.46E-03 | 0.0170 | 19 |
| Q8BH44 | COR2B_MOUSE | QLQLELK | 3.91 | 7.04E-03 | 0.0175 | 19 |
| Q8BH44 | COR2B_MOUSE | YYEISTEKPYLSYLMEFF | 7.51 | 9.74E-03 | 0.0204 | 19 |
| Q8BH44 | COR2B_MOUSE | GLIEPISM[+16]IVPR | 4.18 | 1.04E-02 | 0.0211 | 19 |
| Q8BH44 | COR2B_MOUSE | DPVLMSLK | 4.29 | 1.08E-02 | 0.0215 | 19 |
| Q8BH44 | COR2B_MOUSE | FLAIVTESAGGGSFLVIPLEQTGR | 2.67 | 1.11E-02 | 0.0219 | 19 |
| Q8BH44 | COR2B_MOUSE | VGLVEWHPTTNNILFSAGYDYK | 6.00 | 1.69E-02 | 0.0281 | 19 |
| P99029 | PRDX5_MOUSE | ALNVEPDGTGLTC[+57]SLAPNILSQL | 6.64 | 1.85E-02 | 0.0298 | 19 |
| O35643 | AP1B1_MOUSE | YNDPIYVK | 2.45 | 2.90E-02 | 0.0414 | 19 |
| Q9JKK7 | TMOD2_MOUSE | QQLGTAVEM[+16]EIAQM[+16]LEENSR | 3.82 | 2.31E-04 | 0.0143 | 19 |
| Q9JKK7 | TMOD2_MOUSE | NIDEDELLGK | 3.78 | 2.54E-04 | 0.0143 | 19 |
| Q9JKK7 | TMOD2_MOUSE | QQLGTAVEMEIAQM[+16]LEENSR | 4.00 | 4.22E-04 | 0.0143 | 19 |
| Q9JKK7 | TMOD2_MOUSE | AKPVFEEPPNPTNVEASLQQM[+16]K | 4.78 | 6.52E-04 | 0.0143 | 19 |
| Q9JKK7 | TMOD2_MOUSE | FGYQFTK | 4.37 | 7.56E-04 | 0.0143 | 19 |
| Q9JKK7 | TMOD2_MOUSE | SNDPVALAFAEMLK | 5.14 | 8.22E-04 | 0.0143 | 19 |
| Q9JKK7 | TMOD2_MOUSE | ANDPSLQEVNLNNIK | 4.99 | 9.62E-04 | 0.0143 | 19 |
| Q9JKK7 | TMOD2_MOUSE | QLENVLDDLDPESATLPAGFR | 6.98 | 1.24E-03 | 0.0143 | 19 |
| Q9JKK7 | TMOD2_MOUSE | FSLAATR | 3.81 | 1.41E-03 | 0.0143 | 19 |
| Q9JKK7 | TMOD2_MOUSE | ENDTLTEIK | 4.15 | 1.44E-03 | 0.0143 | 19 |
| Q9JKK7 | TMOD2_MOUSE | DREDFVPFTGEK | 4.36 | 1.55E-03 | 0.0143 | 19 |
| Q9JKK7 | TMOD2_MOUSE | LSEEELK | 4.81 | 1.56E-03 | 0.0143 | 19 |
| Q9JKK7 | TMOD2_MOUSE | QQLGTAVEM[+16]EIAQMLEENSR | 7.39 | 1.71E-03 | 0.0143 | 19 |
| Q9JKK7 | TMOD2_MOUSE | AKPVFEEPPNPTNVEASLQQMK | 5.08 | 3.04E-03 | 0.0143 | 19 |
| Q9JKK7 | TMOD2_MOUSE | QQLGTAVEMEIAQMLEENSR | 4.56 | 4.20E-03 | 0.0149 | 19 |
| Q9JKK7 | TMOD2_MOUSE | SNDPVALAFAEM[+16]LK | 3.75 | 5.37E-03 | 0.0158 | 19 |
| Q9JKK7 | TMOD2_MOUSE | EHLLMYLEK | 5.75 | 6.01E-03 | 0.0164 | 19 |
| Q9JKK7 | TMOD2_MOUSE | FDEETTNGEGR | 4.28 | 7.65E-03 | 0.0181 | 19 |
| Q9JKK7 | TMOD2_MOUSE | LSEEELKQLENVLDDLDPESATLPAGFR | 8.99 | 8.45E-03 | 0.0188 | 19 |
| O08638 | MYH11_MOUSE | QLVSNLEK | 4.58 | 1.57E-03 | 0.0143 | 19 |
| P05213 | TBA1B_MOUSE | LISQIVSSITASLR | 3.07 | 2.82E-03 | 0.0143 | 19 |
| O08638 | MYH11_MOUSE | KATLQAEQLSNELATER | 4.48 | 3.05E-03 | 0.0143 | 19 |
| P05213 | TBA1B_MOUSE | AYHEQLSVAEITNAC[+57]FEPANQM[+16]VK | 2.42 | 3.29E-03 | 0.0143 | 19 |
| O08638 | MYH11_MOUSE | LQNEVESVTGMLNEAEGK | 3.59 | 3.30E-03 | 0.0143 | 19 |
| O08638 | MYH11_MOUSE | SM[+16]LQDREDQSILC[+57]TGESGAGK | 5.18 | 3.48E-03 | 0.0144 | 19 |
| P05213 | TBA1B_MOUSE | FDGALNVDLTEFQTNLVPYPR | 4.07 | 4.20E-03 | 0.0149 | 19 |
| P05213 | TBA1B_MOUSE | YM[+16]AC[+57]C[+57]LLYR | 1.39 | 4.21E-03 | 0.0149 | 19 |
| P05213 | TBA1B_MOUSE | QLFHPEQLITGK | 2.29 | 5.16E-03 | 0.0155 | 19 |
| P05213 | TBA1B_MOUSE | AVC[+57]MLSNTTAIAEAWAR | 2.86 | 5.17E-03 | 0.0155 | 19 |
| P05213 | TBA1B_MOUSE | TIGGGDDSFNTFFSETGAGK | 2.23 | 5.86E-03 | 0.0163 | 19 |
| O08638 | MYH11_MOUSE | KLEGDASDFHEQIADLQAQIAELK | 3.75 | 6.49E-03 | 0.0170 | 19 |
| O08638 | MYH11_MOUSE | VEDMAELTC[+57]LNEASVLHNLR | 4.46 | 6.57E-03 | 0.0171 | 19 |
| P05213 | TBA1B_MOUSE | EIIDLVLDR | 2.18 | 6.60E-03 | 0.0171 | 19 |
| P05213 | TBA1B_MOUSE | LDHKFDLMYAK | 2.28 | 6.69E-03 | 0.0172 | 19 |
| P05213 | TBA1B_MOUSE | AVC[+57]M[+16]LSNTTAIAEAWAR | 2.00 | 6.71E-03 | 0.0172 | 19 |
| P05213 | TBA1B_MOUSE | NLDIERPTYTNLNR | 2.65 | 6.94E-03 | 0.0174 | 19 |
| P05213 | TBA1B_MOUSE | VGINYQPPTVVPGGDLAK | 2.34 | 7.47E-03 | 0.0180 | 19 |
| P05213 | TBA1B_MOUSE | IHFPLATYAPVISAEK | 2.40 | 7.71E-03 | 0.0181 | 19 |
| O08638 | MYH11_MOUSE | IRELEGHISDLQEDLDSER | 3.81 | 7.72E-03 | 0.0181 | 19 |
| O08638 | MYH11_MOUSE | ELDEATESNEAMGR | 2.68 | 7.73E-03 | 0.0182 | 19 |
| O08638 | MYH11_MOUSE | DVASLGSQLQDTQELLQEETR | 4.62 | 8.97E-03 | 0.0195 | 19 |
| P05213 | TBA1B_MOUSE | AYHEQLSVAEITNAC[+57]FEPANQMVK | 3.10 | 9.28E-03 | 0.0199 | 19 |
| P05213 | TBA1B_MOUSE | QLFHPEQLITGKEDAANNYAR | 3.09 | 1.01E-02 | 0.0207 | 19 |
| O08638 | MYH11_MOUSE | VVSSVLQLGNIVFK | 5.44 | 1.09E-02 | 0.0216 | 19 |
| P05213 | TBA1B_MOUSE | LADQC[+57]TGLQGFLVFHSFGGGTGSGFTSLLM[+16]ER | 6.72 | 1.12E-02 | 0.0219 | 19 |
| O08638 | MYH11_MOUSE | LDAFLVLEQLR | 5.42 | 1.17E-02 | 0.0225 | 19 |
| P05213 | TBA1B_MOUSE | YMAC[+57]C[+57]LLYR | 2.34 | 1.19E-02 | 0.0227 | 19 |
| O08638 | MYH11_MOUSE | SMLQDREDQSILC[+57]TGESGAGK | 5.22 | 1.21E-02 | 0.0229 | 19 |
| P05213 | TBA1B_MOUSE | EDMAALEK | 1.53 | 1.22E-02 | 0.0230 | 19 |
| O08638 | MYH11_MOUSE | KLEVQLQDLQSK | 4.54 | 1.25E-02 | 0.0232 | 19 |

| Protein Accession | Protein Description | Modified Peptide Sequence | co-IP:Ctrl | p value | q value | Sig Peptide Count |
|---|---|---|---|---|---|---|
| Q8BYI9 | TENR_MOUSE | VGFGNLEDEFWLGLDNIHR | 4.04 | 1.74E-02 | 0.0285 | 19 |
| Q8BPN8 | DMXL2_MOUSE | FQLYNWLEK | 8.09 | 2.09E-02 | 0.0326 | 19 |
| Q62261 | SPTB2_MOUSE | EQWANLEQLSAIR | 3.90 | 2.16E-02 | 0.0335 | 19 |
| P62242 | RS8_MOUSE 4( | ELEFYLR | 1.65 | 2.17E-02 | 0.0336 | 19 |
| Q8C0M9 | ASGL1_MOUSE | GNLAYATSTGGIVNK | 2.81 | 2.23E-02 | 0.0343 | 19 |
| Q571F3 | Q571F3_MOUSE | A[+42]AELEYESVLC[+57]VKPDVSVYR | 1.11 | 2.55E-02 | 0.0377 | 19 |
| P57780 | ACTN4_MOUSE | LVSIGAEEIVDGNAK | 2.46 | 2.89E-02 | 0.0414 | 19 |
| P68033 | ACTC_MOUSE | EITALAPSTM[+16]K | 3.38 | 4.93E-04 | 0.0143 | 19 |
| P68033 | ACTC_MOUSE | DLTDYLM[+16]K | 3.39 | 1.27E-03 | 0.0143 | 19 |
| P68033 | ACTC_MOUSE | HQGVM[+16]VGMGQK | 3.40 | 2.66E-03 | 0.0143 | 19 |
| P68033 | ACTC_MOUSE | AVFPSIVGRPR | 4.29 | 2.75E-03 | 0.0143 | 19 |
| P68033 | ACTC_MOUSE | HQGVMVGM[+16]GQK | 3.24 | 2.84E-03 | 0.0143 | 19 |
| P68033 | ACTC_MOUSE | S[+42]YELPDGQVITIGNER | 4.25 | 2.98E-03 | 0.0143 | 19 |
| P68033 | ACTC_MOUSE | YPIEHGIITNWDDM[+16]EK | 3.91 | 3.09E-03 | 0.0143 | 19 |
| P68033 | ACTC_MOUSE | DIKEKLC[+57]YVALDFENEM[+16]ATAASSSSLEK | 10.56 | 3.35E-03 | 0.0143 | 19 |
| P68033 | ACTC_MOUSE | E[+42]ITALAPSTM[+16]K | 2.45 | 3.37E-03 | 0.0143 | 19 |
| P68033 | ACTC_MOUSE | IWHHTFYNELR | 5.33 | 4.33E-03 | 0.0150 | 19 |
| P68033 | ACTC_MOUSE | HQGVMVGMGQK | 4.02 | 4.64E-03 | 0.0153 | 19 |
| P68033 | ACTC_MOUSE | DSYVGDEAQSKR | 4.33 | 5.89E-03 | 0.0163 | 19 |
| P68033 | ACTC_MOUSE | IIAPPERK | 3.91 | 6.15E-03 | 0.0166 | 19 |
| P68033 | ACTC_MOUSE | DLTDYLMK | 4.31 | 6.79E-03 | 0.0172 | 19 |
| P68033 | ACTC_MOUSE | EITALAPSTMK | 3.64 | 7.28E-03 | 0.0177 | 19 |
| P68033 | ACTC_MOUSE | YPIEHGIITNWDDMEK | 5.06 | 8.40E-03 | 0.0188 | 19 |
| P68033 | ACTC_MOUSE | HQGVM[+16]VGM[+16]GQK | 3.38 | 1.24E-02 | 0.0231 | 19 |
| Q8CHC4 | SYNJ1_MOUSE | VSEQTLQSASSK | 3.39 | 2.28E-02 | 0.0348 | 19 |
| Q6ZQ38 | CAND1_MOUSE | C[+57]LDAVVSTR | 3.70 | 3.19E-02 | 0.0447 | 19 |
| P60710 | ACTB_MOUSE | QEYDESGPSIVHR | 3.57 | 8.73E-04 | 0.0143 | 18 |
| P60710 | ACTB_MOUSE | C[+57]PEALFQPSFLGM[+16]ESC[+57]GIHETTFNSIM[+16]K | 4.25 | 1.53E-03 | 0.0143 | 18 |
| P60710 | ACTB_MOUSE | KDLYANTVLSGGTTM[+16]YPGIADR | 4.17 | 1.72E-03 | 0.0143 | 18 |
| P60710 | ACTB_MOUSE | DLYANTVLSGGTTM[+16]YPGIADR | 4.11 | 1.76E-03 | 0.0143 | 18 |
| P60710 | ACTB_MOUSE | C[+57]PEALFQPSFLGMESC[+57]GIHETTFNSIM[+16]K | 5.32 | 1.77E-03 | 0.0143 | 18 |
| P60710 | ACTB_MOUSE | C[+57]PEALFQPSFLGM[+16]ESC[+57]GIHETTFNSIMK | 5.32 | 1.80E-03 | 0.0143 | 18 |
| P60710 | ACTB_MOUSE | TTGIVM[+16]DSGDGVTHTVPIYEGYALPHAILR | 5.89 | 2.01E-03 | 0.0143 | 18 |
| P60710 | ACTB_MOUSE | GYSFTTTAER | 3.61 | 2.04E-03 | 0.0143 | 18 |
| P60710 | ACTB_MOUSE | C[+57]DVDIR | 4.05 | 2.28E-03 | 0.0143 | 18 |
| P60710 | ACTB_MOUSE | EKLC[+57]YVALDFEQEMATAASSSSLEK | 4.51 | 2.52E-03 | 0.0143 | 18 |
| P60710 | ACTB_MOUSE | VAPEEHPVLLTEAPLNPK | 3.91 | 3.12E-03 | 0.0143 | 18 |
| P60710 | ACTB_MOUSE | C[+57]PEALFQPSFLGMESC[+57]GIHETTFNSIMK | 6.16 | 3.54E-03 | 0.0144 | 18 |
| P60710 | ACTB_MOUSE | EKLC[+57]YVALDFEQEM[+16]ATAASSSSLEK | 4.11 | 3.83E-03 | 0.0147 | 18 |
| P60710 | ACTB_MOUSE | G[+42]YSFTTTAER | 4.08 | 3.91E-03 | 0.0147 | 18 |
| P60710 | ACTB_MOUSE | KDLYANTVLSGGTTMYPGIADR | 3.14 | 4.73E-03 | 0.0153 | 18 |
| P60710 | ACTB_MOUSE | DLYANTVLSGGTTMYPGIADR | 5.17 | 5.44E-03 | 0.0158 | 18 |
| H3BJD0 | H3BJD0_MOUSE | NEPLEDAEANVVGSER | 9.94 | 1.99E-02 | 0.0314 | 18 |
| P48962 | ADT1_MOUSE | DFLAGGIAAAVSK | 6.07 | 2.86E-02 | 0.0410 | 18 |
| O08553 | DPYL2_MOUSE | FQLTDSQIYEVLSVIR | 3.53 | 5.46E-04 | 0.0143 | 18 |
| O08553 | DPYL2_MOUSE | DHGVNSFLVYMAFK | 3.29 | 6.09E-04 | 0.0143 | 18 |
| O08553 | DPYL2_MOUSE | DRFQLTDSQIYEVLSVIR | 4.99 | 8.90E-04 | 0.0143 | 18 |
| O08553 | DPYL2_MOUSE | AALAGGTTM[+16]IIDHVVPEPGTSLLAAFDQWR | 10.63 | 9.81E-04 | 0.0143 | 18 |
| O08553 | DPYL2_MOUSE | ILDLGITGPEGHVLSRPEEVEAEAVNR | 2.44 | 2.16E-03 | 0.0143 | 18 |
| O08553 | DPYL2_MOUSE | IVNDDQSFYADIYM[+16]EDGLIK | 1.35 | 3.27E-03 | 0.0143 | 18 |
| O08553 | DPYL2_MOUSE | ISVGSDADLVIWDPDSVK | 2.10 | 3.43E-03 | 0.0144 | 18 |
| O08553 | DPYL2_MOUSE | AALAGGTTMIIDHVVPEPGTSLLAAFDQWR | 9.57 | 3.64E-03 | 0.0146 | 18 |
| O08553 | DPYL2_MOUSE | DIGAIAQVHAENGDIIAEEQQR | 1.55 | 4.61E-03 | 0.0153 | 18 |
| O08553 | DPYL2_MOUSE | THNSALEYNIFEGMEC[+57]R | 2.06 | 5.58E-03 | 0.0160 | 18 |
| O08553 | DPYL2_MOUSE | IVNDDQSFYADIYMEDGLIK | 3.06 | 6.13E-03 | 0.0166 | 18 |
| O08553 | DPYL2_MOUSE | NLHQSGFSLSGAQIDDNIPR | 1.33 | 9.05E-03 | 0.0196 | 18 |
| O08553 | DPYL2_MOUSE | SITIANQTNC[+57]PLYVTK | 1.43 | 9.33E-03 | 0.0199 | 18 |
| O08553 | DPYL2_MOUSE | THNSALEYNIFEGM[+16]EC[+57]R | 1.17 | 9.74E-03 | 0.0204 | 18 |
| O08553 | DPYL2_MOUSE | GLYDGPVC[+57]EVSVTPK | 1.18 | 1.29E-02 | 0.0237 | 18 |
| O08553 | DPYL2_MOUSE | FQMPDQGM[+16]TSADDFFQGTK | 1.08 | 1.49E-02 | 0.0259 | 18 |
| P50518 | VATE1_MOUSE | LDLIAQQM[+16]M[+16]PEVR | 0.88 | 1.81E-02 | 0.0294 | 18 |
| P63330 | PP2AA_MOUSE | YSFLQFDPAPR | 1.05 | 2.84E-02 | 0.0408 | 18 |

| Protein Accession | Protein Description | Modified Peptide Sequence | co-IP:Ctrl | p value | q value | Sig Peptide Count |
|---|---|---|---|---|---|---|
| Q9D6F9 | TBB4A_MOUSE | EVDEQM[+16]LSVQSK | 1.88 | 1.22E-03 | 0.0143 | 17 |
| Q9D6F9 | TBB4A_MOUSE | M[+16]REIVHLQAGQC[+57]GNQIGAK | 2.53 | 1.29E-03 | 0.0143 | 17 |
| Q9D6F9 | TBB4A_MOUSE | GHYTEGAELVDAVLDVVR | 4.66 | 2.22E-03 | 0.0143 | 17 |
| Q9D6F9 | TBB4A_MOUSE | IM[+16]NTFSVVPSPK | 1.68 | 2.90E-03 | 0.0143 | 17 |
| Q9D6F9 | TBB4A_MOUSE | ALTVPELTQQM[+16]FDAK | 2.15 | 2.91E-03 | 0.0143 | 17 |
| Q9D6F9 | TBB4A_MOUSE | ALTVPELTQQMFDAK | 2.50 | 3.40E-03 | 0.0143 | 17 |
| Q9D6F9 | TBB4A_MOUSE | EIVHLQAGQC[+57]GNQIGAK | 2.28 | 4.11E-03 | 0.0148 | 17 |
| Q9D6F9 | TBB4A_MOUSE | MAATFIGNSTAIQELFK | 3.11 | 4.73E-03 | 0.0153 | 17 |
| Q9D6F9 | TBB4A_MOUSE | EVDEQMLSVQSK | 2.21 | 7.45E-03 | 0.0180 | 17 |
| Q9D6F9 | TBB4A_MOUSE | INVYYNEATGGNYVPR | 3.21 | 7.61E-03 | 0.0181 | 17 |
| Q9D6F9 | TBB4A_MOUSE | AVLVDLEPGTM[+16]DSVR | 1.87 | 7.91E-03 | 0.0183 | 17 |
| Q9D6F9 | TBB4A_MOUSE | MREIVHLQAGQC[+57]GNQIGAK | 3.06 | 9.60E-03 | 0.0203 | 17 |
| Q9D6F9 | TBB4A_MOUSE | IMNTFSVVPSPK | 2.38 | 9.89E-03 | 0.0205 | 17 |
| Q9D6F9 | TBB4A_MOUSE | AVLVDLEPGTMDSVR | 2.74 | 1.11E-02 | 0.0219 | 17 |
| Q9D6F9 | TBB4A_MOUSE | EAESC[+57]DC[+57]LQGFQLTHSLGGGTGSGMGTLLISK | 3.66 | 1.13E-02 | 0.0221 | 17 |
| Q9D6F9 | TBB4A_MOUSE | FWEVISDEHGIDPTGTYHGDSDLQLER | 6.20 | 1.25E-02 | 0.0232 | 17 |
| Q9D6F9 | TBB4A_MOUSE | YLTVAAVFR | 2.84 | 1.66E-02 | 0.0277 | 17 |
| P26039 | TLN1_MOUSE T | IPEALAGPPNDFGLFLSDDDPKK | 3.15 | 3.15E-04 | 0.0143 | 16 |
| P26039 | TLN1_MOUSE T | GVAALTSDPAVQAIVLDTASDVLDK | 6.26 | 8.73E-04 | 0.0143 | 16 |
| P26039 | TLN1_MOUSE T | TLAESALQLLYTAK | 4.08 | 1.65E-03 | 0.0143 | 16 |
| P26039 | TLN1_MOUSE T | VGDDPAVWQLK | 2.39 | 1.95E-03 | 0.0143 | 16 |
| P26039 | TLN1_MOUSE T | VAGSVTELIQAAEAMK | 4.26 | 2.26E-03 | 0.0143 | 16 |
| P26039 | TLN1_MOUSE T | ILAQATSDLVNAIK | 4.33 | 3.03E-03 | 0.0143 | 16 |
| P26039 | TLN1_MOUSE T | GVGAAATAVTQALNELLQHVK | 9.34 | 3.60E-03 | 0.0145 | 16 |
| P26039 | TLN1_MOUSE T | LLAALLEDEGGNGRPLLQAAK | 2.43 | 6.34E-03 | 0.0169 | 16 |
| P26039 | TLN1_MOUSE T | LAQAAQSSVATITR | 4.16 | 9.57E-03 | 0.0202 | 16 |
| P26039 | TLN1_MOUSE T | AVASAAAALVLK | 5.68 | 1.27E-02 | 0.0234 | 16 |
| P26039 | TLN1_MOUSE T | LNEAAAGLNQAATELVQASR | 3.22 | 1.60E-02 | 0.0271 | 16 |
| P26039 | TLN1_MOUSE T | GSQAQPDSPSAQLALIAASQSFLQPGGK | 6.17 | 1.68E-02 | 0.0279 | 16 |
| Q99104 | MYO5A_MOUSE | NSLHLLMETLNATTPHYVR | 2.63 | 1.72E-02 | 0.0284 | 16 |
| Q62261 | SPTB2_MOUSE | LAEISDVWEEMK | 5.87 | 2.07E-02 | 0.0324 | 16 |
| P60229 | EIF3E_MOUSE | HLVFPLLEFLSVK | 4.63 | 2.20E-02 | 0.0340 | 16 |
| Q9CXY6 | ILF2_MOUSE In | NQDLAPNSAEQASILSLVTK | 3.51 | 3.28E-02 | 0.0458 | 16 |
| P47757 | CAPZB_MOUSE | SGSGTM[+16]NLGGSLTR | 3.45 | 1.13E-03 | 0.0143 | 15 |
| P47757 | CAPZB_MOUSE | STLNEIYFGK | 3.98 | 1.38E-03 | 0.0143 | 15 |
| P47757 | CAPZB_MOUSE | DYLLC[+57]DYNR | 3.90 | 1.60E-03 | 0.0143 | 15 |
| P47757 | CAPZB_MOUSE | LEVEANNAFDQYR | 4.57 | 2.45E-03 | 0.0143 | 15 |
| P47757 | CAPZB_MOUSE | SPWSNKYDPPLEDGAMPSAR | 5.16 | 2.62E-03 | 0.0143 | 15 |
| P47757 | CAPZB_MOUSE | RLPPQQIEK | 4.39 | 2.85E-03 | 0.0143 | 15 |
| P47757 | CAPZB_MOUSE | KLEVEANNAFDQYR | 4.35 | 3.35E-03 | 0.0143 | 15 |
| P47757 | CAPZB_MOUSE | SGSGTMNLGGSLTR | 4.20 | 3.57E-03 | 0.0145 | 15 |
| P47757 | CAPZB_MOUSE | LTSTVM[+16]LWLQTNK | 6.15 | 3.94E-03 | 0.0147 | 15 |
| P47757 | CAPZB_MOUSE | SPWSNKYDPPLEDGAM[+16]PSAR | 4.92 | 4.08E-03 | 0.0148 | 15 |
| P47757 | CAPZB_MOUSE | GC[+57]WDSIHVVEVQEK | 4.86 | 6.15E-03 | 0.0166 | 15 |
| P47757 | CAPZB_MOUSE | LVEDM[+16]ENK | 3.73 | 9.29E-03 | 0.0199 | 15 |
| P47757 | CAPZB_MOUSE | YDPPLEDGAMPSAR | 4.63 | 1.12E-02 | 0.0219 | 15 |
| P47757 | CAPZB_MOUSE | LTSTVMLWLQTNK | 6.84 | 1.52E-02 | 0.0263 | 15 |
| Q8C0C7 | SYFA_MOUSE I | SLQALGEVIEAELR | 4.87 | 2.32E-02 | 0.0352 | 15 |
| P52480 | KPYM_MOUSE | LNFSHGTHEYHAETIK | 2.27 | 8.78E-04 | 0.0143 | 15 |
| P52480 | KPYM_MOUSE | FGVEQDVDM[+16]VFASFIR | 3.04 | 1.62E-03 | 0.0143 | 15 |
| P52480 | KPYM_MOUSE | RFDEILEASDGIMVAR | 2.17 | 4.88E-03 | 0.0155 | 15 |
| P52480 | KPYM_MOUSE | AEGSDVANAVLDGADC[+57]IM[+16]LSGETAK | 1.18 | 4.94E-03 | 0.0155 | 15 |
| P52480 | KPYM_MOUSE | DAVLNAWAEDVDLR | 1.47 | 5.43E-03 | 0.0158 | 15 |
| P52480 | KPYM_MOUSE | RFDEILEASDGIM[+16]VAR | 1.63 | 6.30E-03 | 0.0168 | 15 |
| P52480 | KPYM_MOUSE | FDEILEASDGIM[+16]VAR | 2.75 | 6.32E-03 | 0.0169 | 15 |
| P01869 | IGH1M_MOUSE | APQVYTIPPPKEQM[+16]AK | 2.09 | 1.84E-02 | 0.0297 | 15 |
| Q8VDD5 | MYH9_MOUSE | VEEEAAQK | 0.73 | 2.02E-02 | 0.0318 | 15 |
| Q91WK2 | EIF3H_MOUSE | TAQGSLSLK | 5.38 | 2.06E-02 | 0.0323 | 15 |
| P61979 | HNRPK_MOUSE | IDEPLEGSEDR | 1.88 | 2.65E-02 | 0.0388 | 15 |
| Q8C8N2 | SCAI_MOUSE F | TSETSYLNEAFSFYSAIR | 1.63 | 2.90E-02 | 0.0414 | 15 |
| Q61171 | PRDX2_MOUSE | EGGLGPLNIPLLADVTK | 1.17 | 3.01E-02 | 0.0427 | 15 |
| P26039 | TLN1_MOUSE T | NLGTALAELR | 2.13 | 3.29E-02 | 0.0460 | 15 |

| Protein Accession | Protein Description | Modified Peptide Sequence | co-IP:Ctrl | p value | q value | Sig Peptide Count |
|---|---|---|---|---|---|---|
| Q8CI94 | PYGB_MOUSE | WLLLC[+57]NPGLAEIIVER | 1.20 | 3.56E-02 | 0.0492 | 15 |
| Q8BYI9 | TENR_MOUSE | ITFTPSSGISSEVTVPR | 2.16 | 9.87E-05 | 0.0143 | 14 |
| Q8BYI9 | TENR_MOUSE | DKEEDMLEVLLDATKR | 3.66 | 1.47E-03 | 0.0143 | 14 |
| Q8BYI9 | TENR_MOUSE | LEGLSENTDYTVLLQAAQEATR | 3.00 | 6.40E-03 | 0.0169 | 14 |
| Q8BYI9 | TENR_MOUSE | AAIENYVLTYK | 2.92 | 7.23E-03 | 0.0177 | 14 |
| Q8BYI9 | TENR_MOUSE | DKEEDMLEVLLDATK | 4.11 | 8.44E-03 | 0.0188 | 14 |
| Q8BYI9 | TENR_MOUSE | SPPTSASVSTVIDGPTQILVR | 2.85 | 8.61E-03 | 0.0190 | 14 |
| Q8BYI9 | TENR_MOUSE | LYPATEYEISLNSVR | 2.47 | 9.88E-03 | 0.0205 | 14 |
| Q8BYI9 | TENR_MOUSE | YEVSISAVR | 2.15 | 1.22E-02 | 0.0230 | 14 |
| Q8BYI9 | TENR_MOUSE | DVSDTVAFVEWTPPR | 2.79 | 1.22E-02 | 0.0230 | 14 |
| Q8BYI9 | TENR_MOUSE | SSLTSTVFTTGGR | 2.06 | 1.42E-02 | 0.0251 | 14 |
| Q8BYI9 | TENR_MOUSE | YGLVGGEGGK | 1.73 | 1.51E-02 | 0.0262 | 14 |
| D0U286 | D0U286_MUSCE E | LLGNVIM[+16]IVLGHHLGK | 8.73 | 1.73E-02 | 0.0284 | 14 |
| Q99104 | MYO5A_MOUSE | QQQLLAQNLQLPPEAR | 2.64 | 1.82E-02 | 0.0295 | 14 |
| Q8C0M9 | ASGL1_MOUSE | LQAGIDLC[+57]ETR | 3.00 | 3.36E-02 | 0.0468 | 14 |
| P47857 | PFKAM_MOUSE | SEWSDLLNDLQK | 2.78 | 2.16E-04 | 0.0143 | 14 |
| P47857 | PFKAM_MOUSE | GITNLC[+57]VIGGDGSLTGADTFR | 2.98 | 3.29E-04 | 0.0143 | 14 |
| P47857 | PFKAM_MOUSE | LPLM[+16]EC[+57]VQVTK | 1.61 | 2.30E-03 | 0.0143 | 14 |
| P47857 | PFKAM_MOUSE | VLVVHDGFEGLAK | 2.04 | 2.88E-03 | 0.0143 | 14 |
| P47857 | PFKAM_MOUSE | LNIIIVAEGAIDK | 4.51 | 3.99E-03 | 0.0148 | 14 |
| P47857 | PFKAM_MOUSE | MGVEAVMALLEGTPDTPAC[+57]VVSLSGNQAVR | 5.60 | 5.54E-03 | 0.0159 | 14 |
| P47857 | PFKAM_MOUSE | NLEQISANITK | 1.65 | 5.74E-03 | 0.0161 | 14 |
| P47857 | PFKAM_MOUSE | DLQVNVEHLVQK | 2.30 | 6.28E-03 | 0.0168 | 14 |
| P47857 | PFKAM_MOUSE | ALVFQPVTELK | 1.83 | 8.07E-03 | 0.0184 | 14 |
| P47857 | PFKAM_MOUSE | VFFVHEGYQGLVDGGEHIR | 1.60 | 9.85E-03 | 0.0205 | 14 |
| P47857 | PFKAM_MOUSE | VGIFTGAR | 1.63 | 1.18E-02 | 0.0226 | 14 |
| P47857 | PFKAM_MOUSE | IGLIQGNR | 1.65 | 1.19E-02 | 0.0227 | 14 |
| P47857 | PFKAM_MOUSE | C[+57]NENYTTDFIFNLYSEEGK | 6.13 | 1.19E-02 | 0.0227 | 14 |
| P47857 | PFKAM_MOUSE | TFVLEVMGR | 2.60 | 1.34E-02 | 0.0242 | 14 |
| Q6A087 | Q6A087_MOUSE | SLDDFQAWLGR | 2.51 | 8.52E-04 | 0.0143 | 13 |
| Q6A087 | Q6A087_MOUSE | VIESTQGLGNDLAGVLALQR | 3.64 | 2.34E-03 | 0.0143 | 13 |
| Q6A087 | Q6A087_MOUSE | DVEDEILWVTER | 5.64 | 3.01E-03 | 0.0143 | 13 |
| Q6A087 | Q6A087_MOUSE | LVSQDNFGLELAAVEAAVR | 4.52 | 3.47E-03 | 0.0144 | 13 |
| Q6A087 | Q6A087_MOUSE | IIGTQEQLNQR | 2.51 | 5.18E-03 | 0.0156 | 13 |
| Q6A087 | Q6A087_MOUSE | KHEAIETDIVAYSGR | 4.36 | 5.33E-03 | 0.0158 | 13 |
| Q6A087 | Q6A087_MOUSE | TLGTAAAGPELAELQEMWK | 5.04 | 9.49E-03 | 0.0202 | 13 |
| Q6A087 | Q6A087_MOUSE | VTAVNDIAEQLLK | 4.38 | 1.09E-02 | 0.0216 | 13 |
| Q6A087 | Q6A087_MOUSE | LTLEQGQQLVAEGHPGANQASTR | 3.20 | 1.13E-02 | 0.0221 | 13 |
| Q6A087 | Q6A087_MOUSE | LLLNLELQK | 4.44 | 1.24E-02 | 0.0231 | 13 |
| Q6A087 | Q6A087_MOUSE | TQTAVASEEGPATLPEAEALLAQHAALR | 4.08 | 1.31E-02 | 0.0239 | 13 |
| Q6A087 | Q6A087_MOUSE | VGELTQEANALAAGHPAQAPAINTR | 3.62 | 1.45E-02 | 0.0255 | 13 |
| P27323 | HS901_ARATH He | SLTNDWEDHLAVK | 7.14 | 3.05E-02 | 0.0431 | 13 |
| Q8C8N2 | SCAI_MOUSE F | NFTNLFGQPLVC[+57]LLSPTAYPK | 8.81 | 2.28E-03 | 0.0143 | 13 |
| Q8C8N2 | SCAI_MOUSE F | DIAQLLTHSR | 4.10 | 3.14E-03 | 0.0143 | 13 |
| Q8C8N2 | SCAI_MOUSE F | FSELTVDMFR | 3.80 | 3.29E-03 | 0.0143 | 13 |
| Q8C8N2 | SCAI_MOUSE F | TVTDFC[+57]YLLDK | 5.14 | 3.46E-03 | 0.0144 | 13 |
| Q8C8N2 | SCAI_MOUSE F | FSELTVDM[+16]FR | 2.12 | 3.78E-03 | 0.0147 | 13 |
| Q8C8N2 | SCAI_MOUSE F | KPLFIVVDSSNSVAYK | 4.17 | 5.88E-03 | 0.0163 | 13 |
| Q8C8N2 | SCAI_MOUSE F | SDGEGPYDFGGVLTNSNR | 7.28 | 8.57E-03 | 0.0190 | 13 |
| Q8C8N2 | SCAI_MOUSE F | SYYSQVNKEDRPELVVK | 7.68 | 1.04E-02 | 0.0211 | 13 |
| Q8C8N2 | SCAI_MOUSE F | QWQSYFGR | 3.73 | 1.36E-02 | 0.0244 | 13 |
| Q8C8N2 | SCAI_MOUSE F | FIVVC[+57]LLLNK | 9.14 | 1.38E-02 | 0.0247 | 13 |
| Q8C8N2 | SCAI_MOUSE F | IGQLYYHYYLR | 3.45 | 1.55E-02 | 0.0266 | 13 |
| Q8C8N2 | SCAI_MOUSE F | HILELASILDVR | 9.08 | 1.68E-02 | 0.0279 | 13 |
| Q8BH44 | COR2B_MOUSE | HGLDVSAC[+57]EVFR | 5.19 | 2.90E-02 | 0.0414 | 13 |
| O35643 | AP1B1_MOUSE | LASQANIAQVLAELK | 1.85 | 1.73E-03 | 0.0143 | 13 |
| O35643 | AP1B1_MOUSE | LGAPISSGLSDLFDLTSGVGTLSGSYVAPK | 3.62 | 2.44E-03 | 0.0143 | 13 |
| O35643 | AP1B1_MOUSE | KPTETQELVQQVLSLATQDSDNPDLR | 4.24 | 3.83E-03 | 0.0147 | 13 |
| O35643 | AP1B1_MOUSE | LAPPLVTLLSAEPELQYVALR | 2.79 | 4.09E-03 | 0.0148 | 13 |
| O35643 | AP1B1_MOUSE | LVYLYLMNYAK | 2.67 | 4.63E-03 | 0.0153 | 13 |
| O35643 | AP1B1_MOUSE | M[+16]EPLNNLQVAVK | 4.25 | 4.91E-03 | 0.0155 | 13 |
| O35643 | AP1B1_MOUSE | C[+57]VSTLLDLIQTK | 2.30 | 5.87E-03 | 0.0163 | 13 |

| Protein Accession | Protein Description | Modified Peptide Sequence | co-IP:Ctrl | p value | q value | Sig Peptide Count |
|---|---|---|---|---|---|---|
| O35643 | AP1B1_MOUSE | EYATEVDVDFVR | 3.06 | 1.23E-02 | 0.0231 | 13 |
| O35643 | AP1B1_MOUSE | AAMIWIVGEYAER | 3.12 | 1.27E-02 | 0.0234 | 13 |
| O35643 | AP1B1_MOUSE | LVYLYLM[+16]NYAK | 2.00 | 1.38E-02 | 0.0247 | 13 |
| O35643 | AP1B1_MOUSE | LSHANSAVVLSAVK | 1.44 | 1.52E-02 | 0.0263 | 13 |
| P46460 | NSF_MOUSE V | FSNLVLQALLVLLKK | 1.58 | 2.40E-02 | 0.0361 | 13 |
| Q99JY9 | ARP3_MOUSE / | DITYFIQQLLR | 1.36 | 2.89E-02 | 0.0414 | 13 |
| Q9JJ28 | FLII_MOUSE Pr | LTSLEEFMAANNNLELIPESLC[+57]R | 7.95 | 2.14E-03 | 0.0143 | 13 |
| Q9JJ28 | FLII_MOUSE Pr | TQSNLPTSLEGLSNLSDVDLSC[+57]NDLTR | 5.99 | 3.06E-03 | 0.0143 | 13 |
| Q9JJ28 | FLII_MOUSE Pr | M[+42]EATGVLPFVR | 5.16 | 3.45E-03 | 0.0144 | 13 |
| Q9JJ28 | FLII_MOUSE Pr | VGLGLGYLELPQINYK | 5.88 | 4.43E-03 | 0.0151 | 13 |
| Q9JJ28 | FLII_MOUSE Pr | LLQSLLDTR | 5.32 | 4.67E-03 | 0.0153 | 13 |
| Q9JJ28 | FLII_MOUSE Pr | ADLTALFLPR | 5.74 | 5.26E-03 | 0.0157 | 13 |
| Q9JJ28 | FLII_MOUSE Pr | LAEDILNTMFDASYSK | 5.96 | 5.88E-03 | 0.0163 | 13 |
| Q9JJ28 | FLII_MOUSE Pr | NAEAVLQGQGLSGK | 5.64 | 7.36E-03 | 0.0178 | 13 |
| Q9JJ28 | FLII_MOUSE Pr | TGLC[+57]YLPEELAALQK | 3.16 | 7.74E-03 | 0.0182 | 13 |
| Q9JJ28 | FLII_MOUSE Pr | NQLTSLPSAIC[+57]K | 2.98 | 7.97E-03 | 0.0183 | 13 |
| Q9JJ28 | FLII_MOUSE Pr | LDFDGLPSGIGK | 3.36 | 8.49E-03 | 0.0189 | 13 |
| Q9JJ28 | FLII_MOUSE Pr | VPEC[+57]LYTLPSLR | 5.48 | 8.88E-03 | 0.0194 | 13 |
| Q9JJ28 | FLII_MOUSE Pr | LAGASPATVAAAAAVGSGSKDPLAF | 6.09 | 9.42E-03 | 0.0201 | 13 |
| P39053 | DYN1_MOUSE I | RIEGSGDQIDTYELSGGAR | 4.20 | 1.61E-03 | 0.0143 | 13 |
| P39053 | DYN1_MOUSE I | ALLQMVQQFAVDFEK | 5.88 | 4.56E-03 | 0.0152 | 13 |
| P39053 | DYN1_MOUSE I | IEGSGDQIDTYELSGGAR | 1.36 | 4.78E-03 | 0.0154 | 13 |
| P39053 | DYN1_MOUSE I | HIFALFNTEQR | 2.07 | 5.75E-03 | 0.0161 | 13 |
| P39053 | DYN1_MOUSE I | QLELAC[+57]ETQEEVDSWK | 2.59 | 8.34E-03 | 0.0188 | 13 |
| P39053 | DYN1_MOUSE I | C[+57]VDMVISELISTVR | 3.91 | 1.07E-02 | 0.0214 | 13 |
| O08638 | MYH11_MOUSE | QADLEKEELAEELASSLSGR | 2.58 | 2.16E-02 | 0.0335 | 13 |
| A0A0B6VMB2 | A0A0B6VMB2_MC | NTQPIM[+16]DTDGSYFVYSK | 2.00 | 2.61E-02 | 0.0384 | 13 |
| P35980 | RL18_MOUSE 6 | TAVVVGTVTDDVR | 1.24 | 2.64E-02 | 0.0387 | 13 |
| Q5SYD0 | MYO1D_MOUSE | HQVEYLGLLENVR | 0.71 | 2.72E-02 | 0.0395 | 13 |
| Q76MZ3 | 2AAA_MOUSE S | VLAM[+16]SGDPNYLHR | 2.13 | 3.07E-02 | 0.0434 | 13 |
| Q9JMH9 | MY18A_MOUSE | TEEQIAAEEAWYETEK | 1.05 | 3.23E-02 | 0.0452 | 13 |
| P48036 | ANXA5_MOUSE | TPEELSAIK | 1.68 | 3.39E-02 | 0.0472 | 13 |
| Q8VDN2 | AT1A1_MOUSE | QAADM[+16]ILLDDNFASIVTGVEEGR | 2.72 | 5.34E-04 | 0.0143 | 12 |
| Q8VDN2 | AT1A1_MOUSE | SPDFTNENPLETR | 2.11 | 7.34E-04 | 0.0143 | 12 |
| Q8VDN2 | AT1A1_MOUSE | GVGIISEGNETVEDIAAR | 3.62 | 2.59E-03 | 0.0143 | 12 |
| Q8VDN2 | AT1A1_MOUSE | DGPNALTPPPTTPEWVK | 2.87 | 4.28E-03 | 0.0150 | 12 |
| Q8VDN2 | AT1A1_MOUSE | AVAGDASESALLK | 3.61 | 4.30E-03 | 0.0150 | 12 |
| Q8VDN2 | AT1A1_MOUSE | QGAIVAVTGDGVNDSPALKK | 2.01 | 5.07E-03 | 0.0155 | 12 |
| Q8VDN2 | AT1A1_MOUSE | LNIPVNQVNPR | 1.63 | 5.95E-03 | 0.0163 | 12 |
| Q8VDN2 | AT1A1_MOUSE | QAADMILLDDNFASIVTGVEEGR | 4.00 | 6.75E-03 | 0.0172 | 12 |
| Q8VDN2 | AT1A1_MOUSE | NM[+16]VPQQALVIR | 1.93 | 7.26E-03 | 0.0177 | 12 |
| Q8VDN2 | AT1A1_MOUSE | QGAIVAVTGDGVNDSPALK | 2.45 | 7.84E-03 | 0.0182 | 12 |
| Q8VDN2 | AT1A1_MOUSE | VDNSSLTGESEPQTR | 1.37 | 8.05E-03 | 0.0184 | 12 |
| Q8VDN2 | AT1A1_MOUSE | NLEAVETLGSTSTIC[+57]SDK | 2.73 | 1.38E-02 | 0.0247 | 12 |
| Q60875 | ARHG2_MOUSE | LGDLLISQFSGSNAEQMR | 3.48 | 1.52E-03 | 0.0143 | 12 |
| Q60875 | ARHG2_MOUSE | AAVASVTPEK | 1.73 | 2.94E-03 | 0.0143 | 12 |
| Q60875 | ARHG2_MOUSE | NNTALQSVSLR | 3.68 | 3.18E-03 | 0.0143 | 12 |
| Q60875 | ARHG2_MOUSE | SVSTTNIAGHFNDESPLGLR | 3.78 | 3.74E-03 | 0.0147 | 12 |
| Q60875 | ARHG2_MOUSE | YIFTSLDKPSVVSLQNLIVR | 8.54 | 4.11E-03 | 0.0148 | 12 |
| Q60875 | ARHG2_MOUSE | LQEIYNR | 2.58 | 6.03E-03 | 0.0164 | 12 |
| Q60875 | ARHG2_MOUSE | ILQNSHGVEEEYQDLASALGLVK | 6.82 | 7.42E-03 | 0.0179 | 12 |
| Q60875 | ARHG2_MOUSE | FLNQLLER | 6.49 | 9.06E-03 | 0.0196 | 12 |
| Q60875 | ARHG2_MOUSE | SLPAGDALYLSFNPPQPSR | 3.18 | 1.00E-02 | 0.0206 | 12 |
| Q60875 | ARHG2_MOUSE | LESFESLR | 2.24 | 1.60E-02 | 0.0271 | 12 |
| P02088 | HBB1_MOUSE I | VITAFNDGLNHLDSLK | 3.88 | 2.11E-02 | 0.0329 | 12 |
| Q61656 | DDX5_MOUSE I | EANQAINPK | 3.25 | 2.46E-02 | 0.0367 | 12 |
| Q8CI94 | PYGB_MOUSE | VIPAADLSQQISTAGTEASGTGNM[+16]K | 1.74 | 1.23E-04 | 0.0143 | 12 |
| Q8CI94 | PYGB_MOUSE | LKQEYFVVAATLQDIIR | 4.55 | 3.90E-03 | 0.0147 | 12 |
| Q8CI94 | PYGB_MOUSE | LKDFNVGDYIEAVLDR | 4.45 | 4.40E-03 | 0.0150 | 12 |
| Q8CI94 | PYGB_MOUSE | IGEGFLTDLSQLKK | 2.04 | 4.45E-03 | 0.0151 | 12 |
| Q8CI94 | PYGB_MOUSE | LLSLVDDEAFIR | 1.79 | 7.44E-03 | 0.0179 | 12 |
| Q8CI94 | PYGB_MOUSE | GIAGLGDVAEVR | 1.53 | 7.57E-03 | 0.0180 | 12 |

| Protein Accession | Protein Description | Modified Peptide Sequence | co-IP:Ctrl | p value | q value | Sig Peptide Count |
|---|---|---|---|---|---|---|
| Q8CI94 | PYGB_MOUSE | VAIQLNDTHPALSIPELMR | 2.07 | 7.76E-03 | 0.0182 | 12 |
| Q8CI94 | PYGB_MOUSE | VLYPNDNFFEGK | 1.40 | 1.02E-02 | 0.0208 | 12 |
| Q8CI94 | PYGB_MOUSE | VIFLENYR | 1.48 | 1.24E-02 | 0.0231 | 12 |
| Q8CI94 | PYGB_MOUSE | LAAC[+57]FLDSMATLGLAAYGYGIR | 8.24 | 1.61E-02 | 0.0272 | 12 |
| P07901 | HS90A_MOUSE | TLTIVDTGIGMTK | 1.24 | 2.43E-02 | 0.0364 | 12 |
| P51410 | RL9_MOUSE 60 | TGVAC[+57]SVSQAQK | 1.86 | 3.56E-02 | 0.0492 | 12 |
| P97427 | DPYL1_MOUSE | NLHQSNFSLSGAQIDDNNPR | 1.37 | 2.95E-03 | 0.0143 | 12 |
| P97427 | DPYL1_MOUSE | IAVGSDADVVIWDPDK | 3.53 | 3.31E-03 | 0.0143 | 12 |
| P97427 | DPYL1_MOUSE | STVEYNIFEGMEC[+57]HGSPLVVISQGK | 3.03 | 4.68E-03 | 0.0153 | 12 |
| P97427 | DPYL1_MOUSE | GM[+16]YDGPVYEVPATPK | 2.28 | 7.90E-03 | 0.0183 | 12 |
| P97427 | DPYL1_MOUSE | GLGAVILVHAENGDLIAQEQK | 1.76 | 8.37E-03 | 0.0188 | 12 |
| P97427 | DPYL1_MOUSE | IINDDQSFYADVYLEDGLIK | 6.55 | 9.52E-03 | 0.0202 | 12 |
| P97427 | DPYL1_MOUSE | DNFTLIPEGVNGIEER | 1.50 | 1.01E-02 | 0.0207 | 12 |
| P97427 | DPYL1_MOUSE | DLYQMSDSQLYEAFTFLK | 5.43 | 1.23E-02 | 0.0231 | 12 |
| P47963 | RL13_MOUSE 6 | GFSLEELR | 1.07 | 1.82E-02 | 0.0295 | 12 |
| Q91V92 | ACLY_MOUSE / | TIAIIAEGIPEALTR | 1.37 | 1.87E-02 | 0.0300 | 12 |
| Q6ZQ38 | CAND1_MOUSE | LGTLSALDILIK | 3.57 | 3.11E-02 | 0.0438 | 12 |
| H3BJD0 | H3BJD0_MOUSE | LYDSVSSTDGEDSLER | 0.96 | 3.13E-02 | 0.0440 | 12 |
| Q76MZ3 | 2AAA_MOUSE S | QLSQSLLPAIVELAEDAK | 3.29 | 1.73E-04 | 0.0143 | 11 |
| P58252 | EF2_MOUSE El | ALLELQLEPEELYQTFQR | 4.30 | 2.69E-04 | 0.0143 | 11 |
| Q76MZ3 | 2AAA_MOUSE S | SALASVIM[+16]GLSPILGK | 3.70 | 3.70E-04 | 0.0143 | 11 |
| P58252 | EF2_MOUSE El | EGIPALDNFLDKL | 3.02 | 3.71E-04 | 0.0143 | 11 |
| Q76MZ3 | 2AAA_MOUSE S | SALASVIMGLSPILGK | 2.92 | 4.18E-04 | 0.0143 | 11 |
| P58252 | EF2_MOUSE El | VFDAIMNFR | 3.43 | 9.43E-04 | 0.0143 | 11 |
| Q76MZ3 | 2AAA_MOUSE S | EFC[+57]ENLSADC[+57]R | 2.15 | 1.14E-03 | 0.0143 | 11 |
| P58252 | EF2_MOUSE El | TFC[+57]QLILDPIFK | 2.57 | 1.73E-03 | 0.0143 | 11 |
| Q76MZ3 | 2AAA_MOUSE S | LNIISNLDC[+57]VNEVIGIR | 2.93 | 2.16E-03 | 0.0143 | 11 |
| P58252 | EF2_MOUSE El | YVEPIEDVPC[+57]GNIVGLVGVDQFLVK | 3.23 | 3.16E-03 | 0.0143 | 11 |
| P13020 | GELS_MOUSE ( | SEDC[+57]FILDHGR | 4.30 | 3.32E-03 | 0.0143 | 11 |
| Q76MZ3 | 2AAA_MOUSE S | DNTIEHLLPLFLAQLK | 7.56 | 3.90E-03 | 0.0147 | 11 |
| P13020 | GELS_MOUSE ( | HVVPNEVVVQR | 5.25 | 4.35E-03 | 0.0150 | 11 |
| P58252 | EF2_MOUSE El | WLPAGDALLQMITIHLPSPVTAQK | 9.27 | 4.48E-03 | 0.0151 | 11 |
| P13020 | GELS_MOUSE ( | DSQEEEKTEALTSAK | 3.92 | 4.59E-03 | 0.0152 | 11 |
| Q76MZ3 | 2AAA_MOUSE S | YFAQEALTVLSLA | 2.31 | 4.68E-03 | 0.0153 | 11 |
| P58252 | EF2_MOUSE El | C[+57]LYASVLTAQPR | 1.76 | 6.00E-03 | 0.0164 | 11 |
| P13020 | GELS_MOUSE ( | EVQGFESSTFSGYFK | 4.97 | 6.55E-03 | 0.0171 | 11 |
| Q76MZ3 | 2AAA_MOUSE S | DNTIEHLLPLFLAQLKDEC[+57]PEVR | 7.20 | 6.96E-03 | 0.0174 | 11 |
| P13020 | GELS_MOUSE ( | YIETDPANR | 4.03 | 7.03E-03 | 0.0175 | 11 |
| P13020 | GELS_MOUSE ( | SGALNSNDAFVLK | 3.23 | 8.00E-03 | 0.0183 | 11 |
| Q76MZ3 | 2AAA_MOUSE S | LSTIALALGVER | 2.46 | 8.03E-03 | 0.0184 | 11 |
| P13020 | GELS_MOUSE ( | VSNGAGSMSVSLVADENPFAQGALR | 6.14 | 8.32E-03 | 0.0187 | 11 |
| P13020 | GELS_MOUSE ( | QTQVSVLPEGGETPLFK | 4.94 | 8.64E-03 | 0.0191 | 11 |
| P58252 | EF2_MOUSE El | ETVSEESNVLC[+57]LSK | 3.98 | 1.10E-02 | 0.0217 | 11 |
| Q76MZ3 | 2AAA_MOUSE S | ENVIMTQILPC[+57]IK | 2.67 | 1.26E-02 | 0.0233 | 11 |
| P13020 | GELS_MOUSE ( | VPVDPATYGQFYGGDSYIILYNYR | 4.49 | 1.48E-02 | 0.0258 | 11 |
| P58252 | EF2_MOUSE El | STLTDSLVC[+57]K | 2.88 | 1.54E-02 | 0.0265 | 11 |
| P58252 | EF2_MOUSE El | STAISLFYELSENDLNFIK | 6.53 | 1.59E-02 | 0.0270 | 11 |
| P13020 | GELS_MOUSE ( | DGGQTAPASIR | 3.49 | 1.59E-02 | 0.0270 | 11 |
| Q8CGF6 | WDR47_MOUSE | LIHDTANIHTSTPR | 3.26 | 1.81E-02 | 0.0294 | 11 |
| O70194 | EIF3D_MOUSE | LGDDIDLIVR | 2.04 | 2.25E-02 | 0.0345 | 11 |
| P48453 | PP2BB_MOUSE | YENNVMNIR | 4.53 | 3.06E-02 | 0.0432 | 11 |
| P16858 | G3P_MOUSE G | VIHDNFGIVEGLM[+16]TTVHAITATQK | 2.56 | 2.09E-04 | 0.0143 | 11 |
| P16858 | G3P_MOUSE G | VIHDNFGIVEGLMTTVHAITATQK | 5.88 | 1.77E-03 | 0.0143 | 11 |
| P16858 | G3P_MOUSE G | WGEAGAEYVVESTGVFTTMEK | 1.87 | 2.65E-03 | 0.0143 | 11 |
| P16858 | G3P_MOUSE G | LVINGKPITIFQER | 1.69 | 4.81E-03 | 0.0154 | 11 |
| P16858 | G3P_MOUSE G | LISWYDNEYGYSNR | 2.03 | 5.36E-03 | 0.0158 | 11 |
| P16858 | G3P_MOUSE G | RVIISAPSADAPMFVMGVNHEK | 2.53 | 6.25E-03 | 0.0168 | 11 |
| P16858 | G3P_MOUSE G | RVIISAPSADAPMFVM[+16]GVNHEK | 1.53 | 1.42E-02 | 0.0251 | 11 |
| P16858 | G3P_MOUSE G | RVIISAPSADAPM[+16]FVMGVNHEK | 1.50 | 1.60E-02 | 0.0271 | 11 |
| P97427 | DPYL1_MOUSE | IFNLYPR | 1.59 | 1.87E-02 | 0.0300 | 11 |
| P17426 | AP2A1_MOUSE | EM[+16]GEAFAADIPR | 1.41 | 2.01E-02 | 0.0317 | 11 |
| P63101 | 1433Z_MOUSE | DNLTLWTSDTQGDEAEAGEGGEN | 1.02 | 2.97E-02 | 0.0422 | 11 |

| Protein Accession | Protein Description | Modified Peptide Sequence | co-IP:Ctrl | p value | q value | Sig Peptide Count |
|---|---|---|---|---|---|---|
| Q9JLM8 | DCLK1_MOUSE | DASGMLYNLASAIK | 5.02 | 3.79E-04 | 0.0143 | 10 |
| Q9JLM8 | DCLK1_MOUSE | SFEALLADLTR | 4.13 | 5.43E-04 | 0.0143 | 10 |
| Q2QL88 | CAZA2_MICMU F- | IVEAAENEYQTAISENYQTM[+16]SDTTFK | 4.30 | 9.14E-04 | 0.0143 | 10 |
| Q9ERD7 | TBB3_MOUSE T | M[+16]SSTFIGNSTAIQELFK | 2.00 | 1.01E-03 | 0.0143 | 10 |
| Q9ERD7 | TBB3_MOUSE T | LATPTYGDLNHLVSATM[+16]SGVTTSLR | 2.35 | 1.29E-03 | 0.0143 | 10 |
| Q9JLM8 | DCLK1_MOUSE | TAHSFEQVLTDITDAIK | 6.04 | 1.45E-03 | 0.0143 | 10 |
| Q9ERD7 | TBB3_MOUSE T | ISVYYNEASSHK | 1.93 | 1.49E-03 | 0.0143 | 10 |
| Q2QL88 | CAZA2_MICMU F- | LLLNNDNLLR | 4.37 | 1.62E-03 | 0.0143 | 10 |
| Q9ERD7 | TBB3_MOUSE T | MSSTFIGNSTAIQELFK | 2.80 | 1.75E-03 | 0.0143 | 10 |
| Q2QL88 | CAZA2_MICMU F- | IEGYEDQVLITEHGDLGNGK | 4.50 | 1.76E-03 | 0.0143 | 10 |
| Q2QL88 | CAZA2_MICMU F- | TSVETALR | 3.99 | 1.84E-03 | 0.0143 | 10 |
| Q9ERD7 | TBB3_MOUSE T | EVDEQM[+16]LAIQSK | 1.64 | 1.96E-03 | 0.0143 | 10 |
| Q2QL88 | CAZA2_MICMU F- | IQVHYYEDGNVQLVSHK | 4.74 | 2.28E-03 | 0.0143 | 10 |
| Q2QL88 | CAZA2_MICMU F- | EHYPNGVC[+57]TVYGK | 4.29 | 2.29E-03 | 0.0143 | 10 |
| Q2QL88 | CAZA2_MICMU F- | EATDPRPYEAENAIESWR | 4.78 | 2.54E-03 | 0.0143 | 10 |
| Q9JLM8 | DCLK1_MOUSE | ISQHGGSSTSLSSTK | 2.78 | 2.87E-03 | 0.0143 | 10 |
| Q9JLM8 | DCLK1_MOUSE | SPSPSPTSPGSLR | 4.40 | 2.98E-03 | 0.0143 | 10 |
| Q2QL88 | CAZA2_MICMU F- | FIIHAPPGEFNEVFNDVR | 5.70 | 3.00E-03 | 0.0143 | 10 |
| Q9ERD7 | TBB3_MOUSE T | LATPTYGDLNHLVSATMSGVTTSLR | 3.75 | 3.15E-03 | 0.0143 | 10 |
| Q2QL88 | CAZA2_MICMU F- | EGAAHAFAQYNLDQFTPVK | 4.61 | 3.32E-03 | 0.0143 | 10 |
| Q2QL88 | CAZA2_MICMU F- | IVEAAENEYQTAISENYQTMSDTTFK | 4.98 | 3.80E-03 | 0.0147 | 10 |
| Q9JLM8 | DCLK1_MOUSE | ISSLDQLVEGESYVC[+57]GSIEPFK | 5.23 | 4.04E-03 | 0.0148 | 10 |
| Q9JLM8 | DCLK1_MOUSE | GIVYAISPDR | 2.84 | 4.32E-03 | 0.0150 | 10 |
| Q9JLM8 | DCLK1_MOUSE | GGDLFDAITSTSK | 3.34 | 6.74E-03 | 0.0172 | 10 |
| Q9JLM8 | DCLK1_MOUSE | TLSDNVNLPQGVR | 3.24 | 8.12E-03 | 0.0185 | 10 |
| Q9ERD7 | TBB3_MOUSE T | FWEVISDEHGIDPSGNYVGDSDLQLER | 4.91 | 8.77E-03 | 0.0192 | 10 |
| Q9ERD7 | TBB3_MOUSE T | EVDEQMLAIQSK | 2.03 | 9.77E-03 | 0.0204 | 10 |
| Q9JLM8 | DCLK1_MOUSE | DIKPENLLVYEHQDGSK | 4.30 | 1.62E-02 | 0.0273 | 10 |
| P46096 | SYT1_MOUSE S | VQVVVTVLDYDK | 2.34 | 1.86E-02 | 0.0300 | 10 |
| Q04447 | KCRB_MOUSE | RGTGGVDTAAVGGVFDVSNADR | 6.43 | 1.95E-02 | 0.0309 | 10 |
| Q8K0U4 | HS12A_MOUSE | LDLTGSGGTAVPAR | 1.89 | 6.83E-04 | 0.0143 | 10 |
| P61161 | ARP2_MOUSE A | HM[+16]VFLGGAVLADIMK | 3.68 | 8.98E-04 | 0.0143 | 10 |
| Q8K0U4 | HS12A_MOUSE | IFGEDFIEQFK | 2.78 | 1.06E-03 | 0.0143 | 10 |
| P61161 | ARP2_MOUSE A | HMVFLGGAVLADIMK | 4.55 | 1.80E-03 | 0.0143 | 10 |
| Q8K0U4 | HS12A_MOUSE | ATAVDITTSK | 1.78 | 2.08E-03 | 0.0143 | 10 |
| Q8K0U4 | HS12A_MOUSE | SPLTYGVGVLNR | 1.75 | 2.16E-03 | 0.0143 | 10 |
| Q8K0U4 | HS12A_MOUSE | IIIPQDVGLTILK | 1.75 | 2.20E-03 | 0.0143 | 10 |
| Q99JY9 | ARP3_MOUSE A | TLTGTVIDSGDGVTHVIPVAEGYVIGSC[+57]IK | 4.32 | 2.47E-03 | 0.0143 | 10 |
| Q99JY9 | ARP3_MOUSE A | NIVLSGGSTM[+16]FR | 3.46 | 2.53E-03 | 0.0143 | 10 |
| Q8K0U4 | HS12A_MOUSE | TNPLNITLPFSFIDYYK | 3.98 | 2.95E-03 | 0.0143 | 10 |
| P61161 | ARP2_MOUSE A | ILLTEPPM[+16]NPTK | 2.98 | 3.53E-03 | 0.0144 | 10 |
| Q8K0U4 | HS12A_MOUSE | VGIDFLNY | 1.92 | 3.63E-03 | 0.0146 | 10 |
| P61161 | ARP2_MOUSE A | VVVC[+57]DNGTGFVK | 2.34 | 4.00E-03 | 0.0148 | 10 |
| Q99JY9 | ARP3_MOUSE A | QYTGVNAISK | 2.42 | 4.00E-03 | 0.0148 | 10 |
| Q8K0U4 | HS12A_MOUSE | FISADQSVALGELVK | 2.57 | 4.05E-03 | 0.0148 | 10 |
| Q99JY9 | ARP3_MOUSE A | LSEELSGGR | 2.48 | 4.07E-03 | 0.0148 | 10 |
| Q99JY9 | ARP3_MOUSE A | DYEEIGPSIC[+57]R | 2.75 | 4.08E-03 | 0.0148 | 10 |
| P61161 | ARP2_MOUSE A | GYAFNHSADFETVR | 3.32 | 4.85E-03 | 0.0154 | 10 |
| Q99JY9 | ARP3_MOUSE A | LPAC[+57]VVDC[+57]GTGYTK | 3.44 | 6.81E-03 | 0.0172 | 10 |
| P61161 | ARP2_MOUSE A | LALETTVLVESYTLPDGR | 5.68 | 7.03E-03 | 0.0175 | 10 |
| Q99JY9 | ARP3_MOUSE A | GVDDLDFFIGDEAIEKPTYATK | 4.05 | 7.35E-03 | 0.0178 | 10 |
| P61161 | ARP2_MOUSE A | DLMVGDEASELR | 2.55 | 7.67E-03 | 0.0181 | 10 |
| Q99JY9 | ARP3_MOUSE A | FMEQVIFK | 2.60 | 7.77E-03 | 0.0182 | 10 |
| Q8K0U4 | HS12A_MOUSE | GAVLFGLDPAVIK | 1.76 | 8.07E-03 | 0.0184 | 10 |
| Q99JY9 | ARP3_MOUSE A | NIVLSGGSTMFR | 2.98 | 8.38E-03 | 0.0188 | 10 |
| Q9WUM4 | COR1C_MOUSE | NDQC[+57]YDDIR | 3.83 | 1.07E-02 | 0.0214 | 10 |
| Q9WUM4 | COR1C_MOUSE | YFEITDESPYVHYLNTFSSK | 5.49 | 1.10E-02 | 0.0217 | 10 |
| Q9WUM4 | COR1C_MOUSE | AIFLADGNVFTTGFSR | 4.09 | 1.11E-02 | 0.0219 | 10 |
| P61161 | ARP2_MOUSE A | KVVVC[+57]DNGTGFVK | 3.50 | 1.32E-02 | 0.0240 | 10 |
| P61161 | ARP2_MOUSE A | ILLTEPPMNPTK | 2.92 | 1.35E-02 | 0.0243 | 10 |
| Q9WUM4 | COR1C_MOUSE | NGSLIC[+57]TASK | 3.43 | 1.52E-02 | 0.0263 | 10 |
| Q8K0U4 | HS12A_MOUSE | ETAPTSTYSSPAR | 1.53 | 1.58E-02 | 0.0269 | 10 |

| Protein Accession | Protein Description | Modified Peptide Sequence | co-IP:Ctrl | p value | q value | Sig Peptide Count |
|---|---|---|---|---|---|---|
| Q9WUM4 | COR1C_MOUSE | NADPILISLK | 3.31 | 1.63E-02 | 0.0274 | 10 |
| Q9CPV4 | GLOD4_MOUSE | VAEGIFETEAPGGYK | 3.82 | 1.73E-02 | 0.0284 | 10 |
| P62281 | RS11_MOUSE | DVQIGDIVTVGEC[+57]RPLSK | 3.31 | 1.74E-02 | 0.0285 | 10 |
| E9QPE7 | E9QPE7_MOUSE | QLEEAEEESQR | 4.26 | 2.19E-02 | 0.0338 | 10 |
| P19253 | RL13A_MOUSE | C[+57]EGINISGNFYR | 3.27 | 2.63E-02 | 0.0386 | 10 |
| P62830 | RL23_MOUSE | LPAAGVGDMVMATVK | 3.30 | 2.71E-02 | 0.0394 | 10 |
| P60710 | ACTB_MOUSE | TTGIVMDSGDGVTHTVPIYEGYALPHAILR | 5.94 | 2.87E-02 | 0.0411 | 10 |
| Q03265 | ATPA_MOUSE | LKEIVTNFLAGFEP | 2.38 | 2.94E-02 | 0.0419 | 10 |
| Q9WUA3 | PFKAP_MOUSE | M[+16]GVEAVIALLEATPETPAC[+57]VVSLR | 3.21 | 7.23E-04 | 0.0143 | 10 |
| Q9WUA3 | PFKAP_MOUSE | EWSGLLEELAR | 2.16 | 1.56E-03 | 0.0143 | 10 |
| Q9WUA3 | PFKAP_MOUSE | WDC[+57]VSSILQVGGTIIGSAR | 2.59 | 1.66E-03 | 0.0143 | 10 |
| Q9WUA3 | PFKAP_MOUSE | LGITNLC[+57]VIGGDGSLTGANLFR | 2.45 | 2.50E-03 | 0.0143 | 10 |
| Q9WUA3 | PFKAP_MOUSE | VYFIYEGYQGLVDGGSNIVEAK | 3.15 | 2.74E-03 | 0.0143 | 10 |
| Q9WUA3 | PFKAP_MOUSE | MGVEAVIALLEATPETPAC[+57]VVSLR | 7.33 | 3.35E-03 | 0.0143 | 10 |
| Q9WUA3 | PFKAP_MOUSE | NESC[+57]SVNYTTDFIYQLYSEEGK | 3.43 | 5.17E-03 | 0.0155 | 10 |
| Q9WUA3 | PFKAP_MOUSE | FVSDDSIC[+57]VLGIC[+57]K | 1.72 | 5.37E-03 | 0.0158 | 10 |
| Q9WUA3 | PFKAP_MOUSE | IIEVVDAIMTTAQSHQR | 2.50 | 8.10E-03 | 0.0184 | 10 |
| Q9WUA3 | PFKAP_MOUSE | DLLFQPVAELK | 1.91 | 1.05E-02 | 0.0212 | 10 |
| P50516 | VATA_MOUSE | ADYAQLLEDMQNAFR | 3.20 | 7.34E-04 | 0.0143 | 10 |
| Q8K1M6 | DNM1L_MOUSE | LHDAIVEVVTC[+57]LLR | 3.25 | 8.57E-04 | 0.0143 | 10 |
| P17427 | AP2A2_MOUSE | TSVSLAVSR | 2.14 | 1.71E-03 | 0.0143 | 10 |
| P17427 | AP2A2_MOUSE | IIGFGSALLEEVDPNPANFVGAGIIHTK | 4.93 | 2.10E-03 | 0.0143 | 10 |
| P50516 | VATA_MOUSE | EILQEEEDLAEIVQLVGK | 2.59 | 2.20E-03 | 0.0143 | 10 |
| P50516 | VATA_MOUSE | YSNSDVIIYVGC[+57]GER | 2.24 | 2.23E-03 | 0.0143 | 10 |
| Q8K1M6 | DNM1L_MOUSE | LQDVFNTVGADIIQLPQIVVVGTQSSGK | 3.14 | 2.30E-03 | 0.0143 | 10 |
| P17427 | AP2A2_MOUSE | LSTVASTDILATVLEEM[+16]PPFPER | 3.70 | 2.82E-03 | 0.0143 | 10 |
| Q8K1M6 | DNM1L_MOUSE | LYTDFDEIRQEIENETER | 2.82 | 2.95E-03 | 0.0143 | 10 |
| P17427 | AP2A2_MOUSE | ILVAGDTM[+16]DSVK | 1.23 | 3.82E-03 | 0.0147 | 10 |
| Q8K1M6 | DNM1L_MOUSE | TLESVDPLGGLNTIDILTAIR | 2.27 | 4.29E-03 | 0.0150 | 10 |
| P17427 | AP2A2_MOUSE | QLSNPQQEVQNIFK | 1.87 | 4.36E-03 | 0.0150 | 10 |
| P17427 | AP2A2_MOUSE | FVNLFPEVK | 2.06 | 5.32E-03 | 0.0158 | 10 |
| P17427 | AP2A2_MOUSE | AVDLLYAMC[+57]DR | 2.37 | 5.34E-03 | 0.0158 | 10 |
| Q8K1M6 | DNM1L_MOUSE | YIETSELC[+57]GGAR | 2.35 | 5.48E-03 | 0.0159 | 10 |
| P50516 | VATA_MOUSE | FSMVQVWPVR | 1.76 | 7.96E-03 | 0.0183 | 10 |
| P50516 | VATA_MOUSE | VLDALFPC[+57]VQGGTTAIPGAFGC[+57]GK | 1.55 | 9.78E-03 | 0.0204 | 10 |
| P50516 | VATA_MOUSE | EASIYTGITLSEYFR | 4.64 | 1.16E-02 | 0.0224 | 10 |
| Q8K1M6 | DNM1L_MOUSE | SSLLDDLLTESEDMAQR | 3.08 | 1.16E-02 | 0.0224 | 10 |
| P50516 | VATA_MOUSE | VGSHITGGDIYGIVNENSLIK | 1.66 | 1.17E-02 | 0.0225 | 10 |
| Q8K1M6 | DNM1L_MOUSE | LGIIGVVNR | 1.10 | 1.20E-02 | 0.0228 | 10 |
| P50516 | VATA_MOUSE | LIKDDFLQQNGYTPYDR | 1.42 | 1.28E-02 | 0.0235 | 10 |
| Q8K1M6 | DNM1L_MOUSE | FATEYC[+57]NTIEGTAK | 6.30 | 1.32E-02 | 0.0240 | 10 |
| P17427 | AP2A2_MOUSE | LSTVASTDILATVLEEMPPFPER | 4.30 | 1.40E-02 | 0.0249 | 10 |
| P50516 | VATA_MOUSE | GSVTYIAPPGNYDASDVVLELEFEGVK | 5.12 | 1.55E-02 | 0.0266 | 10 |
| P20357 | MTAP2_MOUSE | NKDDLTLSR | 0.83 | 2.10E-02 | 0.0328 | 10 |
| Q80ZK2 | Q80ZK2_MOUSE | VPDTAWDGTQSK | 2.25 | 2.29E-02 | 0.0349 | 10 |
| O35643 | AP1B1_MOUSE | GLEISGTFTR | 8.80 | 2.40E-02 | 0.0361 | 10 |
| Q62261 | SPTB2_MOUSE | NDSFTAC[+57]IELGK | 0.84 | 2.50E-02 | 0.0371 | 10 |
| P59999 | ARPC4_MOUSE | IVAEEFLK | 1.07 | 3.09E-02 | 0.0436 | 10 |
| P62137 | PP1A_MOUSE | IFC[+57]C[+57]HGGLSPDLQSM[+16]EQIR | 3.50 | 2.13E-04 | 0.0143 | 9 |
| Q9WV92 | E41L3_MOUSE | VESTSVGSISPGGAK | 2.16 | 2.52E-04 | 0.0143 | 9 |
| Q9WV92 | E41L3_MOUSE | LM[+16]DGSEILSLLESAR | 5.19 | 2.68E-04 | 0.0143 | 9 |
| Q9WV92 | E41L3_MOUSE | LMDGSEILSLLESAR | 7.22 | 4.87E-04 | 0.0143 | 9 |
| P62137 | PP1A_MOUSE | EIFLSQPILLELEAPLK | 5.04 | 6.15E-04 | 0.0143 | 9 |
| P62137 | PP1A_MOUSE | SREIFLSQPILLELEAPLK | 6.16 | 1.46E-03 | 0.0143 | 9 |
| P62137 | PP1A_MOUSE | NVQLTENEIR | 3.03 | 2.24E-03 | 0.0143 | 9 |
| P62137 | PP1A_MOUSE | TFTDC[+57]FNC[+57]LPIAAIVDEK | 4.63 | 2.69E-03 | 0.0143 | 9 |
| P62137 | PP1A_MOUSE | LNLDSIIGR | 3.81 | 3.30E-03 | 0.0143 | 9 |
| P62137 | PP1A_MOUSE | HDLDLIC[+57]R | 3.22 | 4.51E-03 | 0.0152 | 9 |
| P62137 | PP1A_MOUSE | IC[+57]GDIHGQYYDLLR | 3.69 | 7.06E-03 | 0.0175 | 9 |
| Q9WV92 | E41L3_MOUSE | TEPVEAEVESTPHPQPLSTEK | 2.58 | 7.11E-03 | 0.0176 | 9 |
| Q9WV92 | E41L3_MOUSE | KPTEFIGGVSSTTQSWVQK | 2.98 | 8.72E-03 | 0.0191 | 9 |
| Q9WV92 | E41L3_MOUSE | IRPGEFEQFESTIGFK | 3.83 | 9.68E-03 | 0.0204 | 9 |

| Protein Accession | Protein Description | Modified Peptide Sequence | co-IP:Ctrl | p value | q value | Sig Peptide Count |
|---|---|---|---|---|---|---|
| Q9WV92 | E41L3_MOUSE | IVITGDADIDHDQALAQAIK | 3.02 | 1.55E-02 | 0.0266 | 9 |
| Q9WV92 | E41L3_MOUSE | GISQTNLITTVTPEK | 1.53 | 1.62E-02 | 0.0273 | 9 |
| P16546 | SPTN1_MOUSE | SSLSSAQADFNQLAELDR | 2.41 | 2.64E-02 | 0.0387 | 9 |
| O35737 | HNRH1_MOUSE | EGRPSGEAFVELESEDEVK | 5.79 | 2.76E-02 | 0.0399 | 9 |
| Q8BPN8 | DMXL2_MOUSE | QLQSPLPLPTTLPLLSASIASTK | 5.69 | 1.05E-04 | 0.0143 | 9 |
| P63318 | KPCG_MOUSE | LGSGPDGEPTIR | 4.54 | 8.75E-04 | 0.0143 | 9 |
| Q60737 | CSK21_MOUSE | TPALVFEHVNNTDFK | 5.19 | 2.03E-03 | 0.0143 | 9 |
| P42669 | PURA_MOUSE | DYLGDFIEHYAQLGPSQPPDLAQAQDEPR | 6.15 | 2.27E-03 | 0.0143 | 9 |
| P42669 | PURA_MOUSE | LIDDYGVEEEPAELPEGTSLTVDNKR | 2.91 | 2.60E-03 | 0.0143 | 9 |
| P42669 | PURA_MOUSE | FFFDVGSNK | 2.34 | 3.06E-03 | 0.0143 | 9 |
| Q60737 | CSK21_MOUSE | GGPNIITLADIVK | 4.40 | 3.48E-03 | 0.0144 | 9 |
| Q60737 | CSK21_MOUSE | FVHSENQHLVSPEALDFLDK | 5.11 | 4.12E-03 | 0.0148 | 9 |
| Q60737 | CSK21_MOUSE | VLGTEDLYDYIDKYNIELDPR | 6.45 | 4.17E-03 | 0.0149 | 9 |
| Q60737 | CSK21_MOUSE | GGPNIITLADIVKDPVSR | 5.11 | 4.79E-03 | 0.0154 | 9 |
| Q60737 | CSK21_MOUSE | YSEVFEAINITNNEK | 3.36 | 5.40E-03 | 0.0158 | 9 |
| P42669 | PURA_MOUSE | GPGLGSTQGQTIALPAQGLIEFR | 2.39 | 5.53E-03 | 0.0159 | 9 |
| Q60737 | CSK21_MOUSE | FNDILGR | 4.57 | 5.58E-03 | 0.0160 | 9 |
| Q60737 | CSK21_MOUSE | LIDWGLAEFYHPGQEYNVR | 5.52 | 5.63E-03 | 0.0160 | 9 |
| P42669 | PURA_MOUSE | IAEVGAGGNK | 2.21 | 5.65E-03 | 0.0160 | 9 |
| P63318 | KPCG_MOUSE | TFC[+57]GTPDYIAPEIIAYQPYGK | 3.20 | 5.76E-03 | 0.0161 | 9 |
| Q8BPN8 | DMXL2_MOUSE | LVYSQPLDLPEAVEVIR | 5.85 | 6.33E-03 | 0.0169 | 9 |
| P42669 | PURA_MOUSE | VSEVKPTYR | 2.44 | 6.37E-03 | 0.0169 | 9 |
| Q60737 | CSK21_MOUSE | QLYQTLTDYDIR | 3.89 | 6.39E-03 | 0.0169 | 9 |
| P42669 | PURA_MOUSE | NSITVPYK | 2.18 | 7.63E-03 | 0.0181 | 9 |
| P42669 | PURA_MOUSE | SEFLVR | 2.05 | 1.01E-02 | 0.0207 | 9 |
| P63318 | KPCG_MOUSE | APTSDEIHITVGEAR | 3.78 | 1.12E-02 | 0.0219 | 9 |
| P63318 | KPCG_MOUSE | LVLASIDQADFQGFTYVNPDFVHPDAR | 8.23 | 1.32E-02 | 0.0240 | 9 |
| Q8BPN8 | DMXL2_MOUSE | SIDLVSSVDGTPSLPVSLSWVR | 6.13 | 1.55E-02 | 0.0266 | 9 |
| P42669 | PURA_MOUSE | LIDDYGVEEEPAELPEGTSLTVDNK | 3.69 | 1.61E-02 | 0.0272 | 9 |
| P63318 | KPCG_MOUSE | DVIVQDDDVDC[+57]TLVEK | 5.37 | 1.66E-02 | 0.0277 | 9 |
| P01869 | IGH1M_MOUSE | TTPPSVYPLAPGSAAQTNSMVTLGC[+57]LVK | 2.54 | 1.77E-02 | 0.0289 | 9 |
| P47963 | RL13_MOUSE 6 | LATQLTGPVMPIR | 4.27 | 1.96E-02 | 0.0310 | 9 |
| P10126 | EF1A1_MOUSE | MDSTEPPYSQK | 6.37 | 2.04E-02 | 0.0321 | 9 |
| Q99KX1 | MLF2_MOUSE I | VYQETSEMR | 2.21 | 2.09E-02 | 0.0326 | 9 |
| Q3UM45 | PP1R7_MOUSE | LQNLDALTNLTVLSVQSNR | 3.30 | 2.28E-02 | 0.0348 | 9 |
| P62301 | RS13_MOUSE 4 | GLAPDLPEDLYHLIK | 3.42 | 2.35E-02 | 0.0356 | 9 |
| P17182 | ENOA_MOUSE | LAM[+16]QEFM[+16]ILPVGASSFR | 2.75 | 2.82E-02 | 0.0406 | 9 |
| P39053 | DYN1_MOUSE I | VPVGDQPPDIEFQIR | 4.89 | 3.09E-02 | 0.0436 | 9 |
| Q9JMH9 | MY18A_MOUSE | ISELTSELTDER | 8.52 | 3.23E-02 | 0.0452 | 9 |
| Q9QYR6 | MAP1A_MOUSE | SIEEAC[+57]LTLQHLNR | 2.37 | 3.32E-02 | 0.0463 | 9 |
| P10126 | EF1A1_MOUSE | VETGVLKPGM[+16]VVTFAPVNVTTEVK | 1.40 | 8.66E-04 | 0.0143 | 9 |
| P01837 | IGKC_MOUSE I | DSTYSM[+16]SSTLTLTKDEYER | 6.36 | 1.43E-03 | 0.0143 | 9 |
| P01837 | IGKC_MOUSE I | TSTSPIVK | 6.86 | 2.04E-03 | 0.0143 | 9 |
| P10126 | EF1A1_MOUSE | THINIVVIGHVDSGK | 2.25 | 2.80E-03 | 0.0143 | 9 |
| P10126 | EF1A1_MOUSE | DGSASGTTLLEALDC[+57]ILPPTRPTDKPLR | 4.02 | 2.89E-03 | 0.0143 | 9 |
| P01837 | IGKC_MOUSE I | QNGVLNSWTDQDSKDSTYSM[+16]SSTLTLTK | 8.72 | 3.27E-03 | 0.0143 | 9 |
| P10126 | EF1A1_MOUSE | VETGVLKPGMVVTFAPVNVTTEVK | 2.02 | 3.63E-03 | 0.0146 | 9 |
| P10126 | EF1A1_MOUSE | NM[+16]ITGTSQADC[+57]AVLIVAAGVGEFEAGISK | 1.71 | 5.87E-03 | 0.0163 | 9 |
| P01837 | IGKC_MOUSE I | RADAAPTVSIFPPSSEQLTSGGASVVC[+57]FLNNFYPK | 9.03 | 6.63E-03 | 0.0172 | 9 |
| P10126 | EF1A1_MOUSE | YYVTIIDAPGHR | 1.42 | 7.54E-03 | 0.0180 | 9 |
| P10126 | EF1A1_MOUSE | EHALLAYTLGVK | 2.07 | 7.55E-03 | 0.0180 | 9 |
| P01837 | IGKC_MOUSE I | DSTYSMSSTLTLTKDEYER | 7.00 | 9.22E-03 | 0.0198 | 9 |
| P01837 | IGKC_MOUSE I | QNGVLNSWTDQDSKDSTYSMSSTLTLTK | 8.33 | 1.45E-02 | 0.0255 | 9 |
| P62270 | RS18_MOUSE 4 | IAFAITAIK | 6.09 | 2.03E-02 | 0.0319 | 9 |
| Q6ZQ38 | CAND1_MOUSE | VIRPLDQPSSFDATPYIK | 1.42 | 2.03E-02 | 0.0319 | 9 |
| P62301 | RS13_MOUSE 4 | LILIESR | 6.81 | 2.33E-02 | 0.0354 | 9 |
| P62821 | RAB1A_MOUSE | EFADSLGIPFLETSAK | 7.57 | 2.39E-02 | 0.0360 | 9 |
| P67984 | RL22_MOUSE 6 | ITVTSEVPFSK | 1.97 | 2.94E-02 | 0.0419 | 9 |
| P62814 | VATB2_MOUSE | IYPEEMIQTGISAIDGMNSIAR | 3.32 | 1.32E-03 | 0.0143 | 9 |
| P62814 | VATB2_MOUSE | AVVGEEALTSDDLLYLEFLQK | 2.24 | 1.86E-03 | 0.0143 | 9 |
| P62814 | VATB2_MOUSE | GPVVLAEDFLDIMGQPINPQC[+57]R | 4.74 | 2.18E-03 | 0.0143 | 9 |
| P62814 | VATB2_MOUSE | HVLVILTDMSSYAEALR | 4.02 | 2.93E-03 | 0.0143 | 9 |

| Protein Accession | Protein Description | Modified Peptide Sequence | co-IP:Ctrl | p value | q value | Sig Peptide Count |
|---|---|---|---|---|---|---|
| P62814 | VATB2_MOUSE | LALTTAEFLAYQC[+57]EK | 2.05 | 4.42E-03 | 0.0151 | 9 |
| P62814 | VATB2_MOUSE | IPIFSAAGLPHNEIAAQIC[+57]R | 2.11 | 4.59E-03 | 0.0152 | 9 |
| P62814 | VATB2_MOUSE | TSC[+57]EFTGDILR | 1.29 | 5.85E-03 | 0.0163 | 9 |
| P62814 | VATB2_MOUSE | AVVQVFEGTSGIDAK | 1.50 | 6.57E-03 | 0.0171 | 9 |
| P62814 | VATB2_MOUSE | IPQSTLSEFYPR | 2.46 | 8.93E-03 | 0.0194 | 9 |
| Q9DBG3 | AP2B1_MOUSE | APEVSQYIYQVYDSILK | 3.52 | 4.04E-04 | 0.0143 | 9 |
| Q9DBG3 | AP2B1_MOUSE | LAPPLVTLLSGEPEVQYVALR | 2.73 | 5.52E-04 | 0.0143 | 9 |
| Q9DBG3 | AP2B1_MOUSE | KPSETQELVQQVLSLATQDSDNPDLR | 3.77 | 8.81E-04 | 0.0143 | 9 |
| P84091 | AP2M1_MOUSE | LNYSDHDVIK | 3.07 | 1.36E-03 | 0.0143 | 9 |
| Q9DBG3 | AP2B1_MOUSE | SQPDMAIMAVNSFVK | 3.84 | 2.31E-03 | 0.0143 | 9 |
| P84091 | AP2M1_MOUSE | SNIWLAAVTK | 3.03 | 2.61E-03 | 0.0143 | 9 |
| P84091 | AP2M1_MOUSE | NAVDAFR | 2.43 | 3.17E-03 | 0.0143 | 9 |
| P84091 | AP2M1_MOUSE | QNVNAAM[+16]VFEFLYK | 4.48 | 4.41E-03 | 0.0151 | 9 |
| Q9DBG3 | AP2B1_MOUSE | FLELLPK | 1.42 | 5.36E-03 | 0.0158 | 9 |
| P84091 | AP2M1_MOUSE | EEQSQITSQVTGQIGWR | 1.89 | 5.88E-03 | 0.0163 | 9 |
| P84091 | AP2M1_MOUSE | QSIAIDDC[+57]TFHQC[+57]VR | 2.23 | 6.42E-03 | 0.0169 | 9 |
| P84091 | AP2M1_MOUSE | IPTPLNTSGVQVIC[+57]MK | 5.53 | 7.81E-03 | 0.0182 | 9 |
| P84091 | AP2M1_MOUSE | MC[+57]DVMAAYFGK | 3.47 | 9.76E-03 | 0.0204 | 9 |
| Q9DBG3 | AP2B1_MOUSE | LQNNNVYTIAK | 1.46 | 1.07E-02 | 0.0214 | 9 |
| P84091 | AP2M1_MOUSE | ESQISAEIELLPTNDK | 3.48 | 1.28E-02 | 0.0235 | 9 |
| Q9DBG3 | AP2B1_MOUSE | DIPNENELQFQIK | 2.30 | 1.68E-02 | 0.0279 | 9 |
| Q9JMH9 | MY18A_MOUSE | VASGSDLHLTDIDSDSNR | 0.96 | 2.73E-02 | 0.0396 | 9 |
| Q9Z1N5 | DX39B_MOUSE | GLAITFVSDENDAK | 3.42 | 3.18E-02 | 0.0446 | 9 |
| Q03265 | ATPA_MOUSE / | ILGADTSVDLEETGR | 2.08 | 5.28E-04 | 0.0143 | 9 |
| Q03265 | ATPA_MOUSE / | EVAAFAQFGSDLDAATQQLLSR | 3.31 | 2.16E-03 | 0.0143 | 9 |
| Q03265 | ATPA_MOUSE / | TSIAIDTIINQK | 1.51 | 2.24E-03 | 0.0143 | 9 |
| Q03265 | ATPA_MOUSE / | NVQAEEM[+16]VEFSSGLK | 5.48 | 2.67E-03 | 0.0143 | 9 |
| Q03265 | ATPA_MOUSE / | NVQAEEMVEFSSGLK | 2.37 | 3.69E-03 | 0.0147 | 9 |
| Q03265 | ATPA_MOUSE / | TGAIVDVPVGEELLGR | 1.37 | 9.87E-03 | 0.0205 | 9 |
| Q03265 | ATPA_MOUSE / | TGTAEMSSILEER | 1.81 | 1.18E-02 | 0.0226 | 9 |
| Q03265 | ATPA_MOUSE / | EIVTNFLAGFEP | 1.53 | 1.67E-02 | 0.0278 | 9 |
| Q62261 | SPTB2_MOUSE | SNAHYNLQNAFNLAEQHLGLTK | 3.47 | 2.93E-02 | 0.0418 | 9 |
| P63038 | CH60_MOUSE ( | VGEVIVTK | 2.85 | 1.25E-03 | 0.0143 | 9 |
| P63038 | CH60_MOUSE ( | GVMLAVDAVIAELKK | 2.48 | 3.23E-03 | 0.0143 | 9 |
| P63038 | CH60_MOUSE ( | ALM[+16]LQGVDLLADAVAVTMGPK | 2.24 | 4.44E-03 | 0.0151 | 9 |
| P63038 | CH60_MOUSE ( | GYISPYFINTSK | 3.90 | 8.47E-03 | 0.0189 | 9 |
| P63038 | CH60_MOUSE ( | C[+57]EFQDAYVLLSEK | 2.97 | 9.51E-03 | 0.0202 | 9 |
| P16546 | SPTN1_MOUSE | HQALQAEIAGHEPR | 1.63 | 1.80E-02 | 0.0293 | 9 |
| P47911 | RL6_MOUSE 60 | QLDSGLLLVTGPLVINR | 1.58 | 1.84E-02 | 0.0297 | 9 |
| Q9DBJ1 | PGAM1_MOUSE | YADLTEDQLPSC[+57]ESLK | 1.07 | 2.36E-02 | 0.0357 | 9 |
| P68033 | ACTC_MOUSE . | H[+42]QGVM[+16]VGM[+16]GQK | 1.08 | 3.21E-02 | 0.0450 | 9 |
| Q99KI0 | ACON_MOUSE | VAMQDATAQMAMLQFISSGLPK | 3.34 | 2.31E-03 | 0.0143 | 9 |
| Q99KI0 | ACON_MOUSE | NDANPETHAFVTSPEIVTALAIAGTLK | 7.49 | 3.17E-03 | 0.0143 | 9 |
| Q99KI0 | ACON_MOUSE | AKDINQEVYNFLATAGAK | 2.21 | 4.40E-03 | 0.0150 | 9 |
| Q99KI0 | ACON_MOUSE | NAVTQEFGPVPDTAR | 2.52 | 6.31E-03 | 0.0169 | 9 |
| Q99KI0 | ACON_MOUSE | WVVIGDENYGEGSSR | 3.85 | 7.53E-03 | 0.0180 | 9 |
| Q99KI0 | ACON_MOUSE | DVGGIVLANAC[+57]GPC[+57]IGQWDR | 5.57 | 9.15E-03 | 0.0197 | 9 |
| Q99KI0 | ACON_MOUSE | GHLDNISNNLLIGAINIENGK | 1.54 | 1.28E-02 | 0.0235 | 9 |
| Q99KI0 | ACON_MOUSE | FNPETDFLTGK | 1.25 | 1.51E-02 | 0.0262 | 9 |
| Q923T9 | KCC2G_MOUSE | IC[+57]DPGLTSFEPEALGNLVEGM[+16]DFHK | 1.34 | 1.88E-02 | 0.0301 | 9 |
| P17182 | ENOA_MOUSE | SFVQNYPVVSIEDPFDQDDWGAWQK | 4.18 | 4.76E-05 | 0.0143 | 9 |
| P17182 | ENOA_MOUSE | LAMQEFMILPVGASSFR | 2.17 | 1.81E-03 | 0.0143 | 9 |
| P17182 | ENOA_MOUSE | AGYTDQVVIGMDVAASEFYR | 2.48 | 2.68E-03 | 0.0143 | 9 |
| P17182 | ENOA_MOUSE | HIADLAGNPEVILPVPAFNVINGGSHAGNK | 1.85 | 2.88E-03 | 0.0143 | 9 |
| P17182 | ENOA_MOUSE | LMIEMDGTENK | 1.73 | 4.22E-03 | 0.0149 | 9 |
| P17182 | ENOA_MOUSE | FTASAGIQVVGDDLTVTNPK | 1.42 | 8.35E-03 | 0.0188 | 9 |
| P17182 | ENOA_MOUSE | LAM[+16]QEFMILPVGASSFR | 1.86 | 1.14E-02 | 0.0222 | 9 |
| P17182 | ENOA_MOUSE | DATNVGDEGGFAPNILENKEALELLK | 1.67 | 1.33E-02 | 0.0241 | 9 |
| Q8R071 | IP3KA_MOUSE | C[+57]AAVAAAAAAGEPR | 1.21 | 2.81E-02 | 0.0405 | 9 |
| Q9D8Y0 | EFHD2_MOUSE | SM[+16]IQEVDEDFDSK | 4.38 | 3.38E-04 | 0.0143 | 8 |
| Q9D8Y0 | EFHD2_MOUSE | FEEEIKAEQEER | 4.34 | 9.94E-04 | 0.0143 | 8 |
| P46660 | AINX_MOUSE A | LPASDGLDLSQAAAR | 2.90 | 1.13E-03 | 0.0143 | 8 |

| Protein Accession | Protein Description | Modified Peptide Sequence | co-IP:Ctrl | p value | q value | Sig Peptide Count |
|---|---|---|---|---|---|---|
| Q9D8Y0 | EFHD2_MOUSE | LSEIDVSTEGVK | 4.15 | 1.39E-03 | 0.0143 | 8 |
| Q9D8Y0 | EFHD2_MOUSE | RADLNQGIGEPQSPSR | 4.69 | 1.45E-03 | 0.0143 | 8 |
| V9GX76 | V9GX76_MOUSE | YLTESYGTGQDIDDR | 5.74 | 1.48E-03 | 0.0143 | 8 |
| P46660 | AINX_MOUSE A | FANLNEQAAR | 3.46 | 1.88E-03 | 0.0143 | 8 |
| Q9D8Y0 | EFHD2_MOUSE | ADLNQGIGEPQSPSR | 4.53 | 1.94E-03 | 0.0143 | 8 |
| Q9D8Y0 | EFHD2_MOUSE | SMIQEVDEDFDSK | 4.51 | 1.98E-03 | 0.0143 | 8 |
| V9GX76 | V9GX76_MOUSE | AGSLKDPLLDDHGDFIR | 5.48 | 1.99E-03 | 0.0143 | 8 |
| Q9D8Y0 | EFHD2_MOUSE | RVFNPYTEFK | 4.90 | 2.12E-03 | 0.0143 | 8 |
| Q3UHD9 | AGAP2_MOUSE | YEQLLFLAPLGTTEEPLGR | 2.36 | 2.26E-03 | 0.0143 | 8 |
| V9GX76 | V9GX76_MOUSE | IQAEVEAQLAR | 4.64 | 2.77E-03 | 0.0143 | 8 |
| Q9D8Y0 | EFHD2_MOUSE | VFNPYTEFK | 2.92 | 2.88E-03 | 0.0143 | 8 |
| P46660 | AINX_MOUSE A | VGELFQR | 4.91 | 2.95E-03 | 0.0143 | 8 |
| V9GX76 | V9GX76_MOUSE | VVAGVLHLGNIDFEEAGSTSGGC[+57]NLK | 5.47 | 3.51E-03 | 0.0144 | 8 |
| B9DGT7 | TBA2_ARATH Tub | AFVHWYVGEGM[+16]EEGEFSEAR | 3.70 | 3.87E-03 | 0.0147 | 8 |
| P46660 | AINX_MOUSE A | ALEAELAALR | 5.00 | 4.06E-03 | 0.0148 | 8 |
| Q3UHD9 | AGAP2_MOUSE | AEAAAVAGLSTPGSLHR | 2.60 | 4.29E-03 | 0.0150 | 8 |
| P46660 | AINX_MOUSE A | FSTGGLSISGLNPLPNPSYLLPPR | 7.67 | 4.75E-03 | 0.0154 | 8 |
| V9GX76 | V9GX76_MOUSE | EGLGVNEVHYVDNQDC[+57]IDLIEVK | 3.84 | 5.13E-03 | 0.0155 | 8 |
| P46660 | AINX_MOUSE A | KVESLLDELAFVR | 5.38 | 5.16E-03 | 0.0155 | 8 |
| Q3UHD9 | AGAP2_MOUSE | AVVNSQEWTLSR | 3.90 | 5.51E-03 | 0.0159 | 8 |
| V9GX76 | V9GX76_MOUSE | TFLALINQVFPAEEDSK | 8.25 | 6.13E-03 | 0.0166 | 8 |
| B9DGT7 | TBA2_ARATH Tub | IDHKFDLMYAK | 2.28 | 6.69E-03 | 0.0172 | 8 |
| B9DGT7 | TBA2_ARATH Tub | AVFVDLEPTVIDEVR | 2.10 | 6.70E-03 | 0.0172 | 8 |
| V9GX76 | V9GX76_MOUSE | HFAGAVC[+57]YETTQFVEK | 5.16 | 6.85E-03 | 0.0173 | 8 |
| B9DGT7 | TBA2_ARATH Tub | FDLM[+16]YAK | 1.53 | 7.16E-03 | 0.0176 | 8 |
| P46660 | AINX_MOUSE A | SNVASTAAC[+57]SSASSLGLGLAYR | 3.24 | 8.91E-03 | 0.0194 | 8 |
| Q3UHD9 | AGAP2_MOUSE | ALSTDC[+57]TPSGDLSPLSR | 6.19 | 9.80E-03 | 0.0204 | 8 |
| B9DGT7 | TBA2_ARATH Tub | AFVHWYVGEGMEEGEFSEAR | 4.71 | 1.02E-02 | 0.0208 | 8 |
| B9DGT7 | TBA2_ARATH Tub | TIQFVDWC[+57]PTGFK | 2.70 | 1.05E-02 | 0.0212 | 8 |
| Q3UHD9 | AGAP2_MOUSE | TDSQSEAVAIQAIR | 4.63 | 1.19E-02 | 0.0227 | 8 |
| B9DGT7 | TBA2_ARATH Tub | LSVDYGKK | 2.32 | 1.27E-02 | 0.0234 | 8 |
| Q3UHD9 | AGAP2_MOUSE | TTTYLISLTLVK | 4.28 | 1.41E-02 | 0.0250 | 8 |
| V9GX76 | V9GX76_MOUSE | ALFESSTNNNK | 6.08 | 1.46E-02 | 0.0256 | 8 |
| Q3UHD9 | AGAP2_MOUSE | LGVLGDVR | 4.69 | 1.49E-02 | 0.0259 | 8 |
| P46660 | AINX_MOUSE A | VESLLDELAFVR | 1.96 | 1.54E-02 | 0.0265 | 8 |
| B9DGT7 | TBA2_ARATH Tub | FDLMYAK | 2.33 | 1.60E-02 | 0.0271 | 8 |
| P06745 | G6PI_MOUSE G | EVMQMLVELAK | 1.97 | 2.90E-02 | 0.0414 | 8 |
| Q9R1Q8 | TAGL3_MOUSE | TTDIFQTVDLWEGK | 3.93 | 2.65E-04 | 0.0143 | 8 |
| Q8JZQ9 | EIF3B_MOUSE | NLFNVVDC[+57]K | 2.21 | 2.70E-04 | 0.0143 | 8 |
| Q9R1Q8 | TAGL3_MOUSE | GFSEEQLR | 3.38 | 7.13E-04 | 0.0143 | 8 |
| P62702 | RS4X_MOUSE 4 | TDITYPAGFMDVISIDK | 4.80 | 7.84E-04 | 0.0143 | 8 |
| Q8JZQ9 | EIF3B_MOUSE | GHPSAGAEEEGGSDGSAAEAEPR | 5.22 | 9.57E-04 | 0.0143 | 8 |
| P61205 | ARF3_MOUSE A | LGEIVTTIPTIGFNVETVEYK | 2.59 | 1.34E-03 | 0.0143 | 8 |
| Q61699 | HS105_MOUSE | LLTETEDWLYEEGEDQAK | 2.83 | 1.59E-03 | 0.0143 | 8 |
| Q61699 | HS105_MOUSE | FVVQNVSAQK | 2.24 | 2.45E-03 | 0.0143 | 8 |
| Q8R071 | IP3KA_MOUSE | GNVQLETSEDVGQK | 4.41 | 2.49E-03 | 0.0143 | 8 |
| Q8R071 | IP3KA_MOUSE | DTLEISDFFR | 5.21 | 2.68E-03 | 0.0143 | 8 |
| Q8R071 | IP3KA_MOUSE | MLAVDPEAPTEEEHAQR | 5.02 | 2.72E-03 | 0.0143 | 8 |
| P61205 | ARF3_MOUSE A | M[+16]LAEDELRDAVLLVFANK | 2.56 | 2.85E-03 | 0.0143 | 8 |
| Q8JZQ9 | EIF3B_MOUSE | IINDYYPEEDGK | 4.45 | 3.07E-03 | 0.0143 | 8 |
| Q9R1Q8 | TAGL3_MOUSE | LVDWIILQC[+57]AEDIEHPPPGR | 4.28 | 3.27E-03 | 0.0143 | 8 |
| Q9R1Q8 | TAGL3_MOUSE | TLMALGSVAVTK | 3.43 | 3.40E-03 | 0.0143 | 8 |
| P61205 | ARF3_MOUSE A | ILMVGLDAAGK | 1.62 | 4.03E-03 | 0.0148 | 8 |
| Q61699 | HS105_MOUSE | SVLDAAQIVGLNC[+57]LR | 3.53 | 4.33E-03 | 0.0150 | 8 |
| Q61699 | HS105_MOUSE | VVNVELPVEANLVWQLGR | 5.11 | 4.56E-03 | 0.0152 | 8 |
| Q61699 | HS105_MOUSE | AEDVSAIEIVGGATR | 3.10 | 4.62E-03 | 0.0153 | 8 |
| Q9D8E6 | RL4_MOUSE 60 | GPC[+57]IIYNEDNGIIK | 4.87 | 4.69E-03 | 0.0153 | 8 |
| Q8R071 | IP3KA_MOUSE | GAGPC[+57]SPGLER | 4.83 | 4.98E-03 | 0.0155 | 8 |
| P62702 | RS4X_MOUSE 4 | EC[+57]LPLIIFLR | 3.71 | 5.10E-03 | 0.0155 | 8 |
| Q61699 | HS105_MOUSE | VLGTAFDPFLGGK | 2.66 | 5.27E-03 | 0.0157 | 8 |
| Q61699 | HS105_MOUSE | RGPFELEAFYSDPQGVPYPEAK | 3.58 | 5.51E-03 | 0.0159 | 8 |
| P61205 | ARF3_MOUSE A | MLAEDELRDAVLLVFANK | 4.37 | 5.72E-03 | 0.0161 | 8 |

| Protein Accession | Protein Description | Modified Peptide Sequence | co-IP:Ctrl | p value | q value | Sig Peptide Count |
|---|---|---|---|---|---|---|
| Q8R071 | IP3KA_MOUSE | AGVWLIDFGK | 4.43 | 6.03E-03 | 0.0164 | 8 |
| Q8JZQ9 | EIF3B_MOUSE | AKPAAQSEEETATSPAASPTPQSAEF | 6.90 | 6.36E-03 | 0.0169 | 8 |
| P61205 | ARF3_MOUSE A | NISFTVWDVGGQDK | 2.14 | 6.65E-03 | 0.0172 | 8 |
| Q9R1Q8 | TAGL3_MOUSE | QMEQISQFLK | 3.07 | 6.75E-03 | 0.0172 | 8 |
| Q8R071 | IP3KA_MOUSE | AAGTSGLILK | 3.32 | 7.87E-03 | 0.0183 | 8 |
| Q9R1Q8 | TAGL3_MOUSE | LINSLYPPGQEPIPK | 3.16 | 8.24E-03 | 0.0186 | 8 |
| P62702 | RS4X_MOUSE 4 | HPGSFDVVHVK | 3.37 | 8.29E-03 | 0.0187 | 8 |
| Q61699 | HS105_MOUSE | SQFEELC[+57]AELLQK | 3.82 | 9.66E-03 | 0.0203 | 8 |
| Q8R071 | IP3KA_MOUSE | EGISSSTTLGFR | 2.11 | 9.97E-03 | 0.0206 | 8 |
| P62702 | RS4X_MOUSE 4 | VNDTIQIDLETGK | 3.40 | 1.02E-02 | 0.0208 | 8 |
| P62702 | RS4X_MOUSE 4 | LREC[+57]LPLIIFLR | 6.07 | 1.05E-02 | 0.0212 | 8 |
| Q9D8E6 | RL4_MOUSE 60 | RGPC[+57]IIYNEDNGIIK | 3.44 | 1.13E-02 | 0.0221 | 8 |
| P62702 | RS4X_MOUSE 4 | DANGNSFATR | 4.12 | 1.15E-02 | 0.0223 | 8 |
| Q9R1Q8 | TAGL3_MOUSE | TLM[+16]ALGSVAVTK | 1.54 | 1.20E-02 | 0.0228 | 8 |
| P62702 | RS4X_MOUSE 4 | YALTGDEVK | 2.35 | 1.22E-02 | 0.0230 | 8 |
| P61205 | ARF3_MOUSE A | DAVLLVFANK | 1.40 | 1.33E-02 | 0.0241 | 8 |
| P61205 | ARF3_MOUSE A | QDLPNAMNAAEITDK | 1.27 | 1.46E-02 | 0.0256 | 8 |
| Q9D8E6 | RL4_MOUSE 60 | KLEAAATALATK | 6.01 | 1.56E-02 | 0.0267 | 8 |
| Q9D8E6 | RL4_MOUSE 60 | NIPGITLLNVSK | 4.20 | 1.61E-02 | 0.0272 | 8 |
| Q8JZQ9 | EIF3B_MOUSE | GYIFLEYASPAHAVDAVK | 5.30 | 1.69E-02 | 0.0281 | 8 |
| Q9QY94 | GLNA_ACOCA Glu | RPSANC[+57]DPYAVTEAIVR | 4.88 | 1.72E-02 | 0.0284 | 8 |
| O08553 | DPYL2_MOUSE | FQM[+16]PDQGM[+16]TSADDFFQGTK | 3.77 | 1.81E-02 | 0.0294 | 8 |
| Q64331 | MYO6_MOUSE | NLEISIDALM[+16]AK | 2.65 | 1.93E-02 | 0.0307 | 8 |
| Q9WTM5 | RUVB2_MOUSE | FVQC[+57]PDGELQK | 1.47 | 1.94E-02 | 0.0308 | 8 |
| P13020 | GELS_MOUSE G | DPDQTDGPGLGYLSSHIANVER | 5.91 | 2.26E-02 | 0.0346 | 8 |
| P14115 | RL27A_MOUSE | NQSFC[+57]PTVNLDK | 5.85 | 2.32E-02 | 0.0352 | 8 |
| P01872 | IGHM_MOUSE I | SILEGSDEYLVC[+57]K | 5.08 | 2.54E-02 | 0.0375 | 8 |
| O70194 | EIF3D_MOUSE | YLEVSEPQDIEC[+57]C[+57]GALEYYDK | 6.30 | 2.69E-02 | 0.0392 | 8 |
| P07724 | ALBU_MOUSE S | AADKDTC[+57]FSTEGPNLVTR | 3.02 | 2.80E-02 | 0.0404 | 8 |
| Q8BPN8 | DMXL2_MOUSE | FLLQESQQETTVK | 8.18 | 2.83E-02 | 0.0407 | 8 |
| Q6PIC6 | AT1A3_MOUSE | DFTSEQIDEILQNHTEIVFAR | 3.16 | 3.16E-03 | 0.0143 | 8 |
| Q6PIC6 | AT1A3_MOUSE | M[+16]QVNAEEVVVGDLVEIK | 1.91 | 3.30E-03 | 0.0143 | 8 |
| Q6PIC6 | AT1A3_MOUSE | IATLASGLEVGK | 1.41 | 6.67E-03 | 0.0172 | 8 |
| Q6PIC6 | AT1A3_MOUSE | YQLSIHETEDPNDNR | 1.87 | 7.55E-03 | 0.0180 | 8 |
| Q6PIC6 | AT1A3_MOUSE | LNIPVSQVNPR | 1.68 | 9.22E-03 | 0.0198 | 8 |
| Q6PIC6 | AT1A3_MOUSE | ADIGVAMGIAGSDVSK | 1.46 | 1.41E-02 | 0.0250 | 8 |
| Q6PIC6 | AT1A3_MOUSE | C[+57]IELSSGSVK | 2.91 | 1.60E-02 | 0.0271 | 8 |
| P49722 | PSA2_MOUSE P | LVQIEYALAAVAGGAPSVGIK | 1.47 | 2.21E-02 | 0.0341 | 8 |
| P09405 | NUCL_MOUSE | GFGFVDFNSEEDAK | 3.62 | 1.93E-03 | 0.0143 | 8 |
| P09405 | NUCL_MOUSE | SVSLYYTGEK | 2.52 | 3.43E-03 | 0.0144 | 8 |
| P09405 | NUCL_MOUSE | GLSEDTTEETLK | 3.31 | 6.51E-03 | 0.0171 | 8 |
| P09405 | NUCL_MOUSE | VEGSEPTTPFNLFIGNLNPNK | 4.06 | 7.78E-03 | 0.0182 | 8 |
| P09405 | NUCL_MOUSE | FAISELFAK | 2.88 | 9.86E-03 | 0.0205 | 8 |
| P09405 | NUCL_MOUSE | NLSFNITEDELK | 2.27 | 1.03E-02 | 0.0209 | 8 |
| P52480 | KPYM_MOUSE | LAPITSDPTEAAAVGAVEASFK | 3.51 | 1.84E-02 | 0.0297 | 8 |
| Q60875 | ARHG2_MOUSE | EVEGLKDLLLGPC[+57]VDLPMTSR | 5.88 | 2.46E-02 | 0.0367 | 8 |
| Q04447 | KCRB_MOUSE | NYEFM[+16]WNPHLGYILTC[+57]PSNLGTGLR | 6.62 | 2.26E-03 | 0.0143 | 8 |
| Q04447 | KCRB_MOUSE | LGFSEVELVQMVVDGVK | 2.97 | 2.95E-03 | 0.0143 | 8 |
| Q04447 | KCRB_MOUSE | NYEFMWNPHLGYILTC[+57]PSNLGTGLR | 6.37 | 3.01E-03 | 0.0143 | 8 |
| P08249 | MDHM_MOUSE | LTLYDIAHTPGVAADLSHIETR | 2.18 | 3.07E-03 | 0.0143 | 8 |
| Q04447 | KCRB_MOUSE | FC[+57]TGLTQIETLFK | 1.60 | 8.75E-03 | 0.0192 | 8 |
| P08249 | MDHM_MOUSE | FVFSLVDAMNGK | 2.23 | 9.06E-03 | 0.0196 | 8 |
| P08249 | MDHM_MOUSE | GYLGPEQLPDC[+57]LK | 3.24 | 1.25E-02 | 0.0232 | 8 |
| P08249 | MDHM_MOUSE | ETEC[+57]TYFSTPLLLGK | 1.25 | 1.57E-02 | 0.0268 | 8 |
| P08249 | MDHM_MOUSE | GC[+57]DVVVIPAGVPR | 1.05 | 1.62E-02 | 0.0273 | 8 |
| P08249 | MDHM_MOUSE | VDFPQDQLATLTGR | 1.86 | 1.63E-02 | 0.0274 | 8 |
| E9Q912 | E9Q912_MOUSE | DQEVLLQTGR | 1.23 | 1.71E-02 | 0.0283 | 8 |
| P26039 | TLN1_MOUSE T | AGALQC[+57]SPSDVYTK | 1.01 | 1.72E-02 | 0.0284 | 8 |
| Q91VR5 | DDX1_MOUSE A | DQLSVLDNGVDIVVGTPGR | 1.05 | 1.94E-02 | 0.0308 | 8 |
| P05202 | AATM_MOUSE | ASAELALGENNEVLK | 1.06 | 2.23E-02 | 0.0343 | 8 |
| P06151 | LDHA_MOUSE L | SADTLWGIQK | 1.33 | 3.03E-02 | 0.0429 | 8 |
| Q8BJH1 | ZC21A_MOUSE | HINFC[+57]K | 1.09 | 3.51E-02 | 0.0486 | 8 |

| Protein Accession | Protein Description | Modified Peptide Sequence | co-IP:Ctrl | p value | q value | Sig Peptide Count |
|---|---|---|---|---|---|---|
| P07724 | ALBU_MOUSE | LPC[+57]VEDYLSAILNR | 3.21 | 1.07E-03 | 0.0143 | 8 |
| P17183 | ENOG_MOUSE | NYPVVSIEDPFDQDDWAAWSK | 2.69 | 1.21E-03 | 0.0143 | 8 |
| O08599 | STXB1_MOUSE | AAHVFFTDSC[+57]PDALFNELVK | 2.57 | 1.68E-03 | 0.0143 | 8 |
| P17183 | ENOG_MOUSE | FGANAILGVSLAVC[+57]K | 2.25 | 1.83E-03 | 0.0143 | 8 |
| O08599 | STXB1_MOUSE | DNALLAQLIQDKLDAYK | 2.07 | 2.23E-03 | 0.0143 | 8 |
| O08599 | STXB1_MOUSE | ISEQTYQLSR | 1.47 | 2.25E-03 | 0.0143 | 8 |
| P17183 | ENOG_MOUSE | SGETEDTFIADLVVGLC[+57]TGQIK | 2.60 | 3.40E-03 | 0.0143 | 8 |
| O08599 | STXB1_MOUSE | EVLLDEDDDLWIALR | 5.73 | 4.09E-03 | 0.0148 | 8 |
| P07724 | ALBU_MOUSE | DVFLGTFLYEYSR | 5.45 | 4.13E-03 | 0.0148 | 8 |
| O08599 | STXB1_MOUSE | VLVVDQLSMR | 1.73 | 6.40E-03 | 0.0169 | 8 |
| O08599 | STXB1_MOUSE | DLSQM[+16]LK | 1.77 | 1.09E-02 | 0.0216 | 8 |
| P07724 | ALBU_MOUSE | RHPDYSVSLLLR | 1.81 | 1.24E-02 | 0.0231 | 8 |
| P07724 | ALBU_MOUSE | APQVSTPTLVEAAR | 1.70 | 1.25E-02 | 0.0232 | 8 |
| Q9WTM5 | RUVB2_MOUSE | TQGFLALFSGDTGEIK | 1.12 | 1.76E-02 | 0.0288 | 8 |
| P47963 | RL13_MOUSE | TIGISVDPR | 1.11 | 1.88E-02 | 0.0301 | 8 |
| Q8CHC4 | SYNJ1_MOUSE | TPGPPSSQGSPVDTQPAAQK | 2.31 | 2.14E-02 | 0.0333 | 8 |
| Q8R191 | SNG3_MOUSE | TTPGPGTAQAGDAAR | 1.13 | 2.35E-02 | 0.0356 | 8 |
| Q61879 | MYH10_MOUSE | QEVMISDLEER | 4.83 | 2.50E-02 | 0.0371 | 8 |
| P62242 | RS8_MOUSE | LDVGNFSWGSEC[+57]C[+57]TR | 0.96 | 2.67E-02 | 0.0390 | 8 |
| Q62261 | SPTB2_MOUSE | EIEELQSQAQALSQEGK | 0.83 | 2.80E-02 | 0.0404 | 8 |
| O88456 | CPNS1_MOUSE | M[+58]FLVNSFLK | 1.50 | 3.06E-02 | 0.0432 | 8 |
| P17426 | AP2A1_MOUSE | FFQPTEMAAQDFFQR | 0.91 | 3.16E-02 | 0.0444 | 8 |
| P97351 | RS3A_MOUSE | APAMFNIR | 1.43 | 3.52E-02 | 0.0487 | 8 |
| P52480 | KPYM_MOUSE | KGVNLPGAAVDLPAVSEK | 0.91 | 3.58E-02 | 0.0494 | 8 |
| E9Q557 | DESP_MOUSE | FGDSNTVM[+16]R | 3.39 | 1.70E-02 | 0.0282 | 8 |
| Q9Z2H5 | E41L1_MOUSE | IIITGDEDVDQDQALALAIK | 2.64 | 1.81E-03 | 0.0143 | 7 |
| Q9Z2H5 | E41L1_MOUSE | AETMTVSSLAIR | 2.87 | 2.07E-03 | 0.0143 | 7 |
| Q9Z2H5 | E41L1_MOUSE | DVLTSTYGATAETLSTSTTTHVTK | 2.50 | 2.49E-03 | 0.0143 | 7 |
| Q9Z2H5 | E41L1_MOUSE | SSPWNFAFTVK | 3.50 | 2.57E-03 | 0.0143 | 7 |
| Q6R891 | NEB2_MOUSE | ERPGEQSEVAQLIQQTLEQER | 6.35 | 2.66E-03 | 0.0143 | 7 |
| Q6R891 | NEB2_MOUSE | LEGYWGEAQSLC[+57]QAVDEHLR | 6.22 | 2.70E-03 | 0.0143 | 7 |
| Q80ZK2 | Q80ZK2_MOUSE | SAELGC[+57]TVDEVESLIK | 3.64 | 3.50E-03 | 0.0144 | 7 |
| Q9Z2H5 | E41L1_MOUSE | GQVLFDLVC[+57]EHLNLLEK | 6.47 | 3.75E-03 | 0.0147 | 7 |
| Q80ZK2 | Q80ZK2_MOUSE | AASAGVPYHGEVPVSLAR | 2.01 | 4.94E-03 | 0.0155 | 7 |
| Q9Z2H5 | E41L1_MOUSE | VTLLDASEYEC[+57]EVEK | 3.25 | 7.17E-03 | 0.0177 | 7 |
| Q6R891 | NEB2_MOUSE | ISELEGNLQTLR | 3.64 | 7.88E-03 | 0.0183 | 7 |
| Q80ZK2 | Q80ZK2_MOUSE | ELDDLEQWIQER | 4.76 | 7.96E-03 | 0.0183 | 7 |
| Q9Z2H5 | E41L1_MOUSE | YYLC[+57]LQLR | 2.86 | 8.02E-03 | 0.0184 | 7 |
| P62242 | RS8_MOUSE | NC[+57]IVLIDSTPYR | 5.38 | 9.69E-03 | 0.0204 | 7 |
| Q6R891 | NEB2_MOUSE | LELFPVELEK | 5.06 | 9.74E-03 | 0.0204 | 7 |
| P62242 | RS8_MOUSE | ISSLLEEQFQQGK | 5.26 | 1.05E-02 | 0.0212 | 7 |
| O70194 | EIF3D_MOUSE | IFHTVTTTDDPVIR | 3.80 | 1.12E-02 | 0.0219 | 7 |
| P62242 | RS8_MOUSE | LTPEEEEILNK | 3.72 | 1.15E-02 | 0.0223 | 7 |
| Q80ZK2 | Q80ZK2_MOUSE | DLNAAEALQR | 2.97 | 1.20E-02 | 0.0228 | 7 |
| Q6R891 | NEB2_MOUSE | SAYEAGIQALKPPDAPGPDEAPK | 5.39 | 1.20E-02 | 0.0228 | 7 |
| Q80ZK2 | Q80ZK2_MOUSE | LEQSNVPEGPGSGTGDESSGPR | 4.44 | 1.21E-02 | 0.0229 | 7 |
| O70194 | EIF3D_MOUSE | NMVQFNLQTLPK | 3.29 | 1.21E-02 | 0.0229 | 7 |
| P62242 | RS8_MOUSE | ADGYVLEGK | 3.40 | 1.21E-02 | 0.0229 | 7 |
| O70194 | EIF3D_MOUSE | TQGNVFATDAILATLMSC[+57]TR | 6.60 | 1.34E-02 | 0.0242 | 7 |
| O70194 | EIF3D_MOUSE | TQGNVFATDAILATLM[+16]SC[+57]TR | 6.28 | 1.39E-02 | 0.0248 | 7 |
| Q6R891 | NEB2_MOUSE | VFQPPPPPPAPSGDGATEK | 1.70 | 1.50E-02 | 0.0261 | 7 |
| Q8BPN8 | DMXL2_MOUSE | IPVAFPSGDANSLSK | 2.77 | 1.77E-02 | 0.0289 | 7 |
| P39053 | DYN1_MOUSE | ALLQM[+16]VQQFAVDFEK | 5.76 | 2.16E-02 | 0.0335 | 7 |
| Q9R1R2 | TRIM3_MOUSE | SVLNLGALLTTSATAHETVATGEGLR | 3.18 | 2.25E-02 | 0.0345 | 7 |
| Q8R4U7 | LUZP1_MOUSE | VGNSGDAPELSPR | 3.16 | 2.29E-02 | 0.0349 | 7 |
| P20357 | MTAP2_MOUSE | GVVESVVTIEDDFITVVQTTTDEGESGSHSVR | 2.86 | 2.39E-02 | 0.0360 | 7 |
| Q9WV92 | E41L3_MOUSE | VLQETILVEER | 6.11 | 2.65E-02 | 0.0388 | 7 |
| Q6PHZ2 | KCC2D_MOUSE | DLINKM[+16]LTINPAK | 7.42 | 2.67E-02 | 0.0390 | 7 |
| Q99PU8 | DHX30_MOUSE | AVDEAVILLQEIGVLDQR | 4.10 | 2.68E-02 | 0.0391 | 7 |
| P17426 | AP2A1_MOUSE | DFLTPPLLSVR | 3.72 | 3.43E-02 | 0.0477 | 7 |
| P18760 | COF1_MOUSE | NIILEEGKEILVGDVGQTVDDPYTTFVK | 3.32 | 6.86E-04 | 0.0143 | 7 |
| P61979 | HNRPK_MOUSE | ILSISADIETIGEILK | 2.88 | 9.16E-04 | 0.0143 | 7 |

| Protein Accession | Protein Description | Modified Peptide Sequence | co-IP:Ctrl | p value | q value | Sig Peptide Count |
|---|---|---|---|---|---|---|
| Q9EQH3 | VPS35_MOUSE | IPVDTYNNILTVLK | 1.61 | 1.71E-03 | 0.0143 | 7 |
| Q9EQH3 | VPS35_MOUSE | LFDIFSQQVATVIQSR | 2.65 | 1.85E-03 | 0.0143 | 7 |
| P61979 | HNRPK_MOUSE | IITITGTQDQIQNAQYLLQNSVK | 1.82 | 1.85E-03 | 0.0143 | 7 |
| P18760 | COF1_MOUSE | KEDLVFIFWAPENAPLK | 2.69 | 1.88E-03 | 0.0143 | 7 |
| Q9EQH3 | VPS35_MOUSE | VADLYELVQYAGNIIPR | 4.37 | 2.04E-03 | 0.0143 | 7 |
| P18760 | COF1_MOUSE | YALYDATYETK | 1.19 | 2.25E-03 | 0.0143 | 7 |
| P18760 | COF1_MOUSE | HELQANC[+57]YEEVKDR | 1.70 | 2.31E-03 | 0.0143 | 7 |
| P61979 | HNRPK_MOUSE | GSDFDC[+57]ELR | 1.73 | 2.78E-03 | 0.0143 | 7 |
| P61979 | HNRPK_MOUSE | LLIHQSLAGGIIGVK | 2.04 | 2.79E-03 | 0.0143 | 7 |
| P61979 | HNRPK_MOUSE | IILDLISESPIK | 1.67 | 3.02E-03 | 0.0143 | 7 |
| P18760 | COF1_MOUSE | LGGSAVISLEGKPL | 2.13 | 4.34E-03 | 0.0150 | 7 |
| Q9EQH3 | VPS35_MOUSE | QIVLTGILEQVVNC[+57]R | 2.32 | 4.97E-03 | 0.0155 | 7 |
| P18760 | COF1_MOUSE | AVLFC[+57]LSEDKK | 1.07 | 5.05E-03 | 0.0155 | 7 |
| Q7TPR4 | ACTN1_MOUSE | VGWEQLLTTIAR | 4.27 | 5.71E-03 | 0.0161 | 7 |
| Q9EQH3 | VPS35_MOUSE | LSQLEGVNVER | 2.17 | 6.91E-03 | 0.0174 | 7 |
| Q7TPR4 | ACTN1_MOUSE | LLETIDQLYLEYAK | 6.64 | 7.65E-03 | 0.0181 | 7 |
| Q7TPR4 | ACTN1_MOUSE | LASDLLEWIR | 2.85 | 1.06E-02 | 0.0213 | 7 |
| Q7TPR4 | ACTN1_MOUSE | IDQLEC[+57]DHQLIQEALIFDNK | 4.39 | 1.06E-02 | 0.0213 | 7 |
| Q9EQH3 | VPS35_MOUSE | IREDLPNLESSEETEQINK | 2.07 | 1.15E-02 | 0.0223 | 7 |
| Q7TPR4 | ACTN1_MOUSE | TINEVENQILTR | 3.28 | 1.16E-02 | 0.0224 | 7 |
| P18760 | COF1_MOUSE | EILVGDVGQTVDDPYTTFVK | 2.03 | 1.46E-02 | 0.0256 | 7 |
| P61979 | HNRPK_MOUSE | GSYGDLGGPIITTQVTIPK | 1.87 | 1.47E-02 | 0.0257 | 7 |
| Q8BP47 | SYNC_MOUSE | FLSWILNR | 4.20 | 2.25E-02 | 0.0345 | 7 |
| P56480 | ATPB_MOUSE | SLQDIIAILGMDELSEEDKLTVSR | 3.20 | 2.42E-02 | 0.0364 | 7 |
| P39053 | DYN1_MOUSE | NLVDSYMAIVNK | 1.70 | 2.64E-02 | 0.0387 | 7 |
| Q8R5C5 | ACTY_MOUSE | VQYTLPDGSTLDVGPAR | 0.94 | 3.48E-02 | 0.0483 | 7 |
| P14869 | RLA0_MOUSE | TSFFQALGITTK | 2.31 | 1.09E-03 | 0.0143 | 7 |
| Q01853 | TERA_MOUSE | AIANEC[+57]QANFISIK | 1.92 | 1.60E-03 | 0.0143 | 7 |
| Q8C8R3 | ANK2_MOUSE | YGSLDVAK | 3.21 | 2.15E-03 | 0.0143 | 7 |
| Q01853 | TERA_MOUSE | NAPAIIFIDELDAIAPK | 2.11 | 2.32E-03 | 0.0143 | 7 |
| P14869 | RLA0_MOUSE | GNVGFVFTK | 2.05 | 2.57E-03 | 0.0143 | 7 |
| P14869 | RLA0_MOUSE | AGAIAPC[+57]EVTVPAQNTGLGPEK | 1.76 | 2.65E-03 | 0.0143 | 7 |
| Q01853 | TERA_MOUSE | LDQLIYIPLPDEK | 1.60 | 2.81E-03 | 0.0143 | 7 |
| P14869 | RLA0_MOUSE | GTIEILSDVQLIK | 2.15 | 2.88E-03 | 0.0143 | 7 |
| Q01853 | TERA_MOUSE | DVDLEFLAK | 1.29 | 4.19E-03 | 0.0149 | 7 |
| Q8C8R3 | ANK2_MOUSE | EGHVGLVQELLGR | 3.32 | 4.83E-03 | 0.0154 | 7 |
| Q8C8R3 | ANK2_MOUSE | AGEEEPGEPFEIVER | 1.52 | 5.82E-03 | 0.0162 | 7 |
| P14869 | RLA0_MOUSE | IIQLLDDYPK | 1.61 | 6.54E-03 | 0.0171 | 7 |
| Q01853 | TERA_MOUSE | ELQELVQYPVEHPDK | 1.30 | 7.98E-03 | 0.0183 | 7 |
| Q8C8R3 | ANK2_MOUSE | SGHDQVVELLLER | 5.12 | 8.03E-03 | 0.0184 | 7 |
| P14869 | RLA0_MOUSE | AFLADPSAFAAAAPAAAATTAAPAAAAAPA | 2.26 | 1.01E-02 | 0.0207 | 7 |
| Q8C8R3 | ANK2_MOUSE | VVTEEVTTTTTTITEK | 5.50 | 1.09E-02 | 0.0216 | 7 |
| Q8C8R3 | ANK2_MOUSE | GLVHQAIC[+57]NLNITLPIYAK | 5.66 | 1.09E-02 | 0.0216 | 7 |
| Q8C8R3 | ANK2_MOUSE | VALLLLEK | 3.96 | 1.14E-02 | 0.0222 | 7 |
| Q01853 | TERA_MOUSE | AVANETGAFFFLINGPEIMSK | 4.67 | 1.57E-02 | 0.0268 | 7 |
| P19253 | RL13A_MOUSE | YQAVTATLEEK | 3.47 | 2.16E-02 | 0.0335 | 7 |
| P17183 | ENOG_MOUSE | VNQIGSVTEAIQAC[+57]K | 2.38 | 3.58E-02 | 0.0494 | 7 |
| Q61316 | HSP74_MOUSE | SNLAYDIVQLPTGLTGIK | 2.32 | 1.21E-04 | 0.0143 | 7 |
| Q61316 | HSP74_MOUSE | VLATAFDTTLGGR | 2.23 | 3.92E-03 | 0.0147 | 7 |
| Q61316 | HSP74_MOUSE | AFSDPFVEAEK | 1.71 | 4.95E-03 | 0.0155 | 7 |
| Q61316 | HSP74_MOUSE | SVMDATQIAGLNC[+57]LR | 2.55 | 8.17E-03 | 0.0185 | 7 |
| Q61316 | HSP74_MOUSE | LMNETTAVALAYGIYK | 2.04 | 1.32E-02 | 0.0240 | 7 |
| Q80UX7 | Q80UX7_MOUSE | NEAEVINMSEELAQLEGILK | 1.71 | 2.57E-02 | 0.0379 | 7 |
| P28652 | KCC2B_MOUSE | QTTAPATM[+16]STAASGTTMGLVEQAK | 2.63 | 3.27E-02 | 0.0457 | 7 |
| Q9Z0E0 | NCDN_MOUSE | ILGAWLAEETSSLR | 2.45 | 1.44E-03 | 0.0143 | 7 |
| Q9Z0E0 | NCDN_MOUSE | LQAGEETASHYR | 6.76 | 2.44E-03 | 0.0143 | 7 |
| Q9Z0E0 | NCDN_MOUSE | LLLAANVATLGLLMAR | 2.35 | 3.02E-03 | 0.0143 | 7 |
| Q9Z0E0 | NCDN_MOUSE | FELC[+57]QLLPLFLPPTTVPPEC[+57]HR | 6.76 | 1.17E-02 | 0.0225 | 7 |
| Q9WUM4 | COR1C_MOUSE | QLALWNPK | 1.05 | 1.75E-02 | 0.0287 | 7 |
| P09411 | PGK1_MOUSE | VLPGVDALSNV | 1.17 | 2.77E-02 | 0.0400 | 7 |
| Q60900 | ELAV3_MOUSE | TGQALLTHLYQSSAR | 1.07 | 2.85E-02 | 0.0409 | 7 |
| P17710 | HXK1_MOUSE | GAAMVTAVAYR | 4.55 | 1.55E-03 | 0.0143 | 7 |

| Protein Accession | Protein Description | Modified Peptide Sequence | co-IP:Ctrl | p value | q value | Sig Peptide Count |
|---|---|---|---|---|---|---|
| P23116 | EIF3A_MOUSE | QPALDVLYDVMK | 4.55 | 2.74E-03 | 0.0143 | 7 |
| P63101 | 1433Z_MOUSE | IETELRDIC[+57]NDVLSLLEK | 7.49 | 3.28E-03 | 0.0143 | 7 |
| P63101 | 1433Z_MOUSE | TAFDEAIAELDTLSEESYK | 2.41 | 4.75E-03 | 0.0154 | 7 |
| P23116 | EIF3A_MOUSE | NQLTAMSSVLAK | 4.06 | 7.25E-03 | 0.0177 | 7 |
| P63101 | 1433Z_MOUSE | DIC[+57]NDVLSLLEK | 2.11 | 7.39E-03 | 0.0179 | 7 |
| P17710 | HXK1_MOUSE I | GDFIALDLGGSSFR | 2.22 | 9.93E-03 | 0.0206 | 7 |
| P17710 | HXK1_MOUSE I | LSDEILIDILTR | 4.12 | 1.05E-02 | 0.0212 | 7 |
| P17710 | HXK1_MOUSE I | IDEAVLITWTK | 2.31 | 1.10E-02 | 0.0217 | 7 |
| P23116 | EIF3A_MOUSE | LTSLVPFVDAFQLER | 6.63 | 1.13E-02 | 0.0221 | 7 |
| P63101 | 1433Z_MOUSE | GIVDQSQQAYQEAFEISKK | 1.35 | 1.26E-02 | 0.0233 | 7 |
| P23116 | EIF3A_MOUSE | FSVLQYVVPEVK | 6.89 | 1.28E-02 | 0.0235 | 7 |
| P17710 | HXK1_MOUSE I | MISGMYLGEIVR | 2.10 | 1.35E-02 | 0.0243 | 7 |
| P63101 | 1433Z_MOUSE | GIVDQSQQAYQEAFEISK | 1.36 | 1.40E-02 | 0.0249 | 7 |
| P63101 | 1433Z_MOUSE | YLAEVAAGDDK | 0.62 | 1.44E-02 | 0.0254 | 7 |
| P23116 | EIF3A_MOUSE | VLLATLSIPITPER | 6.83 | 1.52E-02 | 0.0263 | 7 |
| P23116 | EIF3A_MOUSE | DIDIEDLEELDPDFIMAK | 7.13 | 1.59E-02 | 0.0270 | 7 |
| P17710 | HXK1_MOUSE I | SANLVAATLGAILNR | 2.13 | 1.66E-02 | 0.0277 | 7 |
| Q8BP67 | RL24_MOUSE 6 | VFQFLNAK | 4.75 | 1.97E-02 | 0.0311 | 7 |
| P01837 | IGKC_MOUSE I | ADAAPTVSIFPPSSEQLTSGGASVVC[+57]FLNNFYPK | 2.41 | 2.39E-02 | 0.0360 | 7 |
| P10126 | EF1A1_MOUSE | NMITGTSQADC[+57]AVLIVAAGVGEFEAGISK | 1.14 | 2.95E-02 | 0.0420 | 7 |
| P16330 | CN37_MOUSE 2 | AGQVFLEELGNHK | 2.17 | 9.12E-04 | 0.0143 | 6 |
| P47753 | CAZA1_MOUSE | EGAAHAFAQYNMDQFTPVK | 3.37 | 1.11E-03 | 0.0143 | 6 |
| P47753 | CAZA1_MOUSE | EASDPQPEDVDGGLK | 4.86 | 1.17E-03 | 0.0143 | 6 |
| P47753 | CAZA1_MOUSE | IEGYDDQVLITEHGDLGNSR | 5.00 | 2.14E-03 | 0.0143 | 6 |
| P12382 | PFKAL_MOUSE | NEWGSLLEELVK | 2.72 | 2.39E-03 | 0.0143 | 6 |
| P16330 | CN37_MOUSE 2 | AAGAEEYAQQEVVK | 2.46 | 2.45E-03 | 0.0143 | 6 |
| P47753 | CAZA1_MOUSE | FTITPPSAQVVGVLK | 5.57 | 2.51E-03 | 0.0143 | 6 |
| P12382 | PFKAL_MOUSE | VFANAPDSAC[+57]VIGLR | 2.29 | 3.00E-03 | 0.0143 | 6 |
| P16330 | CN37_MOUSE 2 | AHVTLGC[+57]AADVQPVQTGLDLLDILQQVK | 8.44 | 3.03E-03 | 0.0143 | 6 |
| P47753 | CAZA1_MOUSE | FITHAPPGEFNEVFNDVR | 4.50 | 3.33E-03 | 0.0143 | 6 |
| P16330 | CN37_MOUSE 2 | GGSQGEAVGELPR | 2.43 | 5.01E-03 | 0.0155 | 6 |
| P16330 | CN37_MOUSE 2 | LSISALFVTPK | 2.50 | 5.36E-03 | 0.0158 | 6 |
| P60229 | EIF3E_MOUSE | NALSSLWGK | 3.91 | 8.53E-03 | 0.0189 | 6 |
| P12382 | PFKAL_MOUSE | AIGVLTSGGDAQGMNAAVR | 2.27 | 9.98E-03 | 0.0206 | 6 |
| P47753 | CAZA1_MOUSE | ESC[+57]DSALR | 4.98 | 1.01E-02 | 0.0207 | 6 |
| P60229 | EIF3E_MOUSE | LKETIDNNSVSSPLQSLQQR | 4.76 | 1.17E-02 | 0.0225 | 6 |
| P60229 | EIF3E_MOUSE | MLFDYLADK | 4.65 | 1.20E-02 | 0.0228 | 6 |
| P12382 | PFKAL_MOUSE | AIGVLTSGGDAQGM[+16]NAAVR | 2.40 | 1.22E-02 | 0.0230 | 6 |
| P12382 | PFKAL_MOUSE | LAAAYNLLQHGITNLC[+57]VIGGDGSLTGANIFR | 7.91 | 1.28E-02 | 0.0235 | 6 |
| P60229 | EIF3E_MOUSE | LASEILMQNWDAAMEDLTR | 7.35 | 1.33E-02 | 0.0241 | 6 |
| P60229 | EIF3E_MOUSE | LFIFETFC[+57]R | 5.53 | 1.64E-02 | 0.0275 | 6 |
| P48036 | ANXA5_MOUSE | ETSGNLEQLLLAVVK | 7.26 | 1.87E-02 | 0.0300 | 6 |
| P41105 | RL28_MOUSE 6 | TVGVEPAADGK | 8.46 | 2.19E-02 | 0.0338 | 6 |
| Q6URW6 | MYH14_MOUSE | HEVPPHVYAVTEGAYR | 1.75 | 2.47E-02 | 0.0368 | 6 |
| Q60605 | MYL6_MOUSE I | ILYSQC[+57]GDVMR | 5.16 | 6.53E-04 | 0.0143 | 6 |
| P48722 | HS74L_MOUSE | EDINSIEIVGGATR | 2.60 | 7.42E-04 | 0.0143 | 6 |
| Q60605 | MYL6_MOUSE I | NKDQGTYEDYVEGLR | 5.25 | 1.36E-03 | 0.0143 | 6 |
| Q9R111 | GUAD_MOUSE | FTLSC[+57]TETLMSELGNIAK | 2.01 | 1.83E-03 | 0.0143 | 6 |
| Q9R111 | GUAD_MOUSE | GASIAHC[+57]PNSNLSLSSGLLNVLEVLK | 3.88 | 1.90E-03 | 0.0143 | 6 |
| Q60605 | MYL6_MOUSE I | ILYSQC[+57]GDVM[+16]R | 5.86 | 1.91E-03 | 0.0143 | 6 |
| Q60605 | MYL6_MOUSE I | VLDFEHFLPMLQTVAK | 5.69 | 1.94E-03 | 0.0143 | 6 |
| Q60605 | MYL6_MOUSE I | EAFQLFDR | 4.91 | 2.24E-03 | 0.0143 | 6 |
| P48722 | HS74L_MOUSE | EFSITDLVPYSVTLR | 3.60 | 2.43E-03 | 0.0143 | 6 |
| Q9QY94 | GLNA_ACOCA Gl | RLTGFHETSNINDFSAGVANR | 2.03 | 3.24E-03 | 0.0143 | 6 |
| Q9QY94 | GLNA_ACOCA Gl | VQAM[+16]YIWVDGTGEGLR | 0.69 | 3.31E-03 | 0.0143 | 6 |
| Q9R111 | GUAD_MOUSE | LATLGGSQALGLDSEIGNFEVGK | 1.62 | 3.60E-03 | 0.0145 | 6 |
| Q9R111 | GUAD_MOUSE | AVMVSNVLLINK | 1.89 | 3.81E-03 | 0.0147 | 6 |
| Q60605 | MYL6_MOUSE I | HVLVTLGEK | 3.82 | 4.06E-03 | 0.0148 | 6 |
| O54833 | CSK22_MOUSE | QLYQILTDFDIR | 6.97 | 4.30E-03 | 0.0150 | 6 |
| P48722 | HS74L_MOUSE | SIDLPIQSSLYR | 3.20 | 5.23E-03 | 0.0157 | 6 |
| Q9QY94 | GLNA_ACOCA Gl | VQAMYIWVDGTGEGLR | 2.84 | 5.64E-03 | 0.0160 | 6 |
| O54833 | CSK22_MOUSE | LIDWGLAEFYHPAQEYNVR | 5.09 | 6.23E-03 | 0.0167 | 6 |

| Protein Accession | Protein Description | Modified Peptide Sequence | co-IP:Ctrl | p value | q value | Sig Peptide Count |
|---|---|---|---|---|---|---|
| O54833 | CSK22_MOUSE | VLGTDELYGYLK | 3.63 | 7.41E-03 | 0.0179 | 6 |
| P48722 | HS74L_MOUSE | VLATTFDPYLGGR | 2.37 | 7.99E-03 | 0.0183 | 6 |
| O54833 | CSK22_MOUSE | TPALVFEYINNTDFK | 6.08 | 8.31E-03 | 0.0187 | 6 |
| Q9QY94 | GLNA_ACOCA Gl | LTGFHETSNINDFSAGVANR | 1.43 | 8.52E-03 | 0.0189 | 6 |
| P25444 | RS2_MOUSE 4 | GC[+57]TATLGNFAK | 2.82 | 8.82E-03 | 0.0193 | 6 |
| P25444 | RS2_MOUSE 4 | C[+57]GSVLVR | 3.73 | 9.22E-03 | 0.0198 | 6 |
| P48722 | HS74L_MOUSE | LSLTQDPVVK | 1.37 | 1.01E-02 | 0.0207 | 6 |
| P25444 | RS2_MOUSE 4 | GTGIVSAPVPK | 3.12 | 1.01E-02 | 0.0207 | 6 |
| O54833 | CSK22_MOUSE | HLVSPEALDLLDK | 3.33 | 1.07E-02 | 0.0214 | 6 |
| Q9QY94 | GLNA_ACOCA Gl | ITGTNAEVMPAQWEFQIGPC[+57]EGIR | 2.43 | 1.12E-02 | 0.0219 | 6 |
| P25444 | RS2_MOUSE 4 | TYSYLTPDLWK | 2.42 | 1.12E-02 | 0.0219 | 6 |
| O54833 | CSK22_MOUSE | VYAEVNSLR | 2.25 | 1.38E-02 | 0.0247 | 6 |
| Q9R111 | GUAD_MOUSE | STDVAEEVYTR | 1.79 | 1.42E-02 | 0.0251 | 6 |
| Q8R1B4 | EIF3C_MOUSE | ELLGQGLLLR | 5.76 | 1.46E-02 | 0.0256 | 6 |
| Q8R1B4 | EIF3C_MOUSE | GC[+57]ILTLVER | 4.82 | 1.47E-02 | 0.0257 | 6 |
| Q8R1B4 | EIF3C_MOUSE | DAHNALLDIQSSGR | 3.73 | 1.58E-02 | 0.0269 | 6 |
| P48722 | HS74L_MOUSE | NFDEALVDYFC[+57]DEFK | 5.31 | 1.65E-02 | 0.0276 | 6 |
| Q99104 | MYO5A_MOUSE | WTYQEFFSR | 1.14 | 1.72E-02 | 0.0284 | 6 |
| P20357 | MTAP2_MOUSE | ESSKDEEPLKDK | 4.46 | 2.13E-02 | 0.0331 | 6 |
| Q61879 | MYH10_MOUSE | LQQELDDLTVDLDHQR | 5.08 | 2.13E-02 | 0.0331 | 6 |
| P07724 | ALBU_MOUSE S | RPC[+57]FSALTVDETYVPK | 1.07 | 2.35E-02 | 0.0356 | 6 |
| P62259 | 1433E_MOUSE | VAGM[+16]DVELTVEER | 3.17 | 2.74E-02 | 0.0397 | 6 |
| P68404 | KPCB_MOUSE | LSVEIWDWDLTSR | 5.57 | 2.76E-02 | 0.0399 | 6 |
| Q922U2 | K2C5_MOUSE K | VDALMDEINFMK | 7.68 | 3.45E-02 | 0.0479 | 6 |
| P68510 | 1433F_MOUSE | ELETVC[+57]NDVLALLDK | 2.49 | 1.72E-03 | 0.0143 | 6 |
| Q9R1R2 | TRIM3_MOUSE | GRLPQLSAAIALVGGISQQLQER | 7.71 | 2.56E-03 | 0.0143 | 6 |
| Q8C1B7 | SEP11_MOUSE | LTIVDTVGFGDQINK | 1.63 | 4.48E-03 | 0.0151 | 6 |
| Q8C1B7 | SEP11_MOUSE | ELEEEVSNFQK | 3.35 | 4.97E-03 | 0.0155 | 6 |
| Q9R1R2 | TRIM3_MOUSE | LPQLSAAIALVGGISQQLQER | 6.41 | 5.24E-03 | 0.0157 | 6 |
| Q9R1R2 | TRIM3_MOUSE | TEGDLLLSVLLYGQPVR | 4.92 | 6.35E-03 | 0.0169 | 6 |
| Q8C1B7 | SEP11_MOUSE | STSQGFC[+57]FNILC[+57]VGETGIGK | 2.22 | 6.55E-03 | 0.0171 | 6 |
| P68510 | 1433F_MOUSE | IEKELETVC[+57]NDVLALLDK | 7.29 | 6.96E-03 | 0.0174 | 6 |
| P68510 | 1433F_MOUSE | QAFDDAIAELDTLNEDSYK | 1.75 | 8.87E-03 | 0.0194 | 6 |
| Q8C1B7 | SEP11_MOUSE | STLMDTLFNTK | 1.31 | 1.07E-02 | 0.0214 | 6 |
| Q9R1R2 | TRIM3_MOUSE | LGSAPEVLLVR | 1.17 | 1.24E-02 | 0.0231 | 6 |
| P68510 | 1433F_MOUSE | NC[+57]NDFQYESK | 1.25 | 1.34E-02 | 0.0242 | 6 |
| Q9R1R2 | TRIM3_MOUSE | QFLVC[+57]SIC[+57]LDR | 4.24 | 1.55E-02 | 0.0266 | 6 |
| Q99104 | MYO5A_MOUSE | QETDQLVSNLK | 0.92 | 1.78E-02 | 0.0290 | 6 |
| Q6PHZ2 | KCC2D_MOUSE | IPTGQEYAAK | 2.92 | 2.24E-02 | 0.0344 | 6 |
| Q9DBR7 | MYPT1_MOUSE | VGQTAFDVADEDILGYLEELQK | 1.00 | 2.26E-02 | 0.0346 | 6 |
| P16330 | CN37_MOUSE 2 | HFISGDEPK | 1.85 | 2.47E-02 | 0.0368 | 6 |
| P16858 | G3P_MOUSE G | IVSNASC[+57]TTNC[+57]LAPLAK | 3.09 | 2.98E-02 | 0.0424 | 6 |
| E9Q912 | E9Q912_MOUSE | M[+16]LIDAQAEAAEQLGK | 4.27 | 2.69E-03 | 0.0143 | 6 |
| E9Q912 | E9Q912_MOUSE | IPC[+57]VDAGLISPLVQLLNSK | 2.81 | 2.91E-03 | 0.0143 | 6 |
| E9Q912 | E9Q912_MOUSE | TEGSLEGC[+57]LDC[+57]LLQALAQNNAETSEK | 8.40 | 3.06E-03 | 0.0143 | 6 |
| E9Q912 | E9Q912_MOUSE | EVQDLAFLDVVSK | 1.16 | 9.54E-03 | 0.0202 | 6 |
| Q9CVB6 | ARPC2_MOUSE | DTDAAVGDNIGYITFVLFPR | 1.47 | 1.71E-02 | 0.0283 | 6 |
| Q61171 | PRDX2_MOUSE | QITVNDLPVGR | 0.87 | 2.37E-02 | 0.0358 | 6 |
| Q8CHC4 | SYNJ1_MOUSE | VLDAYGLLGVLR | 2.45 | 7.59E-04 | 0.0143 | 6 |
| Q8CHC4 | SYNJ1_MOUSE | NQTLTDWLLDAPK | 2.09 | 8.05E-04 | 0.0143 | 6 |
| P62259 | 1433E_MOUSE | VAGMDVELTVEER | 2.63 | 9.89E-04 | 0.0143 | 6 |
| P05202 | AATM_MOUSE | HFIEQGINVC[+57]LC[+57]QSYAK | 2.75 | 1.65E-03 | 0.0143 | 6 |
| P05202 | AATM_MOUSE | DAGMQLQGYR | 1.50 | 1.81E-03 | 0.0143 | 6 |
| Q6ZQ38 | CAND1_MOUSE | ISGSILNELIGLVR | 4.48 | 2.03E-03 | 0.0143 | 6 |
| Q6ZQ38 | CAND1_MOUSE | IDLRPVLGEGVPILASFLR | 9.18 | 3.25E-03 | 0.0143 | 6 |
| Q8CHC4 | SYNJ1_MOUSE | QEAIDVLLLGNTLNSDLADK | 2.03 | 4.83E-03 | 0.0154 | 6 |
| Q8CHC4 | SYNJ1_MOUSE | ALLTTGSLR | 1.75 | 5.22E-03 | 0.0156 | 6 |
| P05202 | AATM_MOUSE | ISVAGVTSGNVGYLAHAIHQVTK | 2.02 | 7.19E-03 | 0.0177 | 6 |
| P62259 | 1433E_MOUSE | M[+58]DDREDLVYQAK | 1.02 | 7.57E-03 | 0.0180 | 6 |
| P62259 | 1433E_MOUSE | LIC[+57]C[+57]DILDVLDK | 1.84 | 8.72E-03 | 0.0191 | 6 |
| P05202 | AATM_MOUSE | TC[+57]GFDFSGALEDISK | 1.49 | 1.24E-02 | 0.0231 | 6 |
| P61358 | RL27_MOUSE 6 | VVLVLAGR | 1.64 | 1.71E-02 | 0.0283 | 6 |

| Protein Accession | Protein Description | Modified Peptide Sequence | co-IP:Ctrl | p value | q value | Sig Peptide Count |
|---|---|---|---|---|---|---|
| P01837 | IGKC_MOUSE | QNGVLNSWTDQDSK | 1.31 | 2.03E-02 | 0.0319 | 6 |
| P25444 | RS2_MOUSE | SLEEIYLFSLPIK | 1.11 | 2.14E-02 | 0.0333 | 6 |
| Q71LX4 | TLN2_MOUSE | TYGVSFFLVK | 1.28 | 2.23E-02 | 0.0343 | 6 |
| P68510 | 1433F_MOUSE | YLAEVASGEK | 1.26 | 2.27E-02 | 0.0347 | 6 |
| P68033 | ACTC_MOUSE | AGFAGDDAPR | 2.69 | 2.28E-02 | 0.0348 | 6 |
| P09405 | NUCL_MOUSE | GYAFIEFASFEDAK | 1.23 | 2.46E-02 | 0.0367 | 6 |
| Q9WUM4 | COR1C_MOUSE | KSDLFQDDLYPDTAGPEAALEAEEWFEGK | 1.56 | 2.72E-02 | 0.0395 | 6 |
| Q7TMM9 | TBB2A_MOUSE | TAVC[+57]DIPPR | 0.81 | 2.74E-02 | 0.0397 | 6 |
| Q7TSJ2 | MAP6_MOUSE | TTEGPSATKPDDKEQSK | 1.12 | 3.10E-02 | 0.0437 | 6 |
| P40142 | TKT_MOUSE | TSRPENAIIYSNNEDFQVGQAK | 1.55 | 3.18E-02 | 0.0446 | 6 |
| P50396 | GDIA_MOUSE | FDLGQDVIDFTGHALALYR | 7.95 | 3.63E-04 | 0.0143 | 6 |
| P06151 | LDHA_MOUSE | DLADELALVDVMEDK | 2.22 | 6.07E-04 | 0.0143 | 6 |
| P11499 | HS90B_MOUSE | HSQFIGYPITLYLEK | 2.37 | 6.45E-04 | 0.0143 | 6 |
| Q61598 | GDIB_MOUSE | FDLGQDVIDFTGHSLALYR | 6.63 | 1.82E-03 | 0.0143 | 6 |
| P06745 | G6PI_MOUSE | KIEPELEGSSAVTSHDSSTNGLISFIK | 2.04 | 2.11E-03 | 0.0143 | 6 |
| P06745 | G6PI_MOUSE | TLASLSPETSLFIIASK | 1.65 | 2.27E-03 | 0.0143 | 6 |
| P06745 | G6PI_MOUSE | ILLANFLAQTEALMK | 3.29 | 2.64E-03 | 0.0143 | 6 |
| P06745 | G6PI_MOUSE | ELFEADPER | 1.49 | 3.11E-03 | 0.0143 | 6 |
| P06745 | G6PI_MOUSE | SITDIINIGIGGSDLGPLMVTEALKPYSK | 8.88 | 3.65E-03 | 0.0146 | 6 |
| P06151 | LDHA_MOUSE | DYC[+57]VTANSK | 3.31 | 5.07E-03 | 0.0155 | 6 |
| P06151 | LDHA_MOUSE | LLIVSNPVDILTYVAWK | 4.74 | 5.82E-03 | 0.0162 | 6 |
| P50396 | GDIA_MOUSE | MLLYTEVTR | 1.40 | 6.35E-03 | 0.0169 | 6 |
| P11499 | HS90B_MOUSE | NPDDITQEEYGEFYK | 1.32 | 6.51E-03 | 0.0171 | 6 |
| Q61598 | GDIB_MOUSE | VPSTEAEALASSLMGLFEK | 3.83 | 7.58E-03 | 0.0180 | 6 |
| P11499 | HS90B_MOUSE | APFDLFENK | 1.74 | 8.23E-03 | 0.0186 | 6 |
| P11499 | HS90B_MOUSE | LVSSPC[+57]C[+57]IVTSTYGWTANMER | 1.96 | 9.47E-03 | 0.0202 | 6 |
| P50396 | GDIA_MOUSE | FLVFVANFDENDPK | 1.43 | 9.53E-03 | 0.0202 | 6 |
| P06151 | LDHA_MOUSE | VIGSGC[+57]NLDSAR | 1.05 | 1.02E-02 | 0.0208 | 6 |
| P50396 | GDIA_MOUSE | FQILEGPPESMGR | 1.34 | 1.07E-02 | 0.0214 | 6 |
| P50396 | GDIA_MOUSE | KFDLGQDVIDFTGHALALYR | 5.18 | 1.20E-02 | 0.0228 | 6 |
| P06151 | LDHA_MOUSE | DQLIVNLLKEEQAPQNK | 2.89 | 1.34E-02 | 0.0242 | 6 |
| Q61598 | GDIB_MOUSE | QLIC[+57]DPSYVK | 4.42 | 1.50E-02 | 0.0261 | 6 |
| P50396 | GDIA_MOUSE | VPSTETEALASNLMGMFEK | 2.69 | 1.53E-02 | 0.0264 | 6 |
| Q8BFR5 | EFTU_MOUSE | LLDAVDTYIPVPTR | 6.00 | 1.87E-02 | 0.0300 | 6 |
| Q921I1 | TRFE_MOUSE | AVLTSQETLFGGSDC[+57]TGNFC[+57]LFK | 1.38 | 2.29E-02 | 0.0349 | 6 |
| P46460 | NSF_MOUSE | THPSVVPGC[+57]IAFSLPQR | 1.30 | 2.43E-02 | 0.0364 | 6 |
| Q9DB20 | ATPO_MOUSE | VSLAVLNPYIK | 2.60 | 2.49E-02 | 0.0370 | 6 |
| Q9CQV8 | 1433B_MOUSE | TAFDEAIAELDTLNEESYK | 2.13 | 2.66E-02 | 0.0389 | 6 |
| O88935 | SYN1_MOUSE | VLLVIDEPHTDWAK | 1.24 | 2.90E-02 | 0.0414 | 6 |
| Q99104 | MYO5A_MOUSE | SHENEAEALRGEIQSLKEENNR | 1.62 | 3.03E-02 | 0.0429 | 6 |
| P05201 | AATC_MOUSE | TPGTWSHITEQIGMFSFTGLNPK | 3.92 | 1.12E-03 | 0.0143 | 6 |
| P05201 | AATC_MOUSE | IANDNSLNHEYLPILGLAEFR | 2.30 | 3.86E-03 | 0.0147 | 6 |
| P05201 | AATC_MOUSE | FLFPFFDSAYQGFASGDLEK | 7.27 | 5.01E-03 | 0.0155 | 6 |
| P05201 | AATC_MOUSE | YFVSEGFELFC[+57]AQSFSK | 4.34 | 5.18E-03 | 0.0156 | 6 |
| P05201 | AATC_MOUSE | INMC[+57]GLTTK | 1.79 | 5.90E-03 | 0.0163 | 6 |
| Q68FD5 | CLH1_MOUSE | AFMTADLPNELIELLEK | 1.29 | 2.49E-02 | 0.0370 | 6 |
| P07901 | HS90A_MOUSE | HGLEVIYMIEPIDEYC[+57]VQQLK | 3.51 | 1.63E-03 | 0.0143 | 6 |
| P07901 | HS90A_MOUSE | HNDDEQYAWESSAGGSFTVR | 3.61 | 2.19E-03 | 0.0143 | 6 |
| P07901 | HS90A_MOUSE | HSQFIGYPITLFVEK | 1.75 | 4.98E-03 | 0.0155 | 6 |
| P07901 | HS90A_MOUSE | HFSVEGQLEFR | 2.64 | 5.04E-03 | 0.0155 | 6 |
| Q61879 | MYH10_MOUSE | GDEVM[+16]VELAENGK | 1.30 | 2.43E-02 | 0.0364 | 6 |
| O54983 | CRYM_MOUSE | SSSLLIPPLEAALANFSK | 1.45 | 2.54E-02 | 0.0375 | 6 |
| Q8R5C5 | ACTY_MOUSE | TLFSNIVLSGGSTLFK | 2.99 | 3.62E-04 | 0.0143 | 5 |
| P68254 | 1433T_MOUSE | AVTEQGAELSNEER | 2.81 | 7.05E-04 | 0.0143 | 5 |
| Q6P9K8 | CSKI1_MOUSE | TLSGPVTGLLATAR | 1.38 | 7.59E-04 | 0.0143 | 5 |
| Q8R5C5 | ACTY_MOUSE | IWQYVYSK | 1.66 | 7.76E-04 | 0.0143 | 5 |
| P40124 | CAP1_MOUSE | GAVPYVQAFDSLLANPVAEYLK | 3.85 | 7.93E-04 | 0.0143 | 5 |
| Q9R0P5 | DEST_MOUSE | C[+57]IVVEEGKEILVGDVGATITDPFK | 2.44 | 8.16E-04 | 0.0143 | 5 |
| Q9R0P5 | DEST_MOUSE | LGGSLIVAFEGSPV | 2.41 | 8.51E-04 | 0.0143 | 5 |
| Q923G3 | Q923G3_MOUSE | SVQTFADK | 3.08 | 1.21E-03 | 0.0143 | 5 |
| P59999 | ARPC4_MOUSE | ELLLQPVTISR | 2.60 | 1.23E-03 | 0.0143 | 5 |
| Q6P9K8 | CSKI1_MOUSE | SVSESSPGDSPVKPPEGSSGAAR | 5.54 | 1.27E-03 | 0.0143 | 5 |

| Protein Accession | Protein Description | Modified Peptide Sequence | co-IP:Ctrl | p value | q value | Sig Peptide Count |
|---|---|---|---|---|---|---|
| P59999 | ARPC4_MOUSE | ATLQAALC[+57]LENFSSQVVER | 3.47 | 1.57E-03 | 0.0143 | 5 |
| P68254 | 1433T_MOUSE | SIC[+57]TTVLELLDK | 3.15 | 1.63E-03 | 0.0143 | 5 |
| P59999 | ARPC4_MOUSE | AENFFILR | 3.29 | 1.70E-03 | 0.0143 | 5 |
| P68254 | 1433T_MOUSE | TAFDEAIAELDTLNEDSYK | 4.21 | 1.71E-03 | 0.0143 | 5 |
| P40124 | CAP1_MOUSE | ALLATASQC[+57]QQPAGNK | 1.55 | 1.82E-03 | 0.0143 | 5 |
| Q8R5C5 | ACTY_MOUSE | DQLQTFSEEHPVLLTEAPLNPSK | 2.54 | 1.83E-03 | 0.0143 | 5 |
| P80317 | TCPZ_MOUSE | AQLGVQAFADALLIIPK | 4.13 | 2.01E-03 | 0.0143 | 5 |
| P99029 | PRDX5_MOUSE | ATDLLLDDSLVSLFGNR | 2.46 | 2.10E-03 | 0.0143 | 5 |
| P99029 | PRDX5_MOUSE | VGDAIPSVEVFEGEPGK | 1.66 | 2.13E-03 | 0.0143 | 5 |
| P80317 | TCPZ_MOUSE | NAIDDGC[+57]VVPGAGAVEVALAEALIK | 2.50 | 2.22E-03 | 0.0143 | 5 |
| Q9CVB6 | ARPC2_MOUSE | ASHTAPQVLFSHR | 3.13 | 2.27E-03 | 0.0143 | 5 |
| Q6P9K8 | CSKI1_MOUSE | NTYSQTALDIVHQFTTSQASK | 4.35 | 2.30E-03 | 0.0143 | 5 |
| Q6P9K8 | CSKI1_MOUSE | GEASAEGPPLAR | 4.44 | 2.40E-03 | 0.0143 | 5 |
| Q8BJH1 | ZC21A_MOUSE | ASSVNSPLGNKPQTLSPSHR | 4.69 | 2.58E-03 | 0.0143 | 5 |
| P80317 | TCPZ_MOUSE | VATAQDDITGDGTTSNVLIIGELLK | 1.70 | 2.60E-03 | 0.0143 | 5 |
| P80317 | TCPZ_MOUSE | GIDPFSLDALAK | 1.69 | 3.10E-03 | 0.0143 | 5 |
| Q923G3 | Q923G3_MOUSE | QEALKNDLVEALKR | 3.59 | 3.93E-03 | 0.0147 | 5 |
| Q923G3 | Q923G3_MOUSE | NDLVEALK | 3.02 | 4.00E-03 | 0.0148 | 5 |
| Q9R0P5 | DEST_MOUSE | HEYQANGPEDLNR | 1.47 | 4.10E-03 | 0.0148 | 5 |
| Q9R0Q6 | ARC1A_MOUSE | TLESSIQGLR | 2.87 | 4.23E-03 | 0.0149 | 5 |
| Q9R0Q6 | ARC1A_MOUSE | STVLSLDWHPNNVLLAAGSC[+57]DFK | 4.64 | 4.34E-03 | 0.0150 | 5 |
| Q9CVB6 | ARPC2_MOUSE | VYGSFLVNPEPGYNVSLLYDLENLPASK | 9.51 | 4.58E-03 | 0.0152 | 5 |
| Q923G3 | Q923G3_MOUSE | SKQEALKNDLVEALK | 4.51 | 4.81E-03 | 0.0154 | 5 |
| Q8R5C5 | ACTY_MOUSE | AGFAGDQIPK | 1.58 | 4.89E-03 | 0.0155 | 5 |
| Q923G3 | Q923G3_MOUSE | NDLVEALKR | 3.12 | 5.00E-03 | 0.0155 | 5 |
| Q9R0Q6 | ARC1A_MOUSE | FC[+57]TTGIDGAMTIWDFK | 4.96 | 5.07E-03 | 0.0155 | 5 |
| P40124 | CAP1_MOUSE | LSDLLAPISEQIQEVITFR | 5.99 | 5.46E-03 | 0.0158 | 5 |
| Q8BJH1 | ZC21A_MOUSE | AIAAPQAGANTK | 3.42 | 5.66E-03 | 0.0160 | 5 |
| P99029 | PRDX5_MOUSE | THLPGFVEQAGALK | 1.76 | 6.99E-03 | 0.0175 | 5 |
| Q9CVB6 | ARPC2_MOUSE | MILLEVNNR | 3.28 | 7.21E-03 | 0.0177 | 5 |
| P68254 | 1433T_MOUSE | YLIANATNPESK | 2.22 | 7.67E-03 | 0.0181 | 5 |
| P99029 | PRDX5_MOUSE | VNLAELFK | 2.01 | 8.41E-03 | 0.0188 | 5 |
| Q9R0Q6 | ARC1A_MOUSE | LAWVSHDSTVSVADASK | 3.43 | 8.61E-03 | 0.0190 | 5 |
| Q6P9K8 | CSKI1_MOUSE | LLLDSGINAQVR | 4.42 | 9.31E-03 | 0.0199 | 5 |
| P59999 | ARPC4_MOUSE | VLIEGSINSVR | 2.83 | 1.02E-02 | 0.0208 | 5 |
| P40124 | CAP1_MOUSE | NSLDC[+57]EIVSAK | 1.14 | 1.06E-02 | 0.0213 | 5 |
| Q9R0P5 | DEST_MOUSE | EILVGDVGATITDPFK | 2.02 | 1.12E-02 | 0.0219 | 5 |
| Q9R0Q6 | ARC1A_MOUSE | DGIWKPTLVILR | 4.24 | 1.32E-02 | 0.0240 | 5 |
| P68254 | 1433T_MOUSE | YLAEVAC[+57]GDDRK | 2.55 | 1.41E-02 | 0.0250 | 5 |
| Q9R0P5 | DEST_MOUSE | YALYDASFETK | 3.82 | 1.60E-02 | 0.0271 | 5 |
| Q8C0M9 | ASGL1_MOUSE | GLGGLILVNK | 5.83 | 1.71E-02 | 0.0283 | 5 |
| P63038 | CH60_MOUSE | ALMLQGVDLLADAVAVTM[+16]GPK | 1.73 | 1.85E-02 | 0.0298 | 5 |
| Q9D0F9 | PGM1_MOUSE | LSGTGSAGATIR | 4.19 | 1.94E-02 | 0.0308 | 5 |
| Q9D8E6 | RL4_MOUSE | FC[+57]IWTESAFR | 2.52 | 2.32E-02 | 0.0352 | 5 |
| Q9D8E6 | RL4_MOUSE | QPYAVSELAGHQTSAESWGTGR | 1.90 | 2.70E-02 | 0.0393 | 5 |
| P62137 | PP1A_MOUSE | IRYPENFFLLR | 4.75 | 2.77E-02 | 0.0400 | 5 |
| Q9Z1G3 | VATC1_MOUSE | VFVESVLR | 0.90 | 2.92E-02 | 0.0417 | 5 |
| P07724 | ALBU_MOUSE | ENYGELADC[+57]C[+57]TK | 3.34 | 3.08E-02 | 0.0435 | 5 |
| A0A0B6VMB2 | A0A0B6VMB2_MC | NTQPIMDTDGSYFVYSK | 2.56 | 3.46E-02 | 0.0480 | 5 |
| Q6URW6 | MYH14_MOUSE | DQADFSVLHYAGK | 4.74 | 3.48E-02 | 0.0483 | 5 |
| Q7TQD2 | TPPP_MOUSE | LSLESEGANEGATAAPELSALEEAFRR | 2.99 | 4.70E-05 | 0.0143 | 5 |
| O08539 | BIN1_MOUSE | LVDQALLTMDTYLGQFPDIK | 2.99 | 5.96E-05 | 0.0143 | 5 |
| Q9WV60 | GSK3B_MOUSE | VTTVVATPGQGPDRPQEVSYTDTK | 3.80 | 1.07E-03 | 0.0143 | 5 |
| P80314 | TCPB_MOUSE | LALVTGGEIASTFDHPELVK | 2.33 | 1.30E-03 | 0.0143 | 5 |
| Q8R570 | SNP47_MOUSE | NLPLFSEGEAQELTQILSK | 5.25 | 1.58E-03 | 0.0143 | 5 |
| P04370 | MBP_MOUSE | YLATASTMDHAR | 1.99 | 1.61E-03 | 0.0143 | 5 |
| P04370 | MBP_MOUSE | HRDTGILDSIGR | 2.07 | 1.63E-03 | 0.0143 | 5 |
| P80314 | TCPB_MOUSE | MLPTIIADNAGYDSADLVAQLR | 2.76 | 1.72E-03 | 0.0143 | 5 |
| Q8R570 | SNP47_MOUSE | TEEVLVGLPLSSIIEIR | 6.56 | 2.07E-03 | 0.0143 | 5 |
| Q7TQD2 | TPPP_MOUSE | TITFEQFQEALEELAK | 4.40 | 2.10E-03 | 0.0143 | 5 |
| Q3THE2 | ML12B_MOUSE | DGFIDKEDLHDMLASLGK | 5.98 | 2.51E-03 | 0.0143 | 5 |
| Q9WV60 | GSK3B_MOUSE | VIGNGSFGVVYQAK | 3.45 | 2.58E-03 | 0.0143 | 5 |

| Protein Accession | Protein Description | Modified Peptide Sequence | co-IP:Ctrl | p value | q value | Sig Peptide Count |
|---|---|---|---|---|---|---|
| P80314 | TCPB_MOUSE T | LSSFIGAIAIGDLVK | 1.73 | 2.71E-03 | 0.0143 | 5 |
| Q9WV60 | GSK3B_MOUSE | DTPALFNFTTQELSSNPPLATILIPPHAF | 7.29 | 2.78E-03 | 0.0143 | 5 |
| Q3THE2 | ML12B_MOUSE | FTDEEVDELYR | 3.63 | 2.83E-03 | 0.0143 | 5 |
| P80314 | TCPB_MOUSE T | QVLLSAAEAAEVILR | 2.43 | 2.94E-03 | 0.0143 | 5 |
| Q9WV60 | GSK3B_MOUSE | IQAAASPPANATAASDTNAGDF | 3.21 | 3.09E-03 | 0.0143 | 5 |
| Q3THE2 | ML12B_MOUSE | ATSNVFAMFDQSQIQEFK | 6.12 | 3.17E-03 | 0.0143 | 5 |
| Q3THE2 | ML12B_MOUSE | ATSNVFAM[+16]FDQSQIQEFK | 3.72 | 3.18E-03 | 0.0143 | 5 |
| Q7TQD2 | TPPP_MOUSE T | VDLVDESGYVPGYK | 1.82 | 3.55E-03 | 0.0145 | 5 |
| O08539 | BIN1_MOUSE M | LQAHLVAQTNLLR | 1.36 | 3.80E-03 | 0.0147 | 5 |
| Q9QYC0 | ADDA_MOUSE | VNLQGDIVDR | 2.04 | 4.29E-03 | 0.0150 | 5 |
| Q7TQD2 | TPPP_MOUSE T | NVTVTDVDIVFSK | 1.87 | 4.58E-03 | 0.0152 | 5 |
| P04370 | MBP_MOUSE M | TQDENPVVHFFK | 2.70 | 5.05E-03 | 0.0155 | 5 |
| Q8R570 | SNP47_MOUSE | IELLEDALVLR | 4.57 | 5.35E-03 | 0.0158 | 5 |
| Q7TQD2 | TPPP_MOUSE T | LSLESEGANEGATAAPELSALEEAFR | 2.33 | 5.35E-03 | 0.0158 | 5 |
| Q8R570 | SNP47_MOUSE | DLQQQSEQLDSVLK | 2.63 | 6.02E-03 | 0.0164 | 5 |
| O08539 | BIN1_MOUSE M | LNQNLNDVLVSLEK | 1.58 | 6.53E-03 | 0.0171 | 5 |
| P47963 | RL13_MOUSE 6 | LATQLTGPVM[+16]PIR | 6.00 | 7.34E-03 | 0.0178 | 5 |
| Q9QYC0 | ADDA_MOUSE | EYQPHVIVSTTGPNPFNTLTDR | 3.72 | 8.27E-03 | 0.0187 | 5 |
| O08539 | BIN1_MOUSE M | VQAQHDYTATDTDELQLK | 2.61 | 8.33E-03 | 0.0187 | 5 |
| Q8R570 | SNP47_MOUSE | FIGKPDVAYQLISAK | 3.80 | 1.02E-02 | 0.0208 | 5 |
| Q9QYC0 | ADDA_MOUSE | TLASAGGPDNLVLLDPGK | 3.79 | 1.05E-02 | 0.0212 | 5 |
| Q9QYC0 | ADDA_MOUSE | LADLFGWSQLIYNHITTR | 6.08 | 1.11E-02 | 0.0219 | 5 |
| Q3THE2 | ML12B_MOUSE | NAFAC[+57]FDEEATGTIQEDYLR | 3.49 | 1.65E-02 | 0.0276 | 5 |
| Q8BYI9 | TENR_MOUSE | VDFILLK | 3.90 | 1.82E-02 | 0.0295 | 5 |
| Q7TMM9 | TBB2A_MOUSE | YLTVAAIFR | 4.39 | 1.88E-02 | 0.0301 | 5 |
| P17183 | ENOG_MOUSE | M[+16]VIGM[+16]DVAASEFYR | 9.50 | 1.88E-02 | 0.0301 | 5 |
| Q99104 | MYO5A_MOUSE | KTDDDAEAIC[+57]SMC[+57]NALTTAQIVK | 4.40 | 1.89E-02 | 0.0302 | 5 |
| Q9JII6 | AK1A1_MOUSE | ALGLSNFNSR | 4.90 | 1.96E-02 | 0.0310 | 5 |
| O88935 | SYN1_MOUSE S | SQSLTNAFNLPEPAPPRPSLSQDEVK | 4.09 | 2.18E-02 | 0.0337 | 5 |
| P97351 | RS3A_MOUSE 4 | VFEVSLADLQNDEVAFRK | 2.08 | 2.47E-02 | 0.0368 | 5 |
| Q7TQI3 | OTUB1_MOUSE | IQQEIAVQNPLVSER | 3.59 | 3.04E-02 | 0.0430 | 5 |
| P80315 | TCPD_MOUSE T | M[+16]PENVASRSGAPTAGPGSR | 1.29 | 3.58E-02 | 0.0494 | 5 |
| P20357 | MTAP2_MOUSE | SEVQAHSPSR | 1.55 | 3.58E-02 | 0.0494 | 5 |
| P27659 | RL3_MOUSE 60 | LEQQVPVNQVFGQDEMIDVIGVTK | 2.76 | 2.70E-05 | 0.0143 | 5 |
| Q9DBJ1 | PGAM1_MOUSE | DAGYEFDIC[+57]FTSVQK | 2.92 | 7.64E-04 | 0.0143 | 5 |
| Q9DBJ1 | PGAM1_MOUSE | HLEGLSEEAIM[+16]ELNLPTGIPIVYELDK | 6.69 | 1.02E-03 | 0.0143 | 5 |
| Q3UHL1 | CAMKV_MOUSE | EVFDWILDQGYYSER | 5.63 | 1.08E-03 | 0.0143 | 5 |
| Q62167 | DDX3X_MOUSE | DLLDLLVEAK | 3.93 | 1.13E-03 | 0.0143 | 5 |
| Q62167 | DDX3X_MOUSE | SFLLDLLNATGK | 4.54 | 1.31E-03 | 0.0143 | 5 |
| P61982 | 1433G_MOUSE | TAFDDAIAELDTLNEDSYK | 2.56 | 1.93E-03 | 0.0143 | 5 |
| Q9DBJ1 | PGAM1_MOUSE | ALPFWNEEIVPQIK | 1.54 | 2.06E-03 | 0.0143 | 5 |
| Q9D0E1 | HNRPM_MOUSE | QGGGGAGGSVPGIER | 4.65 | 2.31E-03 | 0.0143 | 5 |
| Q3UHL1 | CAMKV_MOUSE | HPNILQLVDVFVTR | 6.58 | 2.98E-03 | 0.0143 | 5 |
| Q9D0E1 | HNRPM_MOUSE | GNFGGSFAGSFGGAGGHAPGVAR | 4.29 | 3.13E-03 | 0.0143 | 5 |
| P61982 | 1433G_MOUSE | NVTELNEPLSNEER | 1.29 | 4.22E-03 | 0.0149 | 5 |
| Q9D0E1 | HNRPM_MOUSE | INEILSNALK | 3.85 | 4.34E-03 | 0.0150 | 5 |
| Q9D0E1 | HNRPM_MOUSE | AFITNIPFDVK | 3.66 | 5.08E-03 | 0.0155 | 5 |
| Q3UHL1 | CAMKV_MOUSE | AAATPEPAVAQPDSTALEGATGQAPPSSK | 2.97 | 5.53E-03 | 0.0159 | 5 |
| Q62167 | DDX3X_MOUSE | HVINFDLPSDIEEYVHR | 4.73 | 5.56E-03 | 0.0159 | 5 |
| P61982 | 1433G_MOUSE | NC[+57]SETQYESK | 1.62 | 6.27E-03 | 0.0168 | 5 |
| P61982 | 1433G_MOUSE | LGLALNYSVFYYEIQNAPEQAC[+57]HLAK | 6.83 | 6.73E-03 | 0.0172 | 5 |
| Q3UHL1 | CAMKV_MOUSE | ATPATEESTVPATQSSALPAAK | 1.80 | 7.08E-03 | 0.0175 | 5 |
| P27659 | RL3_MOUSE 60 | HGSLGFLPR | 2.57 | 1.08E-02 | 0.0215 | 5 |
| P27659 | RL3_MOUSE 60 | SINPLGGFVHYGEVTNDFIM[+16]LK | 4.01 | 1.30E-02 | 0.0238 | 5 |
| Q9DBJ1 | PGAM1_MOUSE | YADLTEDQLPSC[+57]ESLKDTIAR | 1.12 | 1.44E-02 | 0.0254 | 5 |
| Q62167 | DDX3X_MOUSE | LEQELFSGGNTGINFEK | 3.05 | 1.48E-02 | 0.0258 | 5 |
| P61982 | 1433G_MOUSE | YLAEVATGEK | 1.07 | 1.50E-02 | 0.0261 | 5 |
| Q9D0E1 | HNRPM_MOUSE | VGEVTYVELLMDAEGK | 7.64 | 1.60E-02 | 0.0271 | 5 |
| P27659 | RL3_MOUSE 60 | NNASTDYDLSDK | 5.12 | 1.64E-02 | 0.0275 | 5 |
| Q3UHL1 | CAMKV_MOUSE | ITAEEAISHEWISGNAASDK | 3.59 | 1.64E-02 | 0.0275 | 5 |
| P61161 | ARP2_MOUSE A | SMLEVNYPMENGIVR | 6.54 | 1.73E-02 | 0.0284 | 5 |
| P23116 | EIF3A_MOUSE | LLQQVAQIYQSIEFSR | 3.39 | 1.97E-02 | 0.0311 | 5 |

| Protein Accession | Protein Description | Modified Peptide Sequence | co-IP:Ctrl | p value | q value | Sig Peptide Count |
|---|---|---|---|---|---|---|
| P63318 | KPCG_MOUSE | LQLEIR | 0.99 | 2.35E-02 | 0.0356 | 5 |
| P29341 | PABP1_MOUSE | SKVDEAVAVLQAHQAK | 2.48 | 1.20E-03 | 0.0143 | 5 |
| P29341 | PABP1_MOUSE | ITGMLLEIDNSELLHM[+16]LESPESLR | 7.38 | 4.47E-03 | 0.0151 | 5 |
| P29341 | PABP1_MOUSE | ITGMLLEIDNSELLHMLESPESLR | 5.42 | 6.22E-03 | 0.0167 | 5 |
| P29341 | PABP1_MOUSE | ALYDTFSAFGNILSC[+57]K | 4.67 | 7.81E-03 | 0.0182 | 5 |
| P29341 | PABP1_MOUSE | IVATKPLYVALAQR | 1.32 | 1.70E-02 | 0.0282 | 5 |
| P56480 | ATPB_MOUSE / | IGLFGGAGVGK | 2.01 | 2.99E-03 | 0.0143 | 5 |
| P56480 | ATPB_MOUSE / | FTQAGSEVSALLGR | 1.44 | 3.08E-03 | 0.0143 | 5 |
| P56480 | ATPB_MOUSE / | TVLIMELINNVAK | 1.72 | 1.65E-02 | 0.0276 | 5 |
| Q9Z2I9 | SUCB1_MOUSE | LHGGTPANFLDVGGGATVQQVTEAFK | 4.42 | 2.41E-02 | 0.0363 | 5 |
| M7PGV1 | M7PGV1_PNEMU | VISNASC[+57]TTNC[+57]LAPLAK | 1.20 | 2.63E-02 | 0.0386 | 5 |
| P17751 | TPIS_MOUSE T | VTNGAFTGEISPGMIK | 1.65 | 1.04E-03 | 0.0143 | 5 |
| P17751 | TPIS_MOUSE T | RHVFGESDELIGQK | 1.51 | 1.95E-03 | 0.0143 | 5 |
| P46096 | SYT1_MOUSE S | NTLNPYYNESFSFEVPFEQIQK | 3.88 | 2.48E-03 | 0.0143 | 5 |
| P46096 | SYT1_MOUSE S | LGDIC[+57]FSLR | 2.26 | 3.84E-03 | 0.0147 | 5 |
| Q9CQV8 | 1433B_MOUSE | IEAELQDIC[+57]NDVLELLDK | 4.44 | 4.74E-03 | 0.0154 | 5 |
| P17751 | TPIS_MOUSE T | ELASQPDVDGFLVGGASLKPEFVDIINAK | 2.44 | 5.04E-03 | 0.0155 | 5 |
| P17751 | TPIS_MOUSE T | VSHALAEGLGVIAC[+57]IGEK | 1.67 | 5.95E-03 | 0.0163 | 5 |
| P46096 | SYT1_MOUSE S | TLNPVFNEQFTFK | 1.51 | 7.03E-03 | 0.0175 | 5 |
| Q9CQV8 | 1433B_MOUSE | YLILNATQAESK | 1.49 | 7.20E-03 | 0.0177 | 5 |
| P46096 | SYT1_MOUSE S | VFVGYNSTGAELR | 1.10 | 8.14E-03 | 0.0185 | 5 |
| P17751 | TPIS_MOUSE T | VVLAYEPVWAIGTGK | 1.29 | 9.47E-03 | 0.0202 | 5 |
| Q9CQV8 | 1433B_MOUSE | VISSIEQK | 1.20 | 1.24E-02 | 0.0231 | 5 |
| Q9CQV8 | 1433B_MOUSE | QTTVSNSQQAYQEAFEISK | 1.38 | 1.26E-02 | 0.0233 | 5 |
| P16546 | SPTN1_MOUSE | EELYQNLTR | 1.76 | 1.85E-02 | 0.0298 | 5 |
| P62242 | RS8_MOUSE 4( | IIDVVYNASNNELVR | 2.13 | 2.65E-02 | 0.0388 | 5 |
| O35295 | PURB_MOUSE | GGGGGGGGPGGEQETQELASK | 4.68 | 2.56E-04 | 0.0143 | 5 |
| O35295 | PURB_MOUSE | DSLGDFIEHYAQLGPSSPEQLAAGAEEGGGPR | 5.58 | 8.36E-04 | 0.0143 | 5 |
| O35295 | PURB_MOUSE | GGGGGGGGGPGGFQPAPR | 3.21 | 8.40E-04 | 0.0143 | 5 |
| O35295 | PURB_MOUSE | GGGGFGGGPGPGGLQSGQTIALPAQGLIEFR | 3.67 | 2.43E-03 | 0.0143 | 5 |
| O35295 | PURB_MOUSE | FGGAFC[+57]R | 2.51 | 2.47E-03 | 0.0143 | 5 |
| P28652 | KCC2B_MOUSE | IC[+57]DPGLTSFEPEALGNLVEGM[+16]DFHR | 6.94 | 1.21E-02 | 0.0229 | 5 |
| P28652 | KCC2B_MOUSE | QTTAPATM[+16]STAASGTTM[+16]GLVEQAK | 6.22 | 1.29E-02 | 0.0237 | 5 |
| Q7TPR4 | ACTN1_MOUSE | KDDPLTNLNTAFDVAER | 4.89 | 2.26E-02 | 0.0346 | 5 |
| P39053 | DYN1_MOUSE I | VLNQQLTNHIR | 6.34 | 3.25E-02 | 0.0455 | 5 |
| Q8BPN8 | DMXL2_MOUSE | DGVAVITLPLGGSIK | 5.79 | 3.34E-02 | 0.0466 | 5 |
| P09411 | PGK1_MOUSE I | VNEM[+16]IIGGGM[+16]AFTFLK | 1.51 | 8.42E-03 | 0.0188 | 5 |
| P09411 | PGK1_MOUSE I | DC[+57]VGPEVENAC[+57]ANPAAGTVILLENLR | 1.74 | 1.07E-02 | 0.0214 | 5 |
| P09411 | PGK1_MOUSE I | QIVWNGPVGVFEWEAFAR | 7.58 | 1.35E-02 | 0.0243 | 5 |
| P09411 | PGK1_MOUSE I | ALESPERPFLAILGGAK | 1.38 | 1.59E-02 | 0.0270 | 5 |
| Q8R1B4 | EIF3C_MOUSE | TEPTAQQNLALQLAEK | 2.08 | 2.76E-02 | 0.0399 | 5 |
| P14152 | MDHC_MOUSE | EVGVYEALKDDSWLK | 1.82 | 6.17E-03 | 0.0166 | 5 |
| P14152 | MDHC_MOUSE | AIADHIR | 3.77 | 6.39E-03 | 0.0169 | 5 |
| P14152 | MDHC_MOUSE | NVIIWGNHSSTQYPDVNHAK | 1.37 | 1.08E-02 | 0.0215 | 5 |
| P14152 | MDHC_MOUSE | DLDVAVLVGSMPR | 1.55 | 1.18E-02 | 0.0226 | 5 |
| Q9DBG3 | AP2B1_MOUSE | VNYVVQEAIVVIR | 0.79 | 3.19E-02 | 0.0447 | 5 |
| P05064 | ALDOA_MOUSE | C[+57]PLLKPWALTFSYGR | 2.32 | 4.68E-03 | 0.0153 | 5 |
| P05064 | ALDOA_MOUSE | FSNEEIAMATVTALR | 1.35 | 1.23E-02 | 0.0231 | 5 |
| P05064 | ALDOA_MOUSE | TVPPAVTGVTFLSGGQSEEEASINLNAINK | 1.49 | 1.31E-02 | 0.0239 | 5 |
| Q9JHU4 | DYHC1_MOUSE | LLNTFLER | 1.30 | 2.31E-02 | 0.0352 | 5 |
| Q8CGF6 | WDR47_MOUSE | LILDFLNSK | 1.37 | 2.81E-02 | 0.0405 | 5 |
| A0A0G2JDN7 | A0A0G2JDN7_MO | LSAGELASLSASQVPTALTFEEPVAK | 3.53 | 3.72E-04 | 0.0143 | 4 |
| P49813 | TMOD1_MOUSE | SNDPVAFALAEMLK | 7.54 | 5.58E-04 | 0.0143 | 4 |
| A0A0G2JDN7 | A0A0G2JDN7_MO | EIEVTATQSIPSLLLEETPR | 4.60 | 6.14E-04 | 0.0143 | 4 |
| A0A0G2JDN7 | A0A0G2JDN7_MO | VDSC[+57]PFIC[+57]LGGEK | 3.49 | 1.11E-03 | 0.0143 | 4 |
| P80316 | TCPE_MOUSE ' | LGFAGVVQEISFGTTK | 1.91 | 1.29E-03 | 0.0143 | 4 |
| Q8C0M9 | ASGL1_MOUSE | TVEEAAQLALDYMK | 2.55 | 1.43E-03 | 0.0143 | 4 |
| Q7TMB8 | CYFP1_MOUSE | DFVSEAYLITLGK | 2.88 | 1.44E-03 | 0.0143 | 4 |
| Q61768 | KINH_MOUSE K | TGAEGAVLDEAK | 2.19 | 1.47E-03 | 0.0143 | 4 |
| P49813 | TMOD1_MOUSE | LADLTGPIIPK | 4.99 | 1.50E-03 | 0.0143 | 4 |
| Q9QZ83 | Q9QZ83_MOUSE | KDLYANTVLSGGTTM[+16]YPGLADR | 4.17 | 1.72E-03 | 0.0143 | 4 |
| Q9QZ83 | Q9QZ83_MOUSE | DLYANTVLSGGTTM[+16]YPGLADR | 4.11 | 1.76E-03 | 0.0143 | 4 |

| Protein Accession | Protein Description | Modified Peptide Sequence | co-IP:Ctrl | p value | q value | Sig Peptide Count |
|---|---|---|---|---|---|---|
| Q9CYT6 | CAP2_MOUSE | LINSMVAEFLK | 2.00 | 1.90E-03 | 0.0143 | 4 |
| P08553 | NFM_MOUSE | EYQDLLNVK | 4.08 | 2.00E-03 | 0.0143 | 4 |
| P47754 | CAZA2_MOUSE | DIQDSLTVSNEVQTAK | 4.59 | 2.05E-03 | 0.0143 | 4 |
| Q5SYD0 | MYO1D_MOUSE | VVSVIAELLSTK | 4.60 | 2.06E-03 | 0.0143 | 4 |
| Q63844 | MK03_MOUSE | IC[+57]DFGLAR | 1.72 | 2.45E-03 | 0.0143 | 4 |
| P47754 | CAZA2_MOUSE | VDGQQTIIAC[+57]IESHQFQAK | 4.94 | 2.47E-03 | 0.0143 | 4 |
| P49813 | TMOD1_MOUSE | TLENELDELDPDNALLPAGLR | 6.64 | 2.49E-03 | 0.0143 | 4 |
| P47754 | CAZA2_MOUSE | FTVTPSTTQVVGILK | 4.88 | 2.58E-03 | 0.0143 | 4 |
| P63330 | PP2AA_MOUSE | GAGYTFGQDISETFNHANGLTLVSR | 3.41 | 2.62E-03 | 0.0143 | 4 |
| Q61768 | KINH_MOUSE | EVLQALEELAVNYDQK | 2.75 | 2.71E-03 | 0.0143 | 4 |
| P15105 | GLNA_MOUSE | TC[+57]LLNETGDEPFQYKN | 1.67 | 2.73E-03 | 0.0143 | 4 |
| Q63844 | MK03_MOUSE | GTAGVVPVVPGEVEVVK | 2.18 | 2.76E-03 | 0.0143 | 4 |
| Q8K183 | PDXK_MOUSE | DIEDPEIVVQATVL | 2.13 | 3.08E-03 | 0.0143 | 4 |
| Q6PIE5 | AT1A2_MOUSE | SPEFTHENPLETR | 1.96 | 3.08E-03 | 0.0143 | 4 |
| Q9CYT6 | CAP2_MOUSE | SALFAQLNQGEAITK | 2.67 | 3.16E-03 | 0.0143 | 4 |
| P51863 | VA0D1_MOUSE | FC[+57]TLLGGTTADAMC[+57]PILEFEADRR | 6.37 | 3.29E-03 | 0.0143 | 4 |
| P80318 | TCPG_MOUSE | AVAQALEVIPR | 1.63 | 3.45E-03 | 0.0144 | 4 |
| Q91V92 | ACLY_MOUSE | DLVSSLTSGLLTIGDR | 2.72 | 3.48E-03 | 0.0144 | 4 |
| P60335 | PCBP1_MOUSE | AITIAGVPQSVTEC[+57]VK | 1.92 | 3.51E-03 | 0.0144 | 4 |
| Q9Z1D1 | EIF3G_MOUSE | ETDLQELFRPFGSISR | 6.32 | 3.63E-03 | 0.0146 | 4 |
| P10630 | IF4A2_MOUSE | VLITTDLLAR | 1.84 | 3.74E-03 | 0.0147 | 4 |
| Q61768 | KINH_MOUSE | ISFLENNLEQLTK | 2.35 | 3.82E-03 | 0.0147 | 4 |
| P10630 | IF4A2_MOUSE | QFYINVER | 1.50 | 3.97E-03 | 0.0148 | 4 |
| Q9Z1D1 | EIF3G_MOUSE | LPGELEPVQAAQSK | 3.61 | 3.97E-03 | 0.0148 | 4 |
| Q5SYD0 | MYO1D_MOUSE | TLFTLEELR | 4.39 | 4.04E-03 | 0.0148 | 4 |
| P80318 | TCPG_MOUSE | IVLLDSSLEYK | 1.87 | 4.20E-03 | 0.0149 | 4 |
| Q641P0 | ARP3B_MOUSE | DYEEYGPSIC[+57]R | 3.32 | 4.57E-03 | 0.0152 | 4 |
| Q9QZ83 | Q9QZ83_MOUSE | KDLYANTVLSGGTTMYPGLADR | 3.14 | 4.73E-03 | 0.0153 | 4 |
| Q9Z1D1 | EIF3G_MOUSE | VTNLSEDTR | 5.66 | 4.76E-03 | 0.0154 | 4 |
| Q61768 | KINH_MOUSE | LYLVDLAGSEK | 1.95 | 4.80E-03 | 0.0154 | 4 |
| P08553 | NFM_MOUSE | VQSLQDEVAFLR | 4.12 | 4.92E-03 | 0.0155 | 4 |
| P62908 | RS3_MOUSE | AELNEFLTR | 2.54 | 5.03E-03 | 0.0155 | 4 |
| P47754 | CAZA2_MOUSE | KVDGQQTIIAC[+57]IESHQFQAK | 3.63 | 5.06E-03 | 0.0155 | 4 |
| P80316 | TCPE_MOUSE | WVGGPEIELIAIATGGR | 5.87 | 5.07E-03 | 0.0155 | 4 |
| P63330 | PP2AA_MOUSE | NVVTIFSAPNYC[+57]YR | 2.60 | 5.10E-03 | 0.0155 | 4 |
| Q641P0 | ARP3B_MOUSE | GVDDLDFFIGDEAIDKPTYATK | 4.41 | 5.37E-03 | 0.0158 | 4 |
| Q9QZ83 | Q9QZ83_MOUSE | DLYANTVLSGGTTMYPGLADR | 5.17 | 5.44E-03 | 0.0158 | 4 |
| P49813 | TMOD1_MOUSE | TLNVESNFISGAGILR | 4.62 | 5.51E-03 | 0.0159 | 4 |
| Q8K183 | PDXK_MOUSE | DKSFLAMVVDIVR | 4.02 | 5.87E-03 | 0.0163 | 4 |
| Q921I1 | TRFE_MOUSE | DFASC[+57]HLAQAPNHVVVSR | 3.64 | 6.10E-03 | 0.0166 | 4 |
| P10630 | IF4A2_MOUSE | GFKDQIYEIFQK | 2.56 | 6.13E-03 | 0.0166 | 4 |
| Q9Z1D1 | EIF3G_MOUSE | ELAEQLGLSTGEK | 6.74 | 6.27E-03 | 0.0168 | 4 |
| Q9CYT6 | CAP2_MOUSE | LEQLSAGLDGPPR | 1.57 | 6.32E-03 | 0.0169 | 4 |
| Q641P0 | ARP3B_MOUSE | NVVLSGGSTMFR | 3.05 | 6.61E-03 | 0.0171 | 4 |
| Q6PIE5 | AT1A2_MOUSE | GIVIATGDR | 4.20 | 6.91E-03 | 0.0174 | 4 |
| Q921I1 | TRFE_MOUSE | DFQLFSSPLGK | 2.14 | 7.34E-03 | 0.0178 | 4 |
| P15105 | GLNA_MOUSE | LVLC[+57]EVFK | 1.36 | 7.47E-03 | 0.0180 | 4 |
| P80318 | TCPG_MOUSE | EMMLSIINSSITTK | 2.69 | 7.81E-03 | 0.0182 | 4 |
| Q5SYD0 | MYO1D_MOUSE | IGELVGVLVNHFK | 4.36 | 7.88E-03 | 0.0183 | 4 |
| Q61937 | NPM_MOUSE | M[+16]SVQPTVSLGGFEITPPVVLR | 6.53 | 7.88E-03 | 0.0183 | 4 |
| P60335 | PCBP1_MOUSE | LVVPATQC[+57]GSLIGK | 1.37 | 7.95E-03 | 0.0183 | 4 |
| P80316 | TCPE_MOUSE | IADGYEQAAR | 2.18 | 8.93E-03 | 0.0194 | 4 |
| P51863 | VA0D1_MOUSE | LLFEGAGSNPGDK | 1.57 | 9.34E-03 | 0.0199 | 4 |
| Q8K183 | PDXK_MOUSE | AEAGEGQKPSPAQLELR | 5.75 | 9.61E-03 | 0.0203 | 4 |
| P62908 | RS3_MOUSE | GGKPEPPAMPQPVPTA | 3.57 | 1.02E-02 | 0.0208 | 4 |
| Q61937 | NPM_MOUSE | GPSSVEDIK | 2.75 | 1.06E-02 | 0.0213 | 4 |
| P15105 | GLNA_MOUSE | TC[+57]LLNETGDEPFQYK | 1.37 | 1.09E-02 | 0.0216 | 4 |
| P08553 | NFM_MOUSE | EIEAEIQALR | 2.60 | 1.15E-02 | 0.0223 | 4 |
| P10630 | IF4A2_MOUSE | GYDVIAQAQSGTGK | 1.80 | 1.18E-02 | 0.0226 | 4 |
| P62908 | RS3_MOUSE | FGFPEGSVELYAEK | 3.65 | 1.18E-02 | 0.0226 | 4 |
| A0A0G2JDN7 | A0A0G2JDN7_MO | LGFFEGK | 2.31 | 1.19E-02 | 0.0227 | 4 |
| Q91V92 | ACLY_MOUSE | LGLVGVNLSLDGVK | 1.53 | 1.21E-02 | 0.0229 | 4 |

| Protein Accession | Protein Description | Modified Peptide Sequence | co-IP:Ctrl | p value | q value | Sig Peptide Count |
|---|---|---|---|---|---|---|
| P62908 | RS3_MOUSE | TEIIILATR | 2.06 | 1.21E-02 | 0.0229 | 4 |
| Q921I1 | TRFE_MOUSE | SAGWVIPIGLLFC[+57]K | 3.02 | 1.22E-02 | 0.0230 | 4 |
| Q9CYT6 | CAP2_MOUSE | FGLVFDHVVGIVEVINSK | 2.96 | 1.25E-02 | 0.0232 | 4 |
| Q61937 | NPM_MOUSE | MSVQPTVSLGGFEITPPVVLR | 6.54 | 1.31E-02 | 0.0239 | 4 |
| P62301 | RS13_MOUSE | GLSQSALPYR | 3.68 | 1.33E-02 | 0.0241 | 4 |
| P80318 | TCPG_MOUSE | TAVETAVLLLR | 3.00 | 1.39E-02 | 0.0248 | 4 |
| Q63844 | MK03_MOUSE | DVYIVQDLMETDLYK | 3.08 | 1.39E-02 | 0.0248 | 4 |
| Q7TMB8 | CYFP1_MOUSE | LGTPQQIAIAR | 1.30 | 1.43E-02 | 0.0253 | 4 |
| Q6PIE5 | AT1A2_MOUSE | AGQENISVSK | 1.37 | 1.45E-02 | 0.0255 | 4 |
| P60335 | PCBP1_MOUSE | IANPVEGSSGR | 1.22 | 1.47E-02 | 0.0257 | 4 |
| Q7TMB8 | CYFP1_MOUSE | FINMFAVLDELK | 5.33 | 1.52E-02 | 0.0263 | 4 |
| P80316 | TCPE_MOUSE | QMAEIAVNAVLTVADMER | 4.63 | 1.57E-02 | 0.0268 | 4 |
| P51863 | VA0D1_MOUSE | AGVLSQADYLNLVQC[+57]ETLEDLK | 3.30 | 1.62E-02 | 0.0273 | 4 |
| P51410 | RL9_MOUSE | DFNHINVELSLLGK | 3.88 | 1.62E-02 | 0.0273 | 4 |
| P62301 | RS13_MOUSE | KGLTPSQIGVILR | 4.32 | 1.65E-02 | 0.0276 | 4 |
| Q91V92 | ACLY_MOUSE | YIC[+57]TTSAIQNR | 2.56 | 1.65E-02 | 0.0276 | 4 |
| P17426 | AP2A1_MOUSE | ALQVGC[+57]LLR | 1.37 | 1.71E-02 | 0.0283 | 4 |
| Q7TMM9 | TBB2A_MOUSE | IMNTFSVMPSPK | 2.34 | 1.73E-02 | 0.0284 | 4 |
| O08638 | MYH11_MOUSE | LQQELDDLVVDLDNQR | 2.00 | 1.74E-02 | 0.0285 | 4 |
| P16546 | SPTN1_MOUSE | GAC[+57]AGSEDAVK | 2.56 | 1.80E-02 | 0.0293 | 4 |
| P28651 | CAH8_MOUSE | GAELVEGC[+57]DGILGDNFRPTQPLSDR | 1.40 | 1.84E-02 | 0.0297 | 4 |
| Q6URW6 | MYH14_MOUSE | MLIAALESK | 1.37 | 1.87E-02 | 0.0300 | 4 |
| P16858 | G3P_MOUSE | VVDLMAYMASKE | 3.24 | 1.88E-02 | 0.0301 | 4 |
| Q9QYC0 | ADDA_MOUSE | VNSEQEHFLIVPFGLLYSEVTASSLVK | 5.03 | 1.88E-02 | 0.0301 | 4 |
| Q9JMH9 | MY18A_MOUSE | TC[+57]WLILASIYHLGAAGATK | 2.61 | 2.05E-02 | 0.0322 | 4 |
| Q7TMK9 | HNRPQ_MOUSE | AIEALKEFNEDGALAVLQQFK | 3.30 | 2.17E-02 | 0.0336 | 4 |
| P48318 | DCE1_MOUSE | MVISNPAATQSDIDFLIEEIER | 1.19 | 2.22E-02 | 0.0342 | 4 |
| Q6ZQ38 | CAND1_MOUSE | TYIQC[+57]IAAISR | 2.38 | 2.28E-02 | 0.0348 | 4 |
| P20357 | MTAP2_MOUSE | VSEGPRPFAPVFFQSDDKVSLQDPSALATSK | 3.62 | 2.32E-02 | 0.0352 | 4 |
| P01837 | IGKC_MOUSE | DSTYSMSSTLTLTK | 2.62 | 2.34E-02 | 0.0355 | 4 |
| Q6ZWN5 | RS9_MOUSE | LIGEYGLR | 0.84 | 2.51E-02 | 0.0372 | 4 |
| Q9JMH9 | MY18A_MOUSE | TTSFQHLVQYLATIAGTSGTK | 6.65 | 2.71E-02 | 0.0394 | 4 |
| Q8BJH1 | ZC21A_MOUSE | LPPPPPPSYDPDYIQC[+57]PYC[+57]QR | 1.73 | 2.77E-02 | 0.0400 | 4 |
| Q9CPR4 | RL17_MOUSE | YSLDPENPTK | 2.37 | 2.83E-02 | 0.0407 | 4 |
| P97427 | DPYL1_MOUSE | IVFEDGNISVSK | 1.69 | 3.13E-02 | 0.0440 | 4 |
| P14152 | MDHC_MOUSE | DLDVAVLVGSM[+16]PR | 2.78 | 3.19E-02 | 0.0447 | 4 |
| P28652 | KCC2B_MOUSE | LTQYIDGQGRPR | 3.60 | 3.35E-02 | 0.0467 | 4 |
| Q501J6 | DDX17_MOUSE | STC[+57]IYGGAPK | 1.55 | 3.37E-02 | 0.0469 | 4 |
| O08599 | STXB1_MOUSE | LAEQIATLC[+57]ATLK | 3.87 | 3.53E-02 | 0.0489 | 4 |
| Q9QYG0 | NDRG2_MOUSE | GIIQHAPNLENIELYWNSYNNR | 5.40 | 2.07E-04 | 0.0143 | 4 |
| Q922F4 | TBB6_MOUSE | M[+16]ASTFIGNSTAIQELFK | 2.45 | 2.35E-04 | 0.0143 | 4 |
| O54983 | CRYM_MOUSE | SLGMAVEDLVAAK | 1.88 | 5.18E-04 | 0.0143 | 4 |
| Q9QXS1 | PLEC_MOUSE | ISLETYNLFR | 2.95 | 5.48E-04 | 0.0143 | 4 |
| Q61548 | AP180_MOUSE | GLGSDLDSSLASLVGNLGISGTTSK | 4.26 | 6.45E-04 | 0.0143 | 4 |
| Q61361 | PGCB_MOUSE | C[+57]EVQHGIDDSSDAVEVK | 1.26 | 7.35E-04 | 0.0143 | 4 |
| Q9QYJ0 | DNJA2_MOUSE | ITFTGEADQAPGVEPGDIVLLLQEK | 4.37 | 9.04E-04 | 0.0143 | 4 |
| Q922F4 | TBB6_MOUSE | MASTFIGNSTAIQELFK | 2.64 | 1.11E-03 | 0.0143 | 4 |
| P99024 | TBB5_MOUSE | M[+16]AVTFIGNSTAIQELFK | 2.25 | 1.18E-03 | 0.0143 | 4 |
| F7B0R9 | F7B0R9_MOUSE | SPEQPGTKPPLPR | 5.13 | 1.58E-03 | 0.0143 | 4 |
| Q9QYJ0 | DNJA2_MOUSE | VIEPGC[+57]VR | 2.78 | 1.66E-03 | 0.0143 | 4 |
| Q9QYG0 | NDRG2_MOUSE | FGDMQEIIQNFVR | 2.80 | 1.98E-03 | 0.0143 | 4 |
| P68368 | TBA4A_MOUSE | AVFVDLEPTVIDEIR | 2.34 | 2.15E-03 | 0.0143 | 4 |
| Q61548 | AP180_MOUSE | DPLADLNIKDFL | 1.81 | 2.19E-03 | 0.0143 | 4 |
| F7B0R9 | F7B0R9_MOUSE | KPLLPSLTLPYPPTGLDTSPGESER | 5.40 | 2.27E-03 | 0.0143 | 4 |
| Q922F4 | TBB6_MOUSE | LHFFM[+16]PGFAPLTAR | 2.80 | 2.58E-03 | 0.0143 | 4 |
| F7B0R9 | F7B0R9_MOUSE | LIPVILPPEPR | 4.71 | 2.69E-03 | 0.0143 | 4 |
| P08551 | NFL_MOUSE | IDSLMDEIAFLK | 6.00 | 2.82E-03 | 0.0143 | 4 |
| Q61548 | AP180_MOUSE | ATNSSWVVVFK | 1.92 | 2.90E-03 | 0.0143 | 4 |
| Q9QYJ0 | DNJA2_MOUSE | NVLC[+57]SAC[+57]SGQGGK | 4.08 | 2.93E-03 | 0.0143 | 4 |
| Q61656 | DDX5_MOUSE | TGTAYTFFTPNNIK | 2.93 | 3.10E-03 | 0.0143 | 4 |
| F7B0R9 | F7B0R9_MOUSE | DEQSHVGAAPALR | 3.39 | 3.14E-03 | 0.0143 | 4 |
| Q9QYG0 | NDRG2_MOUSE | YALNHPDTVEGLVLINIDPNAK | 2.42 | 3.52E-03 | 0.0144 | 4 |

| Protein Accession | Protein Description | Modified Peptide Sequence | co-IP:Ctrl | p value | q value | Sig Peptide Count |
|---|---|---|---|---|---|---|
| Q9QYG0 | NDRG2_MOUSE | TASLTSAASIDGSR | 1.83 | 3.57E-03 | 0.0145 | 4 |
| P08551 | NFL_MOUSE | LAAEDATNEK | 4.16 | 3.76E-03 | 0.0147 | 4 |
| P08551 | NFL_MOUSE | FASFIER | 2.26 | 3.76E-03 | 0.0147 | 4 |
| Q9QXS1 | PLEC_MOUSE | LSVAAQEAAR | 4.56 | 3.90E-03 | 0.0147 | 4 |
| H3BJD0 | H3BJD0_MOUSE | TEAVSPTVSQLSAVFENSEPPGALTSGK | 5.33 | 3.97E-03 | 0.0148 | 4 |
| Q922F4 | TBB6_MOUSE | LHFFMPGFAPLTAR | 3.97 | 4.43E-03 | 0.0151 | 4 |
| Q8C0P5 | COR2A_MOUSE | WNPFNDFEIASC[+57]SEDATIK | 7.12 | 4.68E-03 | 0.0153 | 4 |
| P50518 | VATE1_MOUSE | DDLITDLLNEAK | 2.24 | 4.79E-03 | 0.0154 | 4 |
| Q9JI91 | ACTN2_MOUSE | C[+57]QLEINFNTLQTK | 4.76 | 5.41E-03 | 0.0158 | 4 |
| Q9JI91 | ACTN2_MOUSE | LVSIGAEEIVDGNVK | 4.18 | 6.17E-03 | 0.0166 | 4 |
| P99024 | TBB5_MOUSE | ALTVPELTQQVFDAK | 2.56 | 6.65E-03 | 0.0172 | 4 |
| Q61656 | DDX5_MOUSE | LIDFLEC[+57]GK | 2.74 | 7.22E-03 | 0.0177 | 4 |
| P08551 | NFL_MOUSE | TLEIEAC[+57]R | 2.94 | 7.29E-03 | 0.0177 | 4 |
| Q9JI91 | ACTN2_MOUSE | GYEEWLLNEIR | 1.42 | 7.82E-03 | 0.0182 | 4 |
| Q9QXS1 | PLEC_MOUSE | GFFDPNTHENLTYLQLLER | 5.12 | 8.33E-03 | 0.0187 | 4 |
| P99024 | TBB5_MOUSE | MAVTFIGNSTAIQELFK | 3.85 | 8.38E-03 | 0.0188 | 4 |
| P68368 | TBA4A_MOUSE | EIIDPVLDR | 1.79 | 8.40E-03 | 0.0188 | 4 |
| Q501J6 | DDX17_MOUSE | LIQLMEEIMAEK | 3.40 | 8.44E-03 | 0.0188 | 4 |
| P68368 | TBA4A_MOUSE | TIGGGDDSFTTFFC[+57]ETGAGK | 2.65 | 9.58E-03 | 0.0203 | 4 |
| Q501J6 | DDX17_MOUSE | APILIATDVASR | 2.22 | 9.59E-03 | 0.0203 | 4 |
| O54983 | CRYM_MOUSE | FASTVQGDVR | 2.36 | 9.60E-03 | 0.0203 | 4 |
| Q8C0P5 | COR2A_MOUSE | GLDVSSC[+57]EIFR | 3.19 | 9.96E-03 | 0.0206 | 4 |
| Q9QXS1 | PLEC_MOUSE | DGHNLISLLEVLSGDSLPR | 5.06 | 9.99E-03 | 0.0206 | 4 |
| Q9JI91 | ACTN2_MOUSE | FAIQDISVEETSAK | 3.93 | 1.00E-02 | 0.0206 | 4 |
| Q8C0P5 | COR2A_MOUSE | ENC[+57]YDSVPITR | 4.48 | 1.08E-02 | 0.0215 | 4 |
| P68368 | TBA4A_MOUSE | DVNAAIAAIK | 1.74 | 1.10E-02 | 0.0217 | 4 |
| P50518 | VATE1_MOUSE | ARDDLITDLLNEAK | 1.94 | 1.13E-02 | 0.0221 | 4 |
| Q61548 | AP180_MOUSE | IAAAQYSVTGSAVAR | 1.33 | 1.16E-02 | 0.0224 | 4 |
| Q61361 | PGCB_MOUSE | SWEEAESQC[+57]R | 1.63 | 1.16E-02 | 0.0224 | 4 |
| P12970 | RL7A_MOUSE | AGVNTVTTLVENK | 5.83 | 1.32E-02 | 0.0240 | 4 |
| Q61656 | DDX5_MOUSE | LLQLVEDR | 1.94 | 1.37E-02 | 0.0246 | 4 |
| Q9QYJ0 | DNJA2_MOUSE | IGLVEALC[+57]GFQFTFK | 6.44 | 1.43E-02 | 0.0253 | 4 |
| Q501J6 | DDX17_MOUSE | GVEIC[+57]IATPGR | 2.56 | 1.66E-02 | 0.0277 | 4 |
| P58252 | EF2_MOUSE | AYLPVNESFGFTADLR | 0.90 | 1.81E-02 | 0.0294 | 4 |
| Q8K183 | PDXK_MOUSE | VVPVADIITPNQFEAELLSGR | 1.84 | 1.84E-02 | 0.0297 | 4 |
| Q9DCL9 | PUR6_MOUSE | ASILNTWISLK | 1.84 | 1.90E-02 | 0.0303 | 4 |
| P99024 | TBB5_MOUSE | ISVYYNEATGGK | 1.44 | 1.90E-02 | 0.0303 | 4 |
| Q8VDD5 | MYH9_MOUSE | LTEM[+16]ETM[+16]QSQLMAEK | 4.36 | 1.92E-02 | 0.0306 | 4 |
| Q9CVB6 | ARPC2_MOUSE | VMVSISLK | 5.07 | 1.94E-02 | 0.0308 | 4 |
| P61021 | RAB5B_MOUSE | LVLLGESAVGK | 6.28 | 1.98E-02 | 0.0313 | 4 |
| P17427 | AP2A2_MOUSE | VVHLLNDQHLGVVTAATSLITTLAQK | 1.71 | 2.42E-02 | 0.0364 | 4 |
| Q8QZY1 | EIF3L_MOUSE | LHSLLGDYYQAIK | 2.79 | 2.43E-02 | 0.0364 | 4 |
| Q9CZM2 | RL15_MOUSE | SLQSVAEER | 2.50 | 2.45E-02 | 0.0366 | 4 |
| Q62261 | SPTB2_MOUSE | LLTQHENIKNEIDNYEEDYQK | 1.74 | 2.53E-02 | 0.0374 | 4 |
| Q9CZ04 | CSN7A_MOUSE | VTTAAAAAATSQDPEQHLTELR | 4.85 | 2.61E-02 | 0.0384 | 4 |
| P07356 | ANXA2_MOUSE | TPAQYDASELK | 4.33 | 2.70E-02 | 0.0393 | 4 |
| P05064 | ALDOA_MOUSE | FSNEEIAMATVTALRR | 2.55 | 2.82E-02 | 0.0406 | 4 |
| Q61879 | MYH10_MOUSE | MQAHIQDLEEQLDEEEGAR | 3.53 | 3.11E-02 | 0.0438 | 4 |
| Q8BZ98 | DYN3_MOUSE | RPLVLQLVTSK | 1.27 | 3.35E-02 | 0.0467 | 4 |
| Q9Z1G3 | VATC1_MOUSE | VQENLLASGVDLVTYITR | 3.25 | 1.59E-03 | 0.0143 | 4 |
| Q60932 | VDAC1_MOUSE | VTQSNFAVGYK | 2.08 | 1.65E-03 | 0.0143 | 4 |
| Q60932 | VDAC1_MOUSE | EHINLGC[+57]DVDFDIAGPSIR | 2.99 | 2.93E-03 | 0.0143 | 4 |
| Q9Z1G3 | VATC1_MOUSE | VGTLDVLVGLSDELAK | 2.28 | 3.64E-03 | 0.0146 | 4 |
| Q6PHZ2 | KCC2D_MOUSE | STVASMMHR | 3.79 | 4.31E-03 | 0.0150 | 4 |
| Q60932 | VDAC1_MOUSE | TDEFQLHTNVNDGTEFGGSIYQK | 2.57 | 4.70E-03 | 0.0153 | 4 |
| P14131 | RS16_MOUSE | GPLQSVQVFGR | 3.40 | 5.31E-03 | 0.0158 | 4 |
| Q61171 | PRDX2_MOUSE | KEGGLGPLNIPLLADVTK | 1.81 | 5.63E-03 | 0.0160 | 4 |
| P14131 | RS16_MOUSE | FAGVDIR | 3.05 | 5.71E-03 | 0.0161 | 4 |
| P48036 | ANXA5_MOUSE | GLGTDEDSILNLLTSR | 1.97 | 7.14E-03 | 0.0176 | 4 |
| P14131 | RS16_MOUSE | LLEPVLLLGK | 4.10 | 1.01E-02 | 0.0207 | 4 |
| Q6PHZ2 | KCC2D_MOUSE | IC[+57]DPGLTAFEPEALGNLVEGM[+16]DFHR | 7.38 | 1.02E-02 | 0.0208 | 4 |
| P48036 | ANXA5_MOUSE | GTVTDFPGFDGR | 1.14 | 1.08E-02 | 0.0215 | 4 |

| Protein Accession | Protein Description | Modified Peptide Sequence | co-IP:Ctrl | p value | q value | Sig Peptide Count |
|---|---|---|---|---|---|---|
| Q9Z1G3 | VATC1_MOUSE | FNIPDLK | 1.27 | 1.10E-02 | 0.0217 | 4 |
| Q60932 | VDAC1_MOUSE | GYGFGLIK | 1.64 | 1.14E-02 | 0.0222 | 4 |
| P97351 | RS3A_MOUSE 4 | AC[+57]QSIYPLHDVFVR | 3.01 | 1.21E-02 | 0.0229 | 4 |
| P97351 | RS3A_MOUSE 4 | VFEVSLADLQNDEVAFR | 3.37 | 1.55E-02 | 0.0266 | 4 |
| P14131 | RS16_MOUSE 4 | EIKDILIQYDR | 4.20 | 1.64E-02 | 0.0275 | 4 |
| Q61598 | GDIB_MOUSE F | NTNDANSC[+57]QIIIPQNQVNR | 1.44 | 1.87E-02 | 0.0300 | 4 |
| Q8K1M6 | DNM1L_MOUSE | DTLQSELVGQLYK | 1.32 | 2.12E-02 | 0.0330 | 4 |
| Q04447 | KCRB_MOUSE | VLTPELYAELR | 4.37 | 2.24E-02 | 0.0344 | 4 |
| Q547J4 | Q547J4_MOUSE | AEEAGIGDTPNQEDQAAGHVTQAR | 0.86 | 2.37E-02 | 0.0358 | 4 |
| Q8QZY1 | EIF3L_MOUSE | VYELQASR | 3.34 | 2.47E-02 | 0.0368 | 4 |
| P52480 | KPYM_MOUSE | C[+57]DENILWLDYK | 5.14 | 2.66E-02 | 0.0389 | 4 |
| Q3UHD9 | AGAP2_MOUSE | STAGTGASAAAAGGGGSAAVTTSGGVGAGAGTF | 1.11 | 2.91E-02 | 0.0416 | 4 |
| Q8H156 | RAN3_ARATH GT | LVIVGDGGTGK | 1.02 | 3.00E-02 | 0.0426 | 4 |
| Q8BYI9 | TENR_MOUSE | LDSSVVPNTVTEFAITR | 1.23 | 3.39E-02 | 0.0472 | 4 |
| Q9EQH3 | VPS35_MOUSE | LLDEAIQAVK | 2.39 | 3.50E-02 | 0.0485 | 4 |
| O08709 | PRDX6_MOUSE | DLAILLGM[+16]LDPVEK | 2.43 | 1.51E-03 | 0.0143 | 4 |
| Q91VR5 | DDX1_MOUSE | DLGLAFEIPAHIK | 2.58 | 3.01E-03 | 0.0143 | 4 |
| O08709 | PRDX6_MOUSE | DLAILLGMLDPVEK | 2.51 | 3.22E-03 | 0.0143 | 4 |
| Q91VR5 | DDX1_MOUSE | GHVDVLAPTVQELAALEK | 2.32 | 3.75E-03 | 0.0147 | 4 |
| Q91VR5 | DDX1_MOUSE | FLVLDEADGLLSQGYSDFINR | 4.50 | 6.79E-03 | 0.0172 | 4 |
| O08709 | PRDX6_MOUSE | VVDSLQLTGTKPVATPVDWK | 1.19 | 1.42E-02 | 0.0251 | 4 |
| P12970 | RL7A_MOUSE 6 | HWGGNVLGPK | 2.12 | 1.94E-02 | 0.0308 | 4 |
| Q9WUM4 | COR1C_MOUSE | ETIC[+57]SQDER | 0.59 | 2.94E-02 | 0.0419 | 4 |
| Q64332 | SYN2_MOUSE | VENHYDFQDIASVVALTQTYATAEPFIDAK | 6.91 | 1.72E-03 | 0.0143 | 4 |
| Q64332 | SYN2_MOUSE | QHAFGMAENEDFR | 1.67 | 4.49E-03 | 0.0151 | 4 |
| Q64332 | SYN2_MOUSE | SFRPDFVLIR | 1.42 | 9.75E-03 | 0.0204 | 4 |
| Q9ERD7 | TBB3_MOUSE | M[+16]SM[+16]KEVDEQM[+16]LAIQSK | 1.26 | 1.86E-02 | 0.0300 | 4 |
| Q61781 | K1C14_MOUSE | YC[+57]M[+16]QLAQIQEM[+16]IGSVEEQLAQLR | 8.60 | 4.16E-04 | 0.0143 | 4 |
| P40142 | TKT_MOUSE Tr | NMAEQIIQEIYSQVQSK | 3.52 | 3.75E-03 | 0.0147 | 4 |
| Q61781 | K1C14_MOUSE | ADLEM[+16]QIESLKEELAYLK | 4.65 | 4.25E-03 | 0.0149 | 4 |
| P40142 | TKT_MOUSE Tr | ILTVEDHYYEGGIGEAVSAAVVGEPGVTVTF | 2.07 | 4.43E-03 | 0.0151 | 4 |
| Q61781 | K1C14_MOUSE | GQVGGDVNVEM[+16]DAAPGVDLSR | 2.11 | 6.64E-03 | 0.0172 | 4 |
| Q61781 | K1C14_MOUSE | ADLEMQIESLKEELAYLK | 6.39 | 8.62E-03 | 0.0190 | 4 |
| P40142 | TKT_MOUSE Tr | LDNLVAIFDINR | 1.49 | 1.15E-02 | 0.0223 | 4 |
| Q8QZY1 | EIF3L_MOUSE | LAGFLDLTEQEFR | 4.86 | 1.35E-02 | 0.0243 | 4 |
| Q8C0P5 | COR2A_MOUSE | DSVIAGPVK | 4.54 | 2.42E-02 | 0.0364 | 4 |
| P11499 | HS90B_MOUSE | TLTLVDTGIGMTK | 3.92 | 2.43E-02 | 0.0364 | 4 |
| Q9CPR4 | RL17_MOUSE 6 | GLDVDSLVIEHIQVNK | 1.56 | 2.46E-02 | 0.0367 | 4 |
| P19001 | K1C19_MOUSE | LSVEADINGLR | 1.13 | 3.18E-02 | 0.0446 | 4 |
| Q02053 | UBA1_MOUSE | LAYVAAGDLAPINAFIGGLAAQEVM[+16]K | 2.12 | 3.21E-03 | 0.0143 | 4 |
| Q02053 | UBA1_MOUSE | LAYVAAGDLAPINAFIGGLAAQEVMK | 3.36 | 3.37E-03 | 0.0143 | 4 |
| Q02053 | UBA1_MOUSE | LAGTQPLEVLEAVQR | 1.85 | 4.67E-03 | 0.0153 | 4 |
| Q02053 | UBA1_MOUSE | YFLVGAGAIGC[+57]ELLK | 1.61 | 9.69E-03 | 0.0204 | 4 |
| Q9JMH9 | MY18A_MOUSE | WQALSTLLEAFGNSPTIMNGSATR | 7.76 | 1.77E-02 | 0.0289 | 4 |
| P97427 | DPYL1_MOUSE | QIGENLIVPGGVK | 6.60 | 1.82E-02 | 0.0295 | 4 |
| Q61361 | PGCB_MOUSE | FKDLEALEEEK | 7.08 | 1.84E-02 | 0.0297 | 4 |
| Q2M3X8 | PHAR1_MOUSE | LSQRPTAEELEQR | 6.98 | 3.15E-02 | 0.0443 | 4 |
| P05063 | ALDOC_MOUSE | GILAADESVGSMAK | 2.02 | 5.15E-03 | 0.0155 | 4 |
| P05063 | ALDOC_MOUSE | YASIC[+57]QQNGIVPIVEPEILPDGDHDLK | 1.77 | 1.02E-02 | 0.0208 | 4 |
| P05063 | ALDOC_MOUSE | TVPPAVPGVTFLSGGQSEEEASLNLNAINR | 1.60 | 1.14E-02 | 0.0222 | 4 |
| P05063 | ALDOC_MOUSE | LSQIGVENTEENR | 2.45 | 1.21E-02 | 0.0229 | 4 |
| O88935 | SYN1_MOUSE | QLIVELVVNK | 1.35 | 1.15E-02 | 0.0223 | 4 |
| O88935 | SYN1_MOUSE | LGTEEFPLIDQTFYPNHK | 1.41 | 1.20E-02 | 0.0228 | 4 |
| O08638 | MYH11_MOUSE | IAQLEEQVEQEAR | 1.11 | 2.17E-02 | 0.0336 | 4 |
| O08638 | MYH11_MOUSE | ALEEALEAKEELER | 1.07 | 2.90E-02 | 0.0414 | 4 |
| Q8BFR5 | EFTU_MOUSE | DLDKPFLLPVESVYSIPGR | 3.06 | 2.42E-04 | 0.0143 | 3 |
| P28663 | SNAB_MOUSE | VAAYAAQLEQYQK | 2.02 | 3.67E-04 | 0.0143 | 3 |
| P29515 | TBB7_ARATH Tub | EVDEQMLNVQNK | 2.24 | 3.72E-04 | 0.0143 | 3 |
| Q2PFD7 | PSD3_MOUSE | ELLTTDGNEPVGLK | 2.62 | 4.92E-04 | 0.0143 | 3 |
| P14094 | AT1B1_MOUSE | SYEAYVLNIIR | 2.03 | 6.12E-04 | 0.0143 | 3 |
| Q8K0S0 | PHYIP_MOUSE | M[+42]ELLSTPHSIEINNITC[+57]DSFR | 4.24 | 7.04E-04 | 0.0143 | 3 |
| P29515 | TBB7_ARATH Tub | EVDEQM[+16]LNVQNK | 1.76 | 7.20E-04 | 0.0143 | 3 |

| Protein Accession | Protein Description | Modified Peptide Sequence | co-IP:Ctrl | p value | q value | Sig Peptide Count |
|---|---|---|---|---|---|---|
| O08788 | DCTN1_MOUSE | AFLQGGQEATDIALLLR | 3.21 | 8.31E-04 | 0.0143 | 3 |
| Q5PR73 | DIRA2_MOUSE | ELFQELLNLEK | 2.63 | 8.41E-04 | 0.0143 | 3 |
| P68404 | KPCB_MOUSE | C[+57]SLNPEWNETFR | 2.76 | 8.69E-04 | 0.0143 | 3 |
| Q9Z2X1 | HNRPF_MOUSE | ATENDIYNFFSPLNPVR | 2.84 | 1.24E-03 | 0.0143 | 3 |
| Q91YR1 | TWF1_MOUSE | INEVQTDVSVDTK | 2.54 | 1.39E-03 | 0.0143 | 3 |
| P48318 | DCE1_MOUSE ( | TETDFSNLFAQDLLPAK | 3.28 | 1.42E-03 | 0.0143 | 3 |
| Q8R429 | AT2A1_MOUSE | TASEM[+16]VLADDNFSTIVAAVEEGR | 3.09 | 1.57E-03 | 0.0143 | 3 |
| A2N1N1 | A2N1N1_MOUSE | ESGVPDR | 7.79 | 1.62E-03 | 0.0143 | 3 |
| Q9D4J1 | EFHD1_MOUSE | DGFIDLM[+16]ELK | 3.36 | 1.69E-03 | 0.0143 | 3 |
| Q8R429 | AT2A1_MOUSE | VDQSILTGESVSVIK | 2.15 | 1.76E-03 | 0.0143 | 3 |
| P63037 | DNJA1_MOUSE | NVVHQLSVTLEDLYNGATR | 7.59 | 1.96E-03 | 0.0143 | 3 |
| A2N1N1 | A2N1N1_MOUSE | NYLAWYQQKPGQSPK | 6.93 | 2.39E-03 | 0.0143 | 3 |
| P63276 | RS17_MOUSE 4 | LLDFGSLSNLQVTQPTVGMNFK | 3.41 | 2.57E-03 | 0.0143 | 3 |
| P14094 | AT1B1_MOUSE | VAPPGLTQIPQIQK | 2.17 | 2.67E-03 | 0.0143 | 3 |
| Q6PGN3 | DCLK2_MOUSE | SLSDNVNLPQGVR | 4.78 | 2.75E-03 | 0.0143 | 3 |
| O08788 | DCTN1_MOUSE | GPPPSGIATLVSGIAGEEPQR | 2.58 | 2.80E-03 | 0.0143 | 3 |
| Q9JM76 | ARPC3_MOUSE | LIGNM[+16]ALLPLR | 2.71 | 2.86E-03 | 0.0143 | 3 |
| Q9D4J1 | EFHD1_MOUSE | LGAPQTHLGLK | 4.28 | 2.86E-03 | 0.0143 | 3 |
| Q91YR1 | TWF1_MOUSE | SPLLEIVER | 4.34 | 3.00E-03 | 0.0143 | 3 |
| Q91YR1 | TWF1_MOUSE | YLLSQSSPAPLTAAEEELR | 3.30 | 3.10E-03 | 0.0143 | 3 |
| Q60930 | VDAC2_MOUSE | VNNSSLIGVGYTQTLRPGVK | 3.31 | 3.24E-03 | 0.0143 | 3 |
| P62880 | GBB2_MOUSE ( | LIIWDSYTTNK | 2.14 | 3.32E-03 | 0.0143 | 3 |
| P63276 | RS17_MOUSE 4 | LLDFGSLSNLQVTQPTVGM[+16]NFK | 2.21 | 3.33E-03 | 0.0143 | 3 |
| P62631 | EF1A2_MOUSE | VETGILRPGMVVTFAPVNITTEVK | 2.17 | 3.45E-03 | 0.0144 | 3 |
| O70172 | PI42A_MOUSE | DVEFLAQLK | 2.87 | 3.49E-03 | 0.0144 | 3 |
| O70172 | PI42A_MOUSE | FLDFIGHIL | 5.93 | 3.51E-03 | 0.0144 | 3 |
| Q9Z2X1 | HNRPF_MOUSE | VHIEIGPDGR | 2.62 | 3.69E-03 | 0.0147 | 3 |
| P62880 | GBB2_MOUSE ( | LLVSASQDGK | 2.57 | 3.73E-03 | 0.0147 | 3 |
| Q6PGN3 | DCLK2_MOUSE | GGDLFDAITSSTK | 3.24 | 3.77E-03 | 0.0147 | 3 |
| Q9WTM5 | RUVB2_MOUSE | AVLIAGQPGTGK | 2.17 | 4.09E-03 | 0.0148 | 3 |
| P63037 | DNJA1_MOUSE | ITFHGEGDQEPGLEPGDIIIVLDQK | 6.72 | 4.09E-03 | 0.0148 | 3 |
| Q68FG2 | Q68FG2_MOUSE | LLDPEDVNVDQPDEK | 2.42 | 4.11E-03 | 0.0148 | 3 |
| P48318 | DCE1_MOUSE ( | M[+16]VISNPAATQSDIDFLIEEIER | 4.94 | 4.36E-03 | 0.0150 | 3 |
| Q8R429 | AT2A1_MOUSE | TASEMVLADDNFSTIVAAVEEGR | 3.61 | 4.37E-03 | 0.0150 | 3 |
| Q2PFD7 | PSD3_MOUSE I | IGSTTNPFLDIPHDPNAAVYK | 2.18 | 4.63E-03 | 0.0153 | 3 |
| Q68FG2 | Q68FG2_MOUSE | VPTLEQHYEELQAR | 1.82 | 4.90E-03 | 0.0155 | 3 |
| Q60930 | VDAC2_MOUSE | LTLSALVDGK | 2.04 | 4.92E-03 | 0.0155 | 3 |
| Q8K0S0 | PHYIP_MOUSE | FLTC[+57]SVEDGELIFR | 2.15 | 4.97E-03 | 0.0155 | 3 |
| P28663 | SNAB_MOUSE | LDQWLTTMLLR | 3.65 | 5.16E-03 | 0.0155 | 3 |
| Q9JM76 | ARPC3_MOUSE | LIGNMALLPLR | 3.81 | 5.28E-03 | 0.0157 | 3 |
| Q8K0S0 | PHYIP_MOUSE | TEYSVAVQTAVK | 1.80 | 5.42E-03 | 0.0158 | 3 |
| Q9JM76 | ARPC3_MOUSE | DTDIVDEAIYYFK | 4.17 | 5.55E-03 | 0.0159 | 3 |
| Q99KX1 | MLF2_MOUSE I | QDYINLDESEAAAFDDEWRR | 5.31 | 5.89E-03 | 0.0163 | 3 |
| Q2PFD7 | PSD3_MOUSE I | GTVLYLQK | 1.59 | 6.16E-03 | 0.0166 | 3 |
| Q2M3X8 | PHAR1_MOUSE | ISFNLGAAEEVER | 2.81 | 6.27E-03 | 0.0168 | 3 |
| Q9JIS5 | SV2A_MOUSE S | DREELAQQYETILR | 4.19 | 6.61E-03 | 0.0171 | 3 |
| Q68FG2 | Q68FG2_MOUSE | QLANSLSGVQNQLQSFNSYR | 1.10 | 7.00E-03 | 0.0175 | 3 |
| Q9CR57 | RL14_MOUSE 6 | AAIAAAAAAAAAK | 5.75 | 7.41E-03 | 0.0179 | 3 |
| P28663 | SNAB_MOUSE | AALC[+57]HFIVDELNAK | 1.98 | 7.54E-03 | 0.0180 | 3 |
| P61358 | RL27_MOUSE 6 | YSVDIPLDK | 1.14 | 7.64E-03 | 0.0181 | 3 |
| A0A0B6VMB | A0A0B6VMB2_MC | DDPEVQFSWFVDDVEVHTAQTKPR | 3.84 | 7.74E-03 | 0.0182 | 3 |
| P14094 | AT1B1_MOUSE | YNPNVLPVQC[+57]TGK | 1.69 | 7.75E-03 | 0.0182 | 3 |
| P62281 | RS11_MOUSE 4 | C[+57]PFTGNVSIR | 3.51 | 7.80E-03 | 0.0182 | 3 |
| Q9D4J1 | EFHD1_MOUSE | DGFIDLMELK | 4.55 | 8.50E-03 | 0.0189 | 3 |
| P29515 | TBB7_ARATH Tub | NSSYFVEWIPNNVK | 3.05 | 8.97E-03 | 0.0195 | 3 |
| Q99KX1 | MLF2_MOUSE I | MLSGGFGYSPFLSITDGNMPATRPASR | 4.49 | 9.56E-03 | 0.0202 | 3 |
| Q60930 | VDAC2_MOUSE | LTFDTTFSPNTGK | 2.51 | 9.71E-03 | 0.0204 | 3 |
| P62245 | RS15A_MOUSE | IVVNLTGR | 2.70 | 9.81E-03 | 0.0204 | 3 |
| A2N1N1 | A2N1N1_MOUSE | LLLYFASTR | 4.05 | 9.86E-03 | 0.0205 | 3 |
| P63037 | DNJA1_MOUSE | QISQAYEVLADSK | 1.93 | 1.01E-02 | 0.0207 | 3 |
| P63276 | RS17_MOUSE 4 | DNYVPEVSALDQEIIEVDPDTK | 4.62 | 1.07E-02 | 0.0214 | 3 |
| Q99PT1 | GDIR1_MOUSE | AEEYEFLTPMEEAPK | 1.74 | 1.07E-02 | 0.0214 | 3 |

| Protein Accession | Protein Description | Modified Peptide Sequence | co-IP:Ctrl | p value | q value | Sig Peptide Count |
|---|---|---|---|---|---|---|
| P62245 | RS15A_MOUSE | HGYIGEFEIIDDHR | 2.70 | 1.08E-02 | 0.0215 | 3 |
| Q99PT1 | GDIR1_MOUSE | SIQEIQELDKDDESLRK | 1.08 | 1.10E-02 | 0.0217 | 3 |
| Q9JIS5 | SV2A_MOUSE S | GGLSDGEGPPGGR | 1.40 | 1.12E-02 | 0.0219 | 3 |
| Q5PR73 | DIRA2_MOUSE | EVQSSEAEALAR | 1.52 | 1.16E-02 | 0.0224 | 3 |
| Q99KK2 | NEUA_MOUSE | LAGVPLIGWVLR | 3.55 | 1.18E-02 | 0.0226 | 3 |
| P62245 | RS15A_MOUSE | QFGFIVLTTSAGIMDHEEAR | 2.72 | 1.22E-02 | 0.0230 | 3 |
| P61358 | RL27_MOUSE 6 | NIDDGTSDRPYSHALVAGIDR | 6.51 | 1.24E-02 | 0.0231 | 3 |
| P62631 | EF1A2_MOUSE | VETGILRPGM[+16]VVTFAPVNITTEVK | 1.05 | 1.25E-02 | 0.0232 | 3 |
| Q6PGN3 | DCLK2_MOUSE | VIGDGNFAVVK | 2.45 | 1.25E-02 | 0.0232 | 3 |
| Q99KK2 | NEUA_MOUSE | GLEKPPHLAALVLAR | 2.25 | 1.26E-02 | 0.0233 | 3 |
| P62631 | EF1A2_MOUSE | YYITIIDAPGHR | 1.40 | 1.26E-02 | 0.0233 | 3 |
| P62889 | RL30_MOUSE 6 | LVILANNC[+57]PALR | 4.66 | 1.33E-02 | 0.0241 | 3 |
| Q5PR73 | DIRA2_MOUSE | VAVFGAGGVGK | 1.46 | 1.34E-02 | 0.0242 | 3 |
| P62889 | RL30_MOUSE 6 | VC[+57]TLAIIDPGDSDIIR | 4.53 | 1.43E-02 | 0.0253 | 3 |
| Q8BFR5 | EFTU_MOUSE I | GITINAAHVEYSTAR | 1.66 | 1.53E-02 | 0.0264 | 3 |
| Q9JIS5 | SV2A_MOUSE S | AAPILFASAALALGSSLALK | 3.15 | 1.56E-02 | 0.0267 | 3 |
| P62880 | GBB2_MOUSE ( | TFVSGAC[+57]DASIK | 1.80 | 1.59E-02 | 0.0270 | 3 |
| Q9Z2X1 | HNRPF_MOUSE | ITGEAFVQFASQELAEK | 5.72 | 1.66E-02 | 0.0277 | 3 |
| Q04447 | KCRB_MOUSE | TDLNPDNLQGGDDLDPNYVLSSR | 5.11 | 1.71E-02 | 0.0283 | 3 |
| O88544 | CSN4_MOUSE ( | VISFEEQVASIR | 2.55 | 1.74E-02 | 0.0285 | 3 |
| P68404 | KPCB_MOUSE | EVLIVVVR | 4.20 | 1.74E-02 | 0.0285 | 3 |
| Q9Z0E0 | NCDN_MOUSE | IFDAVGFTFPNR | 5.51 | 1.75E-02 | 0.0287 | 3 |
| P63038 | CH60_MOUSE 6 | NAGVEGSLIVEK | 1.49 | 1.81E-02 | 0.0294 | 3 |
| Q64332 | SYN2_MOUSE S | FPLIEQTYYPNHR | 4.55 | 1.87E-02 | 0.0300 | 3 |
| P12658 | CALB1_MOUSE | LLPVQENFLLK | 3.72 | 1.93E-02 | 0.0307 | 3 |
| P16125 | LDHB_MOUSE I | MVVDSAYEVIK | 2.39 | 1.95E-02 | 0.0309 | 3 |
| P60335 | PCBP1_MOUSE | QVTITGSAASISLAQYLINAR | 1.10 | 2.05E-02 | 0.0322 | 3 |
| Q9D8N0 | EF1G_MOUSE I | TFLVGER | 2.38 | 2.07E-02 | 0.0324 | 3 |
| E9PV24 | FIBA_MOUSE F | GLIDEANQDFTNR | 6.81 | 2.08E-02 | 0.0325 | 3 |
| Q6PIC6 | AT1A3_MOUSE | KYNTDC[+57]VQGLTHSK | 6.66 | 2.21E-02 | 0.0341 | 3 |
| Q9Z1N5 | DX39B_MOUSE | ILVATNLFGR | 3.06 | 2.23E-02 | 0.0343 | 3 |
| P11499 | HS90B_MOUSE | YHTSQSGDEMTSLSEYVSR | 3.33 | 2.30E-02 | 0.0350 | 3 |
| P63038 | CH60_MOUSE 6 | AAVEEGIVLGGGC[+57]ALLR | 5.66 | 2.36E-02 | 0.0357 | 3 |
| P62962 | PROF1_MOUSE | DSLLQDGEFTMDLR | 6.59 | 2.59E-02 | 0.0381 | 3 |
| P61021 | RAB5B_MOUSE | TAMNVNDLFLAIAK | 6.81 | 2.75E-02 | 0.0398 | 3 |
| P14873 | MAP1B_MOUSE | IAELEEER | 6.29 | 2.75E-02 | 0.0398 | 3 |
| Q3UH68 | LIMC1_MOUSE | TGLENGILLC[+57]ELLNAIKPGLVK | 2.24 | 2.81E-02 | 0.0405 | 3 |
| Q8VDD5 | MYH9_MOUSE | DLGEELEALKTELEDTLDSTAAQQELR | 2.10 | 2.97E-02 | 0.0422 | 3 |
| P48774 | GSTM5_MOUSE | SMVLGYWDIR | 3.80 | 3.13E-02 | 0.0440 | 3 |
| P52480 | KPYM_MOUSE | EKGADFLVTEVENGGSLGSK | 6.46 | 3.30E-02 | 0.0461 | 3 |
| Q6R891 | NEB2_MOUSE I | ETQAQYQALER | 5.83 | 3.44E-02 | 0.0478 | 3 |
| Q01853 | TERA_MOUSE T | WALSQSNPSALR | 4.42 | 3.58E-02 | 0.0494 | 3 |
| Q9Z1N5 | DX39B_MOUSE | C[+57]IALAQLLVEQNFPAIAIHR | 4.69 | 2.06E-04 | 0.0143 | 3 |
| P27546 | MAP4_MOUSE | GMVSLSEIEEALAK | 4.37 | 3.36E-04 | 0.0143 | 3 |
| P43006 | EAA2_MOUSE E | NDEVSSLDAFLDLIR | 3.22 | 3.94E-04 | 0.0143 | 3 |
| E9PV24 | FIBA_MOUSE F | SQLQEAPPEWK | 3.35 | 1.22E-03 | 0.0143 | 3 |
| Q8R4U7 | LUZP1_MOUSE | GQVPGHASQGTQAVESSC[+57]SK | 5.41 | 1.37E-03 | 0.0143 | 3 |
| Q9DBR7 | MYPT1_MOUSE | STQGVTLTDLQEAEK | 2.39 | 1.55E-03 | 0.0143 | 3 |
| P43006 | EAA2_MOUSE E | KNDEVSSLDAFLDLIR | 3.95 | 1.64E-03 | 0.0143 | 3 |
| P80313 | TCPH_MOUSE T | SQDAEVGDGTTSVTLLAAEFLK | 2.33 | 1.76E-03 | 0.0143 | 3 |
| Q9D394 | RUFY3_MOUSE | LTEELAVANNR | 2.60 | 1.85E-03 | 0.0143 | 3 |
| P62754 | RS6_MOUSE 40 | DIPGLTDTTVPR | 4.48 | 1.89E-03 | 0.0143 | 3 |
| Q8R4U7 | LUZP1_MOUSE | IEDGISSTLSSK | 2.55 | 1.91E-03 | 0.0143 | 3 |
| P27546 | MAP4_MOUSE | SPATTLPK | 2.01 | 2.01E-03 | 0.0143 | 3 |
| P62082 | RS7_MOUSE 40 | TLTAVHDAILEDLVFPSEIVGK | 3.38 | 2.24E-03 | 0.0143 | 3 |
| P19096 | FAS_MOUSE Fa | LLLPEDPLISGLLNSQALK | 2.63 | 2.27E-03 | 0.0143 | 3 |
| P80313 | TCPH_MOUSE T | IALLNVELELK | 3.92 | 2.52E-03 | 0.0143 | 3 |
| P62962 | PROF1_MOUSE | DSLLQDGEFTM[+16]DLR | 4.02 | 2.88E-03 | 0.0143 | 3 |
| P63085 | MK01_MOUSE I | FRHENIIGINDIIR | 3.84 | 3.00E-03 | 0.0143 | 3 |
| P63085 | MK01_MOUSE I | VADPDHDHTGFLTEYVATR | 3.04 | 3.04E-03 | 0.0143 | 3 |
| P27546 | MAP4_MOUSE | ATSPSTLVSTGPSSR | 2.96 | 3.61E-03 | 0.0145 | 3 |
| P19096 | FAS_MOUSE Fa | SYIITGGLGGFGLELAR | 4.86 | 3.71E-03 | 0.0147 | 3 |

| Protein Accession | Protein Description | Modified Peptide Sequence | co-IP:Ctrl | p value | q value | Sig Peptide Count |
|---|---|---|---|---|---|---|
| P80315 | TCPD_MOUSE | ALIAGGGAPEIELALR | 1.64 | 3.85E-03 | 0.0147 | 3 |
| Q9D394 | RUFY3_MOUSE | LVPEAAEITASVK | 2.08 | 4.37E-03 | 0.0150 | 3 |
| E9PV24 | FIBA_MOUSE F | GDFANANNFDNTYGQVSEDLRR | 2.09 | 4.46E-03 | 0.0151 | 3 |
| P62270 | RS18_MOUSE 4 | VLNTNIDGR | 3.43 | 4.65E-03 | 0.0153 | 3 |
| Q8K310 | MATR3_MOUSE | DLSAAGIGLLAAATQSLSM[+16]PASLGR | 4.97 | 4.66E-03 | 0.0153 | 3 |
| P62082 | RS7_MOUSE 4( | DVNFEFPEFQL | 2.70 | 4.66E-03 | 0.0153 | 3 |
| P80315 | TCPD_MOUSE | IGLIQFC[+57]LSAPK | 2.02 | 4.73E-03 | 0.0153 | 3 |
| Q64531 | HPRT_MUSSP Hy | FFADLLDYIK | 2.60 | 4.98E-03 | 0.0155 | 3 |
| Q62420 | SH3G2_MOUSE | GPGYPQAEALLAEAMLK | 1.85 | 5.07E-03 | 0.0155 | 3 |
| Q62420 | SH3G2_MOUSE | ALYDFEPENEGELGFK | 2.15 | 5.27E-03 | 0.0157 | 3 |
| Q62420 | SH3G2_MOUSE | QAVQILQQVTVR | 1.48 | 5.46E-03 | 0.0158 | 3 |
| Q64531 | HPRT_MUSSP Hy | TM[+16]QTLLSLVK | 1.93 | 5.49E-03 | 0.0159 | 3 |
| P63325 | RS10_MOUSE 4 | IAIYELLFK | 2.53 | 5.67E-03 | 0.0160 | 3 |
| Q9JII6 | AK1A1_MOUSE | GLEVTAYSPLGSSDR | 1.94 | 5.74E-03 | 0.0161 | 3 |
| Q8BRT1 | CLAP2_MOUSE | VLNTGSDVEEAVADALLLGDIR | 7.18 | 5.79E-03 | 0.0162 | 3 |
| Q64531 | HPRT_MUSSP Hy | NVLIVEDIIDTGK | 1.35 | 7.33E-03 | 0.0178 | 3 |
| Q9DBR7 | MYPT1_MOUSE | TGSYGALAEISASK | 3.31 | 7.38E-03 | 0.0178 | 3 |
| Q8K310 | MATR3_MOUSE | YQLLQLVEPFGVISNHLILNK | 8.25 | 7.63E-03 | 0.0181 | 3 |
| Q8BRT1 | CLAP2_MOUSE | EGLLGLQNLLK | 4.67 | 7.71E-03 | 0.0181 | 3 |
| P35979 | RL12_MOUSE 6 | HSGNITFDEIVNIAR | 3.36 | 9.01E-03 | 0.0196 | 3 |
| P62270 | RS18_MOUSE 4 | IPDWFLNR | 4.37 | 9.14E-03 | 0.0197 | 3 |
| P63325 | RS10_MOUSE 4 | AEAGAGSATEFQFR | 3.09 | 1.02E-02 | 0.0208 | 3 |
| P80313 | TCPH_MOUSE | EGTDSSQGIPQLVSNISAC[+57]QVIAEAVR | 6.51 | 1.12E-02 | 0.0219 | 3 |
| Q9DCH4 | EIF3F_MOUSE | VIGLSSDLQQVGGASAR | 4.52 | 1.20E-02 | 0.0228 | 3 |
| Q9DCH4 | EIF3F_MOUSE | VIGTLLGTVDK | 4.66 | 1.21E-02 | 0.0229 | 3 |
| P43006 | EAA2_MOUSE E | NLFPENLVQAC[+57]FQQIQTVTK | 2.36 | 1.22E-02 | 0.0230 | 3 |
| Q8BRT1 | CLAP2_MOUSE | TILLLLLETLGDKEPTIR | 6.59 | 1.25E-02 | 0.0232 | 3 |
| P62082 | RS7_MOUSE 4( | AIIIFVPVPQLK | 2.09 | 1.26E-02 | 0.0233 | 3 |
| P35979 | RL12_MOUSE 6 | EILGTAQSVGC[+57]NVDGR | 3.12 | 1.40E-02 | 0.0249 | 3 |
| P62754 | RS6_MOUSE 4( | LIEVDDER | 3.32 | 1.44E-02 | 0.0254 | 3 |
| P63085 | MK01_MOUSE I | LKELIFEETAR | 2.20 | 1.48E-02 | 0.0258 | 3 |
| P62962 | PROF1_MOUSE | TFVSITPAEVGVLVGK | 1.45 | 1.50E-02 | 0.0261 | 3 |
| Q9D394 | RUFY3_MOUSE | IITLQEEMER | 3.88 | 1.55E-02 | 0.0266 | 3 |
| Q9JII6 | AK1A1_MOUSE | HPDEPVLLEEPVVLALAEK | 1.51 | 1.61E-02 | 0.0272 | 3 |
| Q2M3X8 | PHAR1_MOUSE | LLDVESAQR | 1.42 | 1.96E-02 | 0.0310 | 3 |
| P52480 | KPYM_MOUSE | VNLAM[+16]DVGK | 3.57 | 2.02E-02 | 0.0318 | 3 |
| Q8VDD5 | MYH9_MOUSE | M[+16]QQNIQELEEQLEEEESAR | 4.35 | 2.03E-02 | 0.0319 | 3 |
| P26039 | TLN1_MOUSE T | TEDSGLQTQVIAAATQC[+57]ALSTSQLVAC[+57]TK | 3.18 | 2.07E-02 | 0.0324 | 3 |
| Q9JMH9 | MY18A_MOUSE | VVHFAEPGAGTK | 1.22 | 2.22E-02 | 0.0342 | 3 |
| P28652 | KCC2B_MOUSE | NLINQM[+16]LTINPAK | 8.06 | 2.26E-02 | 0.0346 | 3 |
| P63318 | KPCG_MOUSE | FEAC[+57]NYPLELYER | 3.98 | 2.28E-02 | 0.0348 | 3 |
| P17710 | HXK1_MOUSE I | M[+16]PLGFTFSFPC[+57]K | 2.27 | 2.40E-02 | 0.0361 | 3 |
| P07901 | HS90A_MOUSE | VILHLKEDQTEYLEER | 0.97 | 2.54E-02 | 0.0375 | 3 |
| Q62261 | SPTB2_MOUSE | FESLEPEMNNQASR | 3.65 | 2.58E-02 | 0.0380 | 3 |
| P12970 | RL7A_MOUSE 6 | TC[+57]TTVAFTQVNSEDKGALAK | 3.98 | 2.61E-02 | 0.0384 | 3 |
| Q99104 | MYO5A_MOUSE | RAATIVIQSYLR | 6.03 | 2.79E-02 | 0.0403 | 3 |
| P52480 | KPYM_MOUSE | EATESFASDPILYRPVAVALDTK | 4.27 | 3.02E-02 | 0.0428 | 3 |
| Q71LX4 | TLN2_MOUSE T | VMVTNVTSLLK | 1.29 | 3.18E-02 | 0.0446 | 3 |
| P84099 | RL19_MOUSE 6 | VWLDPNETNEIANANSR | 1.33 | 3.58E-02 | 0.0494 | 3 |
| Q5SQX6 | CYFP2_MOUSE | DFVSEAYLLTLGK | 2.88 | 1.44E-03 | 0.0143 | 3 |
| Q5SQX6 | CYFP2_MOUSE | NTIYAALQDFAQVTLR | 3.39 | 2.46E-03 | 0.0143 | 3 |
| Q5SQX6 | CYFP2_MOUSE | NVLISVLQAIR | 2.64 | 2.95E-03 | 0.0143 | 3 |
| P17742 | PPIA_MOUSE F | IIPGFM[+16]C[+57]QGGDFTR | 1.67 | 4.35E-03 | 0.0150 | 3 |
| P02088 | HBB1_MOUSE I | YFDSFGDLSSASAIMGNAK | 1.88 | 4.42E-03 | 0.0151 | 3 |
| P17742 | PPIA_MOUSE F | VSFELFADK | 1.25 | 1.00E-02 | 0.0206 | 3 |
| Q6ZWN5 | RS9_MOUSE 4( | LFEGNALLR | 2.71 | 1.25E-02 | 0.0232 | 3 |
| P47963 | RL13_MOUSE 6 | STESLQANVQR | 1.52 | 1.89E-02 | 0.0302 | 3 |
| P60710 | ACTB_MOUSE | DIKEKLC[+57]YVALDFEQEMATAASSSSLEK | 1.14 | 1.99E-02 | 0.0314 | 3 |
| P17742 | PPIA_MOUSE F | KITISDC[+57]GQL | 4.64 | 2.00E-02 | 0.0316 | 3 |
| P20357 | MTAP2_MOUSE | ADQGLDFAATK | 1.32 | 2.10E-02 | 0.0328 | 3 |
| O08599 | STXB1_MOUSE | MTDIMTEGITIVEDINK | 2.11 | 2.51E-02 | 0.0372 | 3 |
| P28660 | NCKP1_MOUSE | NLITDIC[+57]TEQC[+57]TLSDQLLPK | 3.05 | 2.55E-04 | 0.0143 | 3 |

| Protein Accession | Protein Description | Modified Peptide Sequence | co-IP:Ctrl | p value | q value | Sig Peptide Count |
|---|---|---|---|---|---|---|
| A2AJI0 | MA7D1_MOUSE | TAEGLLPFAEAEAFLK | 6.66 | 2.40E-03 | 0.0143 | 3 |
| P28660 | NCKP1_MOUSE | AINQIAAALFTIHK | 2.71 | 5.91E-03 | 0.0163 | 3 |
| A2AJI0 | MA7D1_MOUSE | ETAANNSGPDPVK | 5.24 | 7.01E-03 | 0.0175 | 3 |
| P48453 | PP2BB_MOUSE | SQTTGFPSLITIFSAPNYLDVYNNK | 2.94 | 8.76E-03 | 0.0192 | 3 |
| P48453 | PP2BB_MOUSE | YLFLGDYVDR | 1.48 | 9.72E-03 | 0.0204 | 3 |
| P00920 | CAH2_MOUSE | SIVNNGHSFNVEFDDSQDNAVLK | 1.74 | 1.34E-02 | 0.0242 | 3 |
| Q99104 | MYO5A_MOUSE | KTDDDAEAIC[+57]SM[+16]C[+57]NALTTAQIVK | 1.45 | 2.19E-02 | 0.0338 | 3 |
| Q68FD5 | CLH1_MOUSE | QNLQIC[+57]VQVASK | 2.44 | 2.56E-02 | 0.0377 | 3 |
| A0A0N5DP62 | A0A0N5DP62_TRI | GAILTTM[+16]IATR | 1.44 | 2.62E-02 | 0.0385 | 3 |
| Q61598 | GDIB_MOUSE F | VPSTEAEALASSLM[+16]GLFEK | 2.69 | 2.67E-02 | 0.0390 | 3 |
| Q9WV60 | GSK3B_MOUSE | DIKPQNLLLDPDTAVLK | 1.26 | 3.05E-02 | 0.0431 | 3 |
| P20029 | BIP_MOUSE En | ITPSYVAFTPEGER | 2.30 | 1.79E-03 | 0.0143 | 3 |
| Q8VEK3 | HNRPU_MOUSE | YNILGTNTIMDK | 1.50 | 1.88E-03 | 0.0143 | 3 |
| Q8VEK3 | HNRPU_MOUSE | EKPYFPIPEDC[+57]TFIQNVPLEDR | 2.99 | 4.72E-03 | 0.0153 | 3 |
| P20029 | BIP_MOUSE En | IINEPTAAAIAYGLDK | 2.50 | 6.64E-03 | 0.0172 | 3 |
| P20029 | BIP_MOUSE En | VEIIANDQGNR | 1.89 | 6.69E-03 | 0.0172 | 3 |
| Q8VEK3 | HNRPU_MOUSE | NFILDQTNVSAAAQR | 4.69 | 6.76E-03 | 0.0172 | 3 |
| Q80TJ1 | CAPS1_MOUSE | LDALQTFIR | 1.55 | 7.93E-03 | 0.0183 | 3 |
| Q80TJ1 | CAPS1_MOUSE | FVTILEGVLAK | 1.60 | 9.72E-03 | 0.0204 | 3 |
| Q80TJ1 | CAPS1_MOUSE | DVLGSAASGAR | 2.15 | 1.04E-02 | 0.0211 | 3 |
| P63011 | RAB3A_MOUSE | YADDSFTPAFVSTVGIDFK | 2.36 | 1.30E-03 | 0.0143 | 3 |
| P61922 | GABT_MOUSE | TLLTGLLDLQAQYPQFISR | 5.04 | 2.48E-03 | 0.0143 | 3 |
| P61922 | GABT_MOUSE | NLLLAEVINIIK | 2.73 | 2.65E-03 | 0.0143 | 3 |
| P63011 | RAB3A_MOUSE | QLADHLGFEFFEASAK | 2.11 | 3.08E-03 | 0.0143 | 3 |
| P61922 | GABT_MOUSE | M[+16]LDLYSQISSVPIGYNHPALAK | 2.32 | 7.06E-03 | 0.0175 | 3 |
| P02088 | HBB1_MOUSE | KVITAFNDGLNHLDSLK | 1.48 | 1.89E-02 | 0.0302 | 3 |
| P16125 | LDHB_MOUSE | ITVVGVGQVGMAC[+57]AISILGK | 1.67 | 3.53E-03 | 0.0144 | 3 |
| P16125 | LDHB_MOUSE | GMYGIENEVFLSLPC[+57]ILNAR | 4.77 | 1.35E-02 | 0.0243 | 3 |
| Q9ERD7 | TBB3_MOUSE T | SGAFGHLFRPDNFIFGQSGAGNNWAK | 1.21 | 1.95E-02 | 0.0309 | 3 |
| P11798 | KCC2A_MOUSE | MC[+57]DPGM[+16]TAFEPEALGNLVEGLDFHR | 7.09 | 5.48E-03 | 0.0159 | 3 |
| P11798 | KCC2A_MOUSE | VTEQLIEAISNGDFESYTK | 4.53 | 8.55E-03 | 0.0190 | 3 |
| P35979 | RL12_MOUSE 6 | HPHDIIDDINSGAVEC[+57]PAS | 4.53 | 2.62E-02 | 0.0385 | 3 |
| Q8CBE3 | WDR37_MOUSE | YAGHVGSVNSIK | 2.81 | 7.91E-05 | 0.0143 | 2 |
| P32067 | LA_MOUSE Lup | LTTDFNVIVQALSK | 2.32 | 8.52E-05 | 0.0143 | 2 |
| Q9QZX7 | SRR_MOUSE S | DLVDDVFTVTEDEIK | 3.46 | 1.38E-04 | 0.0143 | 2 |
| Q9DCD0 | 6PGD_MOUSE | NPELQNLLLDDFFK | 3.01 | 2.20E-04 | 0.0143 | 2 |
| P13707 | GPDA_MOUSE | LGLMEMIAFAK | 2.71 | 3.05E-04 | 0.0143 | 2 |
| P29516 | TBB8_ARATH Tub | EVDEQMINVQNK | 2.24 | 3.72E-04 | 0.0143 | 2 |
| A0A0N5DKY8 | A0A0N5DKY8_TRI | AFM[+16]TADLPNELIEILEK | 4.22 | 5.52E-04 | 0.0143 | 2 |
| P29516 | TBB8_ARATH Tub | EVDEQM[+16]INVQNK | 1.76 | 7.20E-04 | 0.0143 | 2 |
| Q8H156 | RAN3_ARATH GT | VC[+57]ENIPIVLC[+57]GNK | 2.16 | 7.93E-04 | 0.0143 | 2 |
| O88456 | CPNS1_MOUSE | ILGGVISAISEAAAQYNPEPPPPR | 4.07 | 7.95E-04 | 0.0143 | 2 |
| Q7TPW1 | NEXN_MOUSE | LEINFEQLLR | 7.01 | 8.30E-04 | 0.0143 | 2 |
| F8VPU2 | FARP1_MOUSE | LGAPENSGISTLER | 2.72 | 8.41E-04 | 0.0143 | 2 |
| Q8C845 | Q8C845_MOUSE | AAAGELQEDSGLHVLAR | 4.24 | 9.13E-04 | 0.0143 | 2 |
| P61027 | RAB10_MOUSE | LQIWDTAGQER | 2.11 | 1.06E-03 | 0.0143 | 2 |
| Q7TPW1 | NEXN_MOUSE | NTSVVDSEPVR | 2.44 | 1.10E-03 | 0.0143 | 2 |
| P62827 | RAN_MOUSE G | SNYNFEKPFLWLAR | 3.48 | 1.22E-03 | 0.0143 | 2 |
| P11983 | TCPA_MOUSE | SLLVIPNTLAVNAAQDSTDLVAK | 1.97 | 1.25E-03 | 0.0143 | 2 |
| Q9ESN6 | TRIM2_MOUSE | ASLQVQLDAVNK | 2.97 | 1.29E-03 | 0.0143 | 2 |
| P49615 | CDK5_MOUSE | DLLQNLLK | 2.14 | 1.40E-03 | 0.0143 | 2 |
| G3UZJ2 | G3UZJ2_MOUSE | KTTASGDLAQAPGAFK | 6.14 | 1.43E-03 | 0.0143 | 2 |
| P31938 | MP2K1_MOUSE | LC[+57]DFGVSGQLIDSMANSFVGTR | 3.24 | 1.53E-03 | 0.0143 | 2 |
| Q80VD1 | FA98B_MOUSE | NSEIC[+57]QEVQAVC[+57]DALGVPK | 3.31 | 1.58E-03 | 0.0143 | 2 |
| Q84W47 | Q84W47_ARATH | APNM[+16]ETITESLEK | 5.15 | 1.60E-03 | 0.0143 | 2 |
| Q6P8J7 | KCRS_MOUSE | GWEFMWNERLGYILTC[+57]PSNLGTGLR | 5.96 | 1.66E-03 | 0.0143 | 2 |
| Q8CIE6 | COPA_MOUSE | SILLSVPLLVVDNK | 2.64 | 1.76E-03 | 0.0143 | 2 |
| Q8BP47 | SYNC_MOUSE | NLMFLVLR | 2.08 | 1.78E-03 | 0.0143 | 2 |
| Q9QXY6 | EHD3_MOUSE | FVC[+57]AQLPNAVLESISVIDTPGILSGEK | 4.26 | 1.79E-03 | 0.0143 | 2 |
| A0A178UNP6 | A0A178UNP6_AR | M[+16]ATPPLTPR | 3.06 | 1.90E-03 | 0.0143 | 2 |
| Q9DCL9 | PUR6_MOUSE | ITSC[+57]IFQLLQEAGIK | 3.08 | 1.91E-03 | 0.0143 | 2 |
| P60867 | RS20_MOUSE 4 | TPVEPEVAIHR | 2.68 | 1.95E-03 | 0.0143 | 2 |

| Protein Accession | Protein Description | Modified Peptide Sequence | co-IP:Ctrl | p value | q value | Sig Peptide Count |
|---|---|---|---|---|---|---|
| P53994 | RAB2A_MOUSE | GAAGALLVYDITR | 1.29 | 1.97E-03 | 0.0143 | 2 |
| Q9CPV4 | GLOD4_MOUSE | VTLAVSDLQK | 3.29 | 1.99E-03 | 0.0143 | 2 |
| O54946 | DNJB6_MOUSE | QVAEAYEVLSDAK | 4.39 | 2.11E-03 | 0.0143 | 2 |
| Q9CZ04 | CSN7A_MOUSE | QLEDLVIEAVYADVLR | 4.97 | 2.24E-03 | 0.0143 | 2 |
| G3UZJ2 | G3UZJ2_MOUSE | TTASGDLAQAPGAFK | 6.18 | 2.25E-03 | 0.0143 | 2 |
| Q8CBY8 | DCTN4_MOUSE | AGASISTLAGLSLR | 4.35 | 2.31E-03 | 0.0143 | 2 |
| A0A0N5DFM | A0A0N5DFM1_TR | NAPAILFIDEIDAIAPK | 2.11 | 2.32E-03 | 0.0143 | 2 |
| Q9CQE8 | RTRAF_MOUSE | HILGFDTGDAVLNEAAQILR | 8.32 | 2.38E-03 | 0.0143 | 2 |
| Q80Y86 | MK15_MOUSE I | LC[+57]DFGLAR | 1.72 | 2.45E-03 | 0.0143 | 2 |
| Q8CBE3 | WDR37_MOUSE | STLLELFGQIER | 4.57 | 2.46E-03 | 0.0143 | 2 |
| P53994 | RAB2A_MOUSE | TASNVEEAFINTAK | 1.52 | 2.50E-03 | 0.0143 | 2 |
| Q8C0E9 | Q8C0E9_MOUSE | ISNSSEFSAK | 2.64 | 2.57E-03 | 0.0143 | 2 |
| Q9QZE5 | COPG1_MOUSE | SIATLAITTLLK | 2.50 | 2.60E-03 | 0.0143 | 2 |
| P51881 | ADT2_MOUSE | YFPTQALNFAFK | 2.13 | 2.61E-03 | 0.0143 | 2 |
| Q80Y86 | MK15_MOUSE I | LLDVIPAK | 5.15 | 2.65E-03 | 0.0143 | 2 |
| Q9QXY6 | EHD3_MOUSE I | LFEAEEQDLFR | 4.86 | 2.76E-03 | 0.0143 | 2 |
| P48484 | PP14_ARATH Seri | IYGFYDEC[+57]K | 3.44 | 2.76E-03 | 0.0143 | 2 |
| P49615 | CDK5_MOUSE | SFLFQLLK | 2.84 | 2.85E-03 | 0.0143 | 2 |
| O88685 | PRS6A_MOUSE | VDILDPALLR | 2.01 | 2.87E-03 | 0.0143 | 2 |
| P58771 | TPM1_MOUSE | KLVIIESDLER | 5.14 | 2.89E-03 | 0.0143 | 2 |
| O88685 | PRS6A_MOUSE | TMLELLNQLDGFQPNTQVK | 3.47 | 2.90E-03 | 0.0143 | 2 |
| Q9Z204 | HNRPC_MOUSE | GFAFVQYVNER | 2.07 | 2.94E-03 | 0.0143 | 2 |
| P01831 | THY1_MOUSE | VTSLTAC[+57]LVNQNLR | 3.11 | 3.01E-03 | 0.0143 | 2 |
| Q9EPU0 | RENT1_MOUSE | LVVLGIRPIR | 2.25 | 3.07E-03 | 0.0143 | 2 |
| Q5SV64 | Q5SV64_MOUSE | ASFYDSVSGLHEPPVDR | 4.88 | 3.07E-03 | 0.0143 | 2 |
| P21107 | TPM3_MOUSE | KLVIIEGDLER | 2.50 | 3.10E-03 | 0.0143 | 2 |
| Q9Z1G4 | VPP1_MOUSE | FTHGFQNIVDAYGIGTYR | 3.33 | 3.13E-03 | 0.0143 | 2 |
| Q91WQ3 | SYYC_MOUSE | QVEHPLLSGLLYPGLQALDEEYLK | 5.96 | 3.19E-03 | 0.0143 | 2 |
| P70248 | MYO1F_MOUSE | DIILQSNPLLEAFGNAK | 5.86 | 3.20E-03 | 0.0143 | 2 |
| D3Z4J3 | D3Z4J3_MOUSE | LTNENLDLM[+16]EQLEK | 3.94 | 3.24E-03 | 0.0143 | 2 |
| P0CG50 | UBC_MOUSE P | TLSDYNIQK | 2.38 | 3.24E-03 | 0.0143 | 2 |
| Q80VD1 | FA98B_MOUSE | GPLLEEQALSK | 2.49 | 3.45E-03 | 0.0144 | 2 |
| Q9QZX7 | SRR_MOUSE S | LLIEPTAGVALAAVLSQHFQTVSPEVK | 4.41 | 3.48E-03 | 0.0144 | 2 |
| Q91WQ3 | SYYC_MOUSE | IDVGEAEPR | 1.90 | 3.60E-03 | 0.0145 | 2 |
| Q91Z69 | SRGP1_MOUSE | VQLLQDLQDFFR | 7.07 | 3.78E-03 | 0.0147 | 2 |
| Q6P1F6 | 2ABA_MOUSE | SFFSEIISSISDVK | 5.67 | 3.86E-03 | 0.0147 | 2 |
| P61027 | RAB10_MOUSE | AFLTLAEDILR | 2.13 | 3.92E-03 | 0.0147 | 2 |
| P0CG50 | UBC_MOUSE P | TITLEVEPSDTIENVK | 3.17 | 4.08E-03 | 0.0148 | 2 |
| Q6NS52 | DGKB_MOUSE | DIVC[+57]YLSLLER | 5.75 | 4.13E-03 | 0.0148 | 2 |
| Q91Z69 | SRGP1_MOUSE | VSGSQVEVNDIK | 3.98 | 4.36E-03 | 0.0150 | 2 |
| P60867 | RS20_MOUSE 4 | LIDLHSPSEIVK | 2.85 | 4.40E-03 | 0.0150 | 2 |
| Q9SIP7 | RS31_ARATH 40S | ELAEDGYSGVEVR | 1.78 | 4.40E-03 | 0.0150 | 2 |
| Q9EPN1 | NBEA_MOUSE | ATDAQLC[+57]LESSPK | 4.38 | 4.68E-03 | 0.0153 | 2 |
| P70248 | MYO1F_MOUSE | ISNFLLEK | 4.55 | 4.77E-03 | 0.0154 | 2 |
| A0A178UNP6 | A0A178UNP6_AR | MATPPLTPR | 3.98 | 5.10E-03 | 0.0155 | 2 |
| A0A0N5DF32 | A0A0N5DF32_TRI | EC[+57]LPLILFIR | 3.71 | 5.10E-03 | 0.0155 | 2 |
| Q9EPU0 | RENT1_MOUSE | NVFLLGFIPAK | 4.41 | 5.11E-03 | 0.0155 | 2 |
| Q9Z1W8 | AT12A_MOUSE | LIIVEGC[+57]QR | 1.67 | 5.13E-03 | 0.0155 | 2 |
| P51881 | ADT2_MOUSE | DFLAGGVAAAISK | 1.92 | 5.23E-03 | 0.0157 | 2 |
| P11983 | TCPA_MOUSE | LGVQVVITDPEKLDQIR | 2.95 | 5.32E-03 | 0.0158 | 2 |
| Q84W47 | Q84W47_ARATH U | VVGDGFGR | 1.37 | 5.33E-03 | 0.0158 | 2 |
| Q9ESN6 | TRIM2_MOUSE | VLQSQLDTLLQGQESIK | 2.40 | 5.40E-03 | 0.0158 | 2 |
| P31938 | MP2K1_MOUSE | LPSGVFSLEFQDFVNK | 4.80 | 5.72E-03 | 0.0161 | 2 |
| Q9DBZ5 | EIF3K_MOUSE | YNPENLATLER | 3.94 | 6.07E-03 | 0.0165 | 2 |
| P38647 | GRP75_MOUSE | AQFEGIVTDLIK | 1.81 | 6.28E-03 | 0.0168 | 2 |
| Q6P1F6 | 2ABA_MOUSE | VVIFQQEQENK | 1.90 | 6.37E-03 | 0.0169 | 2 |
| F8VPU2 | FARP1_MOUSE | SLVSQPTAPNSEVPK | 2.48 | 6.37E-03 | 0.0169 | 2 |
| P48486 | PP16_ARATH Seri | AHQVVEDGYEFFAK | 3.33 | 6.47E-03 | 0.0170 | 2 |
| P01872 | IGHM_MOUSE I | LVESGFTTDPVTIENK | 2.20 | 6.53E-03 | 0.0171 | 2 |
| P42932 | TCPQ_MOUSE | FAEAFEAIPR | 1.46 | 6.57E-03 | 0.0171 | 2 |
| M7NIQ0 | M7NIQ0_PNEMU | YPENFFILR | 3.97 | 6.81E-03 | 0.0172 | 2 |
| P48482 | PP12_ARATH Seri | YPENFFLLR | 3.97 | 6.81E-03 | 0.0172 | 2 |

| Protein Accession | Protein Description | Modified Peptide Sequence | co-IP:Ctrl | p value | q value | Sig Peptide Count |
|---|---|---|---|---|---|---|
| P58389 | PTPA_MOUSE S | VDDQVAIVFK | 1.31 | 6.91E-03 | 0.0174 | 2 |
| E9Q634 | MYO1E_MOUSE | VFDFLVDSINK | 4.60 | 6.96E-03 | 0.0174 | 2 |
| P21107 | TPM3_MOUSE T | IQLVEEELDR | 3.75 | 7.21E-03 | 0.0177 | 2 |
| P32067 | LA_MOUSE Lup | GSIFAVFDSIQSAK | 3.79 | 7.21E-03 | 0.0177 | 2 |
| M7NIQ0 | M7NIQ0_PNEMU S | IKYPENFFILR | 4.76 | 7.21E-03 | 0.0177 | 2 |
| P48482 | PP12_ARATH Seri | IKYPENFFLLR | 4.76 | 7.21E-03 | 0.0177 | 2 |
| P49722 | PSA2_MOUSE P | YNEDLELEDAIHTAILTLK | 3.97 | 7.31E-03 | 0.0178 | 2 |
| P46638 | RB11B_MOUSE | GAVGALLVYDIAK | 1.42 | 7.33E-03 | 0.0178 | 2 |
| P48484 | PP14_ARATH Seri | QSLETIC[+57]LLLAYK | 4.33 | 7.34E-03 | 0.0178 | 2 |
| P48486 | PP16_ARATH Seri | QSIETIC[+57]LLLAYK | 4.33 | 7.34E-03 | 0.0178 | 2 |
| Q9ESJ4 | SPN90_MOUSE | TIGAELMELVR | 5.83 | 7.57E-03 | 0.0180 | 2 |
| P67871 | CSK2B_MOUSE | YQQGDFGYC[+57]PR | 6.77 | 7.63E-03 | 0.0181 | 2 |
| Q91ZU6 | DYST_MOUSE I | TREEQVDGATEK | 2.96 | 7.72E-03 | 0.0181 | 2 |
| P62264 | RS14_MOUSE 4 | TPGPGAQSALR | 2.97 | 7.72E-03 | 0.0181 | 2 |
| P01831 | THY1_MOUSE T | VTLSNQPYIK | 2.48 | 8.01E-03 | 0.0184 | 2 |
| Q8C1B1 | CAMP2_MOUSE | SDANNFLILFR | 3.82 | 8.08E-03 | 0.0184 | 2 |
| Q3UGR5 | HDHD2_MOUSE | TFFLEALR | 1.31 | 8.35E-03 | 0.0188 | 2 |
| O88544 | CSN4_MOUSE C | LYNNITFEELGALLEIPAAK | 7.43 | 8.41E-03 | 0.0188 | 2 |
| Q8C854 | MYEF2_MOUSE | VGEVTYVELFK | 2.25 | 8.59E-03 | 0.0190 | 2 |
| Q91VA7 | Q91VA7_MOUSE | NIANPTAMLLSATNMLR | 3.82 | 8.64E-03 | 0.0191 | 2 |
| Q9ERK4 | XPO2_MOUSE I | LLQAFLER | 3.01 | 8.65E-03 | 0.0191 | 2 |
| Q6NS52 | DGKB_MOUSE | HPPVAILPLGTGNDLAR | 3.88 | 8.72E-03 | 0.0191 | 2 |
| Q571F3 | Q571F3_MOUSE | VSGELFAQAPVEQYPGIAVETVTDSSR | 2.24 | 8.77E-03 | 0.0192 | 2 |
| P14106 | C1QB_MOUSE C | VITNANENYEPR | 3.93 | 8.84E-03 | 0.0193 | 2 |
| P61294 | RAB6B_MOUSE | DSTVAVVVYDITNLNSFQQTSK | 2.51 | 9.22E-03 | 0.0198 | 2 |
| Q5SV64 | Q5SV64_MOUSE | FVAELWKDEIQTIQR | 5.88 | 9.26E-03 | 0.0198 | 2 |
| Q8C1B1 | CAMP2_MOUSE | DGTDGC[+57]ALAALIHFYC[+57]PAVVR | 6.99 | 9.29E-03 | 0.0199 | 2 |
| Q9DB20 | ATPO_MOUSE A | LGNTQGIISAFSTIMSVHR | 3.54 | 9.30E-03 | 0.0199 | 2 |
| Q9Z1W8 | AT12A_MOUSE | VAEIPFNSTNK | 1.23 | 9.41E-03 | 0.0201 | 2 |
| E9Q634 | MYO1E_MOUSE | VLQVSIGPGLPK | 6.41 | 9.51E-03 | 0.0202 | 2 |
| Q9DCD0 | 6PGD_MOUSE 6 | GILFVGSGVSGGEEGAR | 1.58 | 9.69E-03 | 0.0204 | 2 |
| P12960 | CNTN1_MOUSE | IFNIQLEDEGLYEC[+57]EAENIR | 4.80 | 9.77E-03 | 0.0204 | 2 |
| Q9DB72 | BTBDH_MOUSE | LVVTPASSGGDAAGVSFQK | 5.43 | 1.00E-02 | 0.0206 | 2 |
| D3Z4J3 | D3Z4J3_MOUSE | LTNENLDLMEQLEK | 5.38 | 1.02E-02 | 0.0208 | 2 |
| P58389 | PTPA_MOUSE S | LVALLDTLDR | 1.22 | 1.03E-02 | 0.0209 | 2 |
| A0A0N5DF32 | A0A0N5DF32_TRI | LREC[+57]LPLILFIR | 6.07 | 1.05E-02 | 0.0212 | 2 |
| Q8C854 | MYEF2_MOUSE | LGSTIFVANLDFK | 5.27 | 1.08E-02 | 0.0215 | 2 |
| P62264 | RS14_MOUSE 4 | IEDVTPIPSDSTR | 2.41 | 1.10E-02 | 0.0217 | 2 |
| O35737 | HNRH1_MOUSE | HTGPNSPDTANDGFVR | 1.76 | 1.12E-02 | 0.0219 | 2 |
| Q60676 | PPP5_MOUSE S | AFLEENQLDYIIR | 2.55 | 1.12E-02 | 0.0219 | 2 |
| P38647 | GRP75_MOUSE | QAASSLQQASLK | 1.90 | 1.15E-02 | 0.0223 | 2 |
| P12960 | CNTN1_MOUSE | VQVTSQEYSAR | 2.93 | 1.17E-02 | 0.0225 | 2 |
| P62717 | RL18A_MOUSE | DLTTAGAVTQC[+57]YR | 5.18 | 1.22E-02 | 0.0230 | 2 |
| P62267 | RS23_MOUSE 4 | VANVSLLALYK | 4.38 | 1.23E-02 | 0.0231 | 2 |
| Q60676 | PPP5_MOUSE S | TEC[+57]YGYALGDATR | 3.70 | 1.32E-02 | 0.0240 | 2 |
| Q8CBY8 | DCTN4_MOUSE | LIEYYQQLAQK | 2.50 | 1.34E-02 | 0.0242 | 2 |
| Q8C845 | Q8C845_MOUSE | KAAAGELQEDSGLHVLAR | 4.19 | 1.35E-02 | 0.0243 | 2 |
| P62855 | RS26_MOUSE 4 | DISEASVFDAYVLPK | 3.60 | 1.35E-02 | 0.0243 | 2 |
| O54946 | DNJB6_MOUSE | VEVEEDGQLK | 2.06 | 1.38E-02 | 0.0247 | 2 |
| P62267 | RS23_MOUSE 4 | ITAFVPNDGC[+57]LNFIEENDEVLVAGFGR | 3.56 | 1.38E-02 | 0.0247 | 2 |
| Q9CXY6 | ILF2_MOUSE In | ILPTLEAVAALGNK | 4.62 | 1.41E-02 | 0.0250 | 2 |
| P13707 | GPDA_MOUSE | LGIPMSVLMGANIASEVAEEK | 2.70 | 1.42E-02 | 0.0251 | 2 |
| Q9DB72 | BTBDH_MOUSE | ITWNVLFSPR | 3.62 | 1.43E-02 | 0.0253 | 2 |
| Q9ERK4 | XPO2_MOUSE I | FLESVEGNQNYPLLLLTLLEK | 4.96 | 1.45E-02 | 0.0255 | 2 |
| P42932 | TCPQ_MOUSE | AIAGTGANVIVTGGK | 1.56 | 1.55E-02 | 0.0266 | 2 |
| P41105 | RL28_MOUSE 6 | QTYSTEPNNLK | 5.17 | 1.55E-02 | 0.0266 | 2 |
| A0A0N5DFM1 | A0A0N5DFM1_TR | AVANETGAFFFLLNGPEIMSK | 4.67 | 1.57E-02 | 0.0268 | 2 |
| Q9EPN1 | NBEA_MOUSE | AVLEQFLSFAK | 1.19 | 1.58E-02 | 0.0269 | 2 |
| P46638 | RB11B_MOUSE | STIGVEFATR | 2.02 | 1.59E-02 | 0.0270 | 2 |
| Q8CIE6 | COPA_MOUSE | LVGQSIIAYLQK | 2.73 | 1.59E-02 | 0.0270 | 2 |
| P62855 | RS26_MOUSE 4 | NIVEAAAVR | 3.52 | 1.60E-02 | 0.0271 | 2 |
| P14106 | C1QB_MOUSE C | TINSPLRPNQVIR | 3.19 | 1.60E-02 | 0.0271 | 2 |

| Protein Accession | Protein Description | Modified Peptide Sequence | co-IP:Ctrl | p value | q value | Sig Peptide Count |
|---|---|---|---|---|---|---|
| P84099 | RL19_MOUSE 6 | LLADQAEAR | 2.73 | 1.67E-02 | 0.0278 | 2 |
| Q9SIP7 | RS31_ARATH 40S | GLC[+57]AIAQAESLR | 3.31 | 1.67E-02 | 0.0278 | 2 |
| P67871 | CSK2B_MOUSE | RPANQFVPR | 4.08 | 1.67E-02 | 0.0278 | 2 |
| Q9D051 | ODPB_MOUSE | VFLLGEEVAQYDGAYK | 3.25 | 1.73E-02 | 0.0284 | 2 |
| P08553 | NFM_MOUSE N | LRDDTEAAIR | 1.85 | 1.74E-02 | 0.0285 | 2 |
| P20357 | MTAP2_MOUSE | QSTEPSIVMPSIGLSAEPPAPK | 1.38 | 1.78E-02 | 0.0290 | 2 |
| P68510 | 1433F_MOUSE | NSVVEASEAAYK | 1.89 | 1.79E-02 | 0.0292 | 2 |
| Q99104 | MYO5A_MOUSE | GDDFETVSFWLSNTC[+57]R | 2.92 | 1.83E-02 | 0.0296 | 2 |
| Q9CZT8 | RAB3B_MOUSE | TYSWDNAQVILVGNK | 2.44 | 1.90E-02 | 0.0303 | 2 |
| Q8CAA7 | PGM2L_MOUSE | LIDALIENFLEPSK | 2.66 | 1.92E-02 | 0.0306 | 2 |
| P46460 | NSF_MOUSE V | DIEAM[+16]DPSILK | 4.68 | 1.92E-02 | 0.0306 | 2 |
| P27659 | RL3_MOUSE 60 | VAFSVAR | 1.17 | 1.97E-02 | 0.0311 | 2 |
| P16858 | G3P_MOUSE G | VVDLMAYMASK | 6.97 | 2.01E-02 | 0.0317 | 2 |
| P17183 | ENOG_MOUSE | LAMQEFMILPVGAESFR | 1.58 | 2.15E-02 | 0.0334 | 2 |
| Q6P8J7 | KCRS_MOUSE | LGYILTC[+57]PSNLGTGLR | 4.83 | 2.16E-02 | 0.0335 | 2 |
| Q9WUM4 | COR1C_MOUSE | VTWDSSFC[+57]AVNPR | 5.14 | 2.19E-02 | 0.0338 | 2 |
| P00920 | CAH2_MOUSE | AVQQPDGLAVLGIFLK | 1.37 | 2.19E-02 | 0.0338 | 2 |
| O70172 | PI42A_MOUSE | FGIDDQDFQNSLTR | 5.89 | 2.23E-02 | 0.0343 | 2 |
| P33175 | KIF5A_MOUSE | SLSALGNVISALAEGTK | 2.30 | 2.24E-02 | 0.0344 | 2 |
| Q8CGF6 | WDR47_MOUSE | AAYADLLTPLISK | 5.25 | 2.30E-02 | 0.0350 | 2 |
| P47757 | CAPZB_MOUSE | QMEKDETVSDC[+57]SPHIANIGR | 1.44 | 2.34E-02 | 0.0355 | 2 |
| Q9CR57 | RL14_MOUSE 6 | C[+57]MQLTDFILK | 1.56 | 2.37E-02 | 0.0358 | 2 |
| Q8CI94 | PYGB_MOUSE | TC[+57]AYTNHTVLPEALER | 3.71 | 2.44E-02 | 0.0365 | 2 |
| Q9D051 | ODPB_MOUSE | TYYMSAGLQPVPIVFR | 2.40 | 2.45E-02 | 0.0366 | 2 |
| Q91VA7 | Q91VA7_MOUSE | KLDLFANVVHVK | 3.93 | 2.45E-02 | 0.0366 | 2 |
| Q9QZE5 | COPG1_MOUSE | AIVDC[+57]IISIIEENSESK | 1.17 | 2.45E-02 | 0.0366 | 2 |
| P04370 | MBP_MOUSE M | DTGILDSIGR | 2.86 | 2.48E-02 | 0.0369 | 2 |
| Q8C1B7 | SEP11_MOUSE | SYELQESNVR | 1.50 | 2.48E-02 | 0.0369 | 2 |
| Q61598 | GDIB_MOUSE F | SPYLYPLYGLGELPQGFAR | 7.49 | 2.49E-02 | 0.0370 | 2 |
| P15105 | GLNA_MOUSE | C[+57]IEEAIDK | 1.31 | 2.53E-02 | 0.0374 | 2 |
| Q8JZQ9 | EIF3B_MOUSE | VDNAYWLWTFQGR | 2.15 | 2.54E-02 | 0.0375 | 2 |
| P14873 | MAP1B_MOUSE | MSISEGTVSDK | 1.54 | 2.59E-02 | 0.0381 | 2 |
| P00920 | CAH2_MOUSE | QSPVDIDTATAQHDPALQPLLISYDK | 3.80 | 2.63E-02 | 0.0386 | 2 |
| Q9JHU4 | DYHC1_MOUSE | DFPLNDLLSATELDK | 4.31 | 2.68E-02 | 0.0391 | 2 |
| P25444 | RS2_MOUSE 40 | LSIVPVR | 4.33 | 2.75E-02 | 0.0398 | 2 |
| P62281 | RS11_MOUSE 4 | RDYLHYIR | 2.14 | 2.75E-02 | 0.0398 | 2 |
| Q9WV54 | ASAH1_MOUSE | ESLDVYELDPK | 3.97 | 2.85E-02 | 0.0409 | 2 |
| Q920I9 | WDR7_MOUSE | LNIWNIADIAEK | 1.09 | 2.93E-02 | 0.0418 | 2 |
| Q8C1B7 | SEP11_MOUSE | STLM[+16]DTLFNTK | 1.14 | 2.99E-02 | 0.0425 | 2 |
| Q9R0Y5 | KAD1_MOUSE | KVNAEGTVDTVFSEVC[+57]TYLDSLK | 1.14 | 2.99E-02 | 0.0425 | 2 |
| Q6A087 | Q6A087_MOUSE | VATLEQSYEALC[+57]ELAATR | 3.23 | 3.05E-02 | 0.0431 | 2 |
| Q8BPN8 | DMXL2_MOUSE | NFFPIAGLEFSELPVTSPLGIAVIK | 4.53 | 3.26E-02 | 0.0456 | 2 |
| Q61316 | HSP74_MOUSE | NFTTEQVTAMLLSK | 1.18 | 3.28E-02 | 0.0458 | 2 |
| Q99L45 | IF2B_MOUSE E | TSFVNFTDIC[+57]K | 2.86 | 3.42E-02 | 0.0475 | 2 |
| P08249 | MDHM_MOUSE | IFGVTTLDIVR | 1.23 | 3.52E-02 | 0.0487 | 2 |
| Q8BL97 | SRSF7_MOUSE | NPPGFAFVEFEDPR | 3.96 | 3.56E-02 | 0.0492 | 2 |
| O08749 | DLDH_MOUSE | RPFTQNLGLEELGIELDPK | 2.99 | 1.51E-04 | 0.0143 | 2 |
| P54609 | CD48A_ARATH C | GVLFYGPPGC[+57]GK | 3.04 | 4.81E-04 | 0.0143 | 2 |
| P28651 | CAH8_MOUSE | AVTEILQDIQYK | 2.35 | 4.92E-04 | 0.0143 | 2 |
| D3YVF0 | AKAP5_MOUSE | VTVDHAEEATVGQAEEATVGQAEK | 2.11 | 7.51E-04 | 0.0143 | 2 |
| P97315 | CSRP1_MOUSE | GFGFGQGAGALVHSE | 1.77 | 1.03E-03 | 0.0143 | 2 |
| P35279 | RAB6A_MOUSE | LQLWDTAGQER | 2.11 | 1.06E-03 | 0.0143 | 2 |
| P97315 | CSRP1_MOUSE | GLESTTLADKDGEIYC[+57]K | 3.90 | 1.13E-03 | 0.0143 | 2 |
| Q7M6Y3 | PICAL_MOUSE | NTLFNLSNFLDK | 1.68 | 1.48E-03 | 0.0143 | 2 |
| Q3UM45 | PP1R7_MOUSE | AIENIDTLTNLESLFLGK | 3.37 | 1.51E-03 | 0.0143 | 2 |
| A0A0J9YUN4 | A0A0J9YUN4_MO | C[+57]VDMVVSELTSTIR | 2.06 | 1.78E-03 | 0.0143 | 2 |
| Q61644 | PACN1_MOUSE | GPQYGSLER | 2.99 | 1.92E-03 | 0.0143 | 2 |
| P14824 | ANXA6_MOUSE | DQAQEDAQVAAEILEIADTPSGDK | 2.57 | 2.01E-03 | 0.0143 | 2 |
| P62852 | RS25_MOUSE 4 | AALQELLSK | 2.20 | 2.64E-03 | 0.0143 | 2 |
| P48962 | ADT1_MOUSE | EQGFLSFWR | 2.27 | 2.77E-03 | 0.0143 | 2 |
| Q9CZ30 | OLA1_MOUSE | IPAFLNVVDIAGLVK | 1.99 | 2.92E-03 | 0.0143 | 2 |
| A0A0J9YUN4 | A0A0J9YUN4_MO | TGLFTPDLAFEATVK | 1.61 | 3.08E-03 | 0.0143 | 2 |

| Protein Accession | Protein Description | Modified Peptide Sequence | co-IP:Ctrl | p value | q value | Sig Peptide Count |
|---|---|---|---|---|---|---|
| P54609 | CD48A_ARATH Ce | IVSQLLTLMDGLK | 2.26 | 3.16E-03 | 0.0143 | 2 |
| O70435 | PSA3_MOUSE P | HVGMAVAGLLADAR | 1.75 | 3.28E-03 | 0.0143 | 2 |
| P55066 | NCAN_MOUSE | AALAELVALPC[+57]FFTLQPR | 4.05 | 3.36E-03 | 0.0143 | 2 |
| O08749 | DLDH_MOUSE | EANLAAAFGKPINF | 2.25 | 3.39E-03 | 0.0143 | 2 |
| P46735 | MYO1B_MOUSE | IFLLTNNNLLLADQK | 7.14 | 3.82E-03 | 0.0147 | 2 |
| D3YVF0 | AKAP5_MOUSE | TSEQYETLLIETASSLVK | 6.07 | 3.88E-03 | 0.0147 | 2 |
| P55066 | NCAN_MOUSE | ELGGEVFYVGPAR | 2.10 | 4.18E-03 | 0.0149 | 2 |
| Q91WK2 | EIF3H_MOUSE | HYQEEGQGTEVVQGVLLGLVVEDR | 5.32 | 4.38E-03 | 0.0150 | 2 |
| P62743 | AP2S1_MOUSE | QLLMLQSLE | 2.06 | 4.43E-03 | 0.0151 | 2 |
| Q9SZN1 | VATB2_ARATH V- | IPLFSAAGLPHNEIAAQIC[+57]R | 2.11 | 4.59E-03 | 0.0152 | 2 |
| G5E8R0 | G5E8R0_MOUSE | ETAEADVASLNR | 3.76 | 4.71E-03 | 0.0153 | 2 |
| Q61553 | FSCN1_MOUSE | YWTLTATGGVQSTASTK | 6.83 | 5.03E-03 | 0.0155 | 2 |
| P48774 | GSTM5_MOUSE | LTFVDFLTYDVLDQNR | 5.35 | 5.32E-03 | 0.0158 | 2 |
| P07356 | ANXA2_MOUSE | SALSGHLETVILGLLK | 5.91 | 5.33E-03 | 0.0158 | 2 |
| P35279 | RAB6A_MOUSE | LVFLGEQSVGK | 1.51 | 5.34E-03 | 0.0158 | 2 |
| Q7M6Y3 | PICAL_MOUSE | FIQYLASR | 1.55 | 6.08E-03 | 0.0165 | 2 |
| Q61553 | FSCN1_MOUSE | VGKDELFALEQSC[+57]AQVVLQAANER | 1.98 | 6.10E-03 | 0.0166 | 2 |
| Q9SZN1 | VATB2_ARATH V- | QIYPPINVLPSLSR | 1.55 | 6.17E-03 | 0.0166 | 2 |
| P08113 | ENPL_MOUSE I | GVVDSDDLPLNVSR | 1.76 | 6.38E-03 | 0.0169 | 2 |
| Q812A2 | SRGP3_MOUSE | VPGSQVEVNDIK | 4.02 | 6.39E-03 | 0.0169 | 2 |
| Q812A2 | SRGP3_MOUSE | TYLSAEYNLETSR | 4.31 | 6.54E-03 | 0.0171 | 2 |
| Q8R4E6 | PURG_MOUSE | DALVQLIEDYGEGDIEER | 5.31 | 6.72E-03 | 0.0172 | 2 |
| Q62188 | DPYL3_MOUSE | IAVGSDSDLVIWDPDALK | 1.97 | 6.99E-03 | 0.0175 | 2 |
| Q9CZ30 | OLA1_MOUSE ( | NYIVEDGDIIFFK | 2.34 | 7.53E-03 | 0.0180 | 2 |
| P14824 | ANXA6_MOUSE | DAFVAIVQSVK | 3.95 | 7.55E-03 | 0.0180 | 2 |
| O70435 | PSA3_MOUSE P | SNFGYNIPLK | 1.58 | 7.79E-03 | 0.0182 | 2 |
| P57722 | PCBP3_MOUSE | INISEGNC[+57]PER | 1.17 | 7.96E-03 | 0.0183 | 2 |
| Q9JJK2 | LANC2_MOUSE | IKDLLQQMEEGLK | 2.87 | 8.56E-03 | 0.0190 | 2 |
| P62852 | RS25_MOUSE 4 | LITPAVVSER | 2.30 | 9.26E-03 | 0.0198 | 2 |
| P61089 | UBE2N_MOUSE | TNEAQAIETAR | 2.25 | 9.51E-03 | 0.0202 | 2 |
| P08113 | ENPL_MOUSE I | FAFQAEVNR | 1.82 | 9.67E-03 | 0.0204 | 2 |
| Q9JJK2 | LANC2_MOUSE | SGNYPSSLSNETDR | 2.38 | 1.01E-02 | 0.0207 | 2 |
| P61089 | UBE2N_MOUSE | YFHVVIAGPQDSPFEGGTFK | 1.93 | 1.02E-02 | 0.0208 | 2 |
| Q9WUB3 | PYGM_MOUSE | VAIQLNDTHPSLAIPELMR | 2.13 | 1.03E-02 | 0.0209 | 2 |
| P46735 | MYO1B_MOUSE | LFSWLVNR | 3.21 | 1.03E-02 | 0.0209 | 2 |
| Q61701 | ELAV4_MOUSE | ILVDQVTGVSR | 2.74 | 1.06E-02 | 0.0213 | 2 |
| P57722 | PCBP3_MOUSE | LVVPASQC[+57]GSLIGK | 1.28 | 1.07E-02 | 0.0214 | 2 |
| Q8VDM4 | PSMD2_MOUSE | AVPLALALISVSNPR | 3.89 | 1.14E-02 | 0.0222 | 2 |
| P51150 | RAB7A_MOUSE | DPENFPFVVLGNK | 1.49 | 1.21E-02 | 0.0229 | 2 |
| Q9CZM2 | RL15_MOUSE 6 | VLNSYWVGEDSTYK | 3.01 | 1.26E-02 | 0.0233 | 2 |
| Q9WUB3 | PYGM_MOUSE | VHINPNSLFDVQVK | 2.32 | 1.31E-02 | 0.0239 | 2 |
| G5E8R0 | G5E8R0_MOUSE | SLQEQADAAEER | 5.35 | 1.31E-02 | 0.0239 | 2 |
| Q8R4E6 | PURG_MOUSE | FGENFIK | 1.13 | 1.32E-02 | 0.0240 | 2 |
| Q61701 | ELAV4_MOUSE | LDNLLNMAYGVK | 1.38 | 1.43E-02 | 0.0253 | 2 |
| P62830 | RL23_MOUSE 6 | ISLGLPVGAVINC[+57]ADNTGAK | 3.67 | 1.57E-02 | 0.0268 | 2 |
| Q61644 | PACN1_MOUSE | AYAQQLTDWAK | 1.42 | 1.61E-02 | 0.0272 | 2 |
| Q62188 | DPYL3_MOUSE | SAADLISQAR | 1.08 | 1.61E-02 | 0.0272 | 2 |
| P35980 | RL18_MOUSE 6 | TNRPPLSLSR | 3.57 | 1.68E-02 | 0.0279 | 2 |
| Q62261 | SPTB2_MOUSE | FLQDC[+57]QELSLWINEK | 2.54 | 1.73E-02 | 0.0284 | 2 |
| O54774 | AP3D1_MOUSE | NYLSLAPLFFK | 6.05 | 1.78E-02 | 0.0290 | 2 |
| P62702 | RS4X_MOUSE 4 | LSNIFVIGK | 3.86 | 1.81E-02 | 0.0294 | 2 |
| Q8C0E9 | Q8C0E9_MOUSE | M[+42]IGVNSVQSASK | 2.19 | 1.83E-02 | 0.0296 | 2 |
| P09405 | NUCL_MOUSE | QKVEGSEPTTPFNLFIGNLNPNK | 5.90 | 1.84E-02 | 0.0297 | 2 |
| Q8VEM8 | MPCP_MOUSE | IQTQPGYANTLR | 2.49 | 1.91E-02 | 0.0305 | 2 |
| P63318 | KPCG_MOUSE | LLNQEEGEYYNVPVADADNC[+57]SLLQK | 5.26 | 1.96E-02 | 0.0310 | 2 |
| Q99PT1 | GDIR1_MOUSE | VAVSADPNVPNVIVTR | 2.19 | 2.06E-02 | 0.0323 | 2 |
| P20357 | MTAP2_MOUSE | RLSNVSSSGSINLLESPQLATLAEDVTAALAK | 2.68 | 2.06E-02 | 0.0323 | 2 |
| Q8JZQ9 | EIF3B_MOUSE | VTLMQLPTR | 1.63 | 2.26E-02 | 0.0346 | 2 |
| Q8VDD5 | MYH9_MOUSE | TEM[+16]EDLM[+16]SSK | 1.57 | 2.28E-02 | 0.0348 | 2 |
| P05064 | ALDOA_MOUSE | IGEHTPSALAIMENANVLAR | 4.61 | 2.32E-02 | 0.0352 | 2 |
| Q8QZY1 | EIF3L_MOUSE I | YGDFFIR | 2.02 | 2.44E-02 | 0.0365 | 2 |
| F6ZIA4 | F6ZIA4_MOUSE | GAYDAQGTLSK | 4.66 | 2.45E-02 | 0.0366 | 2 |

| Protein Accession | Protein Description | Modified Peptide Sequence | co-IP:Ctrl | p value | q value | Sig Peptide Count |
|---|---|---|---|---|---|---|
| Q99JX4 | EIF3M_MOUSE | VAASC[+57]GAIQYIPTELDQVR | 4.86 | 2.46E-02 | 0.0367 | 2 |
| P56480 | ATPB_MOUSE | VALTGLTVAEYFR | 3.39 | 2.64E-02 | 0.0387 | 2 |
| P58771 | TPM1_MOUSE | C[+57]AELEEELK | 1.19 | 2.70E-02 | 0.0393 | 2 |
| P40124 | CAP1_MOUSE | VENQENVSNLVIDDTELK | 2.76 | 2.70E-02 | 0.0393 | 2 |
| O54983 | CRYM_MOUSE | SLGM[+16]AVEDLVAAK | 1.84 | 2.83E-02 | 0.0407 | 2 |
| P14115 | RL27A_MOUSE | TGVAPIIDVVR | 1.28 | 2.85E-02 | 0.0409 | 2 |
| P62827 | RAN_MOUSE | LVLVGDGGTGK | 5.05 | 2.99E-02 | 0.0425 | 2 |
| Q6ZWV3 | RL10_MOUSE | FNADEFEDMVAEK | 1.28 | 3.03E-02 | 0.0429 | 2 |
| P97427 | DPYL1_MOUSE | GVNSFQVYMAYK | 3.16 | 3.13E-02 | 0.0440 | 2 |
| P04370 | MBP_MOUSE | YLATASTM[+16]DHAR | 1.15 | 3.60E-02 | 0.0496 | 2 |
| O54774 | AP3D1_MOUSE | GVPVAEEVSALFAGELNPVAPK | 3.73 | 1.55E-03 | 0.0143 | 2 |
| Q8QZT1 | THIL_MOUSE | QEQDTYALSSYTR | 4.37 | 1.63E-03 | 0.0143 | 2 |
| Q99JX4 | EIF3M_MOUSE | QQWQQLYDTLNAWK | 4.83 | 2.02E-03 | 0.0143 | 2 |
| Q99K85 | SERC_MOUSE | ASLYNAVTTEDVEK | 1.38 | 2.34E-03 | 0.0143 | 2 |
| Q8K0T0 | RTN1_MOUSE | LFLVQDLVDSLK | 4.53 | 2.36E-03 | 0.0143 | 2 |
| Q8BVE3 | VATH_MOUSE | LLEVSDDPQVLAVAAHDVGEYVR | 1.76 | 3.02E-03 | 0.0143 | 2 |
| O70161 | PI51C_MOUSE | GAIQLGIGYTVGNLSSKPER | 3.06 | 3.21E-03 | 0.0143 | 2 |
| P63044 | VAMP2_MOUSE | LSELDDRADALQAGASQFETSAAK | 1.76 | 3.61E-03 | 0.0145 | 2 |
| Q99K85 | SERC_MOUSE | VIFVQGGGSGQFSAVPLNLIGLK | 2.78 | 3.65E-03 | 0.0146 | 2 |
| Q920I9 | WDR7_MOUSE | FRPPLLEMLAR | 4.51 | 4.75E-03 | 0.0154 | 2 |
| Q8BVE3 | VATH_MOUSE | YNIIPVLSDILQESVK | 2.44 | 4.85E-03 | 0.0154 | 2 |
| O70161 | PI51C_MOUSE | ITVQVEPVC[+57]GVGVVPK | 1.47 | 6.12E-03 | 0.0166 | 2 |
| P35700 | PRDX1_MOUSE | QITINDLPVGR | 1.22 | 6.34E-03 | 0.0169 | 2 |
| P35700 | PRDX1_MOUSE | LVQAFQFTDK | 2.35 | 7.12E-03 | 0.0176 | 2 |
| P63044 | VAMP2_MOUSE | LQQTQAQVDEVVDIMR | 2.06 | 8.59E-03 | 0.0190 | 2 |
| Q8QZT1 | THIL_MOUSE | IAAFADAAVDPIDFPLAPAYAVPK | 1.69 | 9.14E-03 | 0.0197 | 2 |
| Q8K0T0 | RTN1_MOUSE | HQAQVDQYLGLVR | 1.27 | 1.26E-02 | 0.0233 | 2 |
| Q80ZK2 | Q80ZK2_MOUSE | VVNAAIATASSAPGESEEPVVPSASR | 2.82 | 1.78E-02 | 0.0290 | 2 |
| P16546 | SPTN1_MOUSE | ALINADELANDVAGAEALLDR | 1.30 | 1.94E-02 | 0.0308 | 2 |
| B9EKR1 | PTPRZ_MOUSE | AIIDGTESVSR | 4.79 | 2.46E-02 | 0.0367 | 2 |
| P50516 | VATA_MOUSE | LPANHPLLTGQR | 1.37 | 2.50E-02 | 0.0371 | 2 |
| Q6URW6 | MYH14_MOUSE | LQQLFNHTM[+16]FVLEQEEYQR | 1.84 | 2.93E-02 | 0.0418 | 2 |
| O55131 | SEPT7_MOUSE | STLINSLFLTDLYSPEYPGPSHR | 2.80 | 1.13E-03 | 0.0143 | 2 |
| Q3UHJ0 | AAK1_MOUSE | VAEDEFDPIPVLITK | 3.78 | 1.62E-03 | 0.0143 | 2 |
| Q3UHJ0 | AAK1_MOUSE | LQTGFTENEVLQIFC[+57]DTC[+57]EAVAR | 6.29 | 1.95E-03 | 0.0143 | 2 |
| O55131 | SEPT7_MOUSE | NLEGYVGFANLPNQVYR | 1.82 | 8.71E-03 | 0.0191 | 2 |
| Q91V12 | BACH_MOUSE | ATLWYVPLSLK | 1.40 | 9.42E-03 | 0.0201 | 2 |
| Q921M7 | FA49B_MOUSE | GLLGALTSTPYSPTQHLER | 1.70 | 1.04E-02 | 0.0211 | 2 |
| Q11011 | PSA_MOUSE | ATFDISLVVPK | 1.26 | 1.24E-02 | 0.0231 | 2 |
| Q921M7 | FA49B_MOUSE | DQPPNSVEGLLNALR | 1.99 | 1.49E-02 | 0.0259 | 2 |
| Q11011 | PSA_MOUSE | YAAVTQFEATDAR | 1.39 | 1.65E-02 | 0.0276 | 2 |
| Q91V12 | BACH_MOUSE | VLEVPPIVYLR | 1.16 | 1.69E-02 | 0.0281 | 2 |
| Q7TMB8 | CYFP1_MOUSE | QLQVVPLFGDMQIELAR | 6.54 | 1.88E-02 | 0.0301 | 2 |
| Q9CR57 | RL14_MOUSE | LVAIVDVIDQNR | 5.79 | 3.32E-02 | 0.0463 | 2 |
| A0A0N5E063 | A0A0N5E063_TRII | EKLC[+57]FVALDFEQEM[+16]ATAASSSSLEK | 4.51 | 2.39E-03 | 0.0143 | 2 |
| Q9Z2T6 | KRT85_MOUSE | LC[+57]EGVGSVNVC[+57]VSSSR | 2.00 | 2.85E-03 | 0.0143 | 2 |
| Q7TMK9 | HNRPQ_MOUSE | EFNEDGALAVLQQFK | 4.28 | 2.90E-03 | 0.0143 | 2 |
| Q9CZX8 | RS19_MOUSE | RVLQALEGLK | 1.99 | 5.73E-03 | 0.0161 | 2 |
| Q9D6R2 | IDH3A_MOUSE | ENTEGEYSGIEHVIVDGVVQSIK | 1.65 | 7.83E-03 | 0.0182 | 2 |
| Q9Z2T6 | KRT85_MOUSE | SDLEANVEALVEESSFLK | 3.25 | 9.83E-03 | 0.0205 | 2 |
| Q9CZX8 | RS19_MOUSE | LKVPEWVDTVK | 1.64 | 1.45E-02 | 0.0255 | 2 |
| Q9D6R2 | IDH3A_MOUSE | TPIAAGHPSMNLLLR | 1.53 | 1.47E-02 | 0.0257 | 2 |
| B9EKR1 | PTPRZ_MOUSE | FAVLYQPLAGNDQAK | 3.75 | 1.79E-02 | 0.0292 | 2 |
| P14869 | RLA0_MOUSE | VLALSVETEYTFPLTEK | 4.61 | 2.17E-02 | 0.0336 | 2 |
| O08638 | MYH11_MOUSE | YEILAANAIPK | 4.43 | 2.23E-02 | 0.0343 | 2 |
| P01869 | IGH1M_MOUSE | DVLTITLTPK | 3.04 | 3.17E-02 | 0.0445 | 2 |
| Q7TQF7 | AMPH_MOUSE | LVDGSLLTLDTYLGQFPDIK | 2.88 | 4.34E-04 | 0.0143 | 2 |
| Q7TQF7 | AMPH_MOUSE | VETLHDFEAANSDELNLQR | 1.34 | 3.02E-03 | 0.0143 | 2 |
| Q61765 | K1H1_MOUSE | ETM[+16]QFLNDR | 1.88 | 3.99E-03 | 0.0148 | 2 |
| Q9R111 | GUAD_MOUSE | TPPLALVFR | 5.15 | 2.36E-02 | 0.0357 | 2 |
| P10649 | GSTM1_MOUSE | VTYVDFLAYDILDQYR | 3.65 | 6.81E-03 | 0.0172 | 2 |
| P10649 | GSTM1_MOUSE | LGLDFPNLPYLIDGSHK | 2.05 | 9.10E-03 | 0.0197 | 2 |

| Protein Accession | Protein Description | Modified Peptide Sequence | co-IP:Ctrl | p value | q value | Sig Peptide Count |
|---|---|---|---|---|---|---|
| Q922U2 | K2C5_MOUSE | TTAENEFVMLK | 2.50 | 7.68E-03 | 0.0181 | 2 |
| Q9DBZ5 | EIF3K_MOUSE | FIC[+57]HVVGITYQHIDR | 3.70 | 3.44E-02 | 0.0478 | 2 |
| Q9ERE2 | KRT81_MOUSE | LLEGEEQR | 3.55 | 5.60E-03 | 0.0160 | 2 |
| Q9ERE2 | KRT81_MOUSE | FAAFIDKVR | 2.49 | 5.67E-03 | 0.0160 | 2 |
| P51150 | RAB7A_MOUSE | EAINVEQAFQTIAR | 1.15 | 3.60E-02 | 0.0496 | 2 |
| Q3TLQ0 | Q3TLQ0_MOUSE | HSAGGGNVQIVTK | 4.60 | 1.15E-05 | 0.0143 | 1 |
| Q8C153 | Q8C153_MOUSE | ALLELQLEPEEIYQTFQR | 4.30 | 2.69E-04 | 0.0143 | 1 |
| P18872 | GNAO_MOUSE | IGAGDYQPTEQDILR | 2.00 | 3.05E-04 | 0.0143 | 1 |
| E9PVU0 | E9PVU0_MOUSE | SVTDYAQQNPAAQLPAR | 4.48 | 4.76E-04 | 0.0143 | 1 |
| Q5SVJ0 | Q5SVJ0_MOUSE | ISDILNSVR | 1.96 | 6.32E-04 | 0.0143 | 1 |
| P62874 | GBB1_MOUSE | LFVSGAC[+57]DASAK | 1.67 | 1.06E-03 | 0.0143 | 1 |
| Q99L75 | Q99L75_MOUSE | TSTVDLPIESQLLWQLDR | 2.58 | 1.29E-03 | 0.0143 | 1 |
| P28738 | KIF5C_MOUSE | ATEILNLLLK | 2.26 | 1.35E-03 | 0.0143 | 1 |
| Q8CG29 | Q8CG29_MOUSE | RPLPLTFSDLLQFR | 5.38 | 1.72E-03 | 0.0143 | 1 |
| M7NKU9 | M7NKU9_PNEMU | LIAQVVSSITASLR | 2.78 | 1.86E-03 | 0.0143 | 1 |
| P15508 | SPTB1_MOUSE | FLDLLEPLGR | 3.72 | 1.97E-03 | 0.0143 | 1 |
| Q3UJV2 | Q3UJV2_MOUSE | DVQDSVTVSNEIQTTK | 5.04 | 1.99E-03 | 0.0143 | 1 |
| Q3THH8 | Q3THH8_MOUSE | SKDAEVGDGTTSVTLLAAEFLK | 2.26 | 2.05E-03 | 0.0143 | 1 |
| Q9CWF2 | TBB2B_MOUSE | INVYYNEATGNK | 2.29 | 2.08E-03 | 0.0143 | 1 |
| Q3U804 | Q3U804_MOUSE | IRIIAPPER | 3.79 | 2.22E-03 | 0.0143 | 1 |
| P84104 | SRSF3_MOUSE | AFGYYGPLR | 3.81 | 2.23E-03 | 0.0143 | 1 |
| P68369 | TBA1A_MOUSE | LIGQIVSSITASLR | 2.96 | 2.33E-03 | 0.0143 | 1 |
| Q8BFZ3 | ACTBL_MOUSE | VAPDEHPILLTEAPLNPK | 3.90 | 2.90E-03 | 0.0143 | 1 |
| A0A0S2UN46 | A0A0S2UN46_9CF | YVGDEAQSKR | 3.84 | 2.92E-03 | 0.0143 | 1 |
| P17156 | HSP72_MOUSE | SINPDEAVAYGAAVQAAILIGDK | 5.29 | 3.45E-03 | 0.0144 | 1 |
| P61164 | ACTZ_MOUSE | DQLQTFSEEHPVLLTEAPLNPR | 2.88 | 3.52E-03 | 0.0144 | 1 |
| Q8C7C3 | Q8C7C3_MOUSE | EQAEAEVASLNR | 3.53 | 4.16E-03 | 0.0149 | 1 |
| Q56WH1 | TBA3_ARATH Tub | FDGAINVDITEFQTNLVPYPR | 4.07 | 4.20E-03 | 0.0149 | 1 |
| P61750 | ARF4_MOUSE | IQEGAAVLQK | 1.67 | 4.22E-03 | 0.0149 | 1 |
| Q61990 | PCBP2_MOUSE | QVTITGSAASISLAQYLINVR | 1.67 | 4.40E-03 | 0.0150 | 1 |
| A0A0N5DS75 | A0A0N5DS75_TRI | DLIGVQNLIK | 2.66 | 4.53E-03 | 0.0152 | 1 |
| M7P3Y2 | M7P3Y2_PNEMU | LPIFSAAGLPHNEIAAQIC[+57]R | 2.11 | 4.59E-03 | 0.0152 | 1 |
| Q9D1G1 | RAB1B_MOUSE | EFADSLGVPFLETSAK | 3.18 | 4.59E-03 | 0.0152 | 1 |
| P45591 | COF2_MOUSE | LGGSVVVSLEGKPL | 3.15 | 4.73E-03 | 0.0153 | 1 |
| A0A087WQ31 | A0A087WQ31_MC | VLSTSTDLEAAVADALLLGDAR | 7.22 | 5.09E-03 | 0.0155 | 1 |
| B2RXX6 | B2RXX6_MOUSE | LQTASDESYKDPTNIQLSK | 2.87 | 5.11E-03 | 0.0155 | 1 |
| A0A158UT74 | A0A158UT74_TRA | KFDEILEVSDGIMVAR | 2.08 | 5.68E-03 | 0.0161 | 1 |
| Q91Z67 | SRGP2_MOUSE | GASLLLYQR | 2.79 | 5.89E-03 | 0.0163 | 1 |
| A3QM89 | A3QM89_MOUSE | SQAIDLLYWR | 1.57 | 5.92E-03 | 0.0163 | 1 |
| M7NI55 | M7NI55_PNEMU C | VEILANDQGNR | 1.89 | 6.69E-03 | 0.0172 | 1 |
| P68372 | TBB4B_MOUSE | INVYYNEATGGK | 2.08 | 7.01E-03 | 0.0175 | 1 |
| Q3U561 | Q3U561_MOUSE | FSVC[+57]VLGDQQHC[+57]DEAK | 2.79 | 7.04E-03 | 0.0175 | 1 |
| A0A0N5DMQ | A0A0N5DMQ1_TR | DLYANTVLSGGSTM[+16]YPGIADR | 2.73 | 8.09E-03 | 0.0184 | 1 |
| D3Z2H9 | D3Z2H9_MOUSE | IQVLQQQADDAEER | 2.60 | 8.09E-03 | 0.0184 | 1 |
| F6VME3 | F6VME3_MOUSE | FSWGAEGQKPGFGYGGR | 3.31 | 8.56E-03 | 0.0190 | 1 |
| A0A0C5PUT3 | A0A0C5PUT3_PEI | VAEALATAAGHLDDLPGALSALSDLHAHK | 6.88 | 1.05E-02 | 0.0212 | 1 |
| Q3UX10 | TBAL3_MOUSE | SFGGGTGSGFTSLLMER | 4.14 | 1.19E-02 | 0.0227 | 1 |
| A0A0N5DXG6 | A0A0N5DXG6_TR | VPTPDVSVVDLTC[+57]R | 1.22 | 1.19E-02 | 0.0227 | 1 |
| O55143 | AT2A2_MOUSE | IGIFGQDEDVTSK | 2.37 | 1.21E-02 | 0.0229 | 1 |
| O70456 | 1433S_MOUSE | VLSSIEQK | 1.20 | 1.24E-02 | 0.0231 | 1 |
| P46097 | SYT2_MOUSE | IFVGSNATGTELR | 1.55 | 1.36E-02 | 0.0244 | 1 |
| B2RRE2 | B2RRE2_MOUSE | EPADLDPEAASPAYSQAK | 5.73 | 1.61E-02 | 0.0272 | 1 |
| Q641P0 | ARP3B_MOUSE | NVVLSGGSTM[+16]FR | 1.70 | 1.73E-02 | 0.0284 | 1 |
| P63011 | RAB3A_MOUSE | TYSWDNAQVLLVGNK | 1.48 | 1.89E-02 | 0.0302 | 1 |
| Q8R1B4 | EIF3C_MOUSE | FEELTNLIR | 8.27 | 2.14E-02 | 0.0333 | 1 |
| P51410 | RL9_MOUSE 60 | TILSNQTVDIPENVEITLK | 5.06 | 2.18E-02 | 0.0337 | 1 |
| Q61879 | MYH10_MOUSE | RHEMPPHIYAISESAYR | 1.20 | 2.24E-02 | 0.0344 | 1 |
| Q61765 | K1H1_MOUSE | LNVEVDAAPTVDLNR | 1.04 | 2.36E-02 | 0.0357 | 1 |
| Q9Z204 | HNRPC_MOUSE | VFIGNLNTLVVK | 2.52 | 2.44E-02 | 0.0365 | 1 |
| Q8CGP0 | H2B3B_MOUSE | AMGIMNSFVNDIFER | 3.71 | 2.56E-02 | 0.0377 | 1 |
| P11798 | KCC2A_MOUSE | GAILTTM[+16]LATR | 1.04 | 2.62E-02 | 0.0385 | 1 |
| Q9D8E6 | RL4_MOUSE 60 | YAIC[+57]SALAASALPALVMSK | 6.66 | 2.83E-02 | 0.0407 | 1 |

| Protein Accession | Protein Description | Modified Peptide Sequence | co-IP:Ctrl | p value | q value | Sig Peptide Count |
|---|---|---|---|---|---|---|
| Q9JMH9 | MY18A_MOUSE | GASFEELC[+57]HNYAQDR | 3.36 | 2.86E-02 | 0.0410 | 1 |
| Q8K310 | MATR3_MOUSE | ITPENLPQILLQLK | 5.52 | 3.03E-02 | 0.0429 | 1 |
| Q923T9 | KCC2G_MOUSE | ITEQLIEAINNGDFEAYTK | 1.40 | 3.34E-02 | 0.0466 | 1 |
| Q63844 | MK03_MOUSE | APEIMLNSK | 4.17 | 3.39E-02 | 0.0472 | 1 |
| Q80TL0 | PPM1E_MOUSE | ETDGTEGTVEIETVK | 2.73 | 5.31E-04 | 0.0143 | 1 |
| Q8BK64 | AHSA1_MOUSE | VFTTQELVQAFTHAPAALEADR | 2.68 | 5.49E-04 | 0.0143 | 1 |
| P61255 | RL26_MOUSE 6 | DDEVQVVR | 2.81 | 6.73E-04 | 0.0143 | 1 |
| Q9QWI6 | SRCN1_MOUSE | QIASLTGLVQSALLR | 5.63 | 7.16E-04 | 0.0143 | 1 |
| P28661 | SEPT4_MOUSE | STLVNSLFLTDLYR | 3.35 | 8.20E-04 | 0.0143 | 1 |
| Q6PER3 | MARE3_MOUSE | VLQAAFK | 2.62 | 1.05E-03 | 0.0143 | 1 |
| Q8K386 | RAB15_MOUSE | IQIWDTAGQER | 2.11 | 1.06E-03 | 0.0143 | 1 |
| P39054 | DYN2_MOUSE | VYSPHVLNLTLIDLPGITK | 2.67 | 1.14E-03 | 0.0143 | 1 |
| Q9Z1W9 | STK39_MOUSE | DAYELQEVIGSGATAVVQAALC[+57]KPR | 4.75 | 1.16E-03 | 0.0143 | 1 |
| Q9CPU0 | LGUL_MOUSE | GLAFIQDPDGYWIEILNPNK | 6.54 | 1.23E-03 | 0.0143 | 1 |
| S4R2F3 | S4R2F3_MOUSE | FLGPVIVEIPHFAALR | 3.12 | 1.49E-03 | 0.0143 | 1 |
| Q9JJV2 | PROF2_MOUSE | SQGGEPTYNVAVGR | 2.26 | 1.74E-03 | 0.0143 | 1 |
| Q8R081 | HNRPL_MOUSE | VFNVFC[+57]LYGNVEK | 3.01 | 1.77E-03 | 0.0143 | 1 |
| Q05512 | MARK2_MOUSE | VPVASPSAHNISSSSGAPDR | 3.83 | 2.02E-03 | 0.0143 | 1 |
| Q8BGQ7 | SYAC_MOUSE | NVGC[+57]LQEALQLATSFAQLR | 5.28 | 2.53E-03 | 0.0143 | 1 |
| Q920Q6 | MSI2H_MOUSE | GFGFVTFADPASVDK | 2.54 | 2.79E-03 | 0.0143 | 1 |
| F8VQC1 | F8VQC1_MOUSE | DIHTLAQLISAYSLVDPEK | 5.23 | 3.09E-03 | 0.0143 | 1 |
| P35922 | FMR1_MOUSE | SFLEFAEDVIQVPR | 2.99 | 3.93E-03 | 0.0147 | 1 |
| Q7TNM2 | TRI46_MOUSE | LLTELSFLR | 1.83 | 4.01E-03 | 0.0148 | 1 |
| P49312 | ROA1_MOUSE | LFIGGLSFETTDESLR | 3.22 | 4.82E-03 | 0.0154 | 1 |
| P62141 | PP1B_MOUSE | IVQMTEAEVR | 3.60 | 4.89E-03 | 0.0155 | 1 |
| O65719 | HSP7C_ARATH H | IINEPTAAAIAYGLDKK | 2.62 | 5.61E-03 | 0.0160 | 1 |
| O08807 | PRDX4_MOUSE | QITLNDLPVGR | 1.22 | 6.34E-03 | 0.0169 | 1 |
| Q9Z0R4 | ITSN1_MOUSE | TPIFLNEVLVK | 2.34 | 6.40E-03 | 0.0169 | 1 |
| P57776 | EF1D_MOUSE | GVVQDLQQAISK | 1.85 | 6.69E-03 | 0.0172 | 1 |
| P20444 | KPCA_MOUSE | LTDFNFLMVLGK | 6.33 | 7.08E-03 | 0.0175 | 1 |
| Q9Z2U0 | PSA7_MOUSE | ALLEVVQSGGK | 1.33 | 7.28E-03 | 0.0177 | 1 |
| P97384 | ANX11_MOUSE | GTITAASGFDPLR | 1.16 | 7.49E-03 | 0.0180 | 1 |
| Q0KK55 | KNDC1_MOUSE | NAGLLGQLEDFISSK | 7.15 | 7.58E-03 | 0.0180 | 1 |
| Q8CGY8 | OGT1_MOUSE | VPNSVLWLLR | 3.25 | 8.77E-03 | 0.0192 | 1 |
| Q9Z0H8 | CLIP2_MOUSE | AQELEGLDVEYR | 3.35 | 9.76E-03 | 0.0204 | 1 |
| Q8BYR5 | CAPS2_MOUSE | LFTESTGVLALEDKELGR | 1.85 | 1.04E-02 | 0.0211 | 1 |
| A0A0N5E1B9 | A0A0N5E1B9_TRI | AIDALHR | 4.42 | 1.05E-02 | 0.0212 | 1 |
| Q9D1A2 | CNDP2_MOUSE | LGGSVELVDIGK | 2.37 | 1.05E-02 | 0.0212 | 1 |
| Q8BGT8 | PHIPL_MOUSE | TEYTVAVQTASK | 1.83 | 1.08E-02 | 0.0215 | 1 |
| Q9DC51 | GNAI3_MOUSE | LLLLGAGESGK | 1.27 | 1.10E-02 | 0.0217 | 1 |
| Q6IFX2 | K1C42_MOUSE | ALEEANADLEVK | 1.31 | 1.15E-02 | 0.0223 | 1 |
| Q8BH66 | ATLA1_MOUSE | SM[+16]LQATAEANNLAAVATAK | 2.93 | 1.15E-02 | 0.0223 | 1 |
| Q64467 | G3PT_MOUSE | VPTPNVSVVDLTC[+57]R | 1.21 | 1.16E-02 | 0.0224 | 1 |
| O55100 | SNG1_MOUSE | DNPLNEGTDAAR | 1.29 | 1.39E-02 | 0.0248 | 1 |
| P48320 | DCE2_MOUSE | GAAALGIGTDSVILIK | 1.51 | 1.42E-02 | 0.0251 | 1 |
| Q91WC3 | ACSL6_MOUSE | LPELSDLGQFFR | 3.85 | 1.42E-02 | 0.0251 | 1 |
| A0A0N5DZK1 | A0A0N5DZK1_TRI | QDLPNAMNAAELTDK | 1.27 | 1.46E-02 | 0.0256 | 1 |
| P62849 | RS24_MOUSE | TTPDVIFVFGFR | 6.40 | 1.46E-02 | 0.0256 | 1 |
| O35098 | DPYL4_MOUSE | QIGENLIVPGGIK | 1.19 | 1.47E-02 | 0.0257 | 1 |
| P19157 | GSTP1_MOUSE | YVTLIYTNYENGK | 1.35 | 1.57E-02 | 0.0268 | 1 |
| Q8C2E7 | WASC5_MOUSE | LSEFIPAVFLLK | 5.74 | 1.66E-02 | 0.0277 | 1 |
| P62806 | H4_MOUSE His | ISGLIYEETR | 3.92 | 1.70E-02 | 0.0282 | 1 |
| P62259 | 1433E_MOUSE | AAFDDAIAELDTLSEESYK | 1.67 | 1.71E-02 | 0.0283 | 1 |
| A0A0N5E063 | A0A0N5E063_TRI | LC[+57]FVALDFEQEM[+16]ATAASSSSLEK | 1.60 | 1.79E-02 | 0.0292 | 1 |
| Q61361 | PGCB_MOUSE | ALGAHLTSIC[+57]TPEEQDFVNDR | 1.06 | 1.91E-02 | 0.0305 | 1 |
| H3BJD0 | H3BJD0_MOUSE | VFNTYSNEDYDR | 2.53 | 1.92E-02 | 0.0306 | 1 |
| Q9R1Q8 | TAGL3_MOUSE | AAEVYGVR | 1.40 | 1.93E-02 | 0.0307 | 1 |
| Q8VDD5 | MYH9_MOUSE | INFDVNGYIVGANIETYLLEK | 1.06 | 2.39E-02 | 0.0360 | 1 |
| O70194 | EIF3D_MOUSE | SVYSWDIVVQR | 3.70 | 2.39E-02 | 0.0360 | 1 |
| P05201 | AATC_MOUSE | NLDYVATSIHEAVTK | 1.97 | 2.50E-02 | 0.0371 | 1 |
| P19096 | FAS_MOUSE Fa | VLEALLPLK | 5.70 | 2.55E-02 | 0.0377 | 1 |
| Q64436 | ATP4A_MOUSE | NAADM[+16]ILLDDNFASIVTGVEQGR | 7.22 | 2.56E-02 | 0.0377 | 1 |

| Protein Accession | Protein Description | Modified Peptide Sequence | co-IP:Ctrl | p value | q value | Sig Peptide Count |
|---|---|---|---|---|---|---|
| P07724 | ALBU_MOUSE S | TC[+57]VADESAANC[+57]DK | 6.92 | 2.68E-02 | 0.0391 | 1 |
| Q9DCH4 | EIF3F_MOUSE | FLMSLVNQVPK | 8.50 | 2.80E-02 | 0.0404 | 1 |
| Q8BWY8 | Q8BWY8_MOUSE | SSLYLLM[+16]ETLNATTPHYVR | 4.94 | 2.83E-02 | 0.0407 | 1 |
| Q9JHU4 | DYHC1_MOUSE | VAEVLFDAADANAIEEVNLAYENVK | 3.37 | 2.85E-02 | 0.0409 | 1 |
| P62743 | AP2S1_MOUSE | VYTVVDEM[+16]FLAGEIR | 2.17 | 3.01E-02 | 0.0427 | 1 |
| A0A0N5DZT( | A0A0N5DZT0_TRI | LLLPGELSK | 3.32 | 3.40E-02 | 0.0473 | 1 |
| Q3UGR5 | HDHD2_MOUSE | LLLDGAPLIAIHK | 3.00 | 3.54E-02 | 0.0490 | 1 |
| P97807 | FUMH_MOUSE | ETAIELGYLTAEQFDEWVKPK | 5.04 | 7.01E-04 | 0.0143 | 1 |
| Q6ZPE2 | MTMR5_MOUSE | ISVQTPVDQLLQDGLQLR | 5.37 | 1.39E-03 | 0.0143 | 1 |
| P15532 | NDKA_MOUSE | EISLWFQPEELVEYK | 3.85 | 1.42E-03 | 0.0143 | 1 |
| Q8R1Q8 | DC1L1_MOUSE | AGATSEGVLANFFNSLLSK | 5.28 | 1.91E-03 | 0.0143 | 1 |
| P28740 | KIF2A_MOUSE | IDILTELR | 2.78 | 2.63E-03 | 0.0143 | 1 |
| P19246 | NFH_MOUSE N | KLLEGEEC[+57]R | 3.78 | 3.62E-03 | 0.0146 | 1 |
| P27773 | PDIA3_MOUSE | ELNDFISYLQR | 4.64 | 3.92E-03 | 0.0147 | 1 |
| Q80U49 | C170B_MOUSE | LGDASDTEAVDGER | 4.83 | 4.64E-03 | 0.0153 | 1 |
| O88569 | ROA2_MOUSE | LFIGGLSFETTEESLR | 2.78 | 5.07E-03 | 0.0155 | 1 |
| Q9QZD9 | EIF3I_MOUSE E | SYSSGGEDGYVR | 2.59 | 8.78E-03 | 0.0192 | 1 |
| O54781 | SRPK2_MOUSE | LKPWSLFDVLVEK | 2.93 | 9.12E-03 | 0.0197 | 1 |
| Q8BG05 | ROA3_MOUSE | LFIGGLSFETTDDSLR | 2.26 | 1.03E-02 | 0.0209 | 1 |
| P51174 | ACADL_MOUSE | AQDTAELFFEDVR | 1.93 | 1.04E-02 | 0.0211 | 1 |
| Q60865 | CAPR1_MOUSE | LNQDQLDAVSK | 3.15 | 1.07E-02 | 0.0214 | 1 |
| Q62277 | SYPH_MOUSE | LHQVYFDAPSC[+57]VK | 1.55 | 1.10E-02 | 0.0217 | 1 |
| Q9Z2Q6 | SEPT5_MOUSE | ESAPFAVIGSNTVVEAK | 1.61 | 1.53E-02 | 0.0264 | 1 |
| Q8BJH1 | ZC21A_MOUSE | FNENAADR | 3.12 | 2.32E-02 | 0.0352 | 1 |
| P63325 | RS10_MOUSE 4 | DYLHLPPEIVPATLR | 5.61 | 2.40E-02 | 0.0361 | 1 |
| P39053 | DYN1_MOUSE | KGWLTINNIGIMK | 4.53 | 2.62E-02 | 0.0385 | 1 |
| O08553 | DPYL2_MOUSE | IVLEDGTLHVTEGSGR | 3.32 | 2.85E-02 | 0.0409 | 1 |
| P08249 | MDHM_MOUSE | AGAGSATLSMAYAGAR | 1.99 | 3.03E-02 | 0.0429 | 1 |
| Q9R0P9 | UCHL1_MOUSE | MPFPVNHGASSEDSLLQDAAK | 1.41 | 5.63E-03 | 0.0160 | 1 |
| P42644 | 14333_ARATH 14- | DSTLIMQLLR | 1.78 | 7.91E-03 | 0.0183 | 1 |
| P11758 | HBB_MYOVE Hem | GTFASLSELHC[+57]DK | 1.75 | 1.46E-02 | 0.0256 | 1 |
| P63328 | PP2BA_MOUSE | QTLQSATVEAIEADEAIK | 4.63 | 1.57E-02 | 0.0268 | 1 |
| P17426 | AP2A1_MOUSE | LLQC[+57]YPPPEDAAVK | 1.76 | 2.06E-02 | 0.0323 | 1 |
| Q99KK2 | NEUA_MOUSE | VGLSAVPADAC[+57]SGAQK | 1.76 | 2.98E-02 | 0.0424 | 1 |
| P70168 | IMB1_MOUSE In | GALQYLVPILTQTLTK | 3.01 | 8.97E-04 | 0.0143 | 1 |
| P26369 | U2AF2_MOUSE | NFAFLEFR | 3.29 | 1.85E-03 | 0.0143 | 1 |
| Q9CZU6 | CISY_MOUSE C | ALGVLAQLIWSR | 3.95 | 2.90E-03 | 0.0143 | 1 |
| P17183 | ENOG_MOUSE | DATNVGDEGGFAPNILENSEALELVK | 2.89 | 3.18E-02 | 0.0446 | 1 |
| E9Q557 | DESP_MOUSE | LLQLQEQM[+16]R | 1.69 | 7.54E-04 | 0.0143 | 1 |
| P26443 | DHE3_MOUSE | VYNEAGVTFT | 1.77 | 1.11E-02 | 0.0219 | 1 |
