## Supplementary material for "MAP2 is Differentially Phosphorylated in Schizophrenia, Altering its Function": Table S5

**Table S5.** The potential protein interactors were entered into DAVID (https://david.ncifcrf.gov/home.jsp) and the functional annotation tool was used to highlight clustering using GOTERM categories. The top 8 annotation clusters (Column A) are provided with their enrichment score (Column B). The top 3 GOTERMS (column C), along with the Benjamini-corrected p-value (Column D), the number of co-IP proteins from our dataset that mapped to the given category (Column E), and the protein accession IDs of the proteins within each GOTERM (Column F) are also provided.

| Annotation cluster | Enrichment Score | GOTERM | Benjamini | # proteins | Protein Accession IDs |
|---|---|---|---|---|---|
| 1 | 28.23 | Intracellular ribonucleoprotein complex (GO:1990904) | $4.6x10^{-43}$ | 75 | P62270, P61979, O35737, P62751, P35980, P61358, P63017, Q9CPR4, P62754, P63325, P12970, Q7TMK9, Q61990, P35979, P62281, P47911, P14131, P62267, P62717, P62264, P29341, Q9D8E6, P62852, P32067, P67984, Q9CZM2, P14115, P97351, P49312, Q9D0E1, Q91VR5, P09405, P19253, P35922, Q8BP67, P51410, Q9CXY6, P58252, P27659, Q3U804, Q9WTM5, P14869, Q6ZWN5, P62301, Q9CR57, P84099, Q9WV60, Q8R081, Q61937, Q8BG05, P57722, Q99104, P62830, P62702, P62908, P62242, P63276, P62849, P60867, P47963, Q3U561, P26369, P62082, Q8C153, Q9Z2X1, P60335, Q9CZX8, O88569, P41105, D3Z4J3, Q9Z204, P60710, P62889, Q61656, P57780, P25444, Q8VEK3, P16858 |
| | | Translation (GO:0006412) | $1.4x10^{-30}$ | 72 | P62270, P60229, P62751, P35980, P61358, Q91WQ3, Q99L45, Q9CPR4, P62754, P12970, P23116, Q8R1B4, P35979, P62281, P47911, P14131, P62267, P62855, P62717, P62264, Q9D8N0, Q9D8E6, P67984, Q9CZM2, P14115, P97351, Q8BFR5, Q8QZY1, O70194, P19253, P10126, Q8BP67, P61255, P51410, P58252, P27659, Q8BP47, P62631, Q9DBZ5, Q8JZQ9, Q6ZWV3, Q99JX4, Q9Z1D1, Q9DCH4, Q6ZWN5, P62301, Q9CR57, Q9QZD9, Q91WK2, P84099, Q8VEM8, P62830, P62245, P62702, P62908, P62242, P63276, P62849, P60867, P57776, P47963, Q3U561, P62082, Q8C153, Q8BGQ7, P10630, Q9CZX8, Q8C0C7, P48962, P41105, P62889, P51881, P25444 |
| | | Ribosome (GO:0005840) | $3.2x10^{-27}$ | 47 | P62270, P27659, P62751, P35980, P61358, Q6ZWV3, Q9CPR4, P62754, P63325, P14869, P12970, Q6ZWN5, P62301, Q9CR57, P84099, P62281, P35979, P47911, P14131, P62267, P62264, P62717, P62855, P62830, P62245, P62702, P62908, P62242, Q9D8E6, P63276, P62852, P62849, P60867, P47963, Q3U561, P67984, P62082, Q9CZM2, P14115, P97351, Q9CZX8, P41105, P62889, P19253, Q8BP67, P25444, P51410 |

| Annotation cluster | Enrichment Score | GOTERM | Benjamini | # proteins | Protein Accession IDs |
|---|---|---|---|---|---|
| 2 | 26.22 | Cadherin binding involved in cell-cell adhesion (GO:0098641) | $6.8x10^{-30}$ | 60 | P62821, P07356, P42932, P60229, Q9CQV8, P61979, Q9Z2H5, Q9CPV4, P63017, Q68FG2, P12970, Q9D8Y0, Q8VDD5, P47911, Q62261, P62855, P62827, Q9D8N0, Q62167, P26039, P11499, P17182, P16546, E9PVU0, Q9CZM2, Q9QYC0, Q9CZ30, P62962, P47753, P47757, Q80ZK2, Q8BP67, P62259, Q9DCL9, P19096, B2RXX6, P58252, Q91YR1, P52480, P61027, P35700, Q9QXS6, Q9CR57, Q64331, O08709, O55131, P63101, Q9QXS1, P46638, Q61768, P46735, Q8C845, Q6A087, P57776, P20029, Q8BK64, Q9WUA3, O70456, P06151, Q8C153, Q923G3, P05064, P60335, Q3UJV2, V9GX76, Q7M6Y3, Q61553, P25444, Q05512 |
| | | Cell-cell adherens junction (GO:0005913) | $1.4x10^{-29}$ | 60 | P62821, P07356, P42932, P60229, Q9CQV8, P61979, Q9Z2H5, Q9CPV4, P63017, Q68FG2, P12970, Q9D8Y0, Q8VDD5, P47911, Q62261, P62855, P62827, Q9D8N0, Q62167, P26039, P11499, P17182, P16546, E9PVU0, Q9CZM2, Q9QYC0, Q9CZ30, P62962, P47753, P47757, Q80ZK2, Q8BP67, P62259, Q9DCL9, P19096, B2RXX6, P58252, Q91YR1, P52480, P61027, P35700, Q9QXS6, Q9CR57, Q64331, O08709, O55131, P63101, Q9QXS1, P46638, Q61768, P46735, Q8C845, Q6A087, P57776, P20029, Q8BK64, Q9WUA3, O70456, P06151, Q8C153, Q923G3, P05064, P60335, Q3UJV2, V9GX76, Q7M6Y3, Q61553, P25444, Q05512 |
| | | Cell-cell adhesion (GO:0098609) | $2.4x10^{-14}$ | 35 | Q91YR1, P60229, Q9CQV8, Q9Z2H5, Q9CPV4, P61027, Q68FG2, P35700, P12970, Q9D8Y0, Q9CR57, Q64331, O08709, P63101, Q9QXS1, P62855, P46735, Q9D8N0, Q8C845, P57776, Q6A087, Q8BK64, P26039, Q9WUA3, O70456, P17182, P06151, E9PVU0, Q9CZM2, Q923G3, Q9CZ30, Q9QYC0, P05064, Q3UJV2, V9GX76, P47753, Q61553, Q80ZK2, P47757, P25444, Q9DCL9, Q05512 |
| 3 | 10.59 | Eukaryotic translation initiation factor 3 complex (GO:0005852) | $1.0x10^{-14}$ | 13 | P60229, Q9DBZ5, Q8JZQ9, Q99JX4, Q9Z1D1, Q9DCH4, P23116, Q9QZD9, Q8R1B4, Q91WK2, Q8QZY1, O70194, Q62167 |
| | | Eukaryotic 48S preinitiation complex (GO:0033290 ) | $2.1x10^{-13}$ | 12 | P60229, O70194, Q9DBZ5, Q8JZQ9, Q99JX4, Q9Z1D1, P23116, Q9DCH4, Q8R1B4, Q9QZD9, Q8QZY1, Q91WK2 |
| | | Eukaryotic 43S preinitiation complex (GO:0016282) | $2.1x10^{-13}$ | 12 | P60229, O70194, Q9DBZ5, Q8JZQ9, Q99JX4, Q9Z1D1, P23116, Q9DCH4, Q8R1B4, Q9QZD9, Q8QZY1, Q91WK2 |

| Annotation cluster | Enrichment Score | GOTERM | Benjamini | # proteins | Protein Accession IDs |
|---|---|---|---|---|---|
| 4 | 7.22 | Vesicle-mediated transport (GO:0016192) | $2.2x10^{-7}$ | 27 | P62821, P28663, O54774, Q8CIE6, P61027, P61205, Q68FG2, P13020, Q9DBG3, Q68FD5, Q99104, P53994, Q6A087, P35279, P84091, Q9QZE5, P62743, P63044, O35643, P33175, D3Z4J3, Q61598, O08599, Q7M6Y3, P17427, Q80ZK2, P17426, P61294, P46460, P61750 |
| | | Protein transport (GO:0015031) | $8.4x10^{-6}$ | 44 | P62821, P28663, O54774, Q8CIE6, P51150, P61205, P61027, Q8K386, P61021, Q9QXY6, P63011, Q571F3, Q64331, Q8VDD5, Q9DBG3, P46638, Q80TJ1, Q8C2E7, Q9CZT8, Q99104, Q61548, Q9R1R2, P62827, P53994, Q8BYR5, P35279, P84091, P63044, P62743, Q9QZE5, Q9ERK4, E9PVU0, O35643, V9GX76, D3Z4J3, Q9EQH3, Q61598, O08599, P50396, P17427, Q9D1G1, P17426, P70168, P57780, P61294, P46460, P61750 |
| | | Intracellular protein transport (GO:0006886) | $1.3x10^{-4}$ | 23 | P62821, P28663, O54774, Q8CIE6, P51150, P35279, P61027, P84091, Q9QZE5, P62743, Q9ERK4, O35643, P49615, Q9EQH3, Q9DBG3, P17427, P68510, Q68FD5, P17426, P70168, P53994, P62827, P46460 |
| 5 | 7.16 | Unfolded protein binding (GO:0051082) | $5.0x10^{-10}$ | 19 | P42932, P20029, O54946, P63017, P11499, P38647, P08113, Q3THH8, P80314, P63037, P80315, P17156, P80313, P80318, P80317, P80316, P07901, Q9QYJ0, Q61937, P11983 |
| | | Positive regulation of protein localization to Cajal body (GO:1904871) | $4.4x10^{-8}$ | 8 | P42932, P80318, P80317, P80316, P11983, Q3THH8, P80314, P80315, P80313 |
| | | Chaperonin-containing T-complex (GO:0005832) | $6.7x10^{-9}$ | 8 | P42932, P80318, P80317, P80316, P11983, Q3THH8, P80314, P80315, P80313 |
| 6 | 5.76 | ATP hydrolysis coupled proton transport (GO:0015991) | $1.6x10^{-9}$ | 14 | Q9Z1W8, Q64436, P56480, P62814, Q6PIE5, Q9Z1G4, P50518, Q9Z1G3, Q03265, P51863, P50516, Q6PIC6, Q8VDN2, Q8BVE3 |
| | | Proton transport (GO:1902600) | $7.7x10^{-5}$ | 12 | Q64436, Q9Z1W8, Q03265, P51863, P56480, P50516, P62814, Q9Z1G4, P50518, Q9Z1G3, Q9DB20, Q8BVE3 |
| | | Proton-transporting ATPase activity, rotational mechanism (GO:0046961) | $1.6x10^{-4}$ | 8 | Q03265, P51863, P56480, P50516, Q9Z1G4, P50518, Q9Z1G3, Q8BVE3 |
| 7 | 5.37 | Membrane coat (GO:0030117) | $6.6x10^{-8}$ | 11 | O54774, Q8CIE6, Q9DBG3, P17427, Q68FD5, P62743, Q9QZE5, P17426, Q8CHC4, A0A0J9YUN4, P39053, O35643 |
| | | Intracellular protein transport (GO:0006886) | $1.3x10^{-4}$ | 23 | P62821, P28663, O54774, Q8CIE6, P51150, P35279, P61027, P84091, Q9QZE5, P62743, Q9ERK4, O35643, P49615, Q9EQH3, Q9DBG3, P17427, P68510, Q68FD5, P17426, P70168, P53994, P62827, P46460 |
| | | Protein transporter activity (GO:0008565) | $5.9x10^{-2}$ | 8 | O54774, Q9EQH3, Q9DBG3, P17427, P62743, P17426, P70168, O35643 |

| Annotation cluster | Enrichment Score | GOTERM | Benjamini | # proteins | Protein Accession IDs |
|---|---|---|---|---|---|
| 8 | 5.37 | Brush border (GO:0005903) | $6.3 \times 10^{-12}$ | 20 | Q64331, Q8VDD5, Q9QXS1, Q6URW6, E9Q634, P46735, Q9JMH9, Q60605, Q99JY9, Q8C0P5, E9PVU0, Q923G3, B2RRE2, E9QPE7, Q9JJ28, Q3UJV2, Q3THE2, V9GX76, P47753, P47754, Q5SV64, P47757, Q7TPR4, P57780, Q8K1M6, O08638, Q5SYD0, Q61879 |
| | | Myosin complex (GO:0016459) | $1.9 \times 10^{-8}$ | 14 | Q8CG29, Q8BWY8, E9PVU0, B2RRE2, E9QPE7, Q3THE2, P70248, Q64331, V9GX76, D3Z4J3, Q8VDD5, Q6URW6, Q5SV64, E9Q634, Q99104, P46735, O08638, Q5SYD0, Q60605, Q9JMH9, Q61879 |
| | | Motor activity (GO:0003774) | $7.0 \times 10^{-7}$ | 16 | Q8CG29, Q8R1Q8, Q8BWY8, E9PVU0, O08788, B2RRE2, E9QPE7, P70248, Q64331, V9GX76, D3Z4J3, Q8VDD5, Q6URW6, Q5SV64, Q9JHU4, E9Q634, Q99104, P46735, O08638, Q5SYD0, Q60605, Q9JMH9, Q61879 |
