## Supplementary material for "MAP2 is Differentially Phosphorylated in Schizophrenia, Altering its Function": Table S6

**Table S6. Primer sequences**

| TARGET | FORWARD (5'-3') | REVERSE (5'-3') |
|---|---|---|
| S1782E mutagenesis | CCACGCTGGATCTGCCTGGTTCCTGT GTAATGATCTCAGCC | N/A |
| MAP2c sanger sequencing | Primer 1: GCGGTAGGCGTGTACGGT<br>Primer 2: TTCACGCACACCAGGCACT<br>Primer3: TAGACCTAAGCCATGTGACAT | Primer 1: AGCAGTCCCCAAGTCAGT<br>Primer 2: AGTGCCTGGTGTGCGTGAA<br>Primer3: GAGAAGGAGGCAGATTAGCTG |
