## Supplementary material for "MAP2 is Differentially Phosphorylated in Schizophrenia, Altering its Function": Table S7

**Table S7. Map2 CRISPR Primer Design and Off Targets**

| Primer | Sequence |
|---|---|
| Map2 sgRNAF | **GAAATTAATACGACTCACTATAGG**ATGCCACGCTGGACCTGCTTGTTTTAGAGCTAGAAATAGC |
| sgRNA common | AAAAGCACCGACTCGGTGCCACTTTTTCAAGTTGATAACGGACTAGCCTTATTTTAACTTGCTATTTCTAGCTCTAAAAC |
| Cas9 Forward | **TATTACGACTCACTATAGG**GAGAATGGACTATAAGGACCACGAC |
| Cas9 Reverse | GCGAGCTCTAGGAATTCTTAC |
| Repair oligo | *T*G*TTTCTCTTCAACAGATTGACAGCCAAAAGTTGAACTTCAGAGAGCATGCAAAGGCCCGGGTAGATCACGGGGCTGAGATCATCACACAGGAGCCAAGCAGGTCCAGCGTGGCATCACCCCGACGA*C*T*C |
| MAP2 F1 | TGAACAGGTGGGAAAGAGCT |
| MAP2 R1 | TGTCCTTGTTGCGGATGGAA |

All sequences are written in the 5’ to 3’ direction.
Underlined sequences mark gRNA sites.
Sequences in **bold** are T7 promoter.
Glu knockin codon sequences are marked in red text in repair oligo.
Phosphorothioate linkages are marked with * in repair oligo.

**Table S7. Map2 CRISPR Primer Design and Off Targets**

| Offtarget Seq | Mismatch Count | Offtarget Score | Chrom | Locus Description |
|---|---|---|---|---|
| ATGCCACACAGGGCCTGCTTGGG | 3 | 0.559006211 | chr17 | intron:Kdm4b |
| ATGCCACACTAGACCAGCTATGG | 4 | 0.511363636 | chr18 | intergenic:Zeb1-Arhgap12 |
| GTGAGACACTGGACCTGCTTGGG | 4 | 0.505263158 | chr14 | intergenic:Lmo7-Gm22347 |
| AGGCCTTGCTGGACCTGCTTTGG | 3 | 0.4875 | chr15 | intergenic:Rims2-Dcstamp |
| CTGCCACTTTGGACCAGCTTTGG | 4 | 0.426136364 | chr19 | intergenic:Slc22a30-Gm6425 |
| CTGCCAAGCTAGACCTGCTAAGG | 4 | 0.361607143 | chr3 | intron:Zfhx4 |
| ATGCCACTGAGGACCTGCTTGGG | 3 | 0.331632653 | chr16 | intergenic:Mrps6-Kcne2 |
| CTGCCCTGCTGGACCTGCTTGGG | 3 | 0.316558442 | chr15 | intron:Cacna1i |
| TTGCCACGCTGGAACAACTTGGG | 4 | 0.296969697 | chr1 | intergenic:Wdr64-Exo1 |
| ATGCCACTCAGCACCTGCTTTGG | 3 | 0.283613445 | chr7 | intergenic:Glrx3-Gm25798 |
